# Fundamental limits to progression of cellular life in frigid environments

**DOI:** 10.1101/2022.06.10.495632

**Authors:** Diederik S. Laman Trip, Théo Maire, Hyun Youk

## Abstract

Life on Earth, including for microbes and cold-blooded animals, often occurs in frigid environments. At frigid temperatures, nearly all intracellular processes slow down which is colloquially said to decelerate life’s pace and, potentially, aging. But even for one cell, an outstanding conceptual challenge is rigorously explaining how the slowed-down intracellular processes collectively sustain a cell’s life and set its pace. Here, by monitoring individual yeast cells for months at near-freezing temperatures, we show how global gene-expression dynamics and Reactive Oxygen Species (ROS) act together as the primary factors that dictate and constrain the pace at which a budding yeast’s life can progresses in frigid environments. We discovered that yeast cells help each other in surviving and dividing at frigid temperatures. By investigating the underlying mechanism, involving glutathione secretion, we discovered that ROS is the primary determinant of yeast’s ability to survive and divide at near-freezing temperatures. Observing days-to-months-long cell-cycle progression in individual cells revealed that ROS inhibits S-G2-M (replicative) phase while elongating G1 (growth) phase up to a temperature-dependent threshold duration, beyond which yeast cannot divide and bursts as an unsustainably large cell. We discovered that an interplay between global gene-expression speed and ROS sets the threshold G1-duration by measuring rates of genome-wide transcription and protein synthesis at frigid temperatures and then incorporating them into a mathematical model. The same interplay yields unbeatable “speed limits” for cell cycling – shortest and longest allowed doubling times – at each temperature. These results establish quantitative principles for engineering cold-tolerant microbes and reveal how frigid temperatures can fundamentally constrain microbial life and cell cycle at the systems-level.

## INTRODUCTION

An important conceptual question in biology is how countless biochemical reactions together dictate a progression of an organism’s life. Although there is a familiar and loosely defined notion of life progressing at some rate, it has been difficult to rigorously define this rate even for a single cell and then quantify the effect of each intracellular process on this rate. Of practical interest is finding any unremovable constraints on slowing down a cell’s life, if they exist, because such constraints are thought be relevant for decelerating aging. Aside from being conceptually and biomedically relevant, finding such constraints and design principles that govern decelerated lives of cells is important for understanding a ubiquitous, natural phenomenon: it is crucial for understanding how organisms that cannot regulate their internal temperatures - including microbes, plants, and cold-blooded animals - survive in frigid environments that they frequently inhabit. One expects that the intracellular processes of these organisms would be extremely slow at frigid temperatures but exactly how slow is unknown for many of these processes^1^. Insightful studies have revealed specific genes, stress responses, and epigenetic mechanisms that contribute to sustaining a cell’s life in frigid environments^2–4^. But currently lacking is a systems-level understanding of how an interplay of myriad processes dictate and may constrain a cell’s progression in life at frigid temperatures which goes beyond a few specific mechanisms^1,5–10^. Here we sought to uncover systems-level design principles that dictate the pace of life for the budding yeast, *Saccharomyces cerevisiae*, at frigid temperatures by combining measurements at single-cell and genome-wide levels and mathematical modeling. As we will show, we found these design principles which revealed fundamental constraints that frigid temperatures can impose on microbial life and their cell cycle.

Conflicting observations have hindered our understanding of whether and how yeast survives in certain cold environments. Two popular views argue that the budding yeast cannot survive and/or proliferate at sufficiently low temperatures. One view proposes that essential processes such as transcription, translation, and molecular transport are too slow for sustaining cell proliferation at sufficiently low temperatures^1,8^. The other, complementary view proposes that yeast cannot proliferate at sufficiently low temperatures due to physical damages caused by their cell membranes becoming too rigid, proteins denaturing, oxidative stresses, and other events^1,5,9,10^. Yeast can repair such damages by, for example, expressing genes to fluidify their membranes, (re-)folding proteins with chaperones, and responding to oxidative stresses^1,8–10^. But at sufficiently low temperatures, the damage from various sources is thought to be too severe that yeast cannot survive and divide^5,9,11^. Importantly, how much each type of damage is responsible for causing yeast to die and fail to divide at frigid temperatures is unclear (i.e., exactly why a yeast fails to divide and survive at sufficiently low temperatures and what determines that a temperature is “sufficiently low”). Furthermore, the known repair mechanisms for yeast and other microbes are autonomous mechanisms: a cell is responsible for its own survival and growth without relying on other cells at frigid temperatures. Despite the two popular views mentioned above, daily experience shows that some baker’s yeast cells in an isogenic colony can evidently proliferate, albeit slowly, when stored in a refrigerator, and that fungal colonies can appear on refrigerated foods after months. Moreover, cellular processes can remain active despite being slow^6,8,11^. These daily-life observations, however, are often made in settings in which temperature is not well controlled. They do not reveal rigorous insights and any constraints on a cell’s life in frigid environments. Moreover, even laboratory studies that have examined cells at near-freezing temperatures have rarely applied quantitative methods to probe and continuously monitor the cell cycle of an individual cell of any organism for weeks to months, which may be necessary to fully observe extremely slow cell cycling that might occur for a small fraction of cells in a yeast population or other microbial populations. Taken together, the conflicting observations mentioned above have hindered our understanding of when and how yeast can survive in frigid environments. We reasoned that continuously monitoring yeast cells at near-freezing temperatures for long times (e.g., months) and applying quantitative methods to examine these cells – without a bias towards a particular damage or repair mechanism – may resolve the conflicts and reveal systems-level design principles that govern yeast’s life in cold environments.

### Outline of this paper

Our paper begins with a discovery of yeast cells helping each other in surviving, dividing, and avoiding extinction at frigid temperatures (Fig. 1). This discovery establishes that microbes can collectively combat damaging effects of frigid temperatures and thereby revises the textbook view in which a cell autonomously combats such effects to survive. We show that an onset of a sufficiently low temperature triggers this cooperative behavior in which yeast secretes and extracellularly accumulates glutathione (GSH), of which a threshold concentration is required for a yeast population to avoid becoming extinct at near-freezing temperatures (Fig. 2). By investigating how GSH, a well-known antioxidant, helps yeast cells survive and divide at frigid temperatures, we discovered the key reason that yeast dies and fails to divide in frigid environments: Reactive Oxygen Species (ROS) being largely created by cold nutrients that are common and essential to many organisms (e.g., amino acids, vitamins) (Fig. 2). Specifically, we discovered how the intracellular ROS level determines a yeast cell’s ability to grow, divide, and survive at frigid temperatures and how it impacts replicative and chronological lifespans in cold environments by continuously monitoring individual yeast cells for weeks to months and implementing quantitative, single-cell-level analyses (Fig. 3). We discovered that all these effects of ROS are due to one mechanism: ROS elongates the G1 (growth) phase of the eukaryotic cell cycle and thereby keeps a cell continuously increasing in size while inhibiting the cell from entering the S-G2-M (replicative) phase at frigid temperatures (Fig. 4). Crucially, we found that frigid temperatures impose a threshold concentration of ROS - and a corresponding threshold amount of time for staying in G1 - such that cells with more than this amount of ROS (and staying longer in G1 than the threshold duration) burst without ever dividing (Fig. 4). We show that an interplay of ROS and global gene-expression dynamics - days-long genome-wide transcription dynamics and weeks-long protein-synthesis dynamics - determines the threshold duration for exiting G1 at frigid temperatures (Fig. 5). Our paper ends with a mathematical model that summarizes all our findings into a coherent, quantitative picture which revealed unbeatable “speed limits” for cell cycling: shortest and longest allowed doubling times at each temperature, down to 0 °C (Fig. 6).

**Figure 1:**
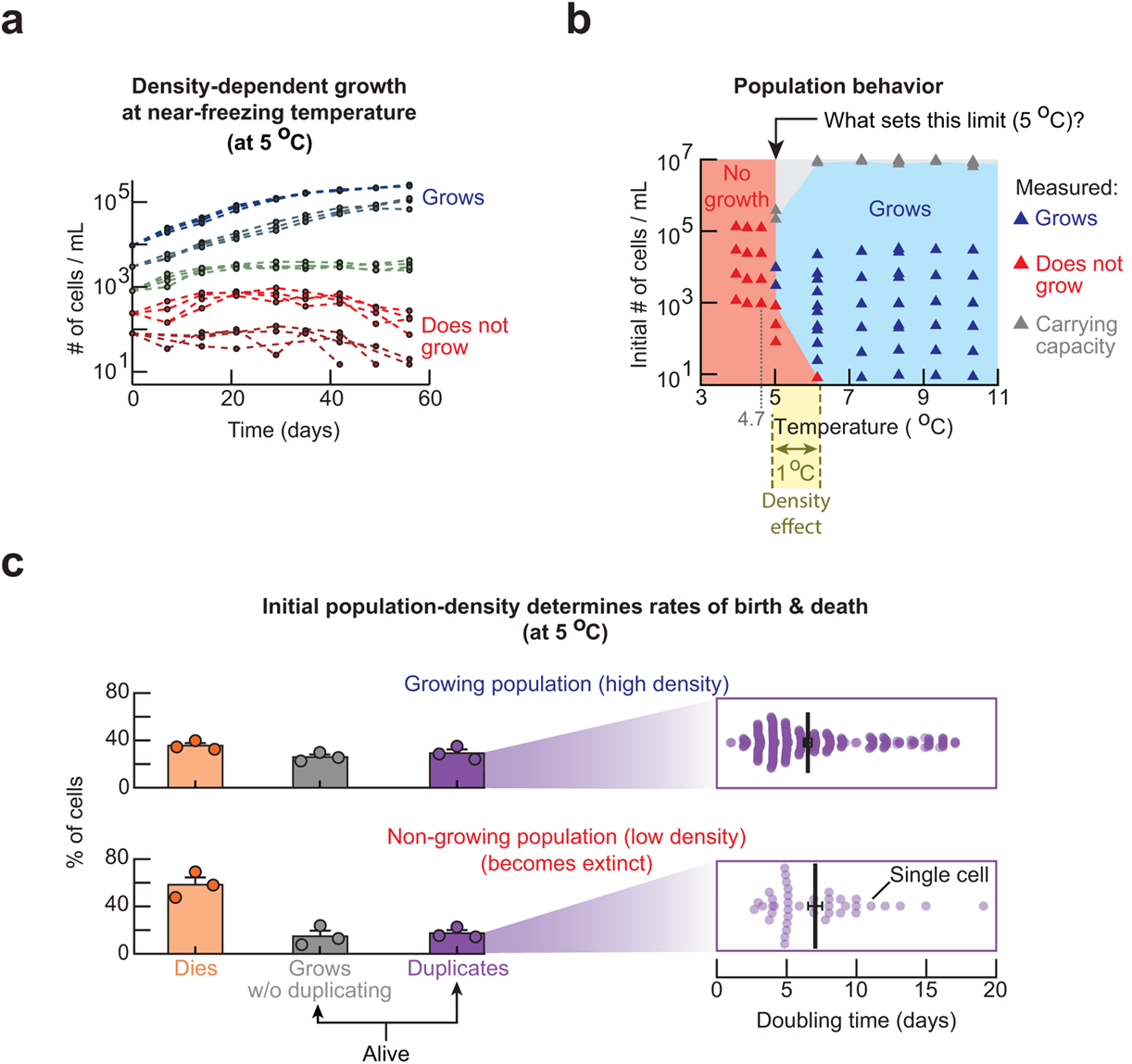
Yeast cells help each other in surviving and duplicating in frigid environments. **(a)** Population-density (number of cells / mL) of wild-type yeast measured over time with a flow cytometer at 5 °C. Each curve shows a different population (*n = 4* duplicate populations) (Also see Fig. S1). Blue: populations that grow to the carrying capacity. Red: populations that do not grow. **(b)** Phase diagram that summarizes all growth experiments of type shown in (a) (see data in Figs. S2–S5). Each triangle represents at least *n = 4* biological replicate populations that exhibit the same behavior as shown in (a). **(c)** Result from monitoring single cells of two populations for 20 days with a wide-field microscope. Upper half is for a growing, high-density population (initially ~6,250 cells / mL). Lower half is for a non-growing, low-density population (initially ~250 cells / mL). Bars represent mean of *n = 3* biological replicates (three circles per bar) with s.e.m. Zoomed out boxes on the right show doubling times of individual, duplicating cells (each dot represents one cell). Black bars denote average doubling time: 6.5 ± 0.3 days for the high-density population and 7.1 ± 0.5 days for the low-density population (error bars are s.e.m., for *n = 3* replicate populations). A doubling time for one cell is the time elapsed, from the moment when a bud appears on a mother cell to the moment when either the daughter (after being born) forms its own bud or when the mother forms another bud. Data also includes cells whose divisions were unfinished at the end of the time-lapse movie.

**Figure 2:**
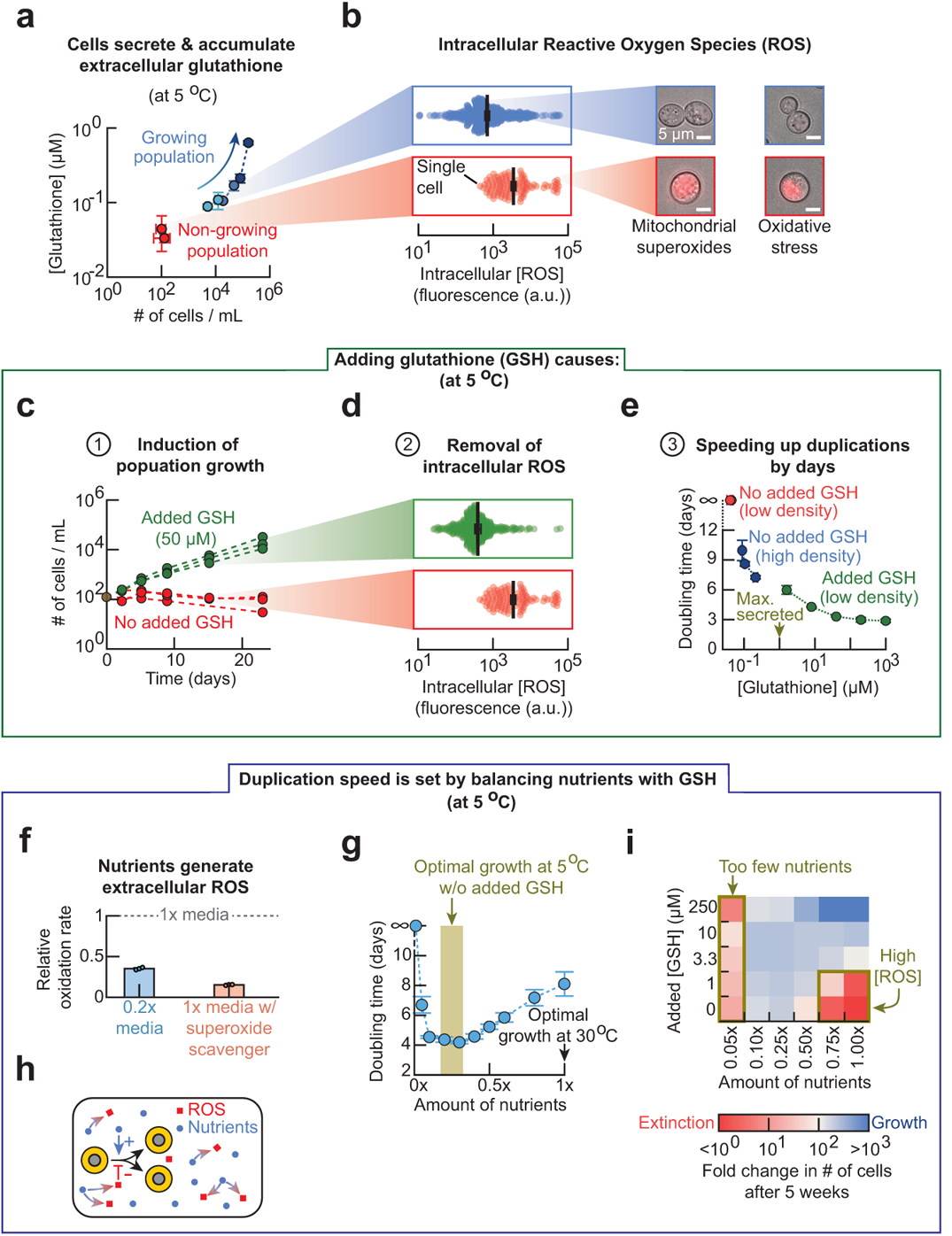
Glutathione (GSH) removes Reactive Oxygen Species (ROS) to enable cells to duplicate in frigid environments. **(a)** At 5 °C, concentration of extracellular glutathione for non-growing, low-density populations (red dots, all initially ~300 cells / mL) and growing, high-density populations (blue dots, initially > 900 cells / mL). Arrow indicates time passing during population growth. Each data point has *n = 3* biological replicates (also see Figs. S6-S8). **(b)** Intracellular ROS level proxied by a fluorescent dye for mitochondrial superoxides, measured after two weeks of incubation at 5 °C. Each dot is a single cell. Blue dots are for a high-density population (initially ~6,250 cells / mL). Red dots are for a non-growing, low-density population (initially ~250 cells / mL). Error bars show mean with s.e.m. for *n = 3* replicate populations. Representative pictures show intracellular ROS levels in cells. Left column: fluorescent dye for mitochondrial superoxides. Right column: fluorescent dye for general cellular ROS. Scale bar is 5 µm. **(c)** Populations incubated with (green curves) or without (red curves) 250 µM of reduced glutathione (“GSH”) (all initially ~120 cells / mL). **(d)** Intracellular ROS level in cells after two weeks of incubation, corresponding to populations shown in (c). Error bars show mean with s.e.m. with *n = 3* replicate populations (Also see Fig. S9). **(e)** Population’s doubling time plotted as a function of glutathione concentration measured in supernatant. Red: non-growing, low-density population (initially ~250 cells / mL) without any GSH supplemented. Blue: growing, high-density populations without any extra GSH supplemented. Green: low-density population (initially ~250 cells / mL) that grows after being given indicated amounts of GSH. *n = 4* for every point (Also see Figs. S10–S11). **(f)** Oxidation rate in a 0.2x medium (blue bar) or in 1x-medium that contained 100 µM of Trolox (superoxide scavenger) (purple bar). Rate is normalized to oxidation rate in 1x-medium. (Also see Fig. S12). **(g)** Population’s doubling time as function of the amount of non-sugar nutrients in the medium (initially ~190 cells / mL). Each data point has *n = 3* biological replicates. **(h)** Summarizing (f-g). **(i)** Each pixel in heatmap represents a fold-change in population density after five weeks of incubation with indicated amount of GSH and nutrients (all started with ~210 cells / mL). Each pixel is an average from *n = 3* replicate populations. (Also see Fig. S13).

**Figure 3:**
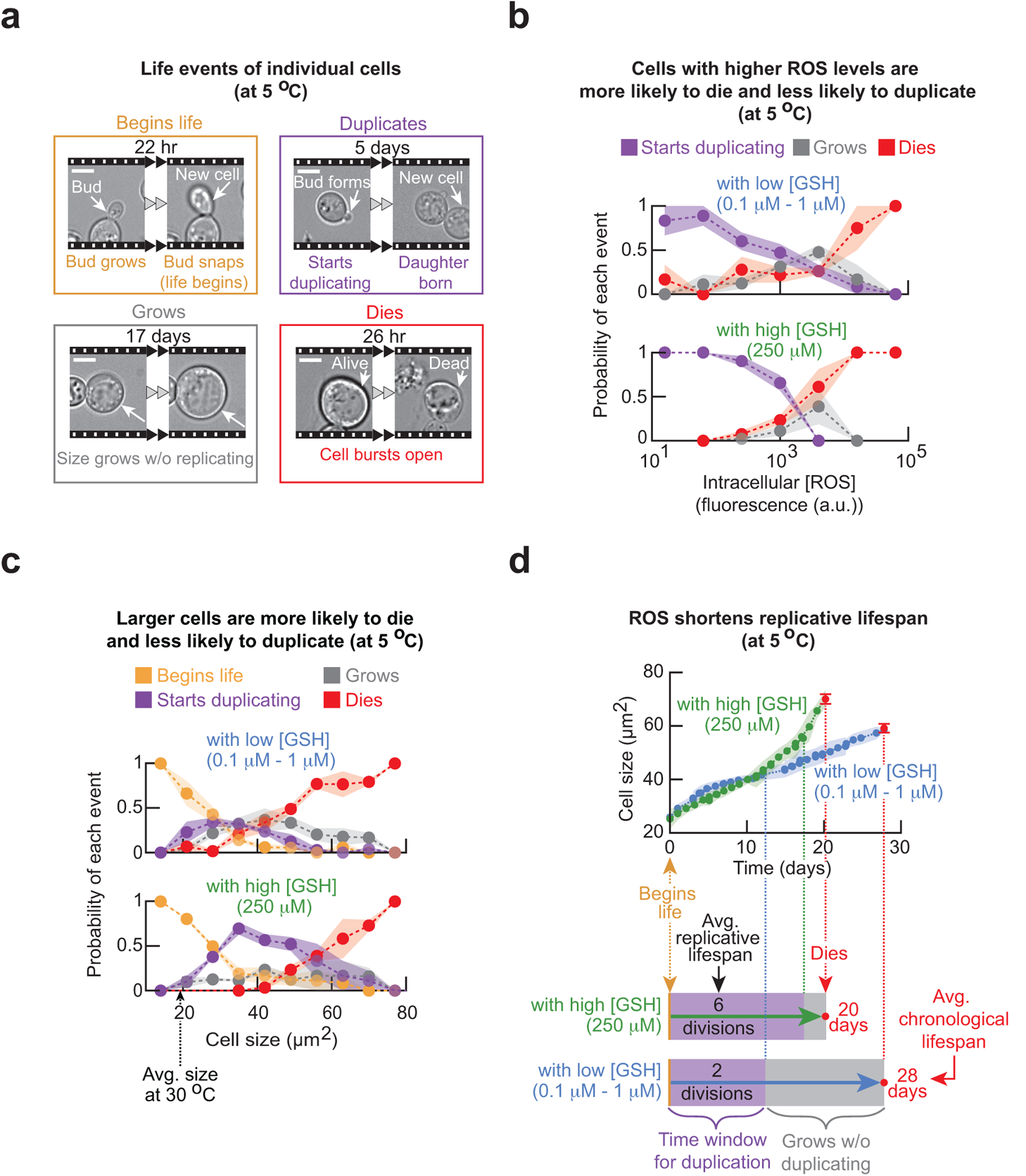
Single-cell analysis of cell-size growth, cell division, and cell death as a function of ROS and cell size. **(a)** Snapshots from time-lapse movies of single cells at 5.0 °C (initially ~8,000 cells / mL). Scale bar is 5 µm. To remove transient effects, populations were incubated for two weeks at 5.0 °C before the start of the movies on which we based (b) and (c). **(b)** Probability of each event (shown in (a)) occurring as a function of intracellular ROS (mitochondrial superoxide) level. Upper half: “with low [GSH]” population is a growing, high-density population that was not given any additional GSH (initially ~6,250 cells / mL). Lower half: “with high [GSH]” population is a growing, low-density population that was incubated with 250 µM of GSH (initially ~250 cells / mL). Dots show the mean, and shaded areas show s.e.m. from *n = 3* replicate populations. **(c)** Probability of each event (shown in (a)) occurring as a function of current cell size. Upper half: “with low [GSH]” population as defined in (b) (initially ~8,000 cells / mL). Lower half: “with high [GSH]” population as defined in (b) (initially ~420 cells / mL). Dots show the mean, and shaded areas show s.e.m. from *n = 3* replicate populations. (Also see Figs. S14–S17). **(d)** Curves show the size of an average cell over time, from birth to death, at 5.0 °C. Blue curve: cell of “with low [GSH]” population (averaged from *n = 330* cells). Green curve: cell of “with high [GSH]” population (averaged from *n = 175* cells). Red dot: death occurs. Shaded area and error bars represent s.e.m. of *n = 3* replicate populations. Details of how these curves and timelines were constructed are in Figs. S18–S19.

**Figure 4:**
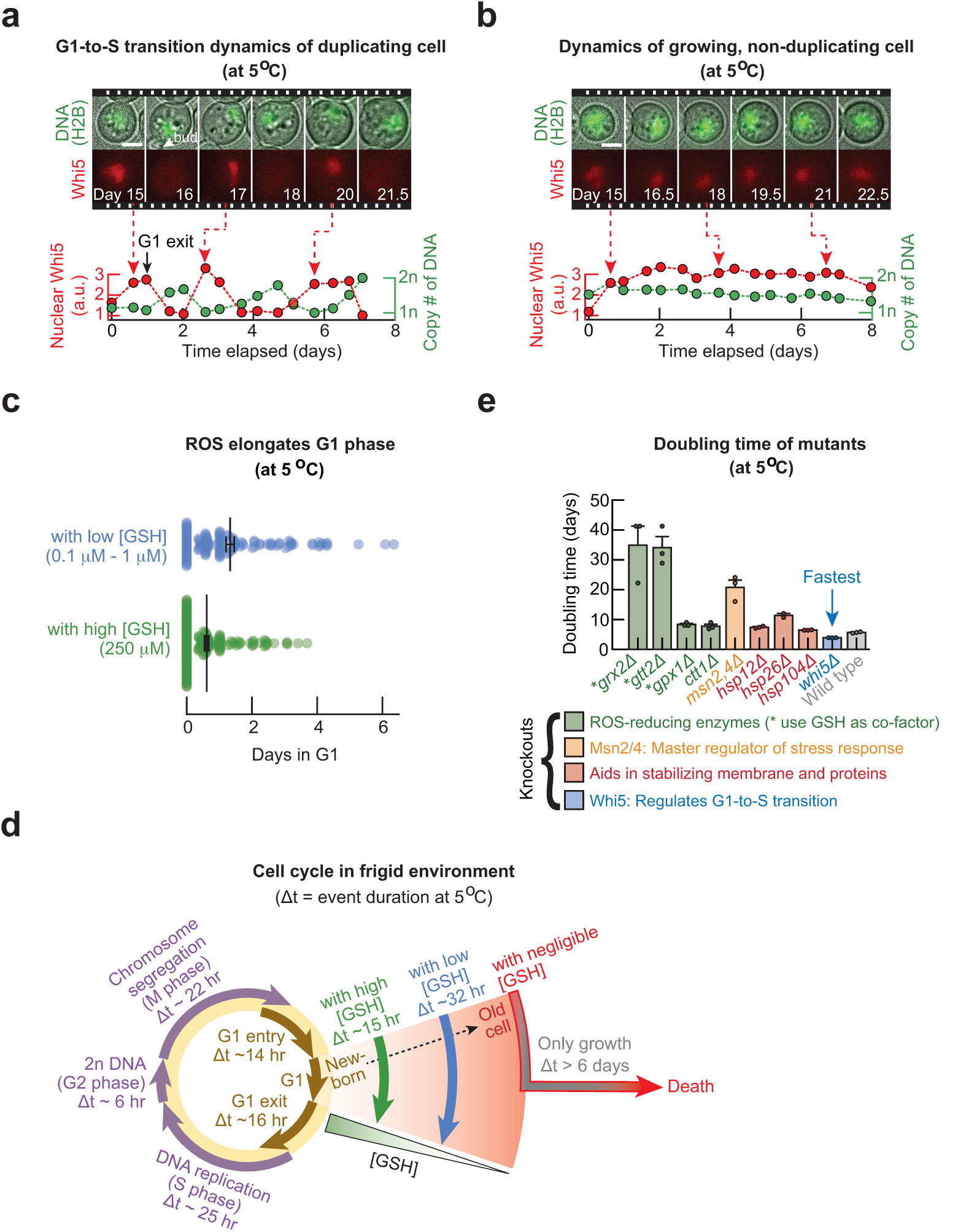
Ultraslow cell-cycle durations tuned with ROS and mutations in frigid environments. **(a-b)** Cells were incubated for two weeks at 5.0 °C before the start of the time-lapse movies to remove any transient effects. Duplicating cell in (a) and a non-duplicating cell in (b). Movie strips show composite of brightfield image with fluorescence in GFP (top row, H2B-GFP) or mCherry (bottom row, Whi5-mCherry). Graphs: normalized nuclear Whi5 fluorescence (red) and the copy number of DNA (green). Scale bars are 3 µm. (Also see Fig. S20). **(c)** Distribution of G1 duration among cells that divided. Each dot represents a cell. Blue: “with low [GSH]” population (initially ~6,250 cells / mL) is a high-density population that grew without any additional GSH (black: average of 32.0 ± 3.0 hours). Green: “with high [GSH]” population (initially ~1,250 cells / mL) is a low-density population that grew while incubated with 250 µM of GSH (black: average of 14.7 ± 1.5 hours). Error bars show mean with s.e.m., *n = 3* replicate populations (Also see Fig. S21) **(d)** Average duration of each cell-cycle phase at 5 °C determined by averaging durations from individual cells. G1 duration increases as cell chronologically ages and ROS level increases (GSH level decreases) with measured examples shown (green, blue, red arrows). Purple arrows show replicate phase: S-G2-M. (Also see Figs. S22–S23). **(e)** Population doubling time for each knockout strain (all initially ~6,250 cells / mL). Gene knocked out is listed and color coded in terms of its function as indicated. Error bars show the mean with s.e.m. of *n = 3* biological replicates. Dots show raw data. (Also see Figs. S24–S25).

**Figure 5:**
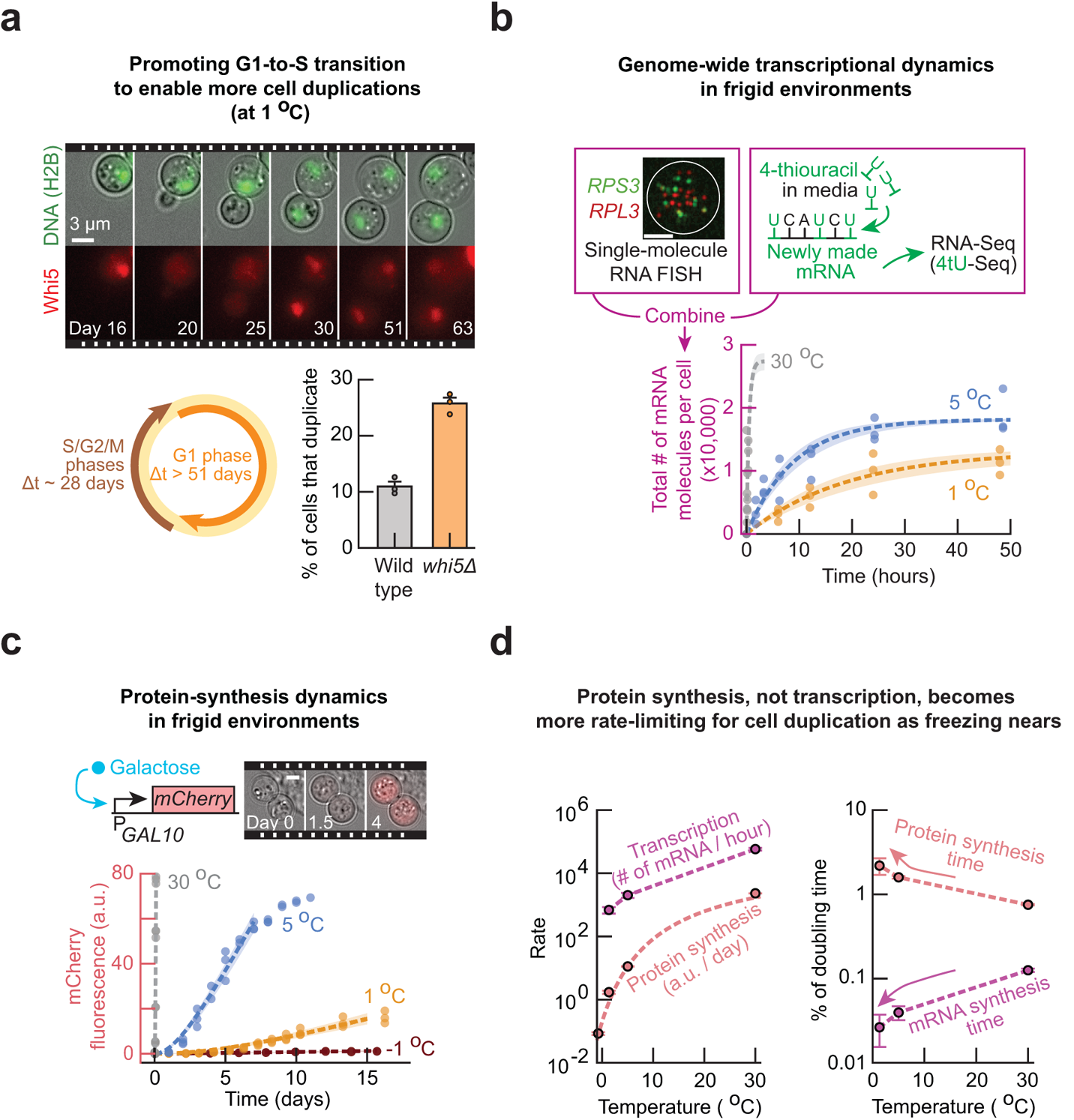
Genome-wide transcription rate and protein-synthesis rate at near-freezing temperatures. **(a)** Time-lapse movie of a duplicating cell at 1.0 °C. Composite of brightfield image and H2B-GFP (top) or Whi5-mCherry (bottom). Scale bar is 3 µm (also see Fig. S26). Circle shows average duration of cell-cycle phases at 1.0 °C: 28.4 ± 3.2 days for S-G2-M phase (n = 10 cells, brown arrow), and at least 51 days for G1 (equal to duration of the time-lapse, orange arrow). Bar graph shows percentage of cells that duplicated at 1.0 °C for *whi5Δ* strain (25.8 ± 1.0%, orange bar) and wild-type strain (10.9 ± 0.9%, grey bar). **(b)** Measuring genome-wide transcriptional dynamics by combining metabolically labelled RNA-sequencing with single-molecule RNA FISH (details in Methods). Graph: total number of newly synthesized mRNA over time at 30.0 °C (grey), 5.0 °C (blue) and 1.0 °C (orange). Image shows mRNAs in a single cell at 1.0 °C as visualized by single-molecule RNA FISH, with a composite of *RPS3* (green) and *RPL3* (red). White circle shows an outline of a cell. Scale bar is 2 µm. (Also see Figs. S27–S29). **(c)** Measuring protein-synthesis dynamics in individual cells. Movie strip: mCherry expression at 5.0 °C after 0, 1.5 and 4 days of incubation with 2% galactose. Images are composite of brightfield and mCherry fluorescence. Scale bar is 5 µm. (Also see Figs. S30–S31 and Supplementary text). In (b-c), dots show raw measurements, dotted lines show a model fitted (details in Supplementary text), and shaded areas show s.e.m. of fitted parameters. *n = 3* biological replicates. **(d)** Plot of rates extracted from (b) and (c): Left graph: transcription rate (# of mRNA / hour) and protein synthesis rate ([mCherry] a.u. / day). Red dotted line shows fit of Arrhenius-type equation to protein synthesis rate (Also see Fig. S32 and details in Supplementary text). Right graph: characteristic time for protein synthesis and transcription, both are relative to the average doubling time at each temperature. Characteristic time is the time to synthesize 1 a.u. mCherry or 100 mRNAs. In both graphs, error bars show mean with s.e.m. of *n = 3* biological replicates.

**Figure 6:**
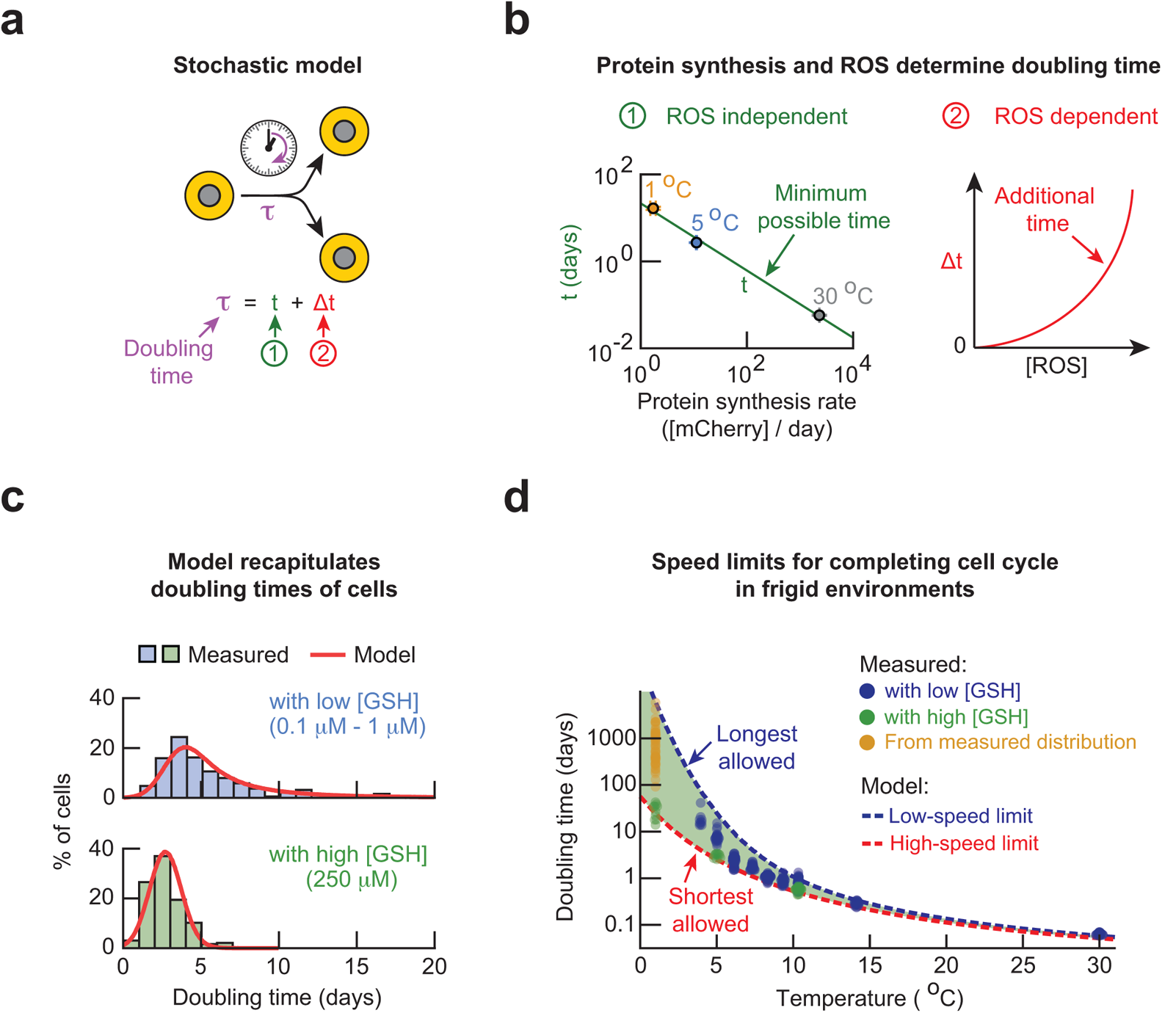
Mathematical model explains origin of low-speed and high-speed limits for completing cell cycle in frigid environments. **(a)** Description of the stochastic model (also see Supplementary text). A cell’s doubling time τ is dictated by a ROS-independent minimum duration *t* (labelled “1”) and a stochastic, ROS-dependent duration Δ*t* (labelled “2”). **(b)** Minimum possible doubling time *t* (taken from populations with 250 µM GSH at 5 °C or the average duration of S-G2-M at 1 °C) as a function of the protein-synthesis rate. Error bars show s.e.m. of *n = 3* biological replicates. Solid green line shows a power-law fit (exponent of 0.77 ± 0.05, Pearson correlation-coefficient β = 0.9979). The additional duration Δ*t* is determined by the intracellular ROS level and varies among cells (red line). **(c)** The model recapitulates the doubling times of cells at 5.0 °C with and without added GSH. Histograms are measured values (Fig. S11). Red curves are predictions of the model. **(d)** Model produces longest allowed doubling time (blue curve) and shorted allowed doubling time (red curve) for each temperature. The longest allowed doubling times are the doubling times of cells with the threshold concentration of ROS. Model predicts that a cell’s doubling time must fall within the green shaded region. Experimental data are from “with low [GSH]” populations as defined previously (blue dots), “with high [GSH]” populations (green dots, >5 °C), and 1 °C (green dots). Yellow dots are doubling times of individual cells that we extrapolated from measured distribution of cell-cycle times at 1 °C (Also see Fig. S33 and Supplementary text).

Through budding yeast, our work revealed how frigid environments can impose constraints on microbes and the eukaryotic cell-cycle. The design principles that govern yeast’s life at frigid temperatures and our systems-level approach that uncovered these principles may serve as a case study for future investigations that aim to find similar design principles for other microbes and microbial communities in frigid environments.

## RESULTS

### Cells help each other survive and duplicate in frigid environments

To examine yeast at frigid temperatures, we prepared liquid cultures of a laboratory-standard (“wild type”) yeast strain at various population densities, ranging from 10 cells/mL to 10^5^ cells/mL. We incubated these cultures at various fixed temperatures (4 °C to 14 °C) in a high-precision, thermostatic incubator that maintained a desired temperature within ± 0.1 °C (Fig. S1). During two months of incubation, we regularly took aliquots from each culture to measure its population density with a flow cytometer. At temperatures above 6 °C, population of every density slowly grew (Figs. S2–S3). But at 6 °C, surprisingly, none of the populations that started with 10 cells/mL grew whereas populations that started with at least 25 cells/mL grew to reach the carrying capacity during the two months. Similarly, at 5 °C, populations that started with at least 3,000 cells/mL grew (Fig. 1a - blue curves) whereas none of the populations that started with 1,000 cells/mL or less grew at all during the two months (Fig. 1a - red curves). Strikingly, at 4.7 °C – just 0.3 °C below 5 °C – no populations grew regardless of their initial density (Figs. S4–S5). In fact, at any temperature below 4.7 °C, we did not observe any populations growing regardless of their initial density. Combining these results yielded a “phase diagram” that indicated for which temperature and initial population-densities a population could grow (Fig. 1b). This diagram showed that the density-dependent population growth only occurs within a narrow, 1 °C-window between 5 °C and 6 °C. It also indicated that 5 °C is the lowest temperature at which yeast populations can grow. But, as we will show, the phase diagram hides the yeast’s true ability to duplicate at even lower temperatures (e.g., 1 °C).

To better understand the origin of the density-dependent population growth at 5 °C, we used a wide-field microscope to continuously monitor individual cells in “high-density” (6,250 cells/mL) and “low-density” (250 cells/mL) populations for three weeks at 5 °C. The high-density population grew towards the carrying capacity whereas the low-density population did not grow at 5 °C. For each cell, we determined whether it eventually died, stayed alive and divided, or stayed alive without duplicating (i.e., continuously increased in size) during the three weeks (Fig. 1c). We found that most cells in the high-density population survived whereas most cells in the low-density population died. Hence, the low-density population was headed towards extinction. Moreover, while both populations had only a minority of cells that divided, the low-density population had fewer dividing cells (17% duplicated) than the high-density population (29% duplicated). The duplicating cells had doubling times that ranged from 2 days to 17 days in both populations (Fig. 1c). Altogether, these results establish that the density-dependent population growth at 5 °C arises from cells helping each other to survive and duplicate.

### Cells collectively remove Reactive Oxygen Species (ROS) by secreting glutathione in frigid environments

We next asked how yeast cells were helping each other in surviving and dividing at low temperatures. At 5 °C, we found that a supernatant of the high-density population induced a growth of the low-density population that would have gone extinct without the supernatant (Fig. S6). Hence, yeast cells were likely secreting molecules that promoted their duplications. Motivated by our previous work^12^ which showed that yeast start to secrete and extracellularly accumulate glutathione, an antioxidant, as temperature increases above 37.5 °C, we hypothesized that yeast also secretes glutathione at sufficiently low temperatures. Indeed, at 5 °C, we detected glutathione gradually accumulating in the growth medium of high-density (growing) populations but not in low-density (non-growing) populations (Figs. 2a & S7). In fact, at every temperature below but not above 8 °C, we found that populations accumulated glutathione as they grew (Fig. S8).

Glutathione is yeast’s key antioxidant^13,14^ that inactivates (reduces) Reactive Oxygen Species (ROS). ROS can damage cellular components (e.g., nucleic acids, proteins and cell membranes^15–17)^. Hence, we used live-cell fluorescent reporter dyes to measure the ROS concentration inside each cell of the high-density (growing) population and the low-density (non-growing) population. We found that cells of the high-density population typically had much less intracellular ROS than cells of the low-density population (e.g., ~10-fold less superoxides) (Fig. 2b). These results establish that lower intracellular ROS concentrations are associated with a population growth.

### Adding reduced glutathione (GSH) enables and accelerates cell duplication by days

Since glutathione accumulates only for a growing population and lower levels of intracellular ROS are associated with growing populations, we hypothesized that ROS is the primary inhibitor of cell division for yeast at frigid temperatures instead of being a mere correlative measure. To test this idea, we added a high concentration (50 μM) of reduced glutathione (GSH) to the growth media of a low-density (non-growing) population at 5 °C. The added GSH caused the low-density population to grow (Fig. 2c - green) whereas the same, low-density population did not grow without the added GSH (Fig. 2c - red). Moreover, the added GSH decreased the ROS concentrations in cells of the low-density population (Fig. 2d - green) to nearly the same levels seen in cells of the high-density (growing) population (Fig. 2b - blue) (Fig. S9). These results establish that GSH alone can decrease ROS concentrations in cells of low-density (non-growing) populations and induce cell proliferation at 5 °C. Specifically, we determined that the low-density populations must be given at least ~1 μM of extracellular GSH to grow at 5 °C (Fig. S10).

We also discovered that increasing the extracellular GSH concentration accelerates population growth at 5 °C (Fig. 2e). For example, without adding any GSH, the low-density populations did not grow whereas sufficiently high-density populations doubled in density once every 7-10 days while accumulating up to ~0.1 μM of extracellular GSH (Fig. 2e - blue). Low-density populations, whose growth media were supplemented by GSH, could grow with a doubling time ranging from 6 days (with 1 μM of GSH) to 3 days (with 1 mM of GSH) (Fig. 2e - green). Thus, we could accelerate and tune the speed of population growth by varying the amount of extracellular GSH. We further confirmed this by using a microscope to monitor individual cells over weeks. We found that giving ample GSH to the low-density population more than tripled the percentage of cells that duplicated to 70% and shortened the average doubling time of a cell by more than half (Fig. S11).

Together, the above results revealed the cooperative mechanism by which yeast cells survive, duplicate, and avoid population extinctions at frigid temperatures: yeast cells collectively build an extracellular pool of GSH that they then use to reduce each other’s intracellular ROS. As we will see further, no populations grow at temperatures below 5 °C because yeast cells are not secreting enough GSH to rescue themselves from extinction. Indeed, adding ample GSH at 5 °C removes the density-dependent growth - populations of every initial density grows - because cells no longer rely on each other for GSH (Fig. 2e). Importantly, above results establish that ROS is the primary inhibitor of cell duplication for yeast at frigid temperatures.

### Cell-duplication speed is set by balancing amounts of GSH and ROS-generating nutrients

We next sought to identify a major source of ROS which we hypothesized to be some of the extracellular nutrients. We used the same fluorescent, oxidation-responsive dye as before to measure ROS levels in the growth medium (“1x medium”). The growth medium consisted of ample (2%) glucose and “non-sugar” nutrients that are common to yeast and other microbes (i.e., essential amino acids, vitamins, etc.). How rapidly the fluorescence of the dye changed over time (i.e., oxidation rate) in media without any yeast was a proxy for the ROS-creation rate by the media components, as demonstrated by the fact that adding scavengers of ROS (superoxides) to the 1x medium caused a 6-fold decrease in the oxidation rate at 5 °C (Fig. 2f - purple). We found that the non-sugar nutrients, but not glucose, were the primary generators of ROS in the media at 5 °C (Fig. S12). Decreasing the amount of non-sugar nutrients by 80% caused a ~85% decrease in the oxidation rate (Fig. 2f – blue; Fig. S12) and shortened the population’s doubling time from 8 days to 4 days at 5 °C (Fig. 2g). Decreasing the non-sugar level further, however, sharply increased the population doubling time (Fig. 2g). Hence the non-sugar nutrients have dual, opposing roles: they both promote and inhibit cell proliferation (Fig. 2h). Incubating yeast with varying amounts of GSH and non-sugar nutrient revealed the full extent to which we could inhibit or enable population growths at 5 °C (Figs. 2i & S13). Together, these results show that non-sugar nutrients are major generators of ROS in frigid temperatures.

### ROS does not affect cell-size growth but prevents cell division and promotes death

Having identified ROS as the primary inhibitor of population growth, we sought to elucidate how ROS affects an individual cell’s ability to divide, grow, and survive. To do so, we used a wide-field microscope to continuously monitor individual cells for three weeks at 5 °C. At the start of the 3-week period, we used a fluorescent ROS-reporter dye as in Fig. 2b to determine the ROS concentration in each cell. Subsequently, over the next 3 weeks, we determined which of the following four events pertained to each cell: 1.) the cell began as a bud at the start of our observation period and successfully separated from its mother to begin its life (Fig. 3a – “Begins life”); 2.) the cell duplicated (Fig. 3a – “Duplicates”); 3.) the cell’s size continuously increased without a bud ever appearing (Fig. 3a – “Grows”); 4.) the cell died (Fig. 3a – “Dies”) (Fig. 3a; Supplementary Movies M1-M2). We found that every cell continued to increase in their size regardless of which of the above events was occurring. Moreover, we observed that virtually all cells that died did so by bursting. By determining what percentage of cells underwent each of the four events for a given concentration of intracellular ROS, we found the probability of each event occurring as a function of intracellular ROS concentration. We first did so for cells of a “low [GSH] population”. This was a high-density population that grew at 5 °C by accumulating relatively “low” amounts of GSH (0.1 μM ~ 1 μM) (Fig. 2a – blue). In this population, cells with more ROS were less likely to divide and more likely to die (Fig. 3b – purple and red curves respectively in upper graph). Intriguingly, we found that nearly every cell had the same chance of increasing further in size regardless of their ROS level (Fig. 3b – grey curve in upper graph). Hence ROS does not affect cell-size growth but decreases the chance of dividing and increases the chance of dying.

Similarly, we determined the probability of each event occurring as a function of ROS level for a “high [GSH] population” (Fig. 3b – lower graph). This was a low-density population to which we gave an abundant (250 μM) GSH to induce its growth at 5 °C. Despite the high GSH concentration, we found that cells of the high [GSH] population spanned the same range of ROS levels as the cells of the low [GSH] population (i.e., 10 ~ 10^5^ fluorescence units in Fig. 3b). This suggested that, even with the high amount of GSH that we gave to this population, cells cannot reduce ROS when ROS is sufficiently abundant. Indeed, in the high [GSH] population, cells that had more than a certain, “threshold concentration” of ROS had a virtually zero chance of dividing (Fig. 3b - bottom purple curve ends at ~6,000 a.u. of ROS). Cells with an above-threshold concentration of ROS could still increase in size or, more likely, die. As in the low [GSH] population, we found that cells with more ROS were more likely to die and less likely to divide. However, while having a less-than threshold concentration of ROS did not guarantee that the cell would divide in the low [GSH] population, it virtually guaranteed that the cell would divide in the high [GSH] population (Fig. 3b - bottom purple curve below ~6,000 a.u. of ROS).

### Larger cells are more likely to die and less likely to divide regardless of their ROS level

In support of our finding that ROS does not affect a cell’s ability to increase in size, we found no correlation between the amount of ROS in a cell and its size (maximum cross-sectional area) (Fig. S14). Hence, we reasoned that cell size, like ROS, may serve as a predictor of a cell’s future^18–22^. To test this idea, we used a wide-field microscope to continuously monitor how each cell’s size changed over three weeks at 5 °C. We could then determine how likely a cell of a certain size engaged in each of the four events mentioned above (Fig. 3c). We examined both the low [GSH] and high [GSH] populations. In both populations, we found that larger cells were more likely to die and less likely to divide (i.e., less likely to form a bud to start cell division) (Fig. 3c – red and purple curves respectively). By comparing the two populations, we discovered that increasing the GSH level - and thereby decreasing the average ROS level in a cell - greatly increased the chance of dividing for cells of nearly all sizes (Fig. 3c – compare two purple curves). In fact, reducing the average ROS level caused even some of the largest cells to divide. Moreover, in both populations, cells of all sizes had nearly the same chance of increasing further in size without dividing (Fig. 3c – grey curves). Thus, neither ROS nor cell size affects a cell’s ability to increase further in size.

### ROS shortens replicative lifespan but increases chronological lifespan

Our continuous monitoring of individual cells established that cell size continuously increases in time, regardless of a cell’s ROS level and current size, until the cell bursts (Figs. S15–S17). It also established the typical (average) sizes of newborn and dying cells. Thus, we could plot the average cell size as a function of time, starting with the typical size at birth and ending with the typical size at death for both the low [GSH] and high [GSH] populations (Fig. 3d & Fig. S18). Additionally, using the probability of a cell dividing as a function its size, we could determine how many times an average cell divides in its lifetime (i.e., replicative lifespan) and how old it was when each division occurred. This analysis revealed that increasing the GSH level – and thereby decreasing the average ROS level in a cell – caused an increase in the replicative lifespan (Fig. 3d). This makes sense given our finding that ROS inhibits cell divisions. Moreover, consistent with our finding that GSH promotes cell division for cells of every size, the above analysis revealed that the typical cell of the high [GSH] population continued to divide until the last moments of its life as a large cell (Fig. 3d – green curve). In contrast, the typical cell of the low [GSH] population could only divide while it is small, during the first half of its 28-day life (Fig. 3d – blue curve). Finally, the same analysis revealed that decreasing the ROS level causes the cell to increase its size more rapidly, meaning that a cell with less ROS typically reached a burst-prone (large) size faster - and thus have a shorter lifespan - than a cell with more ROS (Fig. 3d – timelines, also see Fig. S19).

### ROS elongates G1 (growth) duration and inhibits G1-to-S transition

We have now established that ROS inhibits cell divisions while allowing for the cell size to continuously increase. Cell division pertains to the S-G2-M (replicative) phase of the cell cycle and cell-size growth pertains to the G1 (growth) phase of the cell cycle^23–25^. Hence, examining how ROS affects the durations of each cell-cycle phase may provide a mechanistic understanding of how ROS prevents cell divisions. To that end, we engineered a yeast strain by fusing GFP to histone, H2B, and mCherry to a regulator of G1-to-S transition, Whi5. In this strain, a cell’s GFP level was a proxy for its DNA abundance. The mCherry fluorescence migrating from nucleus to cytoplasm marked a cell exiting G1 to enter S phase^20,26,27^ (Fig. S20). At 5 °C, for one month, we used an epifluorescence microscope to monitor individual cells of the low [GSH] and high [GSH] populations. In both populations, cells that duplicated typically took one day to continuously replicate their DNA (i.e., GFP level continuously increased) after Whi5 exited their nuclei (Fig. 4a; Supplementary Movie M3). In both populations, cells that did not duplicate continuously increased in their size while their DNA level remained constant and their Whi5 remained in their nuclei (Fig. 4b; Supplementary Movie M4). Hence, for both populations, all non-duplicating cells were stuck in G1.

By monitoring many single cells for a month, we determined the average duration of every major cell-cycle event and how ROS affected each duration at 5 °C (Figs. 4c-d). Doing so revealed that increasing the GSH level – and thereby decreasing the average ROS level in a cell – caused more cells to exit G1 (Figs. S21–S22) and shortened the G1 duration of duplicating cells, from an average of 32 hours to 15 hours (Fig. 4c & Fig. 4d – green and blue arrows on right side). Thus, at 5 °C, cells with more ROS spent more time in G1 (growth) phase and were less likely to divide (i.e., less likely to enter S phase). This explains why a cell with an above-threshold concentration of ROS is virtually guaranteed to die by bursting: such a cell cannot leave G1 and thus it increases in size until growing further is physically impossible. We found that nearly every cell that spent more than 6 days in G1 failed to divide and died by bursting (Fig. 4d – red and grey arrows). This suggests - and as we will confirm below - a cell must exit G1 within a certain time window to complete the cell cycle and that this window is set by the threshold concentration of ROS.

We also found that ROS increases the G1 duration in such a way that a cell typically completed its current cell cycle more slowly than its previous cell cycle (Fig. S23). Specifically, a newborn cell typically spent less than 12 hours in G1 (Fig. 4d – green arrow) whereas older cells could stay in G1 for more than 12 days until they burst (Fig. 4d – red arrow).

### ROS does not affect G2-S-M (replicative) duration

Reconstructing the cell cycle also revealed that ROS does not affect the durations of G2, S, and M phases. Indeed, for both populations that differed in their ROS levels, a duplicating cell at 5 °C typically took ~25 hours to replicate its chromosomes, ~6 hours for G2 phase, and ~22 hours for mitosis and cytokinesis combined (Fig. 4d - purple arrows; Fig. S22). Hence, a duplicating cell typically took a total of ~2 days for the combined S-G2-M (replicative) phase regardless of its ROS level at 5 °C (Fig. 4d - left half of the circle). This result also shows that the wide variation among cells in their doubling times, which we previously noted (Fig. 1c), is due to the variation in their G1 duration but not in their S-G2-M duration.

### Mutations hinder or promote cell duplications by controlling G1-to-S transition in frigid environments

We have now established that ROS inhibits the G1-to-S transition and thereby prevents cells from dividing while letting their size continue to increase until the cell bursts. Since ROS decreases a cell’s chance of exiting G1, we reasoned that eliminating the genes for ROS-reducing enzymes that use GSH as a co-factor, *GRX2* and *GTT2*^8,9^, would either elongate the G1 or cause more cells to die. Both effects would manifest as an increase in a population’s doubling time. Indeed, we found that knocking out either one of the two enzymes (*grx2Δ* and *gtt2Δ*) greatly increased the doubling time to 35 days for a high-density population whose doubling time is 6 days when both genes are present (Fig.4e – compare left two green bars with “wild type” bar). Importantly, by constructing several other gene-knockout strains such as knockouts of Msn2 and Msn4^28^, we found that the most severe increases in a population’s doubling time was caused by knocking out the two ROS-reducing enzymes (*grx2Δ* and *gtt2Δ*) that required GSH as a co-factor (Figs. 4e & S24). Hence, disrupting a cell’s ROS-reducing ability severely inhibits its ability to duplicate at 5 °C.

Another way to promote cell duplications may be enabling cells with abundant ROS to exit G1. Indeed, we found that knocking out Whi5, an important regulator of G1-to-S transition, caused more cells to duplicate without changing their ROS levels at 5 °C (Fig. S25). In fact, the only mutant that we found to have a shorter average doubling time than the wild-type cells at 5 °C was a knockout of Whi5 (*whi5Δ*) (Fig. 4e). These results also held at 1 °C: by using a microscope to continuously monitor individual *whi5Δ* cells for two months at 1 °C, we found that knocking out Whi5 increased a cell’s chance of duplicating by 2.5 times. Specifically, 26% of *whi5Δ* cells could duplicate whereas 11% of wild-type cells of the same population density could duplicate at 1 °C (Fig. 5a; Supplementary Movie M5). At 1 °C, the duplicating cells took ~28 days to complete their S-G2-M phases and G1 duration was typically longer than 51 days (Figs. 5a & S26).

### Existence of speed limits for completing eukaryotic cell cycle in frigid environments

Taken together, our findings thus far establish that a cell cannot take an arbitrarily long time to complete the cell cycle at a near-freezing temperature. This is because we have shown that a longer doubling time arises from a longer G1 duration which, in turn, arises from having more ROS. But once the ROS level increases to an above-threshold concentration, a cell has virtually no chance of completing the cell cycle and is guaranteed to burst without dividing. According to this reasoning, there must be a longest allowed doubling time for each temperature. There must also be a shortest allowed doubling time for each temperature because the S-G2-M phase has a fixed duration that is independent of ROS. In the remainder of our paper, we focus on combining all our findings thus far to determine the possible pace at which a yeast cell can cycle and the “speed limits” for this pace – the longest and shortest allowed doubling times – at every temperature above 0 °C. We reasoned that global gene-expression machineries (e.g., ribosomes, RNA polymerases) would affect a cell’s doubling time^29,30^ because we found that ROS-reducing enzymes, which require gene expression to produce, are critical for cell duplications. Moreover, we reasoned that a cell needs time to build a daughter cell and replace key cellular components that ROS damaged. These processes involve the same, global gene-expression machineries. Hence, we hypothesized that the combined working speed of global gene-expression machineries are major factors in determining the doubling time and sought to measure the genome-wide transcription rate and protein-synthesis rate at near-freezing temperatures.

### Genome-wide transcription rate at near-freezing temperatures

To determine a genome-wide transcription rate in yeast at frigid temperatures, we incubated the low [GSH] population at various temperatures (1 °C, 5 °C, or 30 °C) in a growth medium containing the nucleotide analogue, 4-thiouracil (“4tU”). 4tU incorporates into every newly synthesized RNA. We quantified the abundance of all newly made transcripts at different time points by using next-generation sequencing of the 4tU-labelled mRNA (“4tU-seq”, see Methods and Figs. S27–S28). Additionally, we used single-molecule RNA FISH^31,32^ on endogenous yeast genes to measure the integer copy-number of transcripts, at their steady-state expression levels, for each temperature. We then converted the abundance of mRNA from the 4tU-seq to an integer copy-number of mRNA per cell (Fig. S29). This revealed a genome-wide transcription dynamics at each temperature (in “# of mRNA per cell per hour”) (Fig. 5b – points). A rate equation^33,34^ with a constitutive synthesis and degradation of mRNA recapitulated the data (Fig. 5b – curves; details in Supplementary Text). This fitting revealed that, on average, a cell synthesized 700 mRNA molecules per hour at 1 °C, 2,000 mRNA molecules per hour at 5 °C, and 58,000 mRNA molecules per hour at 30 °C. Moreover, the fitting revealed that the average half-life of mRNA at each temperature: 14 hours at 1 °C, 7 hours at 5 °C, and 20 minutes at 30 °C. The values at 30 °C closely resemble those of recent studies^35,36^. These measurements established that transcription occurs on the order of hours to days at near-freezing temperatures.

### Protein-synthesis rate at near-freezing temperatures

To determine the timescale for making a functional protein, starting from transcription initiation, we constructed a strain in which galactose induced an expression of mCherry. After adding galactose to the growth medium of the low [GSH] population, we measured mCherry protein abundance in individual cells for up to two weeks at various temperatures (Fig. 5c; Supplementary Movie M6). The global machineries for gene expression, such as RNA polymerases and ribosomes, must function at least as rapidly as the time taken for mCherry fluorescence to increase in cells. A rate equation for gene induction recapitulated our data at all temperatures to yield a mCherry-synthesis rate at each temperature (Fig. 5c – curves; Figs. S30–S31; details in Supplementary Text). By normalizing this rate for each temperature to that of 30 °C, we obtained a fold-reduction in the mCherry-synthesis rate (“protein-synthesis rate”) which represents a slowing down of the global gene-expression machineries (e.g., ribosomes, RNA polymerases). In support of our result, we found that an Arrhenius-type function recapitulated how the protein-synthesis rate decreased as temperature decreased towards 0 °C (Fig. 5d – curve in left graph; see Supplementary Text). These measurements established that protein synthesis, starting from transcription initiation, occurred on the order of a week or longer at near-freezing temperatures.

### Protein synthesis, but not transcription, becomes more rate limiting for cell duplication as environment freezes

The above measurements of transcription and protein-synthesis rates were performed with the low [GSH] population. By measuring the rates again but now in the high [GSH] population, we discovered that ROS did not affect either of the rates at any temperature (Fig. S32). Thus, we can make general conclusions about transcription and protein-synthesis rates at near-freezing temperatures by examining the results for the low [GSH] population. By comparing the transcriptional rate with the protein-synthesis rate at various temperatures for the low [GSH] population, we found that the protein-synthesis rate decreased faster compared to the genome-wide transcription rate as temperature decreased towards 0 °C (Fig. 5d –left graph). This was more evident when we compared how the inverse of each rate changed as a percentage of the doubling time at each temperature. The inverse of each rate yields a characteristic time for a given process (transcription or protein synthesis). By plotting the characteristic time as a fraction of the doubling time at each temperature, we found that as the temperature decreases, mRNA-synthesis time becomes more negligible whereas protein-synthesis time becomes a larger fraction of the doubling time (Fig. 5d - right). Since the characteristic time for protein synthesis – as we have defined it – includes the characteristic time for transcription, this result establishes that, as temperature approaches 0 °C, the working speeds of translational and post-translational machineries become more rate limiting for cell duplication whereas transcription becomes less rate limiting for cell duplication. In returning to our original goal, we will now turn to explaining how the protein-synthesis rate and ROS together yield the speed limits.

### Model to establish range of pace at which cell cycle can be completed at each temperature

We can now determine how a cell’s doubling time arises from its ROS level and protein-synthesis rate at each temperature. In doing so, we will determine how the speed limits emerge. We have shown that cells with more ROS have a longer doubling times and that they rely on ROS-reducing enzymes. Hence, as our measurements showed, a cell with a minimal ROS would have the shortest doubling time *t* which would depend only on the protein-synthesis timescale which is not affected by ROS. For a cell with a non-negligible level of ROS, it would need an additional time Δ*t*, on top of *t*, to remove ROS. Cells with higher ROS would have a larger Δ*t*. Thus, a cell’s doubling time τ is (Fig. 6a):

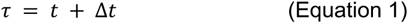

Like the protein-synthesis rate, for all temperatures, we found that an Arrhenius-type function recapitulated the measured doubling times of cells that had the least amounts of ROS. Consequently, we discovered a power-law relationship^37,38^ between *t* and the average protein-synthesis rate *r* : *t* ~ *r*^−0.77^ (Fig. 6b – left, see full details of the model in Supplementary Text). To obtain Δ*t*, which varies among cells as a stochastic variable, we used the fact that the measured distribution of ROS levels among cells was approximately log-normal and that a cell needs time to build ROS-reducing enzymes which depends on the protein-synthesis rate. These two considerations lead to 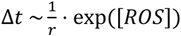 (Fig. 6b – red; see Supplementary Text). With the *t* and Δ*t* now defined, we ran a stochastic simulation in which we used the measured threshold-concentration of ROS to determine whether a cell divides or not and, for a dividing cell, its doubling time (see Supplementary Text). The simulation reproduced the measured distribution of doubling times among cells (shown for 5 °C in Fig. 6c; also see Fig. S33). Importantly, when we plotted together all the measured doubling times of cells for every temperature, we found that all data points lay between the shortest and longest allowed doubling times that the model dictated at every temperature (Fig. 6d). Hence, the data validate our model’s explanation of the origin of speed limits for completing the cell cycle. Specifically, the shortest allowed doubling time *t*, which corresponds to a “high-speed limit”, is defined by the protein-synthesis rate: *t* ~ *r*^−0.77^. The model shows that cells cannot complete the cell cycle any faster than the temperature-specific, high-speed limit because cells cannot synthesize proteins at an arbitrarily fast pace. The longest allowed doubling time *τ_max_* which corresponds to a “low-speed limit”, is determined by the protein-synthesis rate and the threshold concentration of ROS ([*ROS*]*_threshold_*):

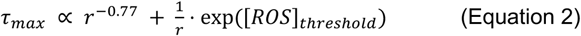

The model shows that cells cannot complete the cell cycle any slower than the temperature-specific, low-speed limit because progressing slower would mean having more than the threshold concentration of ROS which, in turn, causes the cell to keep increasing in size without dividing, until it bursts. The low-speed limit corresponds to an extremely long doubling time at temperatures near 0 °C (e.g., ~3 years at 1 °C). Being stochastic, the model shows that observing such cell duplications is possible but very unlikely (Fig. S26).

## DISCUSSION

Here we investigated how the budding yeast lives and dies at near-freezing temperatures. Our study began without a bias towards any one form of damage or stress that is associated with cold environments. A simple growth experiment – an experiment in which we tested whether an initial number of cells in a population affects the population’s growth at low temperatures – revealed a surprising, cooperative mechanism: yeast cells help each other in surviving and dividing at frigid temperatures by reducing each other’s ROS (by secreting and extracellularly accumulating the antioxidant, glutathione). This discovery establishes that microbes can cooperatively combat harmful effects of cold environments and thereby revises the textbook view in which a microbial cell autonomously fends for its survival at frigid temperatures. This discovery led us to establishing that ROS is the primary determinant of yeast’s ability to survive and divide at frigid temperatures. Specifically, by continuously monitoring individual yeast cells at near-freezing temperatures for unusually long times (i.e., month or longer at 1 °C or 5 °C) and applying quantitative methods to examine individual cells, we discovered how ROS inhibits cell division and causes deaths through its effects on the eukaryotic cell cycle at frigid temperatures. Namely, we discovered that ROS elongates G1 (growth) phase but staying in G1 for longer than a specific, threshold amount of time – determined by a specific, threshold concentration of ROS – makes S2-G2-M (replicative) phase impossible and thus the yeast dies by bursting from being unsustainably large (nearly four times the average size of yeast at 30 °C). We then showed that this threshold G1-duration is temperature specific and set by an interplay of a temperature-dependent, global gene-expression speed and ROS. We then elucidated how the same interplay fundamentally constrains the pace at which a yeast cell can cycle: yeast cannot complete the cell cycle at an arbitrarily fast or slow speed at a given temperature.

Although one can argue without knowing any details that a cell cannot complete its cell cycle at an arbitrarily fast pace at any temperature, it is not obvious - and to date it has not been argued - that there exists a minimum, non-zero speed at which a cell can cycle at a given temperature. Moreover, a detailed, quantitative reasoning is required to explain how any low-speed limit (longest allowed doubling time) and high-speed limit (shortest allowed doubling time) would arise and vary with temperature. We provided such an explanation by discovering that an interplay between the protein-synthesis rate and ROS establishes both the low-speed and high-speed limits for yeast and that changing the temperature tunes this interplay to adjust the speed limits. These results expand our current understanding of how ROS affects cell proliferation^39,40^ which has been unclear in frigid environments.

Finding quantitative “growth laws” for microbes is a goal that has attracted much attention^29,41–48^. We uncovered a temperature-dependent growth law for the budding yeast by examining how temperature tunes quantitative relationships among genome-wide transcription rate, protein-synthesis rate, and cell-proliferation rate. These global relationships have been unclear despite previous studies having examined how temperature affects expression of specific genes in microbes^6,49–51^. By using the budding yeast and using frigid temperatures that are ubiquitous in natural environments, we determined how slowly gene-expression machineries can function and how their unusually slow working speeds can determine and constrain the pace of a cell’s life. Examining this relationship for the budding yeast led us to discovering that speed limits can exist for viably completing the eukaryotic cell cycle in frigid environments, thereby revealing that cold climates can impose unbeatable constraints on a microbe’s life. The systems-level design principles revealed here may be used to engineer potentially other microbes so that they can survive and divide in frigid environments in which they ordinarily die without proliferating. Moreover, the design principles that govern yeast’s life at frigid temperatures and our systems-level approach that uncovered these principles may serve as a case study for future investigations that aim to find similar design principles for other microbes and microbial communities in frigid environments.

## METHODS

### Yeast strains (see Supplementary Text for details)

The “wild-type”, haploid yeast strain w303 that we used is from Euroscarf with the official strain name “20000A”. It is isogenic to another laboratory-standard haploid yeast “W303a”, and has the following genotype: *MATa; his3-11_15; leu2-3_112; ura3-1; trp1*Δ*2; ade2-1; can1-100*. See Supplementary Text for details on engineered and mutant strains.

### Growth media

We cultured all yeasts in defined, minimal media (SC) that consisted of (all from Formedium): Yeast Nitrogen Base (YNB) media (cat. No. CYN0410), Complete Supplement Mixture (CSM, cat. No DCS0019) containing all the essential amino acids and vitamins, and glucose at a saturating concentration (2% *=* 2 g per 100 mL, Melford Biolaboratories Ltd., cat. No. G32040). The agar pads, which we used for growing yeast colonies, contained 2%-agar (VWRChemicals), Yeast Extract and Peptone (YEP) (Melford Biolaboratories Ltd., cat. No. Y20020 and P20240 respectively), and 2% (w/v) glucose.

### Flow cytometry

We used BD FACSCelesta with a High-Throughput Sampler and lasers with the following wave lengths: 405 nm (violet), 488 nm (blue) and 561 nm (yellow/green). We calibrated the FSC and SSC gates to detect only yeast cells (FSC-PMT *=* 681 V, SSC-PMT *=* 264 V, GFP-PMT *=* 485 V, mCherry-PMT *=* 498 V. As a control, flowing dPBS yielded no detected events). The number of cells per mL that we plotted in our growth experiments is proportional to the number of events (yeast cells) that the flow cytometer measured in an aliquot of cells with a defined volume. We measured the GFP fluorescence with a FIT-C channel and the mCherry fluorescence with a mCherry channel. We analysed the flow cytometer data with a custom MATLAB script (MathWorks).

### Growth experiments

In a typical growth experiment, we first picked a single yeast colony from an agar plate and then incubated it at 30 °C for *~*14 hours in 5 mL of minimal medium. Afterwards, we took a 20 µL aliquot from the 5 mL culture, diluted it to a known volume and then flowed it through our flow cytometer to determine the 5 mL culture’s population-density (# of cells / mL). We then serially diluted the culture into fresh minimal media to a desired initial population-density for a growth experiment at the desired temperature. Specifically, we distributed 5 mL of diluted cells to individual wells in a “brick” with twenty-four 10 mL wells (Whatman, “24-well x 10mL assay collection & analysis microplate”). This ensured that we had 4 identical replicate cultures for each initial population-density. We sealed each brick with a breathable film (Diversified Biotech: Breathe-Easy), covered it with a custom-made Styrofoam-cap for insulation, and incubated it in a compressor-cooled, high-precision thermostatic incubators (Memmert ICP260) that stably maintained their target temperature throughout the course of our growth-experiments, with a typical standard deviation of 0.052 °C over time (deviation measured over several days - see Fig. S1). Throughout the incubation, the cultures in the brick were constantly shaken at 400 rpm on a plate shaker(Eppendorf MixMate) that we kept in the incubator. To measure their population densities, we took a small aliquot (typically 250 µL) from each well, diluted it with dPBS (Fisher Bioreagents) into a 96-well plate (Sarstedt, Cat. #9020411), and then flowed it through the flow cytometer which gave us the # of cells / mL. Alternatively, the cultures were distributed into glass tubes, that were kept in the incubator and constantly mixed using a rotator set to 40 rpm.

### Measuring the percentage of dead cells

Cells were incubated as described in the paragraph “growth experiments”. After *~*2 weeks of incubation, we took aliquots of each culture at each time point, and then stained the cells for 20 min with 1 µg / mL of propidium iodide (Thermo Fisher Scientific, cat. No. P3566). We then flowed these (stained) cells through our flow cytometer, and measured the number of cells that were unstained by propidium iodide – these cells have intact membranes, and are assumed to be alive (conversely, stained cells have lost membrane integrity and are assumed dead^54^. We then used the total number of cells and the number of dead or alive cells to extract the growth rate and death rate during several weeks. For this we assumed a simple stochastic growth model that we fitted to our data (see Supplementary Text).

### Microscope sample preparation

All microscopy imaging was performed with 96-well glass-bottom imaging plates (cat. No. 5221-20, Zell-Kontact). Before each sample preparation, the glass bottom of the wells was pre-treated with 0.1 mM concanavalin A for 20 min at room temperature (ConA, Cat. No. C2010, Sigma-Aldrich). We then removed the conA and added an appropriate amount of cells to each well. Typically, we added an aliquot containing *~*7,500 cells and supplemented with sufficient dPBS (Gibco, Life Technologies Limited, cat. No. 14190-144) such that the volume of each well was 200 µL. The plate was then centrifuged at 1,000 rpm for 5 min using a centrifuge (Eppendorf, 5810R) that was precooled at the desired temperature (e.g. 5 °C).

### Microscope data acquisition and time-lapse

We used an Olympus IX81 inverted, epifluorescence, wide-field microscope. Temperature was kept constant during imaging by an incubator cage (OKO Lab) that enclosed the microscope. Fluorescent proteins or fluorescent probes were excited using a wide-spectrum lamp (AMH-600-F6S, Andor) and images were acquired with an EM-CCD Luca R camera (Andor) and IQ3 software (Andor). For time-lapse movies, we prepared yeast cells as described in the paragraph “growth experiments” for two weeks. Aliquots were then transferred to 96-well imaging plates as described in the paragraph “microscope sample preparation”, except that the cultures were not diluted with dPBS. The imaging plates that contained yeast samples were subsequently incubated at the desired temperature throughout the time-lapse (e.g. 5 °C for typically *~*3 weeks), and transported and kept on ice for imaging (typically once every day). We checked that the transport and microscopy – usually less than 15 min during which the imaging plate and samples inevitably warm up – had no observable influence on the samples. To do this, we compared two imaging plates that contained aliquots of the same samples. One plate was imaged once every day for three weeks as described above. The other plate was only imaged once, after the three weeks of incubation and was never transported on ice or warmed up. We found no difference between the cultures in the two plates – in terms of cell density, ROS stress or cell size – after three weeks of incubation.

### Microscope data analysis

We processed the microscope data using ImageJ (1.53c) and MATLAB. Specifically, we segmented the cells by creating oval masks containing the cells, and extracted fluorescence values or cell size from the area inside these masks. We computed the fluorescence for each cell by taking the maximum intensity of 20 images spaced 0.2 µm apart in a z-stack. Finally, we corrected for the background fluorescence by subtracting the average (maximum) background fluorescence in the field-of view from the value obtained for each cell.

### Measuring extracellular reduced and oxidized glutathione

To quantify extracellular glutathione, we isolated the growth media from liquid cultures by flowing liquid cultures through a 0.2 µm pore filter (VWR). We ensured that no cells remained in the filtered media by flowing an aliquot through our flow cytometer. We then measured the total concentration glutathione in the filtered media as described in the manufacturers’ protocol (quantification kit for oxidized and reduced glutathione (cat. No, 38185, sigma aldrich)). To quantify both the concentration of oxidized and reduced glutathion, we took two 200 uL aliquots of the filtered media. To one of the aliquots we then added 4 uL of masking agent provided with the kit (most likely 2-vinylpyridine in ethanol at a final concentration of *~*3 mM). All aliquots were then incubated for 1 hr at 37 °C together with standard curves for reduced glutathione (Cat. No. G4251, sigma Aldrich) without masking agent and oxidized glutathione (G4376, Sigma Aldrich) with masking agent. (During incubation, the masking agent irreversibly binds and thereby removes reduced glutathione. The assay subsequently only detects oxidized glutathione in the samples. We verified that this protocol indeed quantifies the amount of oxidized and reduced glutathione. We found that the masking agent removed *~*90% of the reduced glutathione during incubation, resulting in a false signal of *~*10% reduced glutathione.). We used a spectrophotometer (Spectrostar nano, BMG labtech) to measure the optical absorbance at 415 nm.

### Measuring the concentration of intracellular ROS or intracellular glutathione

We prepared yeast cells as described in the paragraph “growth experiments” and incubated the cultures for two weeks. We then transferred aliquots to 96-well imaging plates as described in the paragraph “Microscope sample preparation”, except that the cultures were not diluted with dPBS. We next removed the supernatant and washed the cells twice with precooled dPBS at 5 °C to remove thiols from the growth media (e.g. cysteine). We then added an indicator dye and incubated the cells for 30 min at 5 °C. As indicator dye, we used either ‘ThiolTracker Violet’ to stain intracellular glutathione (at 20 µM final concentration, Thermo Fisher scientific Cat. No. T10095), ‘mitoSOX red’ to stain intracellular superoxide (at 5 µM final concentration, Thermo Fisher scientific Cat. No. M36008) or ‘cellROX orange’ to stain intracellular ROS (at 5 µM final concentration, Thermo Fisher scientific Cat. No. C10443). Finally, we removed the excess dye by washing the cells twice with dPBS and imaged the cells with a microscope as described in the paragraph “microscope data acquisition and time-lapse”. For a co-stain of multiple indicator dyes, we stained the cells as described above by co-incubating the cells with multiple dyes at the same time. For time-lapse movies of cells stained with the indicator dyes, we followed the above protocol with the following modifications. After preparing the cells and washing away the excess dye, we placed back the supernatant (growth media) that we took away before and kept at 5 °C when incubating the cells with the dye. We then proceeded with the microscopy time-lapse as described in “microscope data acquisition and time-lapse”.

### Measuring extracellular ROS production rate

To measure the extracellular oxidation rate, we used a probe called dihydroethidium (DHE, Thermo Fischer Scientific, Cat. No. D11347) that becomes fluorescent upon oxidation by ROS (superoxide). Dihydroethidium is usually used to detect intracellular ROS, and it intercalates with DNA giving a bright signal in the nucleus. Instead, we used dihydroethidium to detect extracellular ROS where no cellular components were present. To still facilitate the fluorescent signal of dihydroethidium, we added herring sperm DNA (Promega, Cat. D1816) to our samples with a 0.2 mg / mL final concentration. Right before measuring fluorescence, we added dihydroethidium to our samples with a 20 µM final concentration. We then transferred the samples to a flat-bottom 96-well plate (Sarstedt, Cat. #82.1581.001) using 150 µL per well. Directly after, fluorescence was measured using a plate reader (Synergy HTX Multi-Mode Microplate Reader, Biotek) every 3 minutes for 2 hours. Fluorescence was measured with excitation at 500 nm and emission at 620 nm. For measuring the oxidation rate at 30 °C, we pre-warmed all media and set our plate reader to incubate the samples at 30 °C during the measurements. For measuring the oxidation rate at 5 °C, we prepared 5 mL of each sample that we precooled at 5 °C. During the experiment, we incubated the samples at 5 °C for a day. We took 150 µL aliquots of each sample every hour, and transferred these into a 96-well plate. Directly after we measured the fluorescence of these aliquots with our plate reader. For all samples, we measured and averaged the fluorescence of three technical replicates. All measurements included controls consisting of pure water or regular SD media. The oxidation rate was determined by determining the slope (a.u. per second) of the fluorescence curve during *~*1 hour (at 30 °C), starting typically after measuring 10 min with the plate reader. We used the following scavengers of ROS (from Sigma Aldrich): tiron (4,5-dihydroxy-1,3-benzenedisulfonic acid disodium salt monohydrate, cat. No. 172553), trolox ((*±*))-6-Hydroxy-2,5,7,8-tetramethylchromane-2-carboxylic acid, cat. No,238813). Samples with ROS scavengers were compared to a control in appropriate solvent (for example, a sample having trolox that is dissolved in DMSO was compared to a sample having only DMSO).

### Limited nutrients experiment

To test the effect of nutrients on the growth of cells at low temperatures, we prepared fresh wild-type yeast cells as described in the paragraph “growth experiments”, except that we limited the amount of nutrients in the fresh growth media. Specifically, we diluted the minimal media with various amounts of water, and then supplemented each media with 2% glucose. Thus, each media contained a known percentage (0*−*100%) of the nutrients that are in regular minimal media and 2% of glucose. After transferring fresh populations of cells into each media, we incubated the cultures to 5 °C and measured their population density over time as described in the paragraph “growth experiments”.

### Measuring extracellular ROS production during nutrient depletion

To measure the extracellular oxidation rate that cells experience during their incubation at low temperatures, we prepared cultures of our wild-type yeast at various starting densities as described in the paragraph “growth experiments”. We then incubated the cells at the 5 C, and measured the oxidation rate in the growth media over time. To do so, we took aliquots of the cultures that we kept at 5 °C, and flowed them through a 0.2 µm pore filter (VWR, cellulose-acetate membrane). We then directly proceeded to measure the ROS production rate in the supernatant as described in the paragraph “measuring extracellular ROS production rate”.

### Measuring the cell cycle progression

We used the “cell-cycle marker” strain to measure the progression of the cell cycle at low temperatures. We prepared cultures of the “cell-cycle marker” strain as described in the paragraph “growth experiments”. After two weeks, we transferred aliquots of the cultures to a pre-cooled 96-well microscopy plate that we kept on ice. The microscopy plate was further prepared as described in the paragraph “microscope sample preparation”. We kept the plate at the desired temperature (e.g. 5 C) for one day, and then proceeded by taking a snapshot of each sample twice per day for time-lapse movies as described in the paragraph “microscopy data acquisition”. We analysed the time-lapse movies as described in “microscope data analysis”. In short, to quantify the amount of nuclear Whi5-mCherry and H2B-GFP, we first located the nucleus by segmenting the GFP fluorescence of each cell using a threshold GFP fluorescence that we kept fixed for all cells and time points. The nucleus was then the group of pixels whose fluorescences exceeded this threshold. We then determined the total mCherry and GFP fluorescence within the cell’s nucleus. From this fluorescence we subtracted the average background fluorescence in the field-of-view from the value obtained for each cell. Finally, to obtain the copy number of DNA we rescaled the nuclear GFP between the average minimum and maximum GFP fluorescence that we observed for duplicating cells, and to obtain the amount of nuclear Whi5 we took the ratio of the nuclear and cytoplasmic mCherry. (Also see Fig. S20).

### Mutant yeasts

We constructed several mutant strains in which we removed genes involved in (oxidative) stress-response. In short, we designed primers whose ends were homologous to the flanking regions of the desired gene to be knocked out. Using these primers, we amplified a selection marker by PCR, and knocked out the desired gene in the wild-type yeast via homologous recombination. Mutants were selected on YPD selection plates and knockouts were verified by PCR. Specifically, we knocked out the genes for the stress-response transcriptional activators (*MSN2*), membrane organisation (*HSP12*), disaggregase (*HSP104*) glutathione s-transferase (*GTT2*) and glutathione peroxidase (*GPX1*) using the HygB selection marker and YPD plates containing hygromycin B. We also knocked out genes for a suppression of protein aggregation (HSP26), glutaredoxin (*GRX2*), catalase (*CTT1*) and the transcriptional regulator of G1-to-S transition (*WHI5*) using the NatMX selection marker and YPD plates containing nourseothricin. The *MSN2,MSN4* double knockout was constructed by removing, sequentially, first the *MSN2* gene and then the *MSN4* gene. We thus obtained several mutants that lacked genes for transcriptional regulation (*msn2,msn4*Δ-strain, *whi5*Δ-strain) or that lacked genes for the oxidative stress response (*gtt2*Δ-strain, *gpx1*Δ-strain, *grx2*Δ-strain and *ctt1*Δ-strain).

### FISH probes

We designed single-molecule FISH probes to detect mCherry-mRNA. For this we used the Stellaris FISH probe designer (LGC Biosearch Technologies; www.biosearchtech.com). The set of probes (25 probes) were designed to attach to the full length of mCherry RNA and were coupled to Quasar 670 (a Cy5 analog, LGC Biosearch Technologies). We also designed FISH probes to detect mRNA of endogenous yeast genes to convert TPM values from our 4tU RNA-sequencing data to integer numbers of RNA per cell. For this, we used probes for RPS3 (30 probes coupled to Quasar 670), RPL3 (48 probes coupled to Quasar 570, a Cy3 analog LGC Biosearch Technologies), RPB1 (48 probes coupled to Quasar 670) and RPB3 (40 probes coupled to Quasar 570). The excitation and emmission peaks of these fluorophores are ex. 548 / em. 566 nm (Quasar 570) and ex. 647 / em. 760 nm (Quasar 670).

### Single-molecule RNA FISH

We used the standard protocol for single-molecule RNA FISH in yeast, as described in “Protocol for *S. cerevisiae* from Stellaris RNA FISH” (LGC Biosearch Technologies). Finally, we made sure to image the fluorescence of Quasar 670 probes first during our measurements, as this was the dye most sensitive to photo bleaching.

### Measuring single-gene transcription rate

We used the “mCherry-inducible” strain to measure the transcription rate. We prepared cultures of the mCherry-inducible strain as described in the paragraph “growth experiments”, except that we used minimal media (SC) containing 2% raffinose as the growth media. The cultures were incubated in 500 mL Erlenmeyer flasks 5 °C for two weeks on an Eppendorf platform shaker set to 125 rpm. After two weeks, we supplemented the cultures with 2% galactose. After further incubating the cultures at 5 °C for the desired amounts of time, we transferred aliquots of the cultures (typically *~*10 mL) to 15 mL tubes containing 37% formaldehyde such that the final volume formaldehyde was 10%. We then proceeded with RNA FISH as described in the paragraph “single-molecule RNA FISH”.

### Measuring single-gene expression rate

We used the “inducible mCherry” strain to measure the gene expression rate. We prepared cultures of the mCherry-inducible strain as described in the paragraph “growth experiments”, except that the minimal media (SC) contained 2% raffinose as the carbon source. We then incubated the cultures for two weeks in glass tubes at 5 °C in a rotator at 40 rpm. After two weeks, we transferred aliquots of the cultures to a pre-cooled 96-well microscopy plate that we kept on ice. The microscopy plate was further prepared as described in the paragraph “microscope sample preparation”. As media, we used fresh, pre-cooled SC containing 2% raffinose to dilute the aliquots to the desired density on the microscopy plate. We kept the plate at the desired temperature (e.g. 5 °C) for one day, and then took a snapshot of the populations as described in the paragraph “microscope data acquisition and time-lapse”. Finally, we supplemented each sample on the microscopy plate with 2% galactose to induce the expression of mCherry. We then proceeded by taking a snapshot of each sample twice per day for time-lapse movies as described in the paragraph “microscopy data acquisition”. In parallel, we also added 2% galactose to the original cultures that were kept in the rotator at 5 °C. We then measured the average mCherry fluorescence of the population twice per day by flowing aliquots of the cultures through our flow cytometer as described in the paragraphs “growth experiments” and “flow cytometry”.

### Preparing cells and RNA extraction

To prepare fresh cells we first picked a single yeast colony from an agar plate and then incubated it at 30 °C for *~*14 hours in 6 mL of SD media. We then took aliquots of this culture and spun them down using a centrifuge. For each aliquot, we removed the supernatant and resuspended the pellet in fresh media of the desired composition (for example, fresh SD, or fresh SD containing 250 µM glutathione, or 0.25x SD – consisting of 25 volumes of regular SD and 75 volumes of water containing 2% glucose). Typically, we added *~*10 mL of fresh media. Each new culture contained the cells from *~*1 mL of the initial culture (initial density *~*500,000 cells / mL). We incubated the new cultures at the desired temperature for 14 days in glass tubes in a rotator set to 40 rpm. We performed RNA extractions using the RiboPure Yeast RNA extraction kit (Thermo Fischer Scientific, cat. No. AM1926) following the kit instructions. We also performed the DNAse treatment after RNA extraction, and stored the isolated RNA in elution buffer at *−*80 °C before further processing.

### Measuring genome-wide transcription rate with 4tU labelled RNA (see Supplementary Text for details)

In short, we prepared large cultures of our wild-type yeast similarly to the description in “growth experiments” and added 4-thiouracil (4tU) to the growth media at a final 5 mM concentration^55^. Samples were subsequently collected after desired amounts of time at a each temperature. We discarded the supernatant and re-suspended the pellet in 1 mL RNAlater (Cat. No. AM7021, Thermo Fischer Scientific). As a spike-in of 4tU labelled RNA we used a fixed amount of cells from *Schizosaccharomyces pombe* (YFS110) analogously to previous work^55^. We then spun down our samples in a pre-cooled centrifuge, removed the RNAlater, and proceeded with RNA extraction as described in the paragraph “preparing cells and RNA extraction”. After RNA extraction, we proceeded with biotinylation and purification of the 4tU labelled RNA following existing protocols with minor modifications^55^. After sequencing, we processed all sequencing data with the Salmon tool to quantify relative transcript abundance^56^. Finally, we converted the transcript levels for *S. cerevisiae* to gene expression levels (Transcripts Per Million, TPM), merged all samples using the package tximport from Bioconductor^57^ and converted our 4tU time-lapses to “# of RNA / cell” (see Figs. S27–S29 for further details).

### Mathematical model

Derivations of equations, a detailed description of the mathematical model, and the parameter values used for simulations are in the Supplementary Text.

### Code availability

All scripts used for simulations in this work will be publicly available (GitHub diederiklt/YeastLowTemperatures).

### Data Availability

The authors declare that all data supporting the findings of this study are available within the paper and its supplementary information files. RNA-Seq data is available at NCBI GEO. Source data for the main figures is provided. The data that support the findings of this study are available from the corresponding author upon reasonable request.

## ADDITIONAL INFORMATION

Supplementary Figures 1-33

Supplementary Movies 1-6

Supplementary text that describes mathematical modeling

## Supporting information

Supplementary Figs and Model

## ACKNOLWEDGEMENTS

We thank Jennifer Benanti, Amir Mitchell, Job Dekker, Marian Walhout, and Michael Lee for insightful comments and suggestions. We thank Nicholas Rhind for providing the *S. pombe* strain that we used as a control for our 4tU-Seq data. We thank Ezequiel Calvo and Athma Pai for suggesting the 4tU-Seq experiment. H.Y. was supported by the European Research Council (ERC) Starting Grant (MultiCellSysBio, #677972), Netherlands Organisation for Scientific Research (NWO) Vidi Award (#680-47-544), CIFAR Azrieli Global Scholars Program, and EMBO Young Investigator Award.

## AUTHOR CONTRIBUTIONS

D.S.L.T. and H.Y. conceived the project. D.S.L.T. designed and performed all experiments, analyzed all data, and built the mathematical model. T.M. helped with microscopy imaging. D.S.L.T. and H.Y. examined and discussed all data analyses. H.Y. supervised the research. D.S.L.T. and H.Y. wrote the manuscript with input from T.M.

## Supplementary Information

### Supplementary Figures

**Figure S1:**
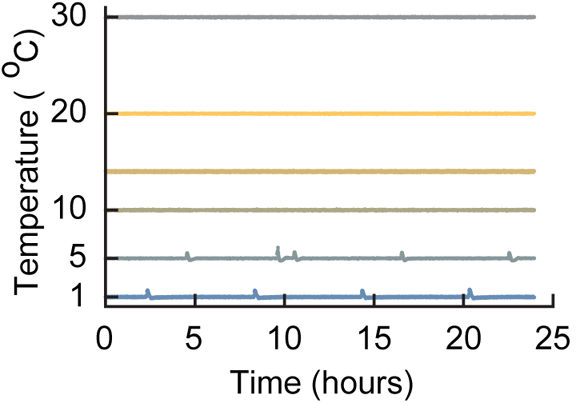
Temperature remains stable during all our growth experiments (Related to Figure 1a). All growth experiments were performed with liquid cultures of cells incubated in compressor-cooled, high-precision thermostatic incubators (Memmert ICPs). The ICPs had a precise temperature-recording device whose temperature readings we additionally verified using a separate thermocouple device and aligned with the temperatures of several other incubators. The temperature was monitored over the entire course of our experiments. As examples, shown here are the temperatures as recorded by the incubator’s temperature sensor for 24 hours during six different growth experiments. Starting from the top, the curves show the temperature of the incubator set at 30.0 °C, 20.0 °C, 14.0 °C, 10.0 °C, 5.0 ^°^C and 1.0 °C. Throughout our experiments, the incubators had a typical standard deviation of 0.033 ^°^C when the incubation temperature was above 10.0 °C. The standard deviation was 0.09 °C when the incubation temperature was below 10.0 °C. This slightly larger standard deviation was due the incubator undergoing short thaw-cycles when the incubation temperature was 5.0 °C to prevent freezing of its components (deviation measured over several days).

**Figure S2:**
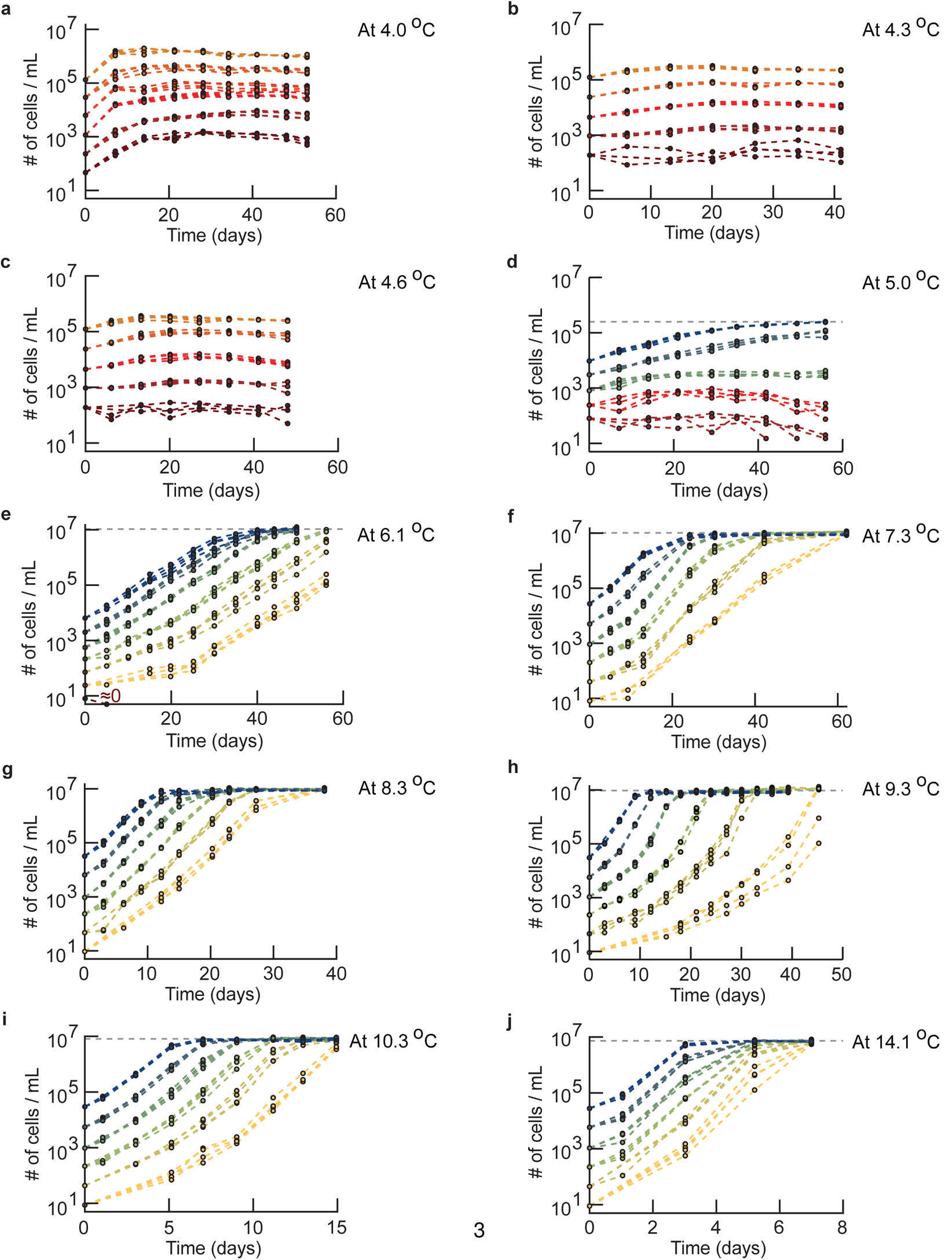
Growth curves for widely varying initial population densities and temperatures (Related to Figure 1a-b). Population density (number of cells / mL) measured over time with a flow cytometer. Shown are populations of wild-type yeast with differing initial densities incubated at 4.0 °C **(a)**, 4.3 °C **(b)**, 4.7 °C **(c)**, 5.0 °C **(d)**, 6.1 °C **(e)**, 7.3 °C **(f)**, 8.3 °C **(g)**, 9.3 °C **(h)**, 10.3 °C **(i)** and 14.1 ^°^C **(j)**. Different colors represent different initial population-densities. The grey line shows the carrying capacity that we estimated from the final densities of the populations. To construct the phase diagram (Figure 1b), we used the growth-kinetics data as shown here to determine whether a population with a given initial population-density should be characterized as growing (”growth” phase) or non-growing (”no growth” phase). An initial population-density was characterized as being in the growth phase (Fig. 1b – blue region) if all replicate populations that started with that density exponentially grow over time and reach the carrying capacity for that temperature. An initial population-density was characterized as being in the no-growth phase (Fig. 1b – red region) if all replicate populations that started with that density did not grow during several weeks of incubation, except for some initial transient growth that results from the cells having been transferred from 30.0 °C. This transient growth typically lasted a few days. For example, every population in (c) belongs to the no-growth phase because no population grew during the ~6 weeks of incubation at 4.6 °C as can be seen by every population density barely increasing over time. As another example, after some lag-time, every population in (g) grew exponentially and identically over time to reach the carrying capacity at 8.0 °C. To draw the phase boundary that separates the growth phase from the no-growth phase in the phase diagram (Fig. 1b), we connected the maximum initial density for which a population does not grow for each temperature. This curve almost overlaps with the minimum initial density that leads to population growth for each temperature because we sampled the initial densities close to each other for each temperature. Similarly, we drew the boundary curve that separates the growth phase from the region of the phase diagram where nutrients are lacking by connecting the data points that represent, for each temperature, the measured carrying capacity (the grey lines in d-j). Finally, we determined the temperature below which no population growth is possible — and thus only the no-growth phase exists below this temperature – by identifying the highest temperature (i.e., 4.6 °C (c)) where populations with different starting densities always reach differing final densities (as opposed to the common, carrying capacity). In fact, these populations do not grow at all beyond the initial, transient growths (a-c). All panels show *n* = 4 replicate populations for each initial population-density (color).

**Figure S3:**
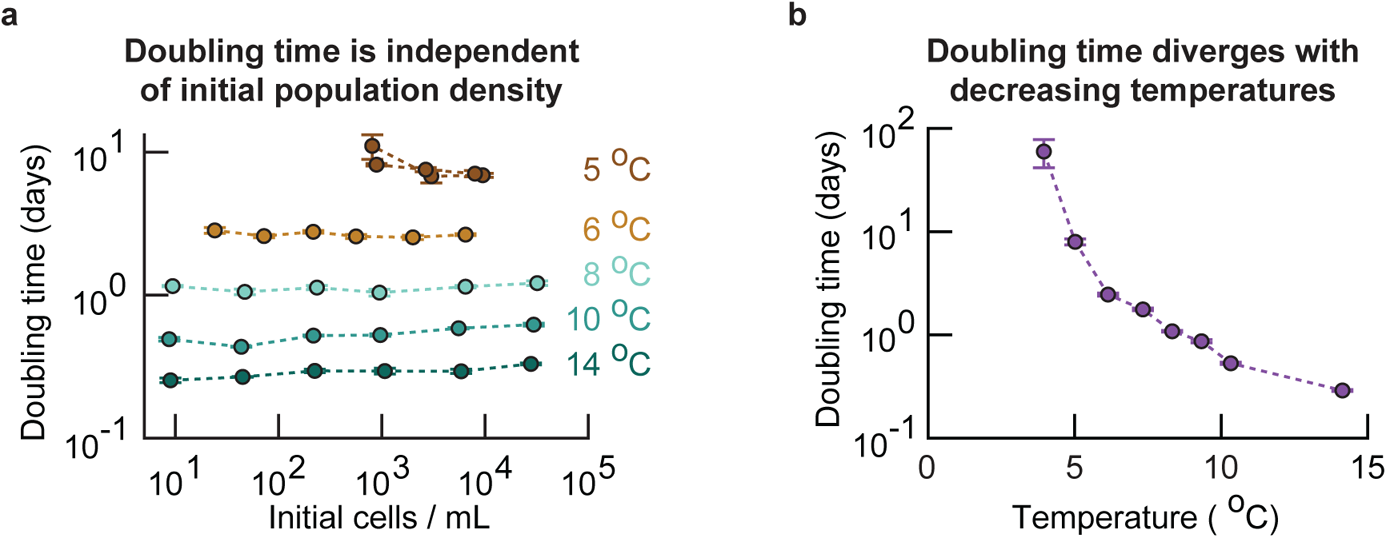
Population’s doubling time is independent of its initial density at near-freezing temperatures (Related to Figure 1b). Summarizing Supplementary Fig. S2. From the growth curves of populations at various temperatures, we determined each population’s doubling time. To obtain the doubling time of the populations that grew (i.e., populations in the ”growth” phase of the phase diagram, Fig. 1b), we excluded the initial transient growth that typically lasted from a few days to one week. For non-growing populations (e.g., populations in the ”no-growth” phase at 4.0 °C) we only took the growth rates that were positive and excluded the initial transient growth (i.e., transient growth in the first week of incubation). **(a)** Population doubling time as a function of initial density and temperature for growing populations. The doubling time does not depend on initial density. Error bars show the mean with s.e.m., and each data point represents at least *n* = 4 replicate populations. **(b)** Since the doubling time does not depend on initial density, we pooled all observed doubling times for each temperature. The population doubling time diverges as the temperature decreases. Error bars represent the mean with s.e.m., having at least *n* = 16 biological replicates per data point.

**Figure S4:**
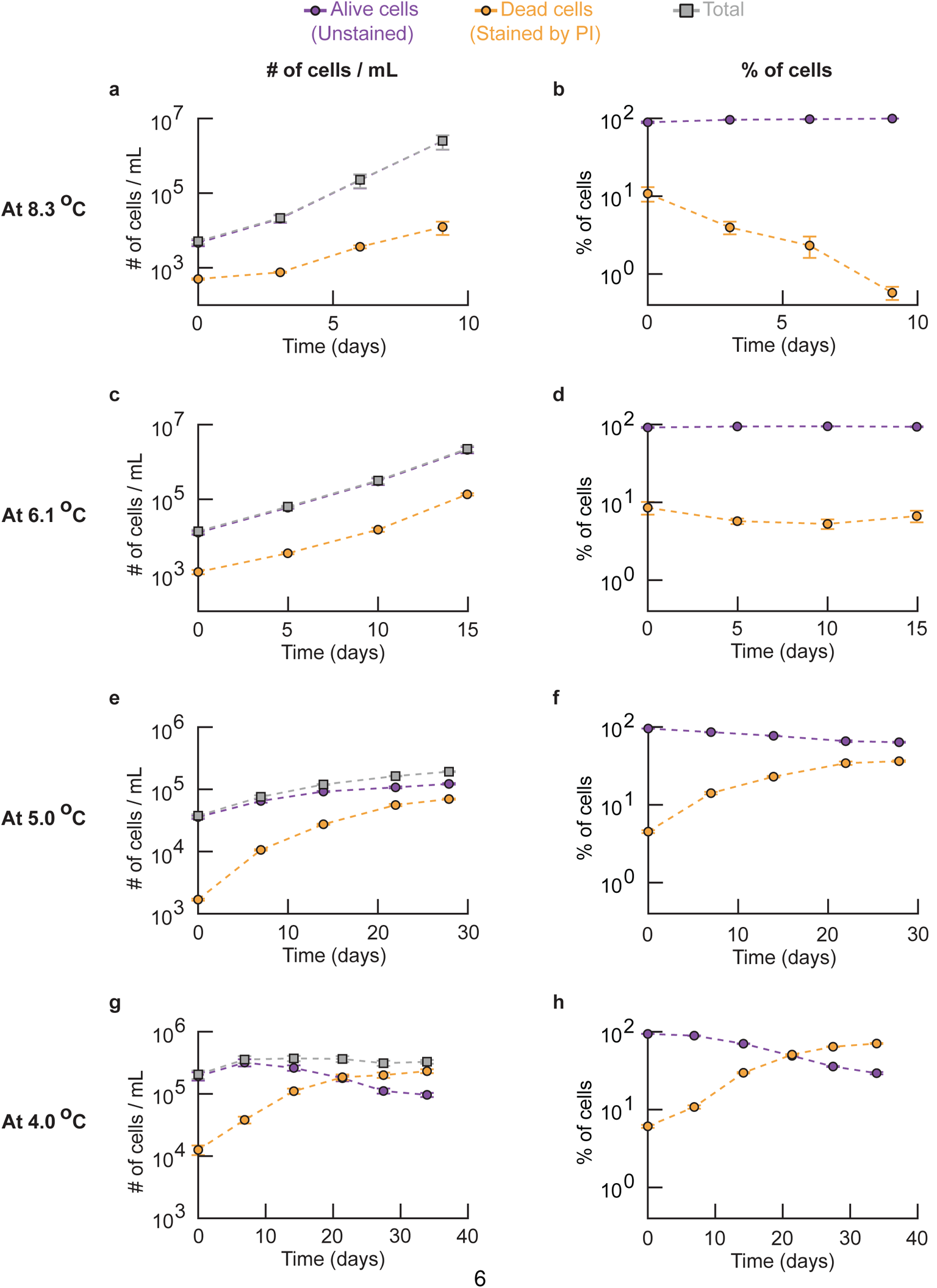
Population extinction occurs below 5 °C because as the temperature decreases, the average doubling time of a cell increases and eventually, just below ~5 °C, becomes larger than the average cell’s lifespan (Related to Fig. 1b). Left column (**a, c, e, g**): number of dead cells (yellow), number of alive cells (purple) and total number of cells (grey) in a population. Right column **(b, d, f, h)**: percentages of dead (yellow) and alive (purple) cells in a population. For each population, we took an aliquot of the liquid culture at various times and incubated it with 1 μg / mL of propidium iodide (PI) for 20 minutes at room temperature. We then flowed this aliquot through a flow cytometer to measure the total number of cells and the number of cells that were stained (red) and unstained (yellow) by PI. PI does not stain cells with an intact membrane [1]. In contrast, PI enters cells with a damaged membrane and stains their DNA. Thus, propidium iodide stains a cell if and only if it’s membrane is permeable. Hence, alive cells are commonly assumed to be impermeable to PI. Shown here are populations of wild-type yeast incubated at 8.3 °C **(a-b)**, 6.1 °C **(c-d)**, 5.0 °C **(e-f)** and 4.0 °C **(g-h)**. The curves show the total population-density and the density of stained and unstained cells over time **(a, c, e** and **f)**, together with the percentage of cells that were stained and unstained in the population **(b, d, f** and **h)**. All populations were already in a steady-state at the respective temperature before the PI staining on day 0 in the graphs (i.e., the populations had already spent ~6 days at 8.3 °C (a-b), ~10 days at 6.1 °C (c-d), and ~14 days at 5.0 °C (e-f) or 4.0 °C (g-h)). Error bars show the mean with s.e.m., with *n* = 4 replicate populations per data point. **(a-b)** Alive and dead cells over time at 8.3 °C. All populations grow exponentially over time. Both the density of alive and dead cells increase exponentially over time, with the density of alive cells increasing faster than the density of dead cells. Specifically, the percentage of dead cells in the population decreases exponentially over time. Thus, at 8.3 °C, alive cells overtake the population as cell duplications occur more frequently than cell deaths (purple curve in (b) stays at ~100%). **(c-d)** Alive and dead cells over time at 6.1 °C. Similar to the results at 8.3 °C. The main difference here is that the density of alive cells and the density of dead cells increases exponentially with the same rate (the two curves in (c) are nearly parallel), whereas the density of alive cells increased faster than the density of dead cells at 8.3 °C (see (a)). Indeed, the percentage of dead cells remains constant over time at 6.1 °C (see d). Like in 8.3 °C, nearly 100% of the population consists of alive cells at 6.1 °C. **(e-f)** Alive and dead cells over time at 5.0 °C. The density of alive cells increases over time. However, the percentage of dead cells in the population increases while the percentage of alive cells decreases over time. Still, the populations consists of more alive cells than dead cells after ~6 weeks of incubation at 5.0 °C (see (f)). **(g-h)** Alive and dead cells over time at 4.0 °C. The population does not grow and the density of alive cells decreases over time (g) (i.e., less than one cells divides per cell that dies). The percentage of dead cells increases exponentially over time while the percentage of alive cells decreases, leading to the population eventually consisting mostly of dead cells (the population becomes extinct as this trend continues over time) (see (h)). Together, (a-h) show that as the temperature decreases, the average doubling time of a cell increases and eventually becomes larger than the average lifespan of a cell. Having a doubling time that is larger than the average lifespan means that the average cell does not have time to duplicate before it dies. This leads to a population extinction. In other words, populations grow more slowly as temperature decreases and eventually, at around 5.0 °C, populations cannot exponentially grow because the average doubling time nearly matches the average lifespan of cells. Finally, at 4.0 °C, the doubling time exceeds the lifespan of cells and thus populations cannot grow.

**Figure S5:**
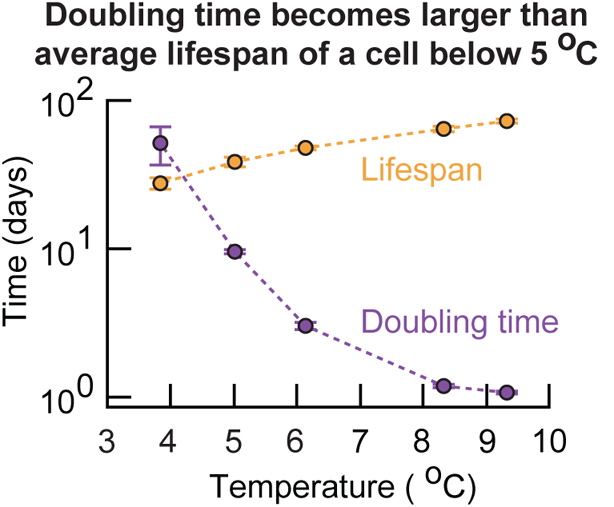
Doubling time of a cell increases and eventually becomes larger than the averace cell’s lifespan as the temperature decreases below 5 °C (Related to Figure 1b). Summarizing Supplementary Fig. S4. We used our measurements of the number of alive and dead cells in populations at each temperature to extract the average doubling time and the average lifespan of cells. Specifically, we fitted a simple growth model to the data (also see Supplementary Theory). Shown is the average cell’s doubling time (purple) and lifespan (orange) at various temperatures. The average doubling time exceeds the average lifespan when the temperature is below 5.0 °C. Consequently, below 5.0 °C, a cell is expected to die before it has a chance to duplicate and hence the population approaches extinction. Error bars represent the mean with s.e.m., having at least *n* = 16 biological replicates per data point.

**Figure S6:**
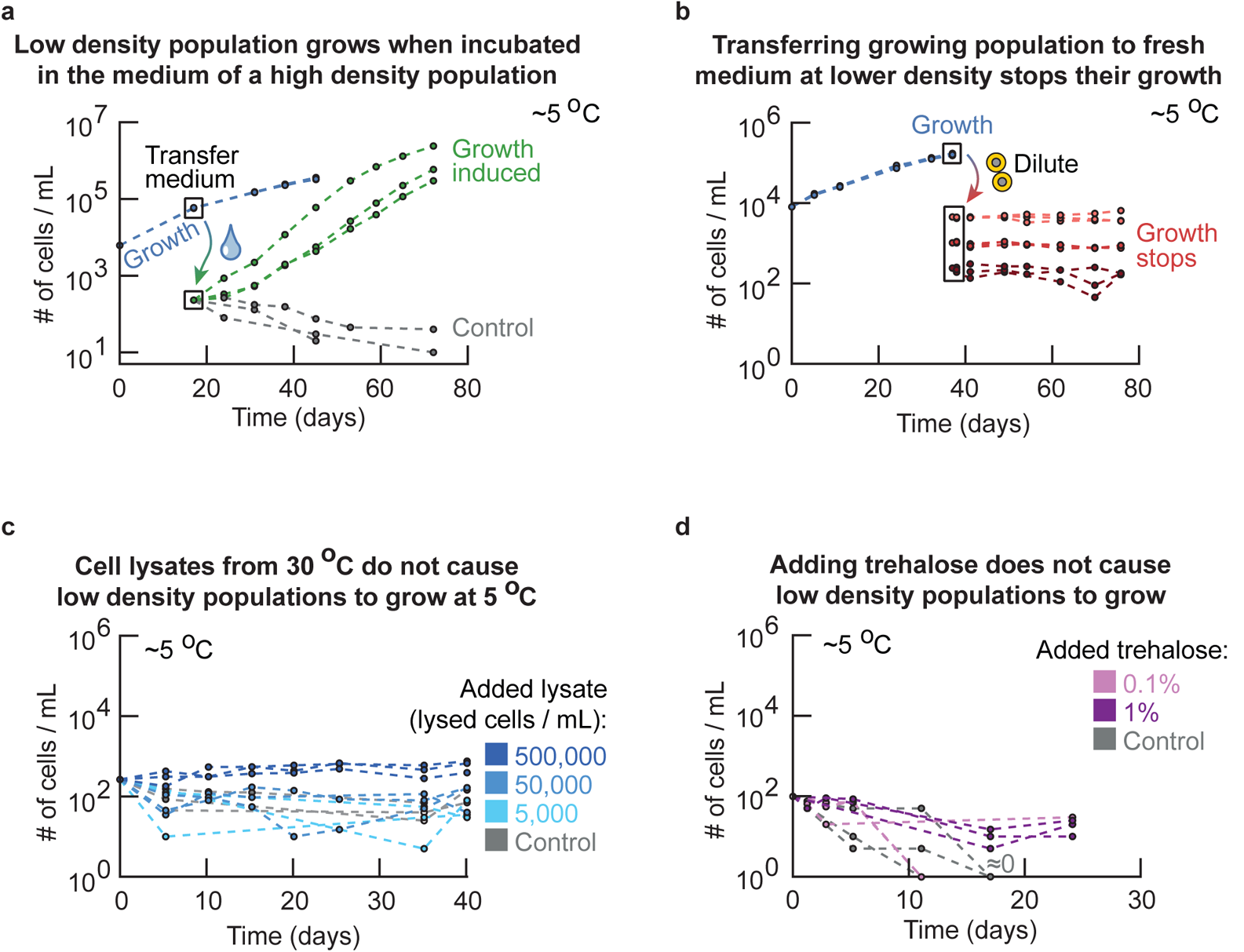
Secreted factor in extracellular medium determines whether a population grows or not at freezing temperatures (Related to Figure 2a). Testing why high-density populations grow whereas low-density populations do not grow at the same near-freezing temperature (5.0 °C). **(a)** To test whether the cell’s ability to duplicate at 5.0 °Cis dictated by secreted factor(s) in the extracellular medium, we took the growth medium of a high-density (growing) population (blue curves: initially ~6, 250 cells / mL) after several weeks of growth at 5.0 °C. Boxed data points on the blue curves show when the growth medium was taken. We flowed the high-density culture through a membrane filter with 0.2 μm pores so that growth medium taken from this culture was free of cells, which we confirmed with a flow cytometer (i.e., no cells were detected). Next, into this filtered medium, we incubated a fresh, low-density population of cells that were growing in 30 °C (green curves: initially ~250 cells / mL). As a control, we also incubated the same, low-density of cells in a fresh growth medium (grey curves: initially ~250 cells / mL). We incubated the two, low-density cultures at 5.0 °C and measured their cell numbers over time. The populations of fresh cells grew in the filtered growth medium of the high-density population at 5.0 °C (green curves). But the low-density (control) populations in the fresh medium did not grow (grey curves). This experiment shows that cells change their growth medium (e.g., secreted factor(s)) such that they can grow at 5.0 °C and that a sufficiently high density of cells is required for the change to be sufficient for a population growth. **(b)** To test whether the ability to duplicate at 5.0 °C is determined solely by intracellular factor(s) (e.g., heritable trait), we took an aliquot of cells from a high-density (growing) population at 5.0 °C and diluted it by various amounts into fresh, pre-cooled media at 5.0 °C to test whether the resulting low-density population could grow. The high-density population of wild-type cells (blue curves, initially ~10, 000 cells / mL) grew at 5.0 °C. After ~35 days of growth, we took an aliquot of cells from this culture and diluted it by ~50x (light red curves, initially ~10, 000 cells / mL), ~250x (red curves) or ~1250x (dark red curves). Ordinarily, populations that start at these ”low densities” do not grow at 5.0 °C (Fig. 1b). We incubated the diluted, low-density populations in fresh medium at 5.0 °C and measured their density over time (red curves). None of these populations grew at all during more than one month of incubation at 5.0 °C. This experiment shows that the ability to duplicate at 5 °C is not determined by any intracellular factor(s). **(c)** To test whether the extracellular factors that enable population growth come from dying cells that may be lysing, we incubated populations in media with cell lysates. We obtained the cell lysates from a population of wild-type cells growing in log-phase at 30.0 °C (~5, 000, 000 cells / mL). We added glass beads to this liquid culture and placed it on a vortex for 20 min to mechanically lyse the cells. After this, we flowed the liquid media containing the lysed cells with the beads through a 0.45 μm pore filter to remove the glass beads and any intact cells that remained. We then added the filtered cell lysate to a fresh medium in amounts dictated by the cell densities before lysis (e.g., fresh medium having 500, 000 lysed cells / mL contains 90% fresh medium and ~10% cell lysate that had ~5, 000, 000 cells / mL before the lysis). Finally, we incubated fresh cell populations in these growth media at ~250 cells / mL and at 5.0 °C. The curves here show populations with ~500, 000 lysed cells / mL (dark blue curves), ~50, 000 lysed cells / mL (blue curves), ~5, 000 cells / mL (light blue curves) and, as control, fresh cells in a fresh medium without any cell lysate (grey curves). None of these populations grew at 5.0 °C. This experiment shows dying cells that may be lysing are not responsible for inducing population growths at 5.0 °C. **(d)** Trehalose is a common cryoprotectant that is abundant in yeasts at low temperatures and is thought to protect the cells against freezing [2]. Its precise mechanism is incompletely understood. We reasoned that yeasts may be secreting trahalose to help them grow at 5.0 °C. To test this idea, we incubated a low-density population at 5.0 °C (initially ~100 cells / mL) with 1% (dark purple curves, percentage in weight per volume), 0.1% (light purple curves) or without trehalose (grey curves). None of the populations grew, indicating that trehalose is not one of the extracellular factor(s) that induce growth at 5.0 °C. All panels show at least *n* = 3 biological replicates per condition (color).

**Figure S7:**
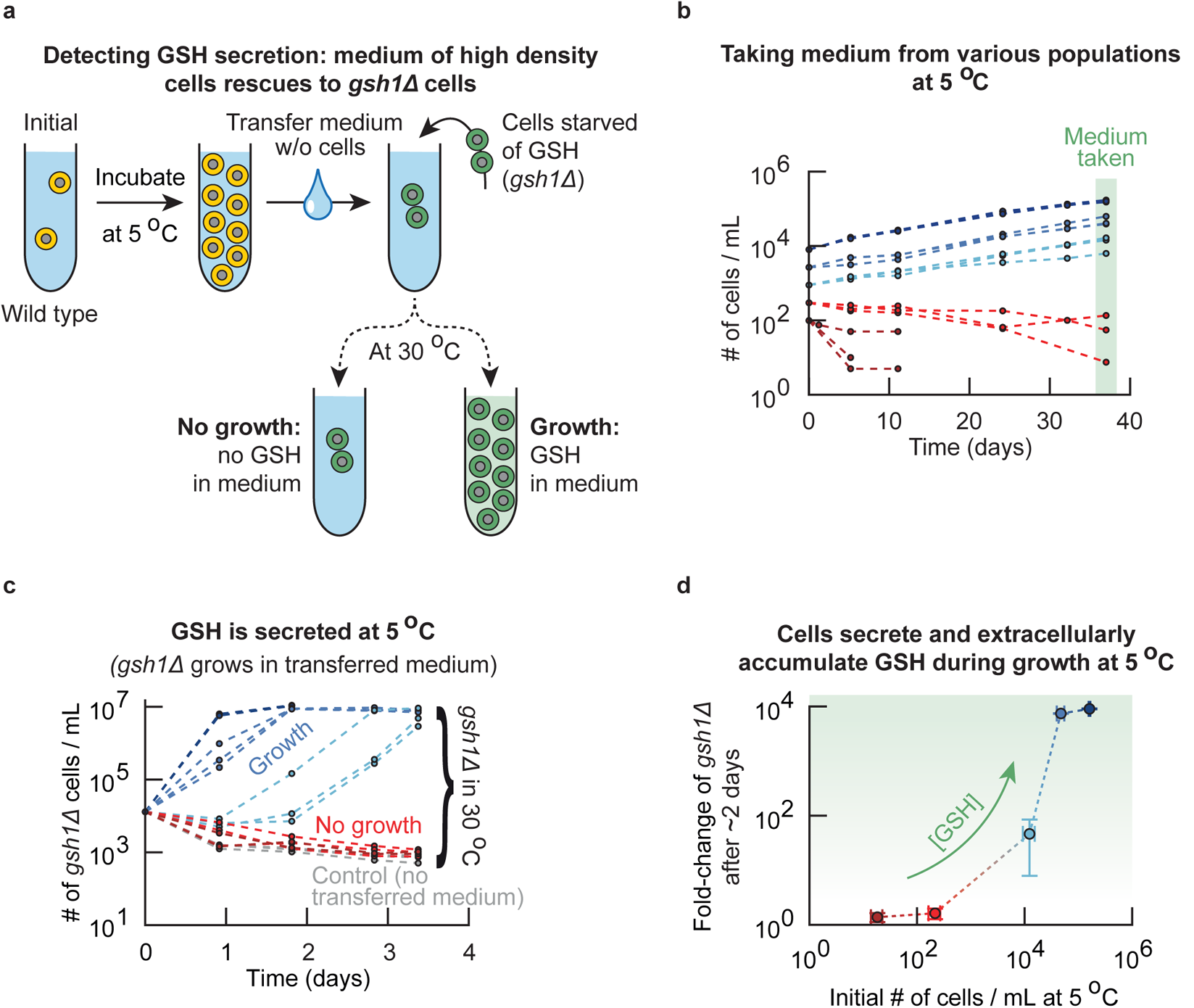
A non-viable mutant that cannot synthesize glutathione (*gsh1*Δ) is rescued by medium from high-density populations, which confirms that cells in high-density populations secrete and extracellularly accumulate glutathione at 5 °C (Related to Figure 2a). **(a)** Schematics of experiments in (b-d). A mutant that cannot synthesize glutathione (*gsh1*Δ-strain) cannot live without supplemented GSH even at 30 °C because GSH is essential for life (e.g., GSH is involved in iron metabolism [3]). The only way for the mutant to survive and grow at any temperature is by supplementing GSH into its growth medium [4,5]. To test whether cells at 5.0 °C secrete glutathione during log-phase growth and stationary phases, we incubated wild-type yeast populations at 5.0 °C at different initial population-densities. After several weeks of growth, we isolated the growth medium from the wild-type populations. Specifically, we took aliquots of the cultures at 5.0 °C and removed the cells from the growth media by spinning down the aliquot and passing the supernatant through a 0.2 μm pore filter to remove all yeasts. We checked that no cells remained in the filtered media by flowing them through a flow cytometer (no events detected). In the filtered media, we incubated a population of *gsh1*Δ cells that we had starved of glutathione by incubating them overnight in fresh medium without GSH at 30.0 °C (These cells therefore did not grow overnight). If the filtered media did not contain any glutathione, then the starved *gsh1*Δ cells would not grow. If the filtered media did contain glutathione secreted by the wild-type cells, the *gsh1*Δ cells would be able to import the GSH and grow. **(b)** As a first part of the experiment described in (a), we incubated wild-type yeast populations at 5.0 °C and determined whether their growth media contained glutathione after ~5 weeks of growth. Shown is the wild-type’s population density over time for different starting densities (~300 cells / mL (red curves), ~900 cells / mL (light blue curves), ~2, 700 cells / mL (blue curves) and ~8, 000 cells / mL (dark blue curves)). Each color shows *n* = 3 replicate populations. **(c)** To detect extracellular glutathione, we isolated and filtered the growth media from the wild-type populations in (b). Shown is the population density over time at 30.0 °C of glutathione-starved *gsh1*Δ cells that received the filtered media (initially ~13, 000 cells / mL). The colors of the curves here match the colors used for the wild-type populations in (b) (e.g., the red populations received the filtered media from the wild-type populations whose growth curves are shown in red in (b)). As a control, we also incubated *gsh1*Δ cells in fresh medium without any glutathione (grey curves). The *gsh1*Δ cells did not grow in the medium transferred from low-density (non-growing) populations at 5.0 °C. The *gsh1*Δ cells always grew to the carrying capacity in the medium from the high-density (growing) populations at 5.0 °C. Each color shows *n* = 3 replicate populations. **(d)** Summary of the data from (b) and (c). The x-axis shows the population density of wild-type cells after ~5 weeks of incubation at 5.0 °C. The y-axis shows, after two days of incubation at 30 °C, the number of *gsh1*Δ cells in transferred media relative to the number of *gsh1*Δ cells in a fresh medium without glutathione. Populations of *gsh1*Δ cells that grew more had more glutathione in their medium (glutathione is the growth limiting factor in the transferred media, see control in (c)). Error bars represent the mean with s.e.m., with *n* = 3 biological replicates per data point. Together, (a-d) show that the amount of extracellular glutathione increases with population density for growing populations, and that barely any glutathione is secreted for non-growing populations. In summary, cells secrete and extracellularly accumulate glutathione during growth at near-freezing temperatures.

**Figure S8:**
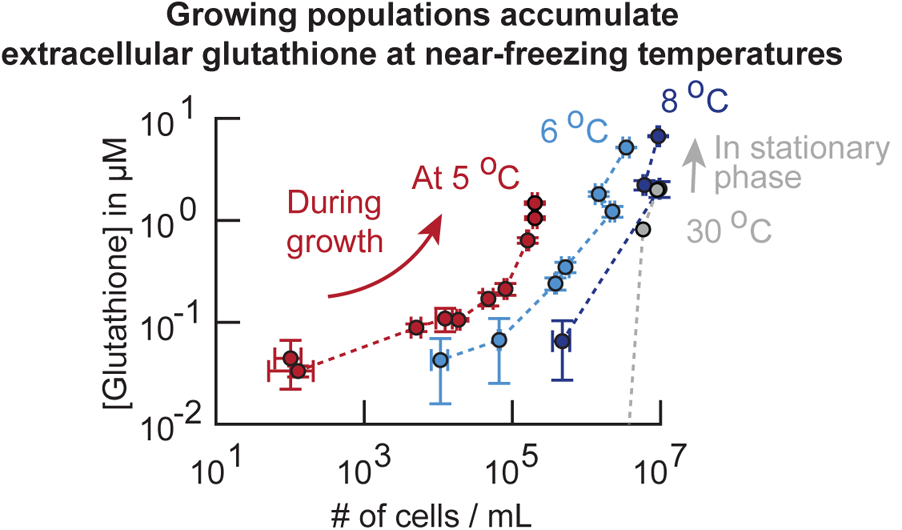
Cells secrete and accumulate glutathione during growth at near-freezing temperatures (Related to Figure 2a). Secretion of glutathione into the extracellular medium at various temperatures. We measured the total glutathione concentration in the growth media of wild-type yeast populations at two different times after ~2 weeks of incubation at various temperatures (see Methods). Shown here is the total extracellular glutathione concentration as function of population density at ~5.0 °C (red curve, one week between sampling), ~6.1 °C (light blue curve), ~8.3 °C (dark blue curve) and ~30.0 °C (grey curve). The extracellular glutathione concentration increases during population growth at near-freezing temperatures (i.e., 5.0 °C - 8.3 °C), whereas glutathione only accumulates in stationary phase at 30.0 °C as we previously found [5] (data reused). Error bars show the mean with s.e.m., having *n* = 3 biological replicates per data point.

**Figure S9:**
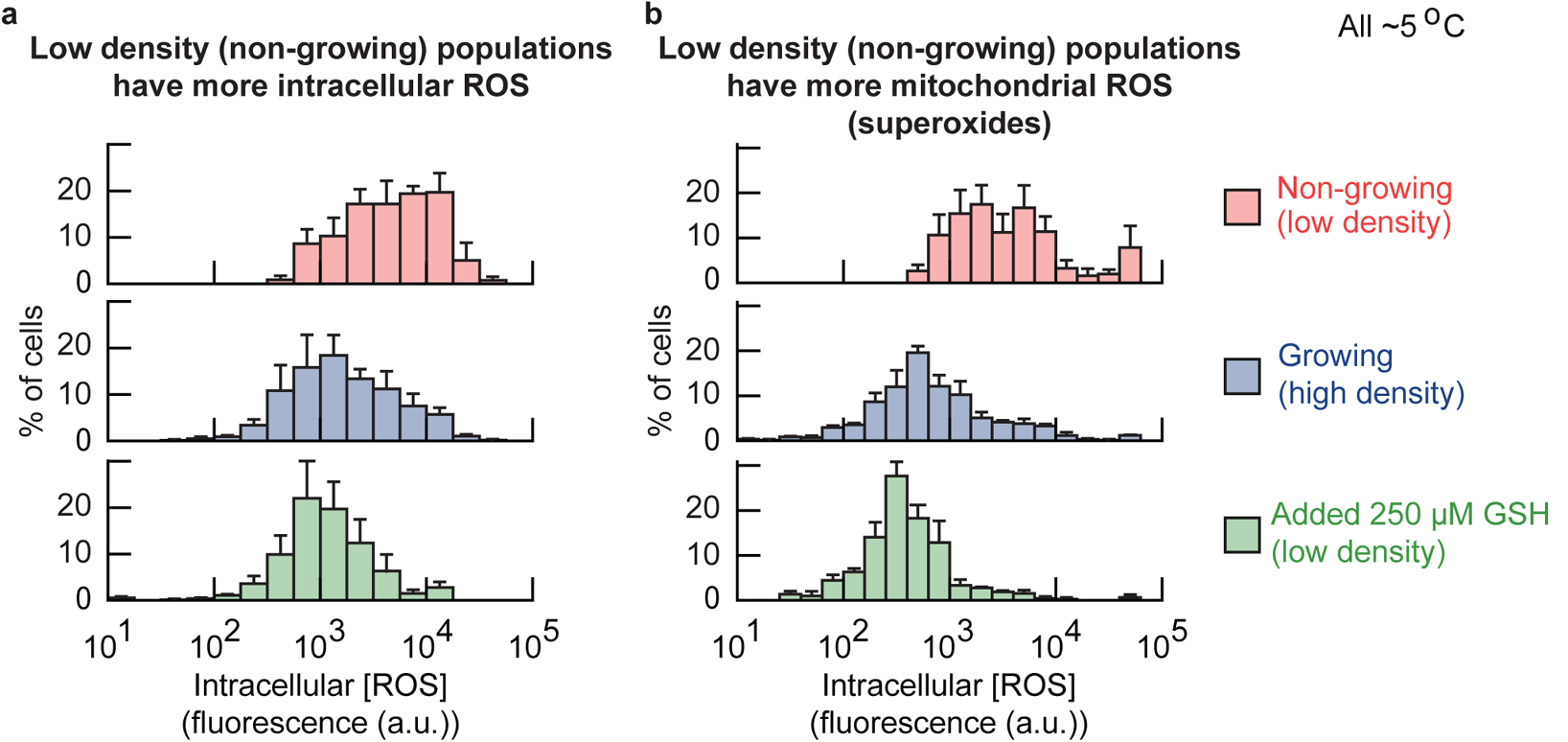
Cells of low-density (non-growing) populations at 5 °C have more intracellular Reactive Oxygen Species (ROS), including superoxides in their mitochondria, which can be removed by supplementing extracellular reduced glutathione (GSH) (Related to Figure 2d). **(a-b)** Intracellular ROS concentrations. We stained intracellular ROS with two different dyes. One dye (cellROX) measures the amount of general, cytoplasmic ROS while another dye (mitoSOX) measures the amount of superoxides in mitochondria. We grew populations of cells at 5.0 °C for two weeks and then stained the cells with the ROS dyes (see Methods). We then used fluorescence microscopy to determine the amounts of intracellular ROS in single cells, as represented by the average fluorescence of each cell (after subtracting background fluorescence). **(a)** Intracellular ROS concentration for general ROS (with cellROX). **(b)** Superoxide concentrations in mitochondria (with mitoSOX). Histograms in (a-b) show cells of low-density (non-growing) populations (red bars, initially ~250 cells / mL), growing populations (blue bars, initially ~6, 250 cells / mL) and populations that had 250 μM added GSH (green bars, initially ~250 cells / mL). These histograms show that the low-density (non-growing) populations have higher intracellular ROS concentrations than high-density (growing) populations. Adding extracellular GSH decreases the intracellular ROS concentrations. Error bars show the mean with s.e.m., having *n* = 3 biological replicates per histogram.

**Figure S10:**
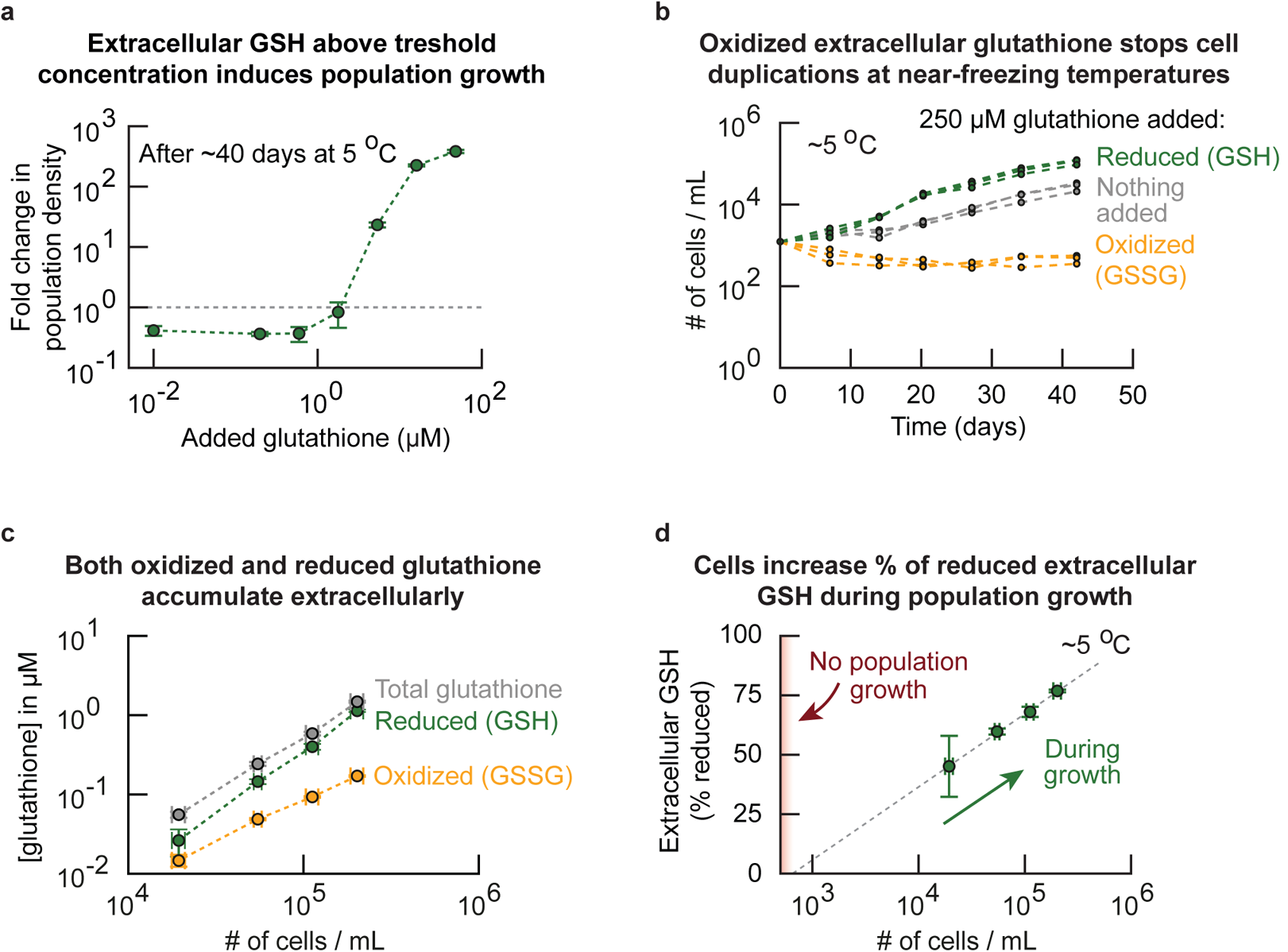
Reduced form of glutathione, not the oxidized form of glutathione, enables and accelerates duplications at 5 °C above threshold concentration of ~1 μM (Related to Figure 2d). Unless stated otherwise, our experiments measured and added the reduced form of glutathione (GSH). But for the experiments in this figure, we used both the reduced glutathione (GSH) and oxidized form of glutathione (GSSG) to determine whether GSSG induces growth at 5.0 °C (answer: it does not). **(a)** We incubated a low-density of wild-type cell population (initially ~260 cells / mL) in various amounts of added (reduced) glutathione at 5.0 °C. The GSH concentration spans almost four orders of magnitude. Shown here is the fold-change of the population density after ~40 days of incubation. Populations expand when the extracellular GSH concentration exceeds ~1 μM, while populations shrink if the extracellular GSH concentration is less than ~1 μM. Thus, low-density populations require at least 1 μM of GSH to grow at 5.0 °C. This threshold concentration is high compared to the ~0.1 μM of GSH that the high-density populations accumulate themselves (Fig. 2a). This result suggests that there are other extracellular factors that induce population growths at 5.0 °C, because the concentration of secreted GSH is an order of magnitude below the concentration of the GSH that must be added to cause a low-density population to grow at 5.0 °C. Error bars show the mean with s.e.m., having *n* = 3 replicate populations per data point. **(b)** So far, we established that extracellular GSH can induce population growths (a). Next, we wondered whether cells at 5.0 °C specifically require glutathione as an antioxidant for the growth induction. The reduced form of glutathione, GSH, functions as an antioxidant whereas the antioxidant function does not exist in the oxidized form, GSSG (two oxidized glutathione molecules). We incubated cells at 5.0 °C with either oxidized or reduced glutathione in their growth medium. Shown here are the population densities over time for 250 μM reduced glutathione (GSH, green curves), 250 μM oxidized glutathione (GSSG, yellow curves) and without any glutathione added to the medium (grey curves, all initially at ~1, 240 cells / mL). All populations grew when either GSH or nothing was added to the medium, with the GSH causing faster growths. In contrast, the populations that had GSSG (oxidized glutathione) added to their medium did not grow at all. These results show that populations require GSH (reduced form of glutathione) to grow at 5.0 °C. Each condition (color) shows *n* = 3 replicate populations. **(c)** We next measured the concentrations of both extracellular oxidized and extracellular reduced glutathione that accumulatd in the medium of high-density (growing) populations (see Methods). Shown here are the total (grey curve), reduced (green curve) and oxidized (yellow curve) extracellular glutathione concentrations in the same medium as a function of density (concentrations quantified after 28 days at 5.0 °C). The total extracellular glutathione concentration increases as the initial density increases (also see Fig. 2a). Both the concentrations of reduced and oxidized glutathione increase with density, with the concentration of reduced glutathione increasing faster. Error bars show the mean with s.e.m., having *n* = 3 biological replicates per density. **(d)** Using the data from (c), we plotted the percentage of extracellular GSH as a function of population density. Grey dotted line shows a linear fit as a visual guide. The percentage of GSH increases with density. Extrapolating the linear fit suggests that all extracellular glutathione is oxidized in the medium of populations with a density below 1, 000 cells / mL (non-growing populations, Fig. 1a). Together, our data ((c-d) and Fig. 2a) establish that cells in high-density populations secrete and accumulate a pool of extracellular GSH, whereas cells in low-density populations accumulate only oxidized extracellular glutathione in low amounts. Moreover, these results show that populations cannot grow when the entire pool of extracellular glutathione becomes oxidized (b). Error bars show the mean with s.e.m, having *n* = 3 biological replicates per data point.

**Figure S11:**
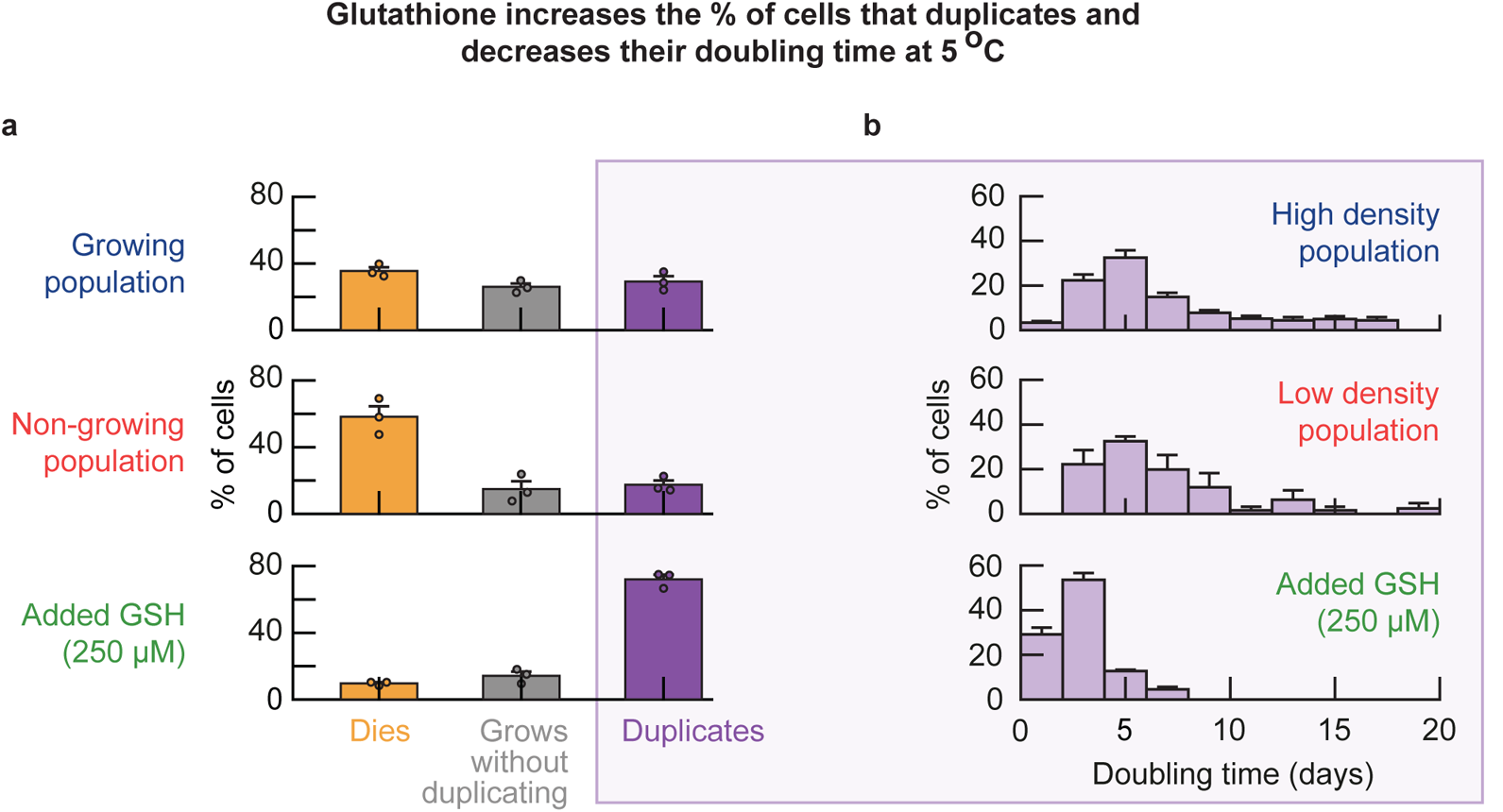
Glutathione increases the percentage of duplicating cells in a population and decreases their doubling time (Related to Figure 2e). Comparing the behavior of single cells at near-freezing temperatures with versus without added glutathione (”GSH”). **(a)** We incubated wild-type cell populations at 5.0 °C with or without adding GSH to their growth medium. After two weeks of incubation, we observed the behavior of individual cells with a microscopy time-lapse (see Methods). Shown here are the percentages of cells that duplicate (purple bar), die (orange bar) and grow without duplicating (grey bar) during the ~20 days of microscope imaging. Three populations were imaged: high-density (growing) population (initially ~6, 250 cells / mL), low-density (non-growing) population (initially ~250 cells / mL), and low-density population with 250 μM of added GSH (initially ~250 cells / mL). Note that 250 μM is higher than the ~1 μM of glutathione that the high-density population accumulates by itself (Fig. 2a). The added GSH decreases the percentage of cells that die in the low-density population to 9.7 *±* 0.7% (compared to ~35% in high-density populations). Similarly, the added GSH increases the percentage of cells that duplicates in the low-density population to 72.1 *±* 2.7% (compared to ~17% in the low-density populations without added GSH). Thus, extracellular GSH decreases cell deaths and increases cell duplications. Bars show the mean with s.e.m., having *n* = 3 biological replicates per condition. Dots show raw data. **(b)** Doubling time for cells that duplicate in each population. Histogram shows the percentage of cells with a given doubling time. The average doubling time is 6.5 days for the high-density populations (*n* = 451 cells) and 7.1 days for the low-density populations (*n* = 42 cells). Thus, the doubling time of duplicating cells is similar regardless of population density. In contrast, the average doubling time is 2.9 *±* 0.1 days for the low-density populations with added GSH (*n* = 444 cells). GSH therefore reduces the time it takes to duplicate by more than half. Data shows the mean with s.e.m. for each bin, having *n* = 3 biological replicates per condition. Together (a-b) show that both high-density (growing) and low-density (non-growing) populations contain duplicating cells, with nearly identical doubling time distributions. The main differences between the two populations are the percentage of cells in each population that duplicates and the percentage of cells that die. Adding extracellular GSH increases the percentage of duplicating cells (a), and decreases the doubling time of these cells (b). Data for growing and non-growing populations in (a-b) is reproduced here from Fig. 1c for comparison.

**Figure S12:**
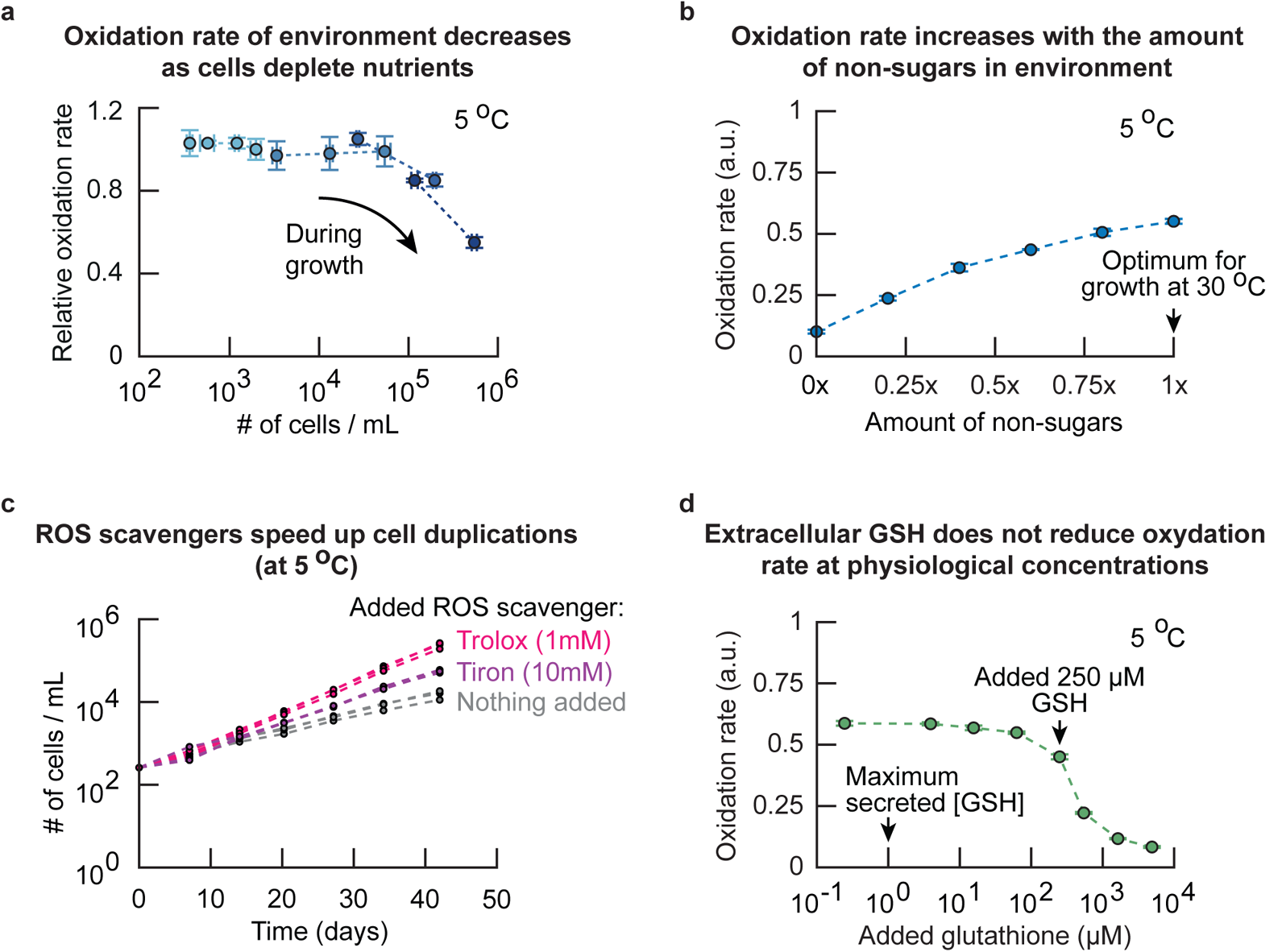
Non-sugar components of growth medium (e.g., vitamins, amino acids) and oxygen create Reactive Oxygen Species without any cells (Related to Figure 2f). Together, our results suggest that more extracellular factor(s) – besides reduced glutathione – dictate population growth at near-freezing temperatures (Supplementary Figs. S6 and S10). Knowing that extracellular factor(s) dictate growth at near-freezing temperatures (Fig. S6), cells are stressed by ROS (Fig. 2a, Supplementary Fig. S9) and that it is the antioxidant action of GSH that induces population growth (Supplementary Fig. S10), we hypothesized that the extracellular environment might also be a source of ROS. **(a)** To test whether ROS are present in the extracellular environment of our yeast populations, we used a dye called dihydroethidium (DHE) that becomes fluorescent upon oxidation by ROS (mainly superoxides). We grew wild-type yeast populations with various starting densities at 5.0 °C, and measured the oxidation rate of the growth media relative to the oxidation rate of fresh media. To do so, at every timepoint, we took aliquots of our liquid cultures and flowed the aliquot through a 0.2 μm pore filter. We then measured the oxidation rate in the flow-through – the growth medium without any cells – and compared with the oxidation rate in fresh medium. Shown is the relative oxidation rate as function of population density. The oxidation rate decreases when the population density goes above ~10, 000 cells / mL. This result shows that ROS is present in the extracellular environment of cells and that the extracellular oxidation rate in the medium decreases with population density. **(b)** Given that the oxidation rate decreases with population density, we further hypothesized that the components of the growth medium – which the cells consume and deplete during population growth – might be a source of ROS. We therefore tested whether the presence of any of the nutrients in medium affects the ROS production of the environment. Specifically, we measured the oxidation rate of DHE at 5.0 °C in growth media having various compositions. We diluted the growth medium with water by various amounts and then supplemented these diluted media with 2% glucose (so a ”0.5x-medium” contains 0.5x SC + 0.5x water, supplemented with 2% glucose). Shown is the oxidation rate in diluted media at 5.0 °C (blue curve, in steps of 0.2x non-sugars). We found that the oxidation rate increased roughly linearly with the amount of non-sugar nutrients in the medium (e.g., amino acids, vitamins, trace elements, salts and the nitrogen source). **(c)** Knowing that the extracellular environment may be a source of ROS through the non-sugar nutrients in the growth medium, we added chemicals to the growth medium that remove (i.e., have scavenging activity against) ROS. Specifically, we used scavengers of superoxide (10 mM tiron) or peroxyl radicals (1 mM trolox) and tested whether the scavenging agents could be beneficial for yeast growth. We grew populations of cells at 5.0 °C (initially ~250 cells / mL). Shown is the population density over time for added trolox (pink curves), added tiron (purple curves) or nothing added to the growth medium (grey curves). Both trolox and tiron increase the growth rate of populations. Thus, scavenging and removing extracellular ROS (such as superoxide and peroxyl radicals) is beneficial for growth. **(d)** Finally, we tested the effect of GSH on the oxidation rate of the growth medium. Shown is the oxidation rate of DHE as function of the GSH concentration in minimal medium at 5.0 °C. Increasing the GSH concentration above a threshold concentration – ~250 μM GSH, at non-phyisiological levels – decreases the oxidation rate. For all panels the error bars are mean with s.e.m., with *n* = 3 replicates per condition. In summary (a-d) show that the non-sugar nutrients in the growth medium increase the oxidation rate of the environment by inducing the generation of ROS. The oxidation rate decreases when non-sugar nutrients are removed from the growth medium. Examples of sources of ROS in the environment are oxygen (leading to superoxide) and the non-sugar nutrients themselves (leading to peroxyl radicals), and removing these ROS is beneficial for yeast growth at near-freezing temperatures. Finally, we found that GSH can decrease the extracellular oxidation rate, albeit at high concentrations (*>* 100 μM).

**Figure S13:**
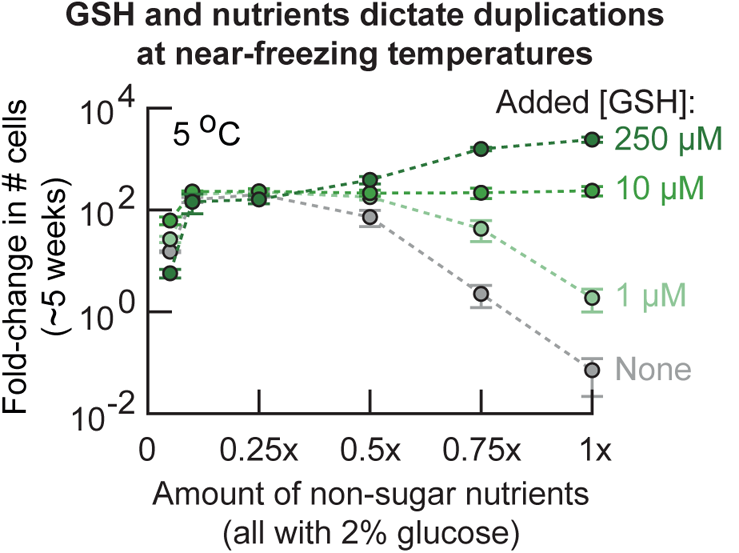
Ability of cells to duplicate is a balance between extracellular GSH concentration and availability of non-sugar nutrients (Related to Figure 2i). Studying the combined effect of non-sugar nutrients and extracellular GSH on population growth. We incubated cells in media that had precise amounts of non-sugar nutrients and added GSH. Specifically, for varying the amount of non-sugar nutrients, we formulated the media to contain 0.05x, 0.1x, 0.25x, 0.5x, 0.75x or 1*x* non-sugar nutrients. Here, 1x equals the amount of non-sugar nutrients that is in the medium for yeast grown under standard conditions (see Methods). Each of these media had 2% glucose. Then, for each of these medium compositions, we added either 0 μM, 1 μM, 10 μM or 250 μM of GSH. This procedure created 24 different media (6 different amounts of non-sugar nutrients each with 4 different added GSH concentrations). In these media we incubated low-density populations of cells (initially ~210 cells / mL) at 5.0 °C. These low-density populations should not grow according to our phase diagram (Fig. 1b). Shown is the fold-change in population density as function of the amount of non-sugar nutrients after ~5 weeks of incubation at 5.0 °C. The curves represent 1 μM (light green curve), 10 μM (green curve), 250 μM (dark green curve) or without (grey curve) added GSH. The population in 1x non-sugar nutrients and without added GSH did not grow at all during the 5 weeks of incubation at 5.0 °C, as expected from our phase diagram (Fig. 1b). Further corroborating our other data, we find that at 1x non-sugar nutrients the fold-change in population density increases with the concentration of added GSH (also see Fig. 2e) and that without added GSH, the fold-change in population density increases when the amount of non-sugar nutrients decreases (grey curve, also see Fig. 2g). Then, starting at 1x non-sugar nutrients, all curves converge to a ~100-fold change in population density as the amount of non-sugar nutrients decreases from 1x to 0.25x. Thus, adding more than 10 μM GSH is only beneficial for population growth when sufficient non-sugar nutrients are available (i.e., more than 0.5x non-sugar nutrients). Finally, populations stop growing when the amount of non-sugar nutrients decreases below 0.1x. In summary, this growth experiment shows that either sufficient extracellular GSH (*>* 1 μM, Supplementary Fig. S10) or low enough amounts of non-sugar nutrients are required for population growth at 5.0 °C. Populations having less non-sugar nutrients require less extracellular GSH for growth, and populations having more extracellular GSH can grow with more non-sugar nutrients. The media with *>* 0.75x non-sugar nutrients and 250 μM added GSH are special because they permit growths that exceed the (~100-fold) growths that a population can achieve by secreting *<* 1 μM glutathione (Fig. 2a) by itself (light green curve). Error bars show the mean with s.e.m., having *n* = 3 biological replicates per condition.

**Figure S14:**
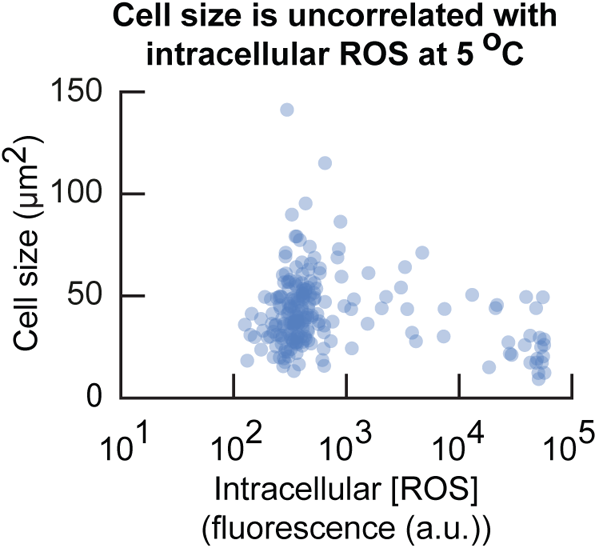
Cell size does not correlate with the intracellular [ROS] at 5 °C (Related to Figure 3c). Cell size as a function of intracellular ROS concentration. We incubated populations of cells for two weeks at 5.0 °C (initially ~6, 250 cells / mL). We then measured the size and intracellular ROS concentration in single cells at 5.0 °C (see Methods). This result shows that the cell size is uncorrelated with intracellular ROS concentration. Dots show single-cell data aggregated from *n* = 3 biological replicates.

**Figure S15:**
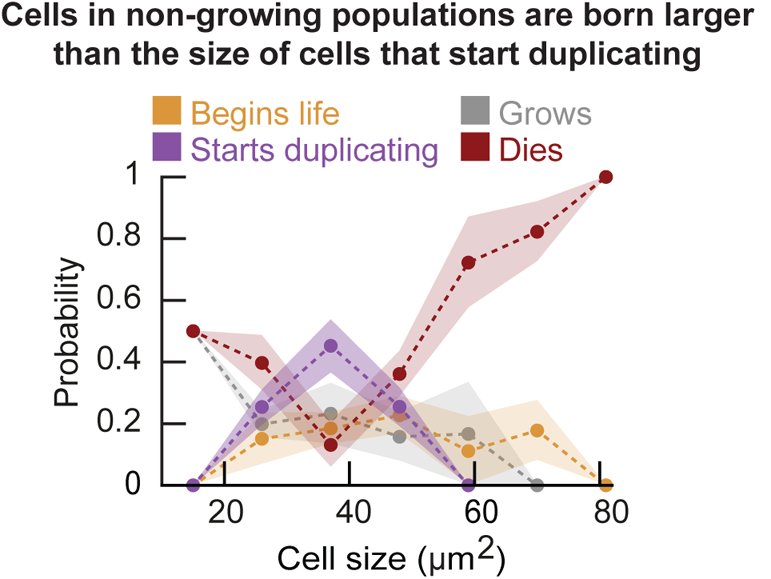
Newborn cells in low-density (non-growing) populations are larger than those in high-density (growing) populations (Related to Figure 3c). We incubated populations of wild-type cells at a low density (initially ~500 cells / mL) for two weeks at 5.0 °C and then imaged these cells in our wide-field microscope for the next several weeks (see Methods). We classified individual cells in the resulting time-lapse movies as being in one of the following classes (see Supplementary Fig. S16): ”begins life” (just-born daughter cell), ”starts duplicating”, ”grows” (without duplicating), or ”dies” (cell just before death). For every cell in each class, we measured its cell size. Specifically, for cells the cells in the ”Begins life” class, we measured the daughter cell’s size just after cytokinesis (when the mother’s bud neck breaks). For the cells in the ”Starts duplicating” class, we measured the mother cell’s size at the time the bud appears. For the cells in the ”Grows” class, we measured their size in the first time point when the movie begins (i.e., while they are in growing). For the cells in the ”Dies” class, we measured their size just before they die (in the last time point before death). We binned the cells according to their sizes. For each bin, we determined the probability that a cell in that size bin would be in each of the four classes. Dots show average probability in each bin for the different events, shaded area represents the s.e.m. for *n* = 3 biological replicates. This graph shows that newborn cells in low-density (non-growing) populations are on average larger than the cells that duplicate in the same population. Thus, newborn cells in low-density (non-growing) populations are less likely to duplicate than the mother cells in the same population (also see Supplementary Fig. S16).

**Figure S16:**
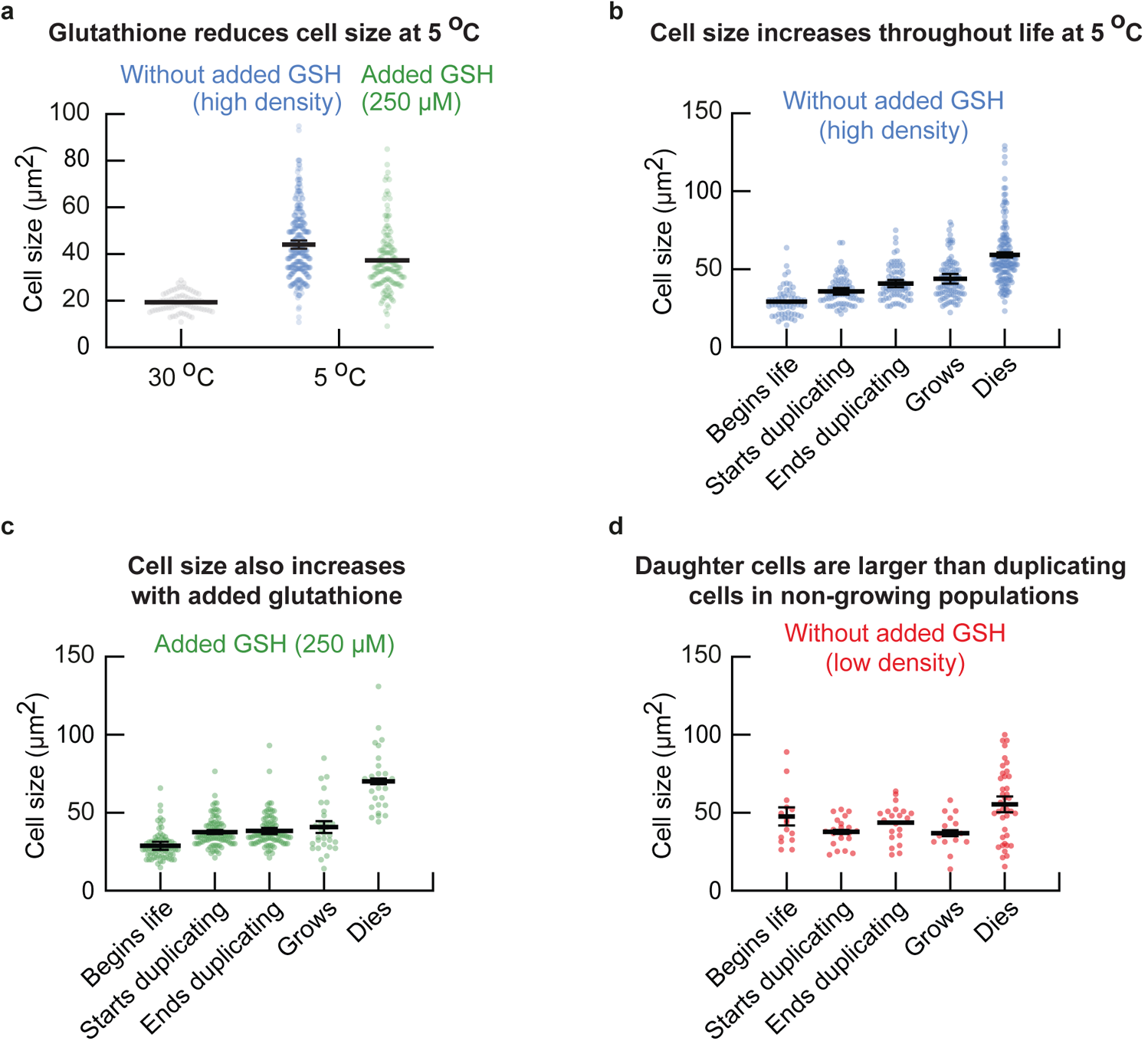
Cell size monotonically increases over time throughout a cell’s life at 5 °C (Related to Figure 3c). **(a)** We incubated wild-type cell populations for two weeks at 5.0 °C and then measured the cell sizes with a wide-field microsope. Shown here are the cell sizes of individual cells from high-density populations without any GSH added (blue dots, initially ~8, 000 cells / mL) and sizes of individual cells from low-density populations that were incubated with 250 μM GSH (green dots, initially ~420 cells / mL). Grey dots show cells in a log-phase growth at 30.0 °C for comparison. Surprisingly, we find that cells are, on average, more than double the size (cross-sectional area) at 5.0 °C (44.0 *±* 1.7 μm^2^) compared to 30.0 °C(19.3 *±* 0.2 μm^2^). Moreover, adding GSH to the growth medium reduces the cell size at 5.0 ^°^C (37.3 *±* 0.1 μm^2^). Error bars represent the mean with s.e.m., having *n* = 3 biological replicates. Dots show data aggregated from the biological replicates. **(b-d)** Next, we incubated populations of wild-type cells for two weeks at 5.0 °C. After two weeks we transferred aliquots of the liquid cultures to microscopy plates that we kept chilled at 5.0 °C before the transfer and took a snapshot with a microscope every day (see Methods). The resulting snapshots form time-lapse movies at 5.0 °C. From these movies, we classified the cells into one of the following events: ”begins life” (size of the daughter cell right after the mother cell finishes cytokinesis), ”start duplicating” (size of the mother cell right after it forms a bud), ”end duplicating” (size of the mother cell right after the mother cell finishes cytokinesis), ”grows” (cells that did not duplicate during the entire time-lapse movie of ~17 days) and ”dies” (cell size just before dying). For all cells in each class, we determined the cell size at the start of each event. Shown here are the cell sizes for populations with high initial density (~8, 000 cells / mL) **(b)**, low initial density (~420 cells / mL) with 250 μM added GSH **(c)**, and low initial density (~420 cells / mL) without any GSH added **(d)**. Dots show size of individual cells in each class. For all panels, the error bar shows the mean with s.e.m., having *n* = 3 biological replicates. Dots are data aggregated from all biological replicates. (b) At high density, newborn cells are the smallest cells in the population (~29.2 *±* 0.5 μm^2^), with cells increasing in size during duplications (from ~35.8 *±* 2.1 μm^2^ at bud formation to ~40.8 *±* 2.3 μm^2^ at cytokinesis). Growing cells are on average 43.8 *±* 3.1 μm^2^. Finally, cells that die are the largest cells in the population (~59.1 *±* 1.6 μm^2^). (c) Similarly, in populations that were incubated in growth medium to which we added 250 μM GSH, we found that newborn cells are the smallest cells (~28.7 *±* 2.5 μm^2^) in the population, with duplicating cells increasing in their size during duplications (starting from ~37.6 *±* 1.3 μm^2^ and ending with ~38.3 *±* 2.0 μm^2^ at the end of division). Growing cells are on average 40.8 *±* 3.7 μm^2^. Finally, with GSH added, cell deaths occur at larger sizes (~70.1 *±* 1.8 μm^2^) compared to cells without added GSH. (d) In non-growing populations (low population density), the newborn cells are on average larger (~47.6 *±* 5.8 μm^2^) than the average size at which the cells start duplicating (~37.9 *±* 1.2 μm^2^), end duplicating (~43.7 *±* 0.4 μm^2^), or are growing (~36.8 *±* 1.9 μm^2^). Finally, cells that die are smaller (~55.3 *±* 5.0 μm^2^) than cells in the growing populations. Together, these results show that newborn cells are the smallest cells in the population. Moreover, the cell size increases monotonically during cell duplications with cells that die being the largest cells in the population. Furthermore, our observations suggest that adding extracellular GSH keeps the cell size approximately constant during cell duplications, but the cell size does increase when cells die. Finally, in low-densty (non-growing) populations without added GSH, the newborn cells are larger than the duplicating cells in the population.

**Figure S17:**
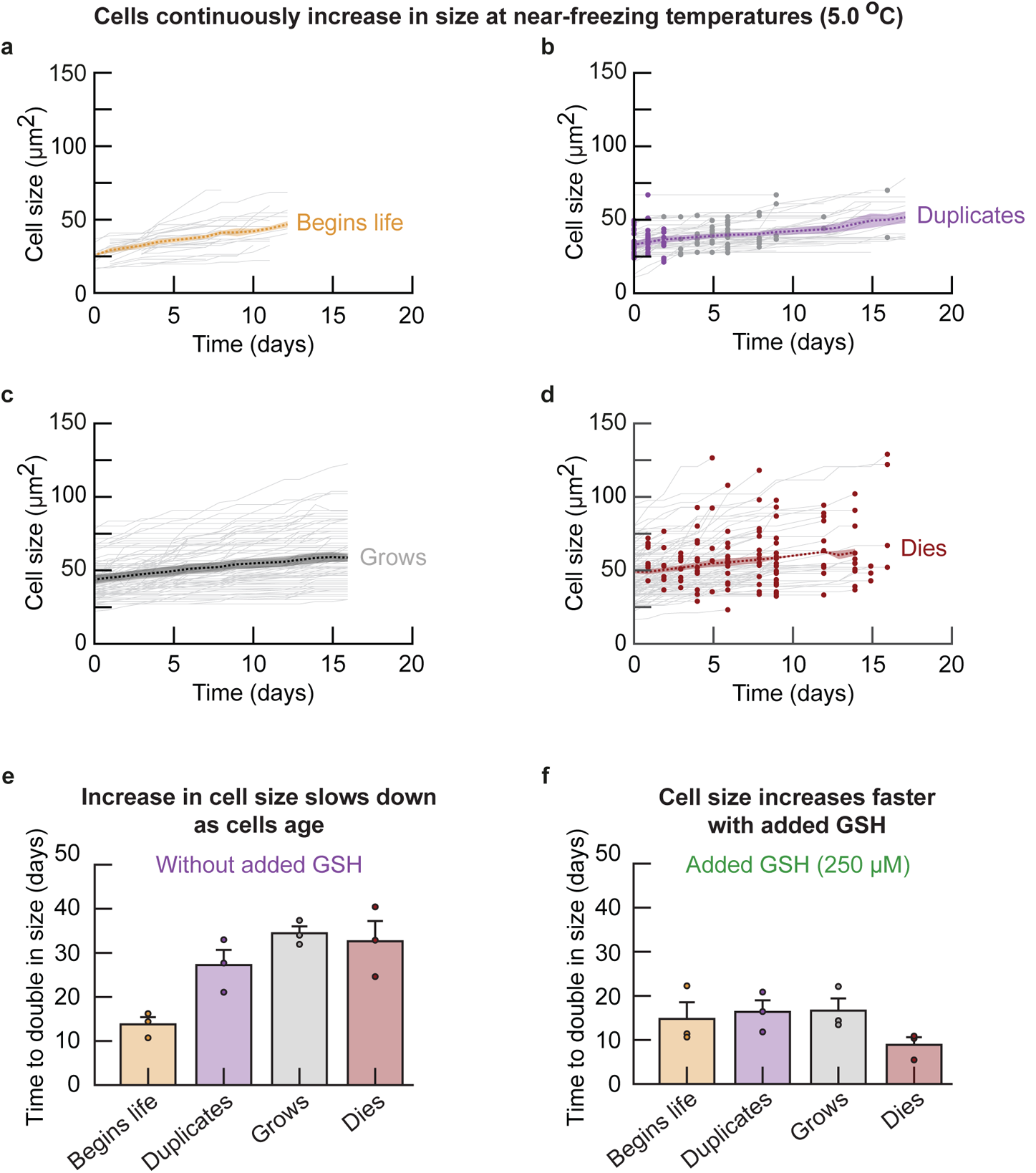
Cell size monotonically increases over time throughout a cell’s life at 5 °C (Related to Figure 3c). Our data shows that the cell size continuously increases at 5.0 °C starting with the smallest cell size at birth and ending with the largest cell size at death, with a continuous increase in size in between the two events (including during cell duplications, Supplementary Fig. S16). We used time-lapse microscopy to measure how the cell size changes in indiviual cells at 5.0 °C. **(a-d)** We used our time-lapse movies of cells in high-density populations (initially ~8, 000 cells / mL) to track the cell size over time in single cells and for all cells that were present at the start of the time-lapse movies. To reconstruct every stage of life for cells at near-freezing temperatures (e.g., 5.0 °C here and 1.0 °C shown in a later figure), we categorized all cells in the first time point of our movies into one of four classes (see Supplementary Fig. S16): newborn daughter cells (labelled, ”begins life”, just after cytokinesis), cells that start to duplicate (”duplicates”), cells that did not duplicate for the entire duration of the time-lapse (~17 days, ”grows”) and cells that died (”dies”). We assumed that the cells in the first time point of the movie are representative samples of populations at 5.0 °C. This is because the cell populations that we imaged were in a steady-state growth when we started the time-lapse of the cells, due to the fact that the population was already incubated at 5.0 °C for two weeks before the first frame of the movie. As an exception, since we did not know which cells were newborn cells when the time-lapse began (”day 0”), we used daughter cells that were newly born within a three-day period during our time-lapse movie. Shown here is the cell size over time for each class: size of a newborn cell **(a)**, size of a cell that begins to duplicate (i.e., when its bud begins to grow) **(b)**, size of a growing cell that does not duplicate **(c)**, and size of a cell just before it dies (almost always by bursting open **(d)**. Grey curves show the cell size over time for individual cells. Blue and grey dots in (b) indicate the time points at which bud formation and cytokinesis occurred respectively. Red dots in (d) indicate the cell size just before death. The dotted line shows the average size of cells each respective class for *n* = 3 biological duplicates, with the shaded area representing the s.e.m. of the average cell size from each biological replicate. Together, (a-d) show that the cells continuously increase their size at 5.0 °C. Almost all cells continuously increase in size during the entire duration of the time-lapse movies. Given that a newborn cell will either eventually duplicate or will grow without duplicating and that every cell will eventually die, these data together show that cells continuously increase in their size throughout their life at 5.0 °C. **(e-f)** From the measurements in (a-d), we determined the time that cells take to double in size without **(e)** and with 250 μM added GSH **(f)**. Shown here are the time taken to double in size for cells right after cytokinesis (yellow bars), cells that begin to duplicate (purple bars), cells that grow without duplicating (grey bars), and cells just before they die (red bars). Dots show the average time for each biological replicate. Error bars show the mean with s.e.m. of *n* = 3 biological replicates. These measurements show that, without added GSH, cells take increasingly more time to double in size. This time is the shortest for a newborn cells (13.8 *±* 1.6 days) and increases in the following order: cells that start to duplicate (27.3 *±* 3.4 days), cells that are growing without duplicating (34.4 *±* 1.6 days), and cells that are about to die (32.6 *±* 4.6 days). In contrast, with added GSH, cells take approximately constant time to double in size for newborn cells (14.8 *±* 3.8 days), cells that begin to duplicate (16.4 *±* 2.6 days), and cells that are growing without duplicating (16.6 *±* 2.8 days). Lastly, cells that die take the shortest time to double in size when there is 250 μM of added GSH (8.9 *±* 1.7 days). Together, (e-f) show that cells double in size more slowly as they (chronologically) age, and that adding GSH accelerates cell growth.

**Figure S18:**
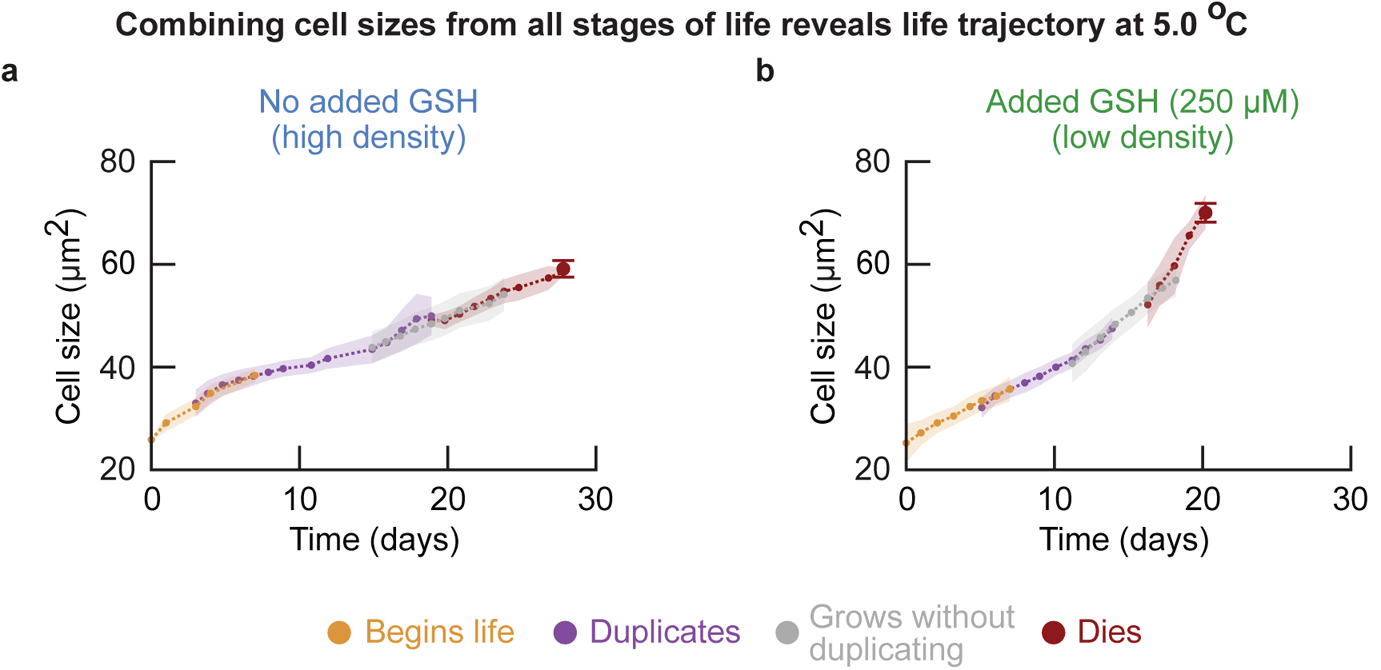
Continuous cell size growth from each life event reveals ”life trajectory” of cells at 5 °C (Related to Figure 3d). We have now established that cells continuously increase in size throughout their life at 5.0 °C (Supplementary Fig. S17). We also found that the cell size increases with consecutive life events (cells in ”begins life” are smaller than cells in ”duplicates”, followed by ”grows” and ”dies”, Supplementary Fig. S16). Moreover, since a newborn cell will either eventually duplicate or will grow without duplicating, and that every cell will eventually die, there exists a temporal order to these life events. We therefore used the cell size over time for each class to reconstruct the life of cells at 5.0 °C as function of cell size. Specifically, we stitched together the cell size curves of the average cell for each of the consecutive events (based on the order of average cell size: ”begins life”, ”duplicates”, ”grows” and ”dies”) by connecting each of the curves with its preceeding curve at the time that the cell sizes at the respective endpoints overlapped most. Shown here are the stitched curves for populations of cells that were incubated without **(a)** and with 250 μM added GSH **(b)**. Each curve overlaps with at least three data points with neighboring curves. The colors of each section indicates the separate life events, showing a cell that begins life (yellow bars), duplicates (purple bars), grows without duplicating (grey bars), and dies (red bars). Large red dot shows average size at just before cell death. Dots show the mean, with shaded areas representing the s.e.m. of *n* = 3 biological replicates.

**Figure S19:**
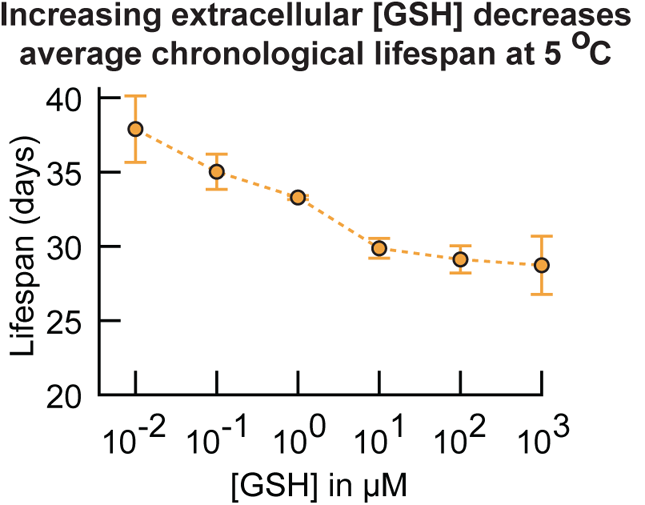
Average chronological lifespan decreases as the added GSH concentration increases at 5 °C (Related to Figure 3d). Cell’s average chronological lifespan (average chronological age at death) as a function of added extracellular GSH concentration. We incubated wild-type cell populations at 5.0 °C with various amounts of GSH added to the growth medium (all initially at ~5, 680 cells / mL). We took an aliquot of each liquid culture every week and measured the number of alive and dead cells using propidium iodide (see Methods). We then fitted these measurements to the same growth model as before to determine a cell’s average chronological lifespan which is plotted here (Supplementary Fig. S5, also see Supplementary Theory). The average chronological lifespan of cells at 5.0 °C decreases from ~37 days without added GSH to ~30 days with 100 μM added extracellular GSH. Extracellular GSH therefore decreases the chronological lifespan of cells at near-freezing temperatures. Error bars represent the mean with s.e.m., having *n* = 4 biological replicates per data point.

**Figure S20:**
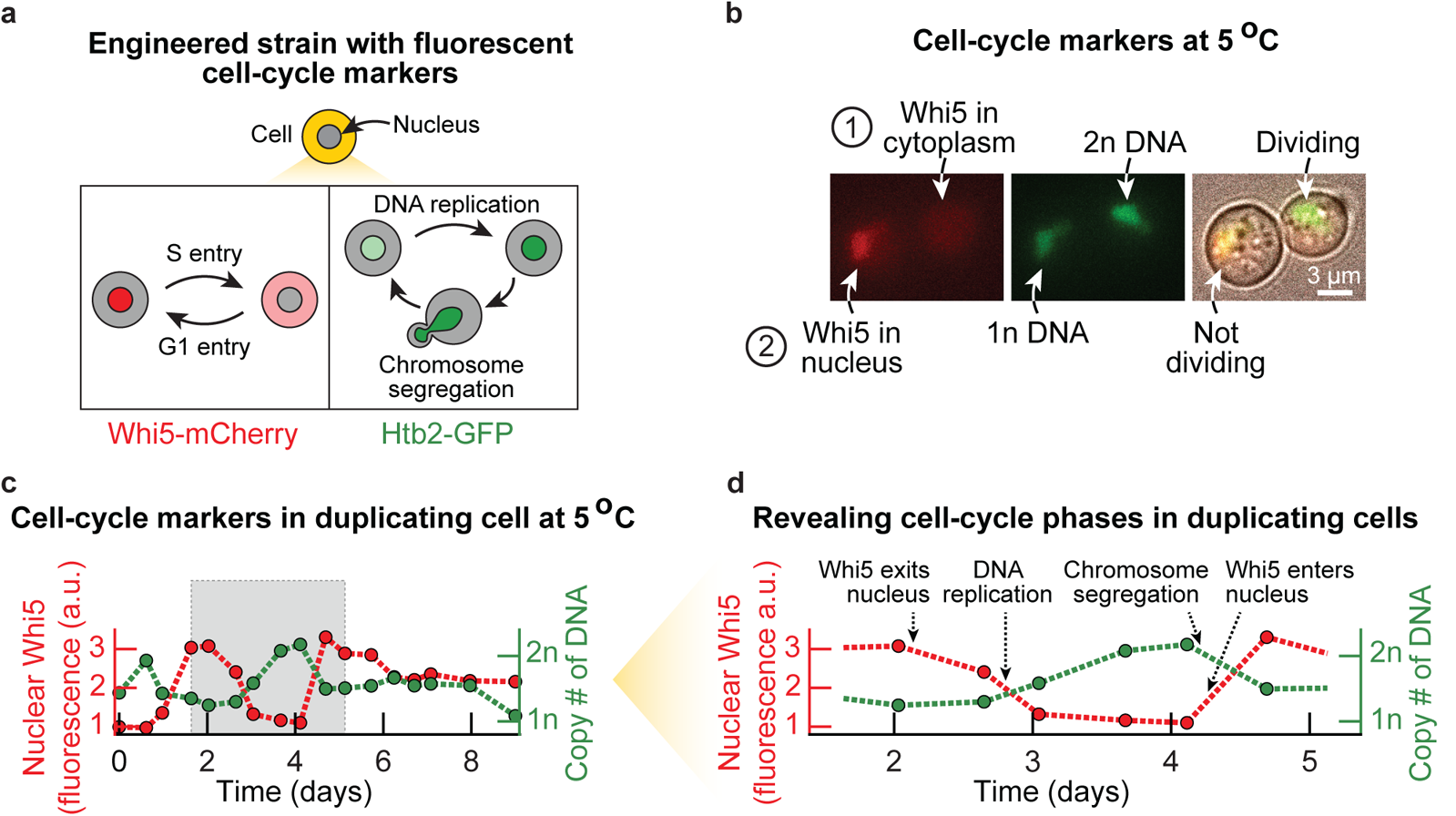
Time-lapse microscopy on engineered cells with fluorescent reporters for DNA replication (H2B histone fused to GFP) and G1-to-S transition (Whi5 protein fused to mCherry) reveals cell-cycle phases at 5 °C (Related to Figure 4). **(a)** Schematic of the engineered strain used for identifying when key cell-cycle phases occur in single cells at 5.0 °C. We engineered yeast to have the G1 transcriptional repressor Whi5 protein fused to mCherry protein (Whi5-mCherry), and the histone protein, H2B (Htb2), fused to GFP (H2B-GFP), also see Methods). Both Whi5 and H2B are well-known cell-cycle markers that indicate G1 phase (Whi5 translocates to the nucleus) and the S-G2-M phases (the amount of H2B is proportional to the amount of DNA) respectively [6,7,8]. We used this engineered strain to track the cell cycle at 5.0 °C. **(b)** Microscopy snapshots of the engineered strain. Nuclear fluorescence from Whi5-mCherry (red, left image) and H2B-GFP (green, middle) are shown for two representative cells at 5.0 °C. Also shown is the composite image that includes brightfield (right). Here, one cell is dividing which we can tell, aside from the brightfield image, by finding almost all of the Whi5-mCherry in the cytoplasm and a bright nucleus with H2B-GFP (contains 2n DNA, labelled ”1” in the top row). In this picture, we see that the other cell is not dividing, which we can also tell by finding almost all the Whi5-mCherry to be localized in the nucleus and a dim nucleus (contains 1n DNA, labelled cell ”2” in the bottom row). The dividing cell has a bud. Scale bar is 3 μm. **(c)** Nuclear Whi5-mCherry and H2B-GFP over time shown for a duplicating cell at 5.0 °C. This cell is from a population that was pre-incubated for two weeks at 5.0 °C before starting the timelapse, after which we tracked cells in these populations with a microscope (initially at ~6, 250 cells / mL). We took aliquots of our liquid cultures that we transferred to an imaging plate. We then kept this plate at 5.0 °C and imaged the cells every ~12 hours. Finally, we used this microscopy time-lapse to quantify the amount of nuclear Whi5-mCherry (left y-axis, red) and H2B-GFP (righ y-axis, green) over time. The grey area indicates a full cell-cycle, starting from G1 (all of Whi5-mCherry in the nucleus). Specifically, we used the fluorescence of H2B-GFP to locate the nucleus and quantify the amount of DNA in the cells. We first located the nucleus by segmenting the GFP fluorescence of each cell using a threshold GFP fluorescence that we kept fixed for all cells and time points. The nucleus was then the group of pixels whose fluorescences exceeded this threshold. To determine the copy number of DNA in a cell, we determined the total GFP fluorescence within the cell’s nucleus. We subsequently rescaled the nuclear GFP between the average minimum and maximum GFP fluorescence that we observed for duplicating cells. Thus, the amount of nuclear H2B was rescaled to a scale with ”1n DNA” (average GFP fluorescence in the nucleus of duplicating cells in G1) and ”2n DNA” (average GFP fluorescence in the nucleus of duplicating cells in G2). Finally, we used the fluorescence of Whi5-mCherry to quantify the relative amount of nuclear Whi5. To do so, we determined the average mCherry fluorescence in the nucleus and cytoplasm of each cell. The amount of nuclear Whi5 was then given by the ratio of measured nuclear and cytoplasmic mCherry fluorescence. **(d)** Nuclear Whi5 and H2B mark the cell-cycle phases of duplicating cells. Shown is a cell during one cell division (from the grey box of the duplicating cell in (c)). Dotted arrows indicate the end of G1 (Whi5-mCherry exits the nucleus), the S phase (the amount of H2B-GFP increases during replication of DNA), the M phase (H2B-GFP decreases during chromosome segregation) and the start of G1 (Whi5-mCherry begins to enter the nucleus).

**Figure S21:**
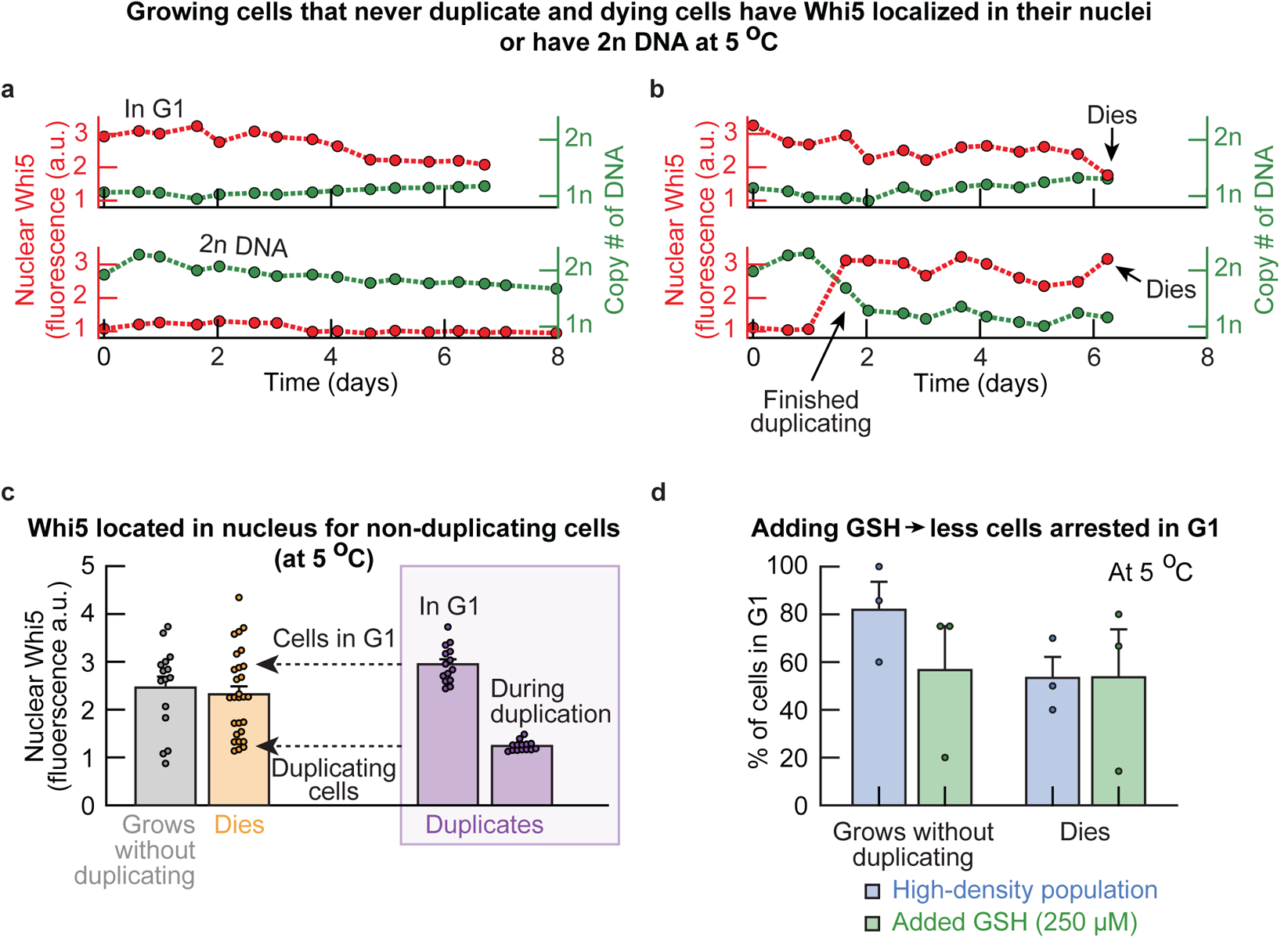
Growing cells that never duplicate until dying are arrested in G1 phase at 5 °C (Related to Figure 4c). **(a-b)** Nuclear Whi5-mCherry and H2B-GFP over time in growing cells that never duplicate until dying and dying cells at 5.0 °C (also see Supplementary Fig. S20). Shown are cells that will never duplicate in G1 or G2 phase **(a)**, and cells just before they die in G1 **(b)**. In (a-b), the amount of nuclear Whi5-mCherry and the amount of DNA remain constant in a growing cell that never duplicates during many days of incubation. **(c)** Cells that will never duplicate and dying cells often have Whi5 located in the nucleus. Shown is the average amount of nuclear Whi5-mCherry in non-duplicating cells (grey bar), in dying cells (orange bar) and for duplicating cells (purple box) when in G1 (left purple bar) or during cell division (S-G2-M phases, right purple bar). Data suggests that most cells that never duplicate stay in G1 (their Whi5-mCherry always stays in the nucleus, compared to the cells in G1 that will duplicate). The average amount of nuclear Whi5-mCherry in cells that will never duplicate was determined by averaging the amount of nuclear Whi5-mCherry for the entire duration of the time-lapse. The average amount of nuclear Whi5-mCherry in dying cells represents the amount of nuclear Whi5-mCherry averaged over the last three frames (~36 hours) before the cell dies. The average amounts of nuclear Whi5-mCherry for cells that can duplicate and are in G1 or are dividing were determined as follows. We determined when Whi5-mCherry was located in the cytoplasm (division) or the nucleus (G1). We then averaged the amount of nuclear Whi5-mCherry for each cell while the cell is either in G1 or duplicating (S-G2-M). Dots show raw data (*n* = 15 non-duplicating cells, *n* = 28 dying cells, *n* = 14 duplicating cells). Bars show the mean with s.e.m. of all cells. **(d)** Cells that never duplicate and dying cells mostly remain in G1. Shown here are the percentages of cells in G1 for high-density (growing) populations (blue bars) and low-density populations supplemented with 250 μM GSH (green bars). This plot shows that 80% of the cells that never duplicate remain in G1 until they die. Extracellular GSH decreases the percentage of these cells in G1. Whether a cell is in G1 or not was determined with the average amount of nuclear Whi5-mCherry (in (c)) and the amount of DNA in the cell (a cell was in G1 when the amount of nuclear Whi5-mCherry was above 1.66 and when the amount of DNA was below 2n). Bars show the mean with s.e.m., having *n* = 3 biological replicates per condition. Dots show raw data.

**Figure S22:**
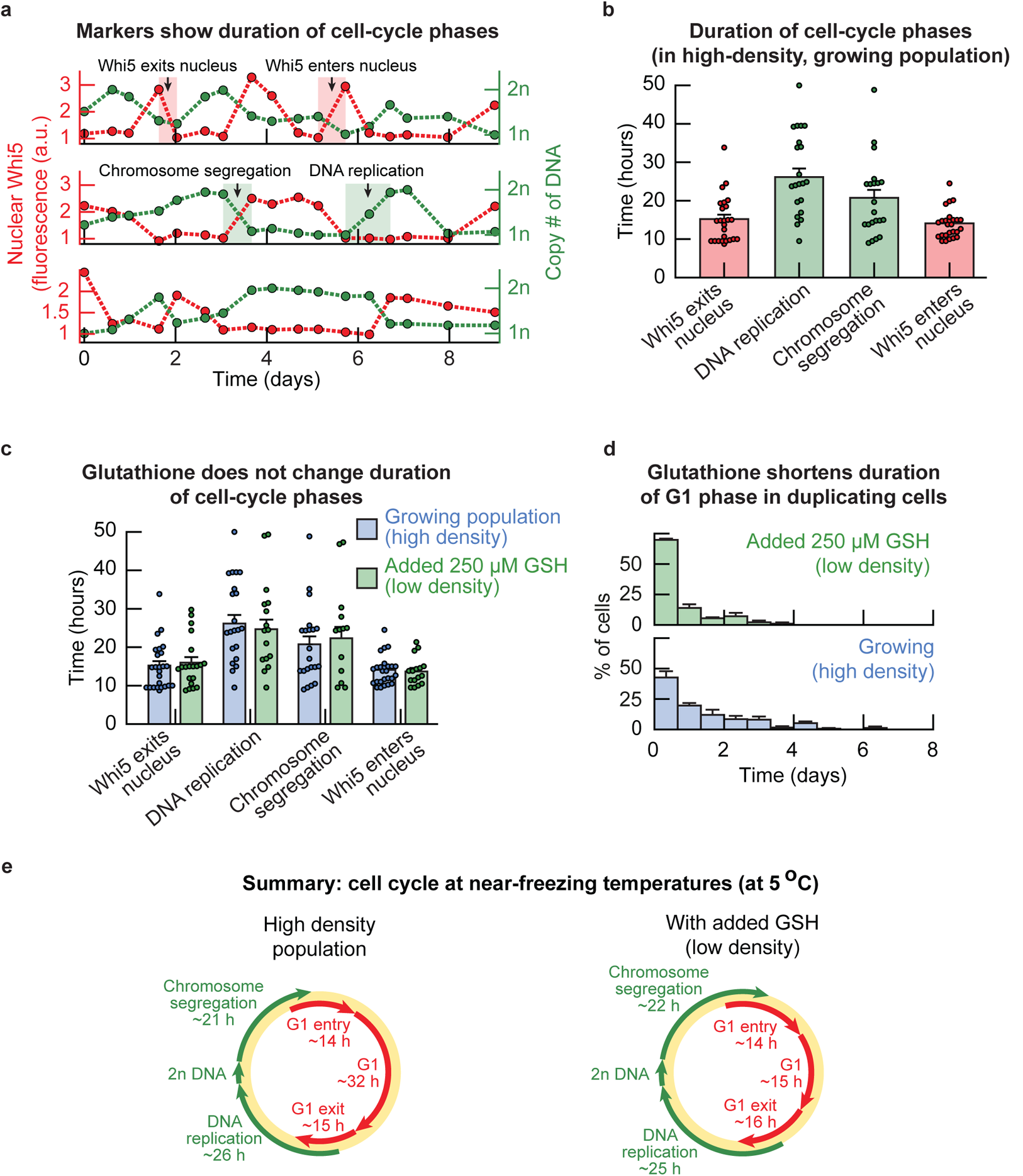
Glutathione shortens duration of G1 phase, but not S-G2-M phases, at 5 °C (Related to Figure 4c-d). **(a)** Nuclear Whi5-mCherry and H2B-GFP over time in duplicating cells that were incubated at 5.0 °C. Red shaded areas indicate examples of the duration of Whi5-mCherry exiting or entering the nucleus. Green shaded areas indicate examples of the duration of DNA replication or chromosome segregation. **(b)** Using time-lapse movies that examined the amounts of Whi5-mCherry and H2B-GFP in single cells, we determined the duration of cell-cycle events in duplicating cells (following the methodology in (a)). Shown here are the durations of Whi5-mCherry exiting the nucleus (15.2 *±* 1.1 hours, red bar), DNA replication (26.1 *±* 2.2 hours, green bar), chromosome segregation (20.8 *±* 2.1 hours, green bar), and Whi5-mCherry entering the nucleus (14.1 *±* 0.8 hours, red bar). Since the time-lapse movies consists of snapshots taken every ~12 hours, this ~12 hours is a lower bound on the true duration of each event. Dots show raw data, having *n* = 28 cells for the location of Whi5-mCherry, *n* = 23 cells with DNA replication and *n* = 24 cells with DNA segregation. Bars show the mean with s.e.m. of all cells. **(c)** Duration of cell-cycle events in cells from low-density populations that were incubated with 250 μM of GSH (green bars). Blue bars show the same data as in (b) for comparison. Shown here are, all with the added GSH, Whi5-mCherry exiting the nucleus (16.0 *±* 1.4 hours), DNA replication (24.7 *±* 2.5 hours), chromosome segregation (22.3 *±* 2.9 hours), and Whi5-mCherry entering the nucleus (13.7 *±* 0.9 hours). There is no reason to assume that the duration of any cell-cycle event changes upon addition of extracellular GSH (p-values are 0.69 for Whi5-mCherry exiting the nucleus, 0.69 for DNA replication, 0.66 for chromosome segregation and 0.72 for Whi5-mCherry entering the nucleus). Blue dots are as in (b). Green dots show raw data, having *n* = 16 cells for Whi5-mCherry entry and *n* = 20 cells for Whi5-mCherry exit, *n* = 17 cells with DNA replication and *n* = 16 cells with DNA segregation. Bars show the mean with s.e.m. of all cells. **(d)** Duration of G1 in duplicating cells from populations without (blue) and with 250 μM added GSH (green). The average duration of G1 was 32 *±* 3.0 hours without added GSH, and 15.0 *±* 1.5 hours with added GSH (median ~24 hours without and ~9 hours with added GSH). Thus, GSH shortens the duration of G1 in duplicating cells at near-freezing temperatures. With added GSH, the majority of G1 phases in duplicating cells take 12 hours or less. In contrast, the G1 phase in duplicating cells takes up to 6 days without added GSH. Error bar shows mean with s.e.m. of *n* = 3 biological replicates. **(e)** Summarizing (b-d). Shown are the durations of the cell cycle phases in duplicating cells in populations without (left) and with (right) added GSH. Length of arrows are proportional to the duration of each phase. Red arrows represent Whi5 (indicating events associated to G1), green arrows represent H2B (indicating events associated to S-G2-M). Together, (b-d) show that adding GSH to the growth medium shortens the duration of G1 in duplicating cells, while the duration of cell-cycle transitions remain unchanged upon incubation with GSH.

**Figure S23:**
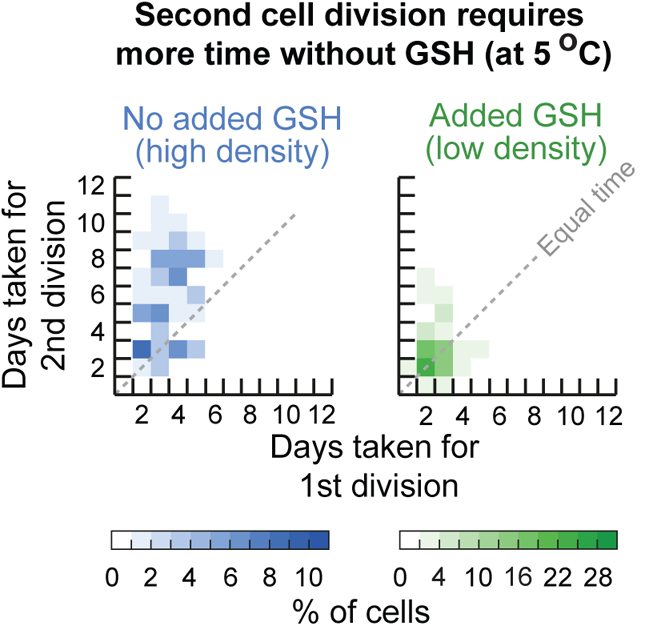
Consecutive divisions in individual cells slow down at 5 °C (Related to Figure 4d). We used the single-cell time-lapse movies at 5.0 °C to measure the doubling times of cells that divided multiple times. Heatmaps showing the relation of consecutive cell divisions of single cells at 5.0 °C without added GSH (blue heatmap, left) and with 250 μM added GSH (green heatmap, right). Shown is the number of days taken for the second cell division (y-axis) as function of the number of days taken for the first division (x-axis). Grey dotted diagonal indicates location where consecutive divisions in the same cell have equal doubling times. Data is aggregated from *n* = 3 biological replicates. These measurements show that the doubling time increases over time in populations without added GSH: the time of a next division is on average larger (above the diagonal) than the time of a preceeding division. With added GSH, the doubling time remains approximately constant for consecutive divisions.

**Figure S24:**
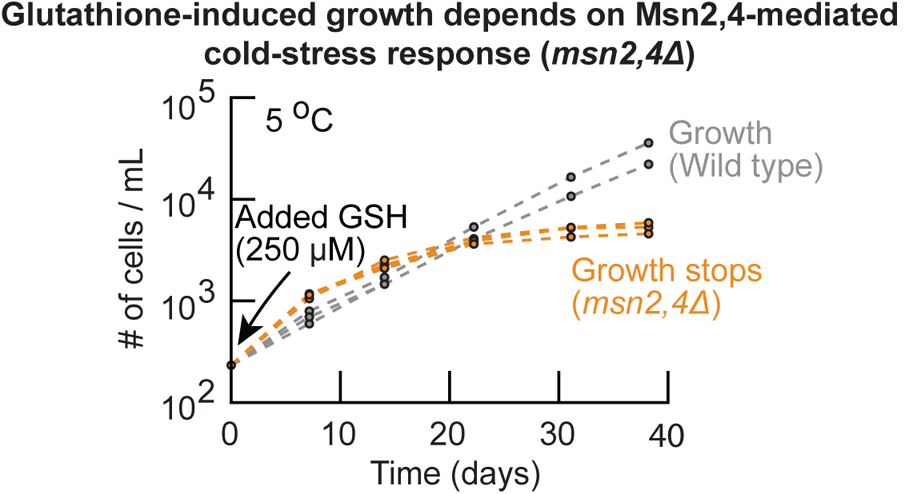
Glutathione-induced growth at near-freezing temperatures depends on Msn2,4-mediate cold-stress response (Related to Figure 4e). We constructed a mutant strain that lacks the major stress-response transcriptional activators, Msn2,4 (*msn2,4*Δ strain) [9,10,11]. Shown here is the population density over time for the *msn2,4*Δ strain at 5.0 °C with 250 μM added GSH (orange curves, initially ~250 cells / mL). Grey curves show the wild-type strain for comparison (initially ~250 cells / mL with 250 μM added GSH). The *msn2,4*Δ strain stops growing after two weeks of incubation with added GSH at 5.0 °C whereas the wild-type strain grows exponentially over time. We already established that GSH removes intracellular ROS and thereby induces the wild-type population to grow (Fig. 2c). However, the *msn2,4*Δ population stops growing at 5.0 °C even with the added GSH. This means that yeasts incubated with extracellular GSH must still experience a low-temperature (cold) stress, despite the added GSH greatly reducing the amount of intracellular ROS that inhibits cell duplications. Together, our results suggest that glutathione-induced population growths at 5.0 °C depend on the Msn2,4 cold-stress response. All colors show *n* = 3 biological replicates.

**Figure S25:**
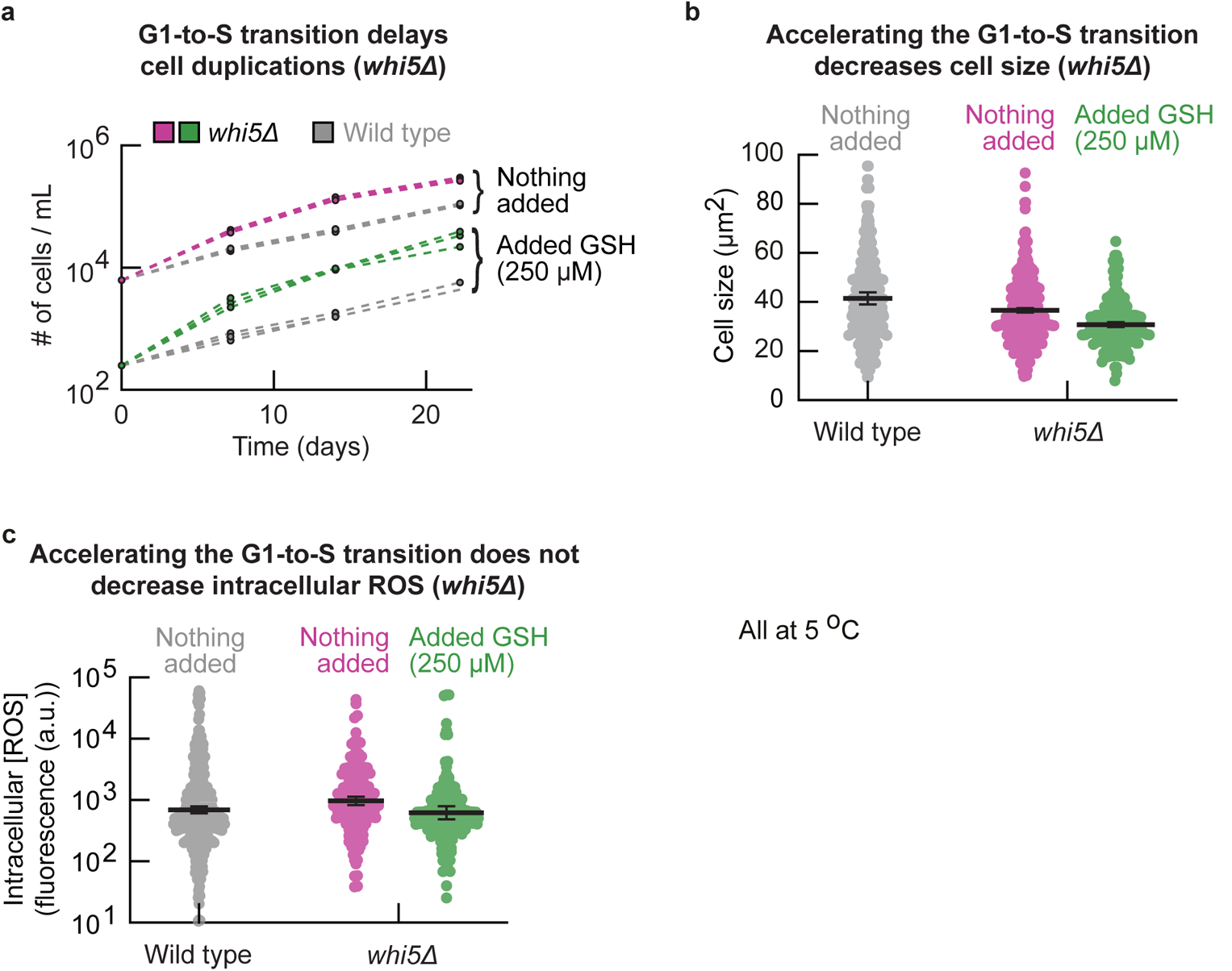
G1-to-S transition delays cell duplications at 5 °C (Related to Figure 4e). **(a)** We constructed a mutant strain (*whi5*Δ strain) whose *WHI5* was knocked out and therefore has an accelerated G1-to-S transition [12]. Shown here are the population densities over time for the *whi5*Δ strain that was incubated at 5.0 °C either without (pink curves, initially ~6, 250 cells / mL) or with 250 μM added GSH (green curves, initially ~250 cells / mL). Both conditions are compared with the wild-type strain (grey curves). The *whi5*Δ strain grows faster at 5.0 °C (3.4 *±* 0.1-fold difference in density after two weeks without added GSH, 5.7 *±* 0.3-fold difference in density with added GSH). Thus, knocking out *WHI5* accelerates population growth, suggesting that the G1-to-S transition delays cell duplications. Each color and initial density shows *n* = 3 replicate populations. **(b)** Cell size of the *whi5*Δ strain. Shown here are the cell sizes of the *whi5*Δ strain in populations without (pink dots) or with 250 added μM GSH (green dots). Wild-type strain without added GSH is shown for comparison (grey). Cell sizes were measured after two weeks of incubation at 5.0 °C. The *whi5*Δ strain has smaller cell size compared to the wild type, with adding extracellular GSH reducing the cell size further. This suggests that accelerating the G1-to-S transition decreases the cell size at 5.0 °C(cells spend less time in the G1 (growth) phase). Error bars show mean with s.e.m., having *n* = 3 biological replicates. **(c)** Intracellular ROS concentration in the *whi5*Δ strain after two weeks of incubation at 5.0 °C (see Methods). Shown here are populations that were incubated either without (pink dots) or with 250 μM added GSH (green dots). Grey dots show wild-type without added GSH for comparison. The *whi5*Δ strain does not have less intracellular ROS compared to the wild-type strain. Error bars show mean with s.e.m., having *n* = 3 biological replicates. Together, (a-c) show that accelerating the G1-to-S transition at 5.0 °C accelerates cell duplications and decreases cell size. Researchers found similar effects at 30.0 °C [12,13]. Simultaneously, accelerating the G1-to-S transition does not decrease intracellular ROS abundance. This suggests that cells do not accumulate intracellular ROS during G1, since shortening G1 does not decrease the intracellular ROS concentration.

**Figure S26:**
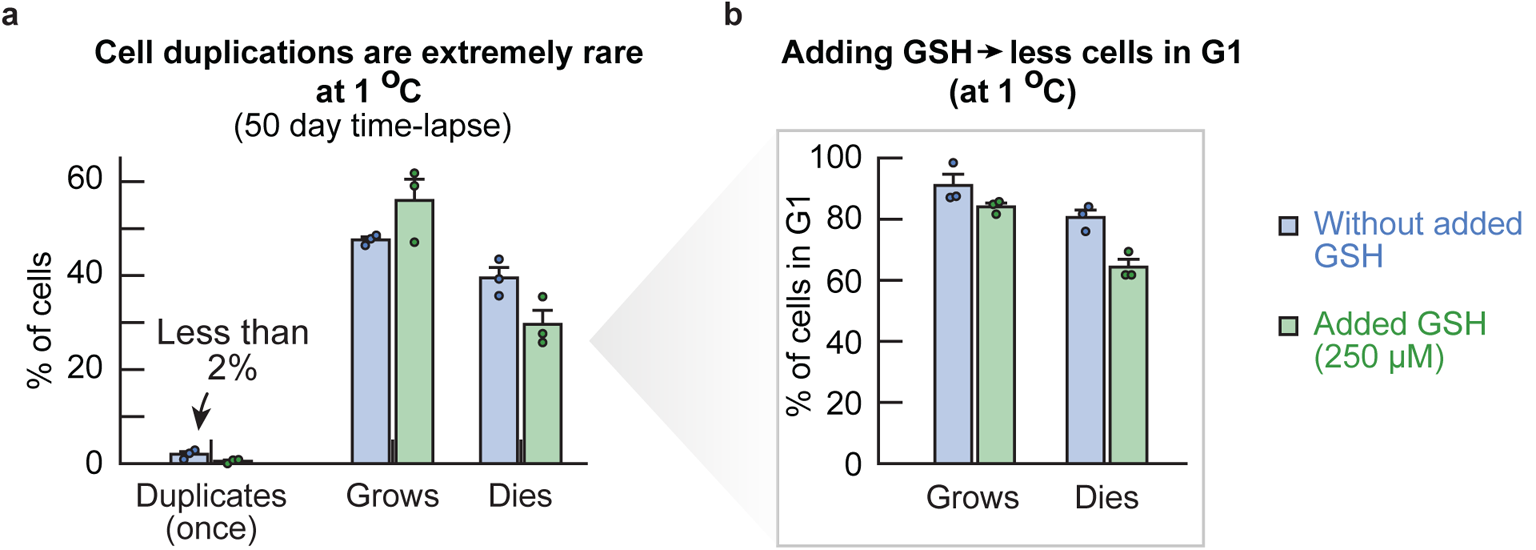
Cells at 1 °C are arrested in G1 and less than 2% of cells proceeds to duplicate during a 51-day time-lapse (Related to Figure 5a). **(a)** Percentage of cells that duplicates, grows and dies during a 51 day microscope time-lapse at 1.0 °C. Bars show populations with (blue bars) and without (green bars) 250 μM added GSH. Less than 2% of cells duplicated once (no cell duplicated more than once). **(b)** Percentage of cells that grows without duplicating or dies and that are in G1 for the duration of the time-lapse. Almost all cells that grow without duplicating or die are in G1. Similar to our measurements at 5.0 °C, we found that adding GSH decreases the percentage of cells that are in G1 (Supplementary Fig. S21).

**Figure S27:**
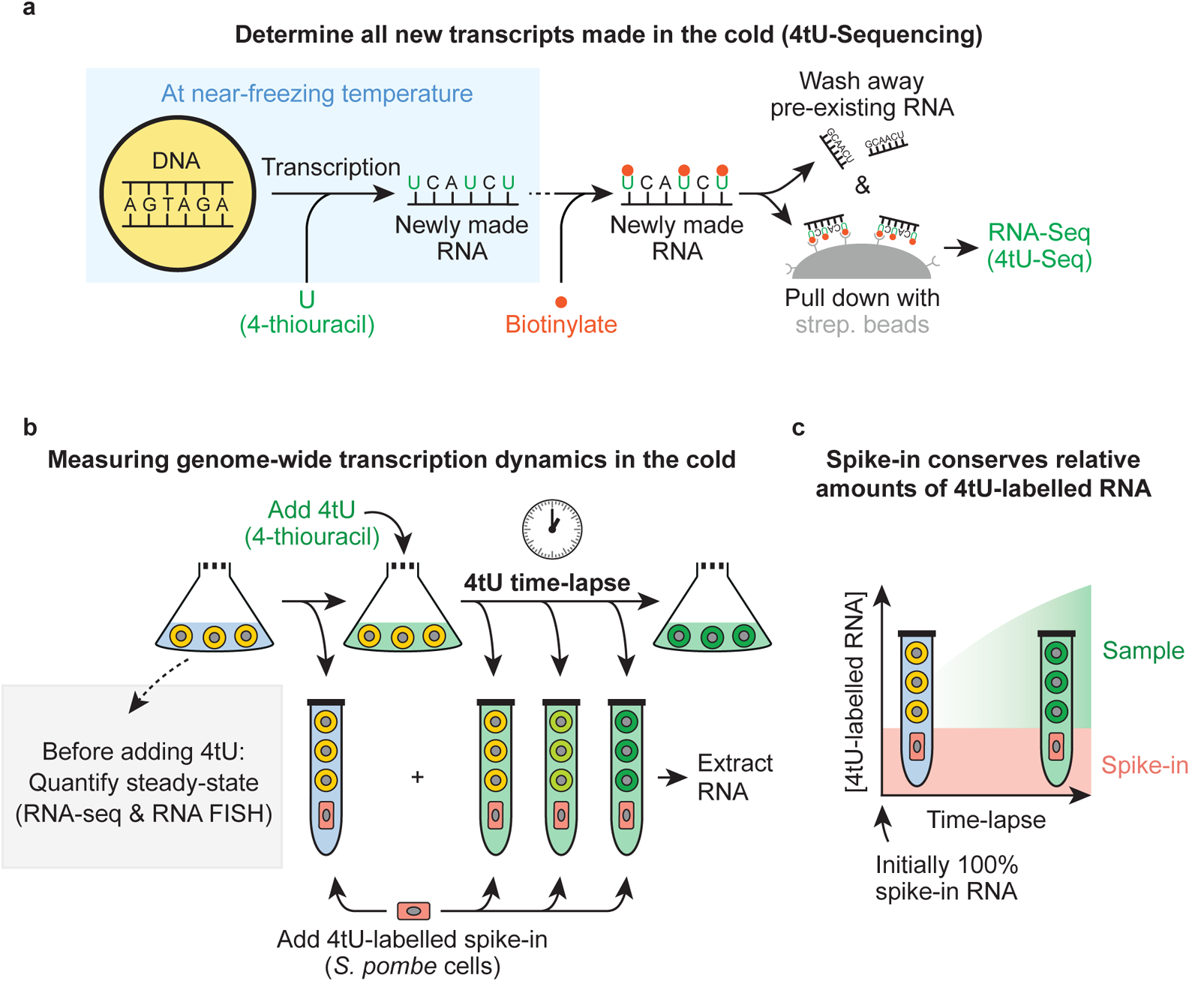
Measuring the genome-wide transcription at near-freezing temperatures (Related to Figure 5b). **(a)** Schematic demonstrating the experimental procedure for performing 4tU-sequencing on newly-made transcripts in populations of *S. cerevisiae* [14,15,16]. A synthetic uracil analog (the nucleotide 4-thiouracil, 4tU) is added to the growth medium of cells. Cells incorporate the 4tU into their newly synthesized RNA. We collect cells after the desired time of incubation with 4tU and extract the total RNA. The newly synthesized RNA is linked to biotin by specifically biotinylating the 4tU (”4tU-labelled” RNA). Simultaneously, the pre-existing RNA that does not contain 4tU is not biotinylated. Finally, the 4tU-labelled RNA is separated from the pre-existing RNA without 4tU through a pull-down with magnetic beads containing streptavidin, after which the purified 4tU-labelled RNA is sequenced (see Methods for experimental details). **(b)** Schematic illustrating the experimental procedure for measuring the genome-wide transcription dynamics at near-freezing temperatures. We incubated liquid cultures with populations of wild-type yeasts at the desired temperature (e.g., 5.0 °C). After two weeks of incubation, we first collected two aliquots of our cultures (as ”time 0” hours of the time-lapse). Directly after, we supplemented the growth medum of the cultures with 4tU at a final 5 mM concentration, and collected aliquots from our cultures at the desired time-points (see Methods for experimental details). After collecting the time-lapse samples, we added a fixed amount of 4*tU*-labelled *S. pombe* (*Schizosaccharomyces pombe*) cells to all 4tU time-lapse samples as a spike-in and proceeded with RNA-extraction. Finally, we used one of the ”time 0” hours aliquots to quantify the steady-state transcript levels (grey box, RNA-seq and RNA FISH). **(c)** The *S. pombe* spike-in that was added to all 4tU time-lapse samples enabled us to normalize the amount of 4tU-labelled transcripts from different time-points. Initially, all 4tU-labelled RNA in the sample is from the spike-in since the cold-incubated cells have not initially synthesized 4tU-labelled RNA. Over time, the cold-incubated cells synthesize new 4tU-labelled RNA and degrade old (unlabelled) RNA, increasing the percentage of 4tU-labelled RNA that is from the cold-incubated cells. Normalizing with the amount of spike-in transcripts therefore ensures that the relative differences between samples in a time-lapse are conserved (since the amount of 4tU-labelled RNA from *S. pombe* cell was the same in every sample). Moreover, this procedure eliminates experimental differences between samples due to sequencing and processing of RNA.

**Figure S28:**
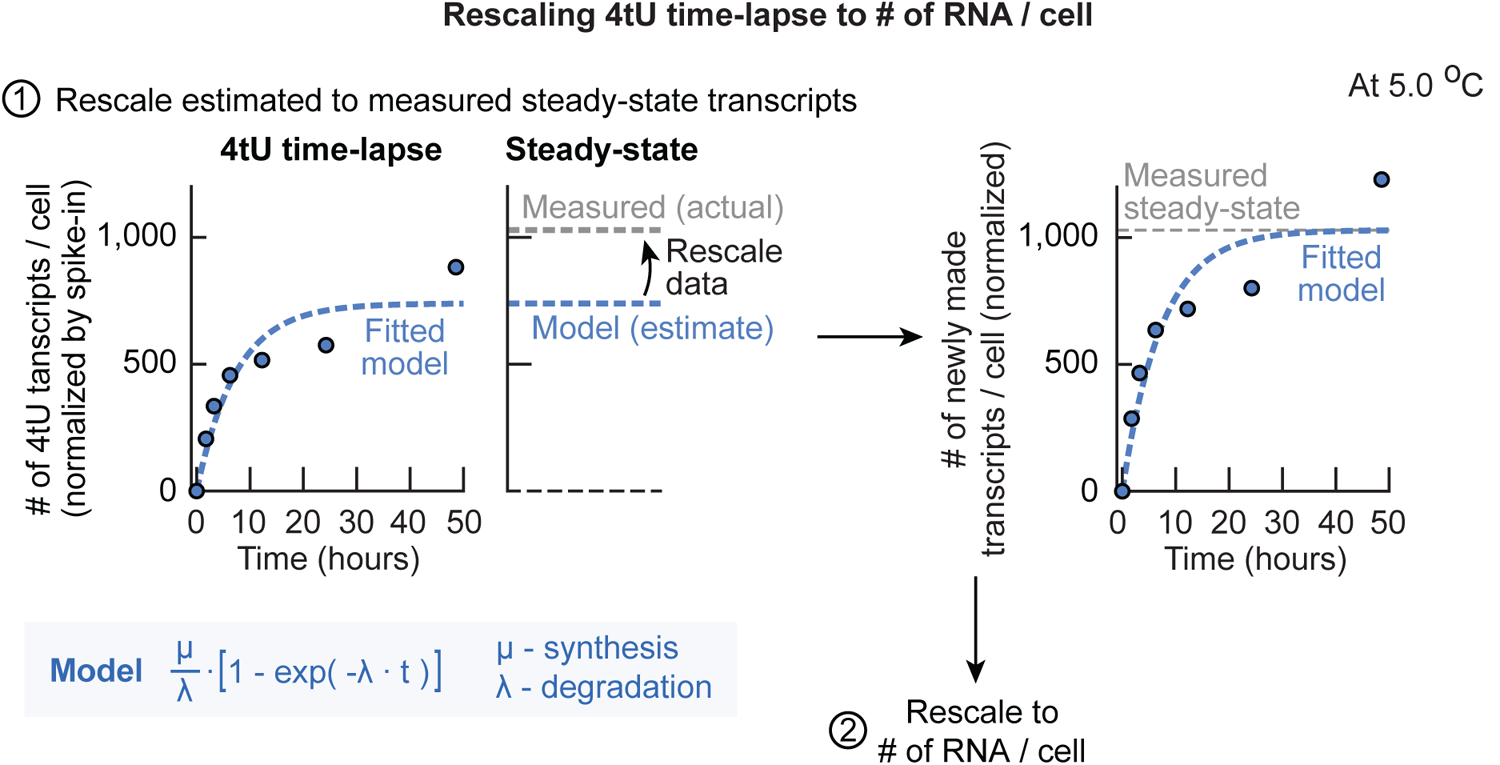
Procedure for rescaling the 4tU time-lapses to # of RNA / cell using steady-state transcript levels (Related to Figure 5b). First, the 4tU time-lapse samples were normalized by the number of spike-in transcripts and the number of alive cells in the population, giving the relative amount of 4tU transcripts per cell over time (amount of 4tU-labelled RNA normalized by 4tU spike-in). Next, these relative values are rescaled to obtain the ”# of RNA / cell”. To do so, the samples are rescaled such that steady-state amount of RNA / cell that is predicted by the 4tU time-lapse is equal to the actual steady-state amount of RNA / cell that we measured through RNA-seq. First, we fitted a mathematical model to the amount of 4tU transcripts / cell. The conventional model to describe RNA synthesis assumes that RNA is synthesized at a constant rate and degraded at a rate that is dependent on the concentration. Thus, the kinetics of 4tU-labelled RNA are given by 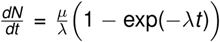 with a RNA synthesis rate *µ* (in ”# of RNA / cell / hour”), a RNA degradation rate *λ* (in ”per hour”) and time *t*. Over time the relative amount of 4tU transcripts per cell converges to a steady-state *µ/λ* (in ”cell / hour”) as *t → ∞* by degradation of old RNA and synthesis of new (4tU-labelled) RNA [17,16]. To rescale the steady-state amount of RNA / cell that is predicted by model to the actual steady-state amount of RNA / cell, we had also sequenced the steady-state transcript levels (total RNA, not 4tU purified RNA) of each time-lapse together with a *S. pombe* spike-in (total RNA) to normalize our time-lapses across temperatures (the steady-state transcript levels can differ per temperature, and the same spike-in RNA was used for all steady-state samples). Thus, we rescaled all 4tU-samples such that the predicted steady-state of the model matched the measured steady-state transcript levels. Shown is a representative example of the rescaling from a 4tU time-lapse without added extracellular GSH at 5.0 °C. Note that this rescaling conserves the relative differences between time points within each time-lapse. Together, the previous normalizations resulted in the relative number of newly-made transcripts (4tU labelled RNA) over time between replicates and across temperatures, and that the predicted steady-state of these time-lapses is equal to the measured steady-state of each time-lapse. As a last step, the relative number of newly-made transcripts / cell are rescaled to the actual integer number of RNA per cell. Dots show raw (normalized) data for one replicate. Blue dotted lines shows model fit for that replicate. Grey dotted lines shows measured steady state values.

**Figure S29:**
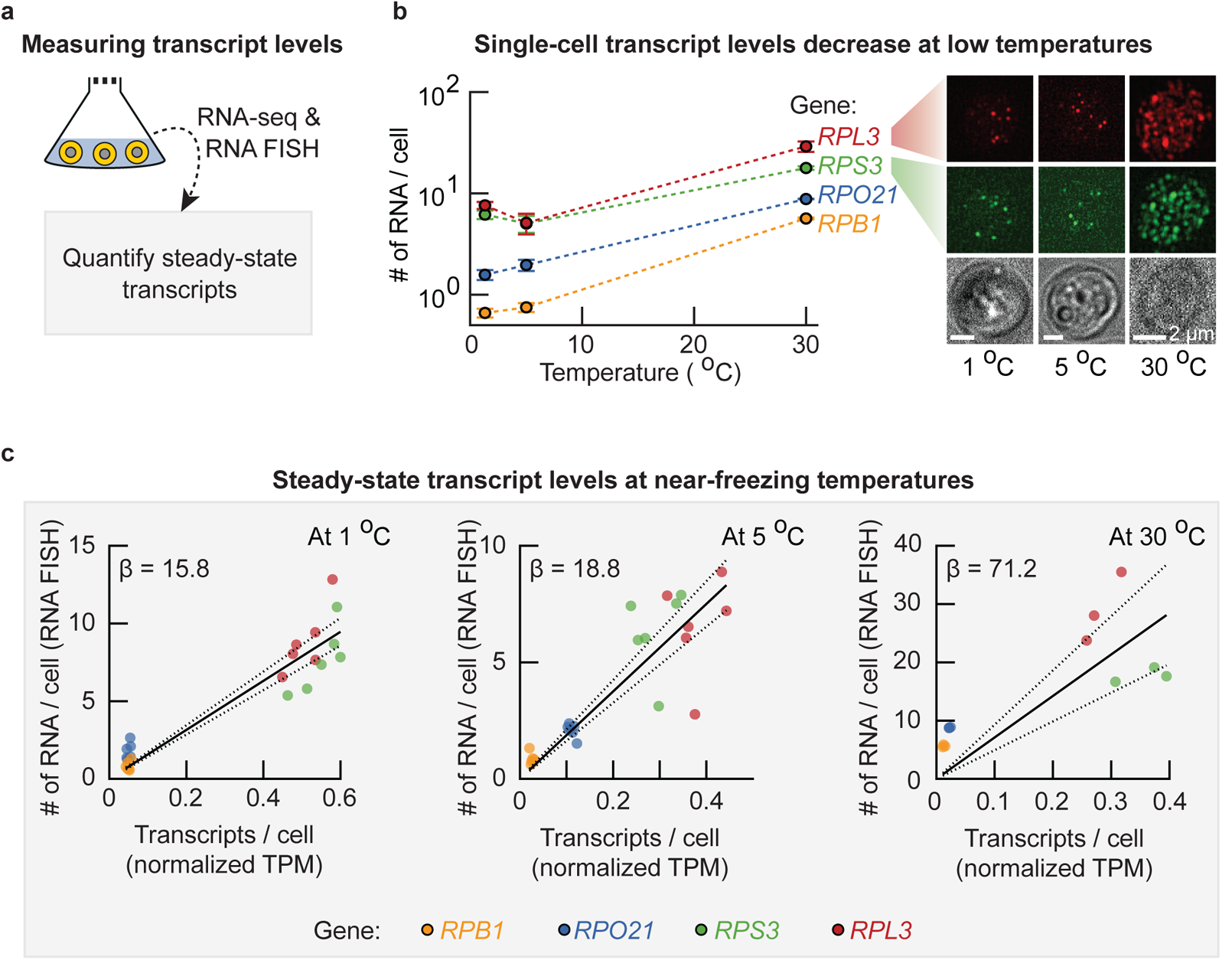
Steady-state transcript levels decrease with temperature (Related to Figure 5b). **(a)** Steady-state transcript levels were quantified in two ways. In one way, we performed regular RNA-seq on the steady-state RNA (normalized across temperatures by a spike-in of total RNA from *S. pombe*). In the another way, we quantified the number of mRNA per cell via single-molecule RNA FISH on cells from the same aliquots on which we performed RNA-seq. These measurements reveal the steady-state transcript levels across temperatures, and can be combined to rescale the number of transcripts / cell (from RNA-seq) to the ”# of RNA / cell” as measured by RNA FISH. **(b)** The single-cell transcript levels of several endogenous yeast genes were quantified at 30 °C, 5.0 °Cand 1.0 °C. Shown is the average number of mRNA per cell as function of temperature for *RPL3* (red curve), *RPS3* (green curve), *RPO21* (blue curve) and *RPB1* (yellow curve curve). The average number of mRNA per cell decreases with temperature. Each data point shows the average of *n* = 3 biological replicates, in total having counted at least *n* = 616 cells (for *RPB1* and *RPO21*) or at least *n* = 421 cells (for *RPL3* and *RPS3*). Images show representative examples of labelled RNA in single-cells at 1.0 °C (left), 5.0 °C (middle) and 30 °C (right). Dots are *RPL3* RNA (red fluorescence, top) and *RPS3* RNA (green fluorescence, bottom). Brightfield images are shown as comparison. Scale bar is 2 μm. **(c)** The average number of RNA per cell (from RNA FISH) as function of the relative steady-state amount of transcripts per cell (from RNA-seq, normalized with the *S. pombe* RNA spike-in) at 1.0 °C (left panel), 5.0 °C (middle panel) and 30 °C (right panel). Shown are the steady-state levels for *RPB1* (yellow dots), *RPO21* (blue dots), *RPS3* (green dots) and *RPL3* (red dots). Black solid line shows linear fit without intercept, dotted lines show 95% confidence interval of fit. Since both RNA FISH and RNA-seq measurements come from the same sample, these linear fits yield a scaling factor to convert the relative amount of transcripts per cell to ”# of RNA / cell” (*β* = 15.8 at 1.0 °C (pearson-correlation coefficient *ρ* = 0.94), *β* = 18.8 at 5.0 °C (*ρ* = 0.87) and *β* = 71.2 at 30 °C (*ρ* = 0.79)). All colors show 6 dots, having *n* = 3 biological replicates for with and without added GSH added.

**Figure S30:**
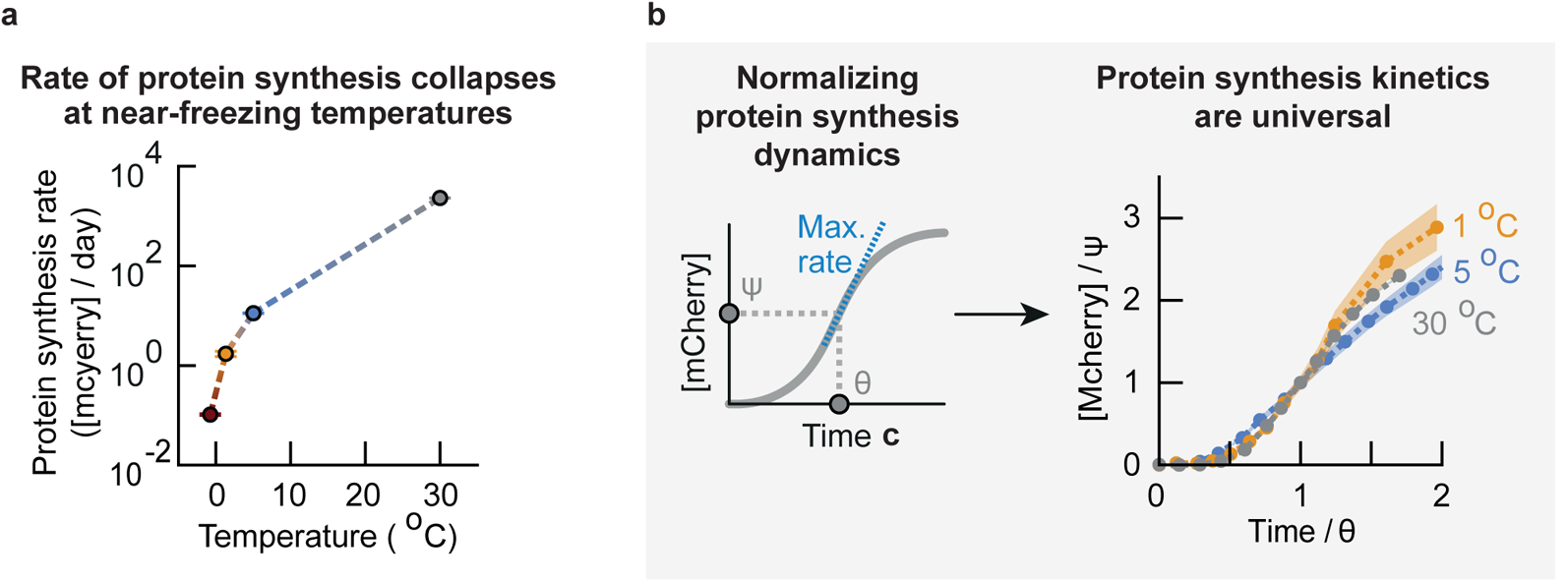
Protein synthesis dynamics are universal with the protein synthesis rate collapsing around 0 °C (Related to Figure 5c). We build a synthetic gene-circuit in our wild-type yeast that makes the cells produce a red fluorescent protein (mCherry) upon induction by galactose. Expression of mCherry is controlled by an inducible promoter, *pGAL10*, which is activated by galactose. We first grow the yeast in raffinose and then add galactose to the raffinose-medium to activate the expression of mCherry. **(a)** For each curve in Fig. 5c we determined the maximum rate of mCherry expression (steepest slope in each curve, also see Supplementary Theory). Shown are the protein synthesis rates as function of temperature (the synthesis rate of general proteins can be at least as fast as the synthesis rate of mCherry measured here). **(b)** Protein synthesis dynamics across temperatures. *Left* : Schematics of relevant parameters. At each temperature, we used the characteristic time Θ and fluorescence Ψ at which the population reaches its maximum rate of protein synthesis to rescale the fluorescence curves. *Right* : Renormalized protein synthesis curves from Fig. 5c. The fluorescence was normalized with the characteristic fluorescence Ψ and time was rescaled with the characteristic time Θ for each temperature. The renormalized curves collapses onto a single master curve. These renormalized curves suggest that one only needs to know the characteristic fluorescence-scale Ψ and time-scale Θ for each temperature to fully describe expression of mCherry at that temperature. In both panels the error bars and shaded areas show the mean with s.e.m., having *n* = 3 replicate populations.

**Figure S31:**
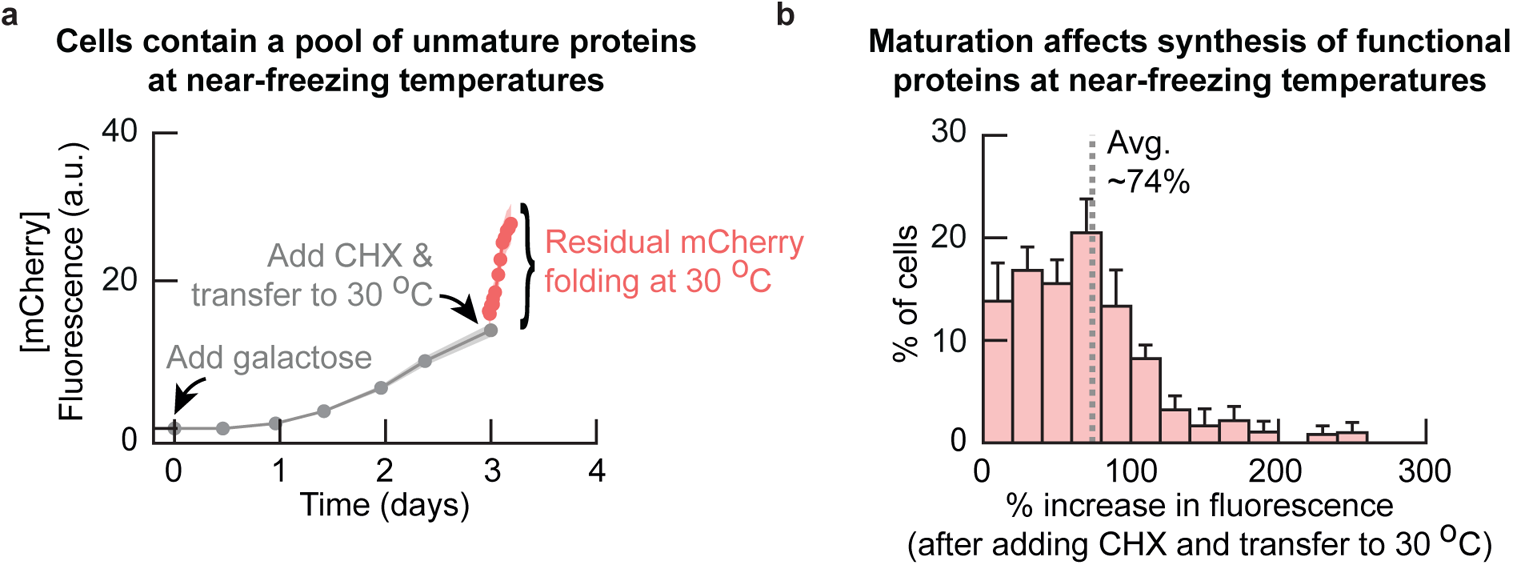
Time to synthesize functional protein affected by maturation (e.g., folding) time at 5 °C (Related to Figure 5c). Using the mCherry-inducible strain to quantify processes affecting time to synthesize functinoal proteins at near-freezing temperatures (also see Supplementary Fig. S30). **(a)** We incubated cells of the inducible mCherry strain for two weeks at 5.0 °C. We then added galactose to the growth media and followed the expression of mcherry at 5 °C in single cells over time with a microscope (see Methods). After three days of incubation with galactose, we added cycloheximide (CHX) to the growth medium at a final 100 μg/mL concentration to instantly stop translation in the cells. We then transferred the cells to 30 °C and continued to measure the mCherry fluorescence of the cells to quantify the maturation (e.g., folding) of residual mCherry. Shown is the fluorescence of mCherry over time at 5 ^°^C (grey curve), and the increase in fluorescence at 30 °C after addition of CHX (red curve). The average fluorescence considerably increases after addition of CHX. Cells therefore contain a pool of unmatured mCherry at 5 °C. Dots show average fluorescence, and shaded area shows s.e.m. of fluorescence from the averages in *n* = 5 populations. **(b)** Histogram shows the increase in fluorescence of single cells from (a) after addition of CHX and transfer to 30 °C. The average fluorescence increases by 74% after addition of CHX, showing that roughly half of mCherry remains unmatured during gene expression. Synthesis of functional proteins is therefore affected by the time required for maturation (e.g., folding) at near-freezing temperatures (also see Supplementary Theory). Histogram contains *n* = 108 cells, error bar shows s.e.m. of *n* = 5 populations.

**Figure S32:**
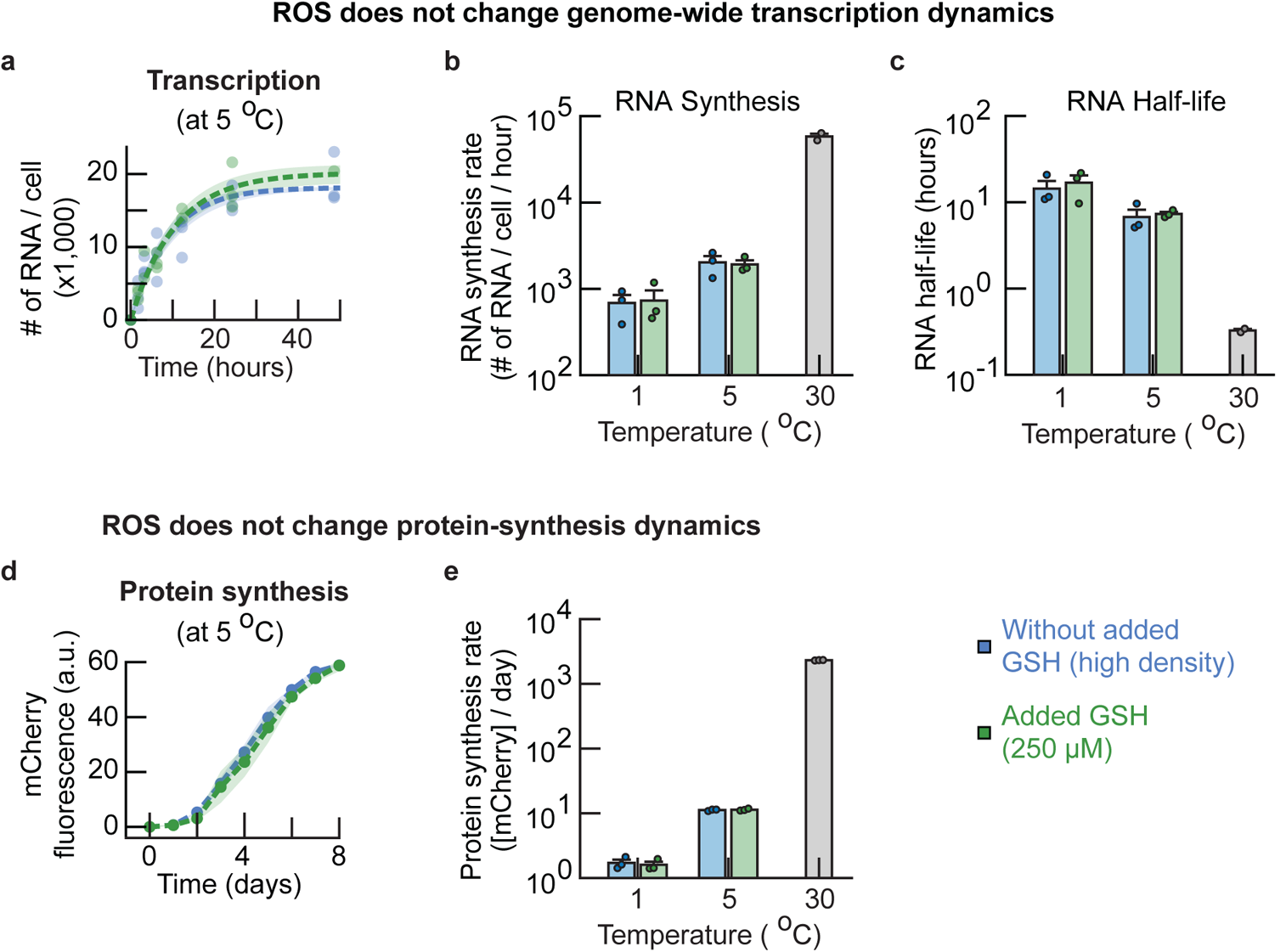
ROS does not change genome-wide transcription or protein synthesis dynamics at near-freezing temperatures (Related to Figure 5d). **(a-c)** ROS does not change rates of genome-wide transcription. **(a)** We used the number of synthesized RNA over time to fit a model for the kinetics of RNA synthesis (also see Supplementary Fig. S28). Shown are the number of newly synthesized RNA per cell over time at 5.0 °C for populations with (blue curve) or without (green curve) 250 μM added GSH. Dots show raw measurements. Dotted line represents kinetics model using average parameter estimates, with n = 3 biological replicates per condition. Shaded area represents the s.e.m. for fitted model parameters. **(b-c)** RNA synthesis rates **(b)** and RNA half-lives **(c)** at 1.0 °C, 5.0 °C as quantified from the fitted model. Included are the rates for populations without (blue bars) and with 250 μM added GSH (green bars). Rates at 30 °C are shown as a comparison (grey bars). The RNA half-lives and RNA synthesis rates decrease with temperature. Without added GSH, the steady-state transcript levels (in # of RNA / cell) are 12, 900 *±* 900 (1.0 °C), 18, 300 *±* 800 (5.0 °C) and 27, 300 *±* 1, 300 (30 °C), the synthesis-rates (in # of RNA / cell / hour) are 690 *±* 160 (1.0 °C), 2020 *±* 370 (5.0 °C) and 58, 100 *±* 5, 500 (30 °C), and the half-lives are (in hours) 14.4 *±* 3.9 (1.0 °C), 6.7 *±* 1.8 (5.0 °C) and 20 *±* 1 (30 °C, in minutes). There is no reason to assume that the RNA synthesis rates are different upon addition of GSH (p-value is 0.82 at 5.0 °C and 0.89 at 1.0 °C). Error bars show mean with s.e.m., having *n* = 3 biological replicates per condition at 1.0 °C and 5.0 °C, and *n* = 2 biological replicates at 30 ^°^C. Dots show raw data. **(d-e)** ROS does not change rate of protein synthesis. **(d)** The synthesis of mCherry over time at 5.0 °C compared for populations without (blue curve) and with (green curve) added 250 μM GSH (also see Supplementary Fig. S30). The dots show average fluorescence, with shaded representing the s.e.m. of n = 3 biological replicates. **(e)** Protein synthesis rates at 5.0 °C and 1.0 °C for populations that were incubated without (blue bars) or with (green bars) 250 μM added GSH. Grey bar shows synthesis rate of mCherry at 30 °C as comparison. The protein synthesis rate decreases with temperature. Without added GSH, the protein synthesis rates (in mCherry fluorescence / day (a.u.)) are 1.7 *±* 0.2 (1.0 °C), 11.2 *±* 0.2 (5.0 °C) and 2314 *±* 10 (30 °C). There is no reason to assume that the protein synthesis rates are different upon addition of GSH (p-value is 0.66 at 5.0 °C and 0.69 at 1.0 °C).

**Figure S33:**
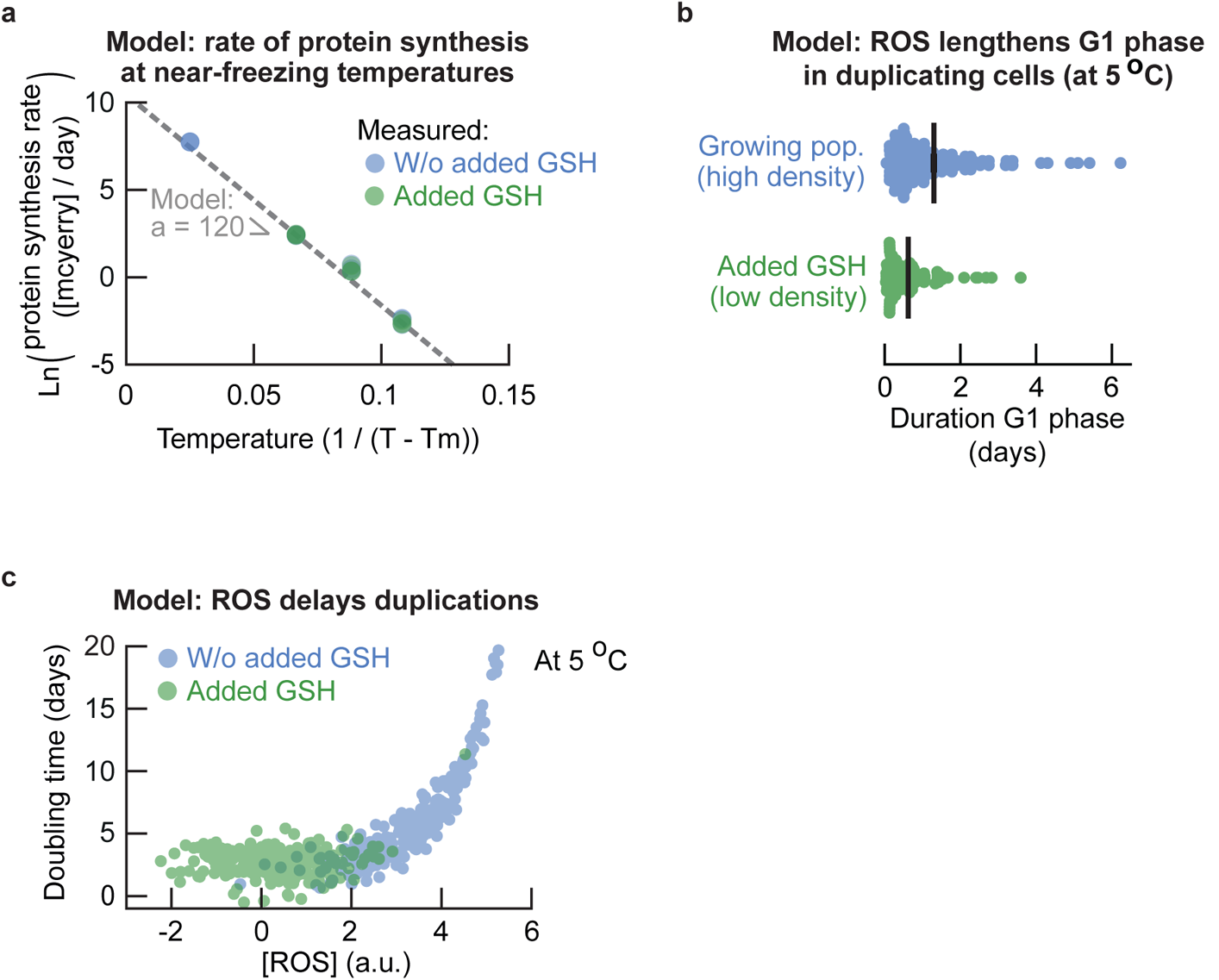
Stochastic model for single-cell duplications at near-freezing temperatures (Related to Figure 6). **(a)** Fitting the protein synthesis rate *r_g_*(*T*) as function of temperature *T* using an Arrhenius-type equation (ln *r_g_*(*T*) *∝ −a/*(*T −T_m_*), see equation S11 and Supplementary Theory for derivation). This model assumes a minimum temperature *T_m_* = *−*10 °C below which protein synthesis does not occur. Fitting a linear model yields the parameter *a ≈* 120.72 that fully describes the protein synthesis rate as function of temperature. **(b)** Model simulation of ROS-dependent G1 duration in single cells at 5.0 °C (equal to the delay in the model, compare with Fig. 4c). The only free model parameter (intracellular ROS concentration) was chosen such that the average delay equals the average measured G1 duration for each condition respectively (Fig. 4c, see Supplementary Theory, parameter *R* in equation S16). This yields *R* = 2.4 for high-density (growing) populations and *R* = 1.55 for low-density populations with added GSH. Error bar shows mean with s.e.m. of *n* = 3 replicate simulations that each have *n* = 50 cells. Dots show raw single-cell delays. **(c)** Model simulation of ROS-dependent doubling times in single cells at 5.0 ^°^C without (blue dots) and with added GSH (green dots). Initially, for relatively low ROS concentrations, the ROS-dependent duration is negligible compared to the minimum doubling time that is set by the protein synthesis rate. Thus, at low ROS concentrations (green dots), the doubling time is approximated by the minimum doubling time with noise (see Supplementary Theory, equation S14). For relatively high ROS concentrations, the ROS-dependent duration becomes of the same size as the minimum doubling time, such that the doubling time becomes dictated by ROS abundance. Each condition shows a simulation of *n* = 250 cells. The free parameter of the model was chosen to be equal to the values used to simulate the single-cell doubling times (Fig. 6c, *R* = 3.2 without added GSH, *R* = 0.35 with added GSH).

## Supplementary Text

### Density effect emerges from the percentage of duplicating and dying cells (related to Fig. 1)

Here we establish that the density-dependent growth at 5.0 °C does not emerge from the density-independent single-cell doubling times at 5.0 °C but instead from the percentage of cells in the population that dies and the percentage of cells that duplicates. This is illustrated through only considering the initial population of cells in the time-lapse. For example, in high-density (growing) populations, ~29% of the population at the start of the time-lapse duplicates with a doubling time of ~6.5 days. These 29% of cells can therefore duplicate at least 3x during the ~20 day time-lapse, thereby expanding the initial population with 88% newborn cells. During the same ~20 days, ~36% of the population at the start of the time-lapse dies. Thus, the number of cells that die within the high-density population at the start of the time-lapse are more than replenished by the number of newborn cells from the cells at the start of the time-lapse. This leads to the high-density populations growing over time. In contrast, in low-density (non-growing) populations, ~17% of the population at the start of the time-lapse duplicates with a doubling time of ~7.1 days. The duplicating cells in non-growing populations can therefore duplicate less than 3x during the ~20 day time-lapse. The duplicating cells then expanded the initial population with at most 52% newborn cells, while during the same time ~58% of the initial cells dies. Then the number of cells that die within the low-density population at the start of the time-lapse exceeds the number of newborn cells from the cells at the start of the time-lapse. As a consequence, more cells die than there are newborns per unit time, leading to the low-density population to go extinct. In summary, the density effect at 5.0 °C emerges from the percentage of cells that dies and the percentage of cells that duplicates depending on initial population density.

### Growth model for expected lifespan and doubling time (related to Fig. 1)

Here we derive a simple growth model that describes the expected lifespan and doubling time of cells at near-freezing temperatures. Assume that the number of cells in the population is given by *N*(*t*) at time *t*. Of these *N*(*t*) cells, *A*(*t*) are alive and *D*(*t*) are dead, such that *N*(*t*) = *A*(*t*) + *D*(*t*). The number of alive cells changes over time through cell duplications and cell deaths. The rate at which cells duplicate is given by *µ* and the rate at which cells die is given by *λ*. Then, the number of alive cells changes over time according to,

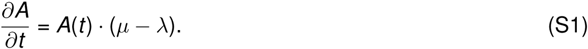

Similarly, the total number of cells changes according to,

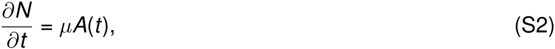

since we add freshly born cells to the total number of cells and dead cells are not removed from the total number of cells in the population. Solving the differential equation S1 yields,

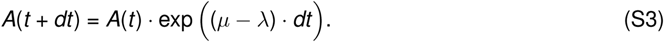

Using equation S3 to solve equation S2 we obtain,

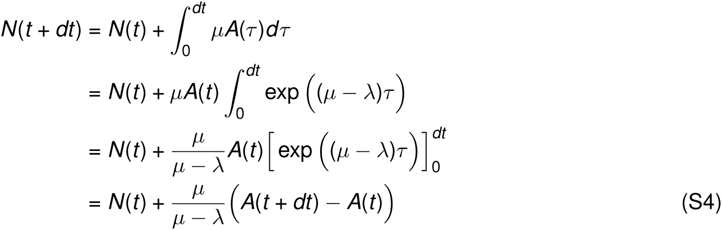

In our experiments we can measure the number of alive cells and the number of dead cells and the total number of cells over time. We therefore have *A*(*t*) and *N*(*t*) available to fit the growth rate *µ* and death rate *λ* with our model. Taking the logarithm of equation S3 describing the number of alive cells in the population and rewriting yields,

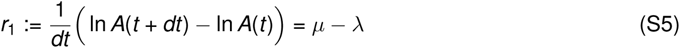

Similarly, we can rewrite equation S4 describing the total number of cells in the population as,

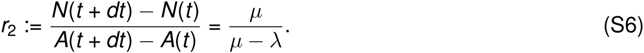

Finally, we use the numbers *r*_1_ (from equation S5) and *r*_2_ (from equation S6) to find the rates *µ* and *λ*,

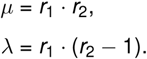

We used the above equations together with equations S5 and S6 to estimate the growth rate *µ* and death rate *λ* from the total number of cells and the number of alive cells in a population.

### Model for protein synthesis rate (Related to Fig. 5)

Here we describe a simple kinetics model for the synthesis of mCherry that we measured experimentally by following the mCherry fluorescence in cells over time. Assume that mCherry is synthesized by first being translated into an unfolded polypeptide chain, followed by folding into a fluorescent protein (Supplementary Fig. S31). We let *u* denote the amount of unfolded mCherry and *f* denote the amount of folded mCherry in a cell. We further assume that unfolded mCherry is synthesized through translation at a constant rate *m* (a.u. / day), and that the unfolded mCherry is folded at a rate *γ* (day*^−^*^1^). Then the amounts of unfolded and folded mCherry over time *t* are described by,

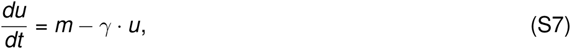

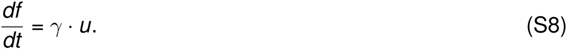

Solving equation S7 yields 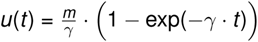, and substitution of *u*(*t*) into equation S8 yields,

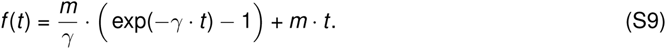

equation S9 describes the amount of folded mCherry over time and provides a model for the experimentally measured mCherry fluorescence in cells. Initially translation of mCherry has not occured, such that a cell does not have unfolded mCherry and no folded mCherry is being produced. At time *t* = 0 the cell starts synthesizing unfolded mCherry, whose concentration is described by 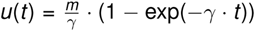. Thus, the amount of unfolded mCherry initially increases by translation and eventually reaches a steady-state that is determined by an inflow from translation and an outflow to folded mCherry (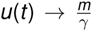 as *t → ∞*). At this steady-state, the equation S7 for unfolded mCherry satisfies 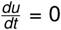, such that *γ · u* = *m* and 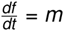 (from equation S8). Thus, this derivation shows that the slope of mCherry fluorescence over time converges to the translation rate *m* (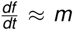 for sufficiently large *t*). In practice, we measured the synthesis rate *m*(*T*) for each temperature *T* by taking the maximum slope of the increase of mCherry fluorescence over time, and then used these *m*(*T*) to further fit the model in equations S7 to the data.

## Stochastic model for the speed limits of single-cell doubling times at near-freezing temperatures

In this section we derive a stochastic model with one free parameter for single-cell doubling times at near-freezing temperatures. The model is derived using our experimental observations. First, we motivate a description of the doubling time in terms of a ROS-independent lower limit. This lower limit is set by the protein synthesis rate, while the doubling time increases above this minimum through an additional time that is ROS-dependent. Using our measurements, we motivate a power-law type scaling between the minimum doubling time and the protein synthesis rate. Next, we describe the protein synthesis rate as function of temperature, such that also the minimum doubling time becomes a function of temperature. We use our experimental data to derive the stochastic, ROS-dependent time above the minimum doubling time. Combining the ROS-independent minimum time and the ROS-dependent additional time then yields the complete stochastic model. Finally, we fit all parts of the model to our experimental data to find the values of all model parameters.

### Doubling time is limited by the protein synthesis rate and ROS-dependent delay

Experimentally we have found that removing ROS with GSH shortens the doubling time at a given temperature (Supplementary Fig. S11), and that the durations of the S-G2-M phases of the cell cycle are independent of ROS (durations of S-G2-M do not change upon addition of GSH, Supplementary Fig. S22). These observations suggest that removing ROS by adding GSH decreases the doubling time to a minimum value for a given temperature (compare Fig. 2e with Fig. 4c-d). Thus, we assume that there is a ROS-independent minimum doubling time for each temperature, and that cells duplicate almost as fast as that minimum doubling time when ROS is removed. Additionally, we have found that removing ROS by adding GSH shortens G1 duration (Fig. 4c). Furthermore, cells with high intracellular ROS concentrations are unlikely to duplicate (Fig. 3b) and are often arrested in G1 (Supplementary Fig. S21). These results suggest that G1 duration is ROS-dependent, which yields an additional time that increases the doubling time above the ROS-independent minimum. Together, these experimental observations suggest that cell duplications can be split into a ROS-independent part (minimum doubling time) and a ROS-dependent part (additional time). We therefore assume that the doubling time *τ* of single cells at a temperature *T* is given by a ROS-independent time *t* and an additional duration Δ*t* such that,

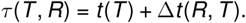

where *R* is some environmental parameter that depends on, for example, ROS and extracellular GSH. Here Δ*t >* 0 such that the doubling time is at least the minimum time for a given temperature (*tau*(*T*, *R*) *> t* (*T*)).

### Modeling temperature dependence of the protein synthesis rate

Recall that the protein synthesis rate decreases as function of temperature (Supplementary Fig. S30). To describe the protein synthesis as function of temperature with a model we assume a lower temperature limit *T_m_*below which no protein synthesis is possible. Then we can describe a rate as function of temperature with a common Arrhenius-type model *r_g_*(*T*) = *c ·* exp(*−a/*(*T − T_m_*)) for some parameter *a* and minimum temperature *T_m_*, such that *r_g_*(*T*) *→* 0 as the temperature decreases to *T_m_* (the Arrhenius equation would have *T_m_* = *−*273 °C). We can rewrite this model using the experimentally measured protein synthesis rate at a known temperature *T*_0_,

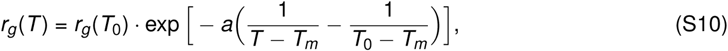

providing a simple equation that describes the protein synthesis rate as function of temperature. Taking the logarithm of equation S10 we find that,

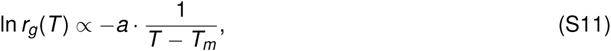

which is a model that is linear in 1*/*(*T − T_m_*) and that we fit to the measured protein synthesis rates at various temperatures to find the slope *a*.

### Model of minimum time limited by the protein synthesis rate

We have experimentally measured both the minimum time *t* (*T*) by taking away ROS with GSH (Supplementary Fig. S11). We have also measured the ROS-independent protein synthesis rates (Fig. 5d, Supplementary Fig. S32). Then both the minimum time *t* (*T*) and protein synthesis rate are independent of ROS, and we assume without loss of generality that the minimum time *t* (*T*) is limited by the protein synthesis rate *r_g_*(*T*) at each temperature *T* (i.e., cell division requires protein synthesis, so that the timescale of cell division must be longer than the timescale of protein synthesis). Here, we seek to describe the relation between the minimum doubling time and the protein synthesis rate.

Experimentally we have found that the observed doubling time slows down with the protein synthesis rate via a power-law type scaling (Fig. 6b). One reason why the relation may be of power-law type is the following. We assume that the rate of each process depends on the temperature according to an Arrhenius-type equation 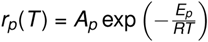, where *r_p_*(*T*) is the rate of the process *p* (protein synthesis or cell division) dependent on and constants *E_p_*, *R* and *A_p_*. Isolating the reciprocal temperature gives 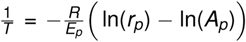. Denoting the rate of cell division as *r_d_* (*T*) and the rate of protein synthesis with *r_g_*(*T*), we can express the rate of cell division in terms of the rate of protein synthesis (temperature dependence of the rates omitted for readability),

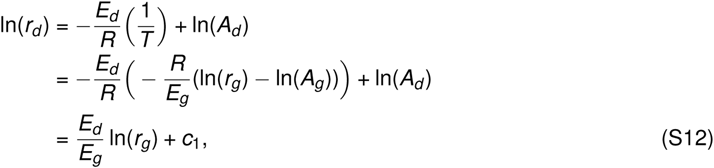

with *c*_1_ a constant of aggregated rest terms. Let *k* = *E_d_ /E_g_* be some factor such that by taking the exponent of equation S12 we obtain, 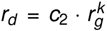. Then, by substituting the minimum time *t* (*T*) for the growth rate via *t* (*T*) = ln(2)*/r_d_* (*T*) we obtain,

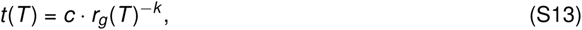

for some constant *c* and exponent *k*. Note that the above derivation is not restricted to the choice for a temperature scale. Taking the logarithm of equation S13 we find that,

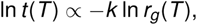

which is a simple linear model that we fit to the measured minimum times and protein synthesis rates at various temperatures to find the value of the exponent *k*. The power law with exponent *k* in equation S13 also illustrates how the doubling time scales with protein synthesis. For example, if the protein synthesis rate changes by two-fold, then the doubling time changes by 2*^−k^*-fold, such that fold-change in the doubling time is smaller than the fold-change in protein synthesis for exponents *k <* 1. Of note, it is only the fold-change of the protein synthesis rate that sets how the doubling time changes, irrespective of their actual values (the doubling time and protein synthesis rate change proportionally).

In summary, with an Arrhenius-type equation describing rates as function of temperature we can derive that cell division and protein synthesis may follow a power-law relation. We have therefore now experimentally measured and theoretically motivated a power-law type scaling of the minimum time *t* (*T*) and the protein synthesis rate. Finally, using equation S13, we rewrite the minimum time *t* (*T*) using a minimum observed time *t* (*T*_0_) at a known temperature *T*_0_,

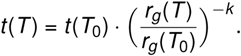

We know from our single-cell data that the experimentally measured minimum doubling time is stochastic and approximately normally distributed (e.g., the single-cell doubling times of the wild type at 5.0 °C with added GSH, see Supplementary Fig. S11). We therefore assume that *t* (*T*_0_) is stochastic and equal to the experimentally measured minimum doubling time *τ* (*T*_0_) plus some normally distributed noise *ɛ_m_ ~ N*(0, 1), such that,

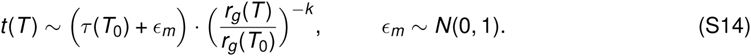

We then use the *t* (*T*) for *ɛ_m_* = 0 as the deterministic minimum doubling time *t*_min_(*T*) from the model. The minimum doubling time is thereby the same for all cells in a population at a given temeperature, and only changes proportionally to the protein synthesis rate as temperature changes. Cell-to-cell variability due to other factors is simulated with the normally distributed noise.

### Stochastic model of the ROS-dependent duration of the doubling time

Given the ROS-independent description of the minimum doubling time (equation S14), we here seek to describe the ROS-dependent additional time Δ*t* (*R*, *T*) that increases the doubling time above this minimum. To describe this ROS-dependent additional time, we assume that Δ*t* (*R*, *T*) is described by the time it takes a cell to remove sufficient intracellular ROS and repair enough damages to divide. For example, we have experimentally measured that cells with low intracellular ROS concentrations are likely to duplicate whereas cells with high intracellular ROS concentrations are unlikely to duplicate (Fig. 3b). We can therefore assume that a cell has some amount of doubling time delaying tasks to perform before it can duplicate and that this amount of time is proportional to the abundance of ROS. We further assume that these tasks can be performed in a time that is proportional to how fast a cell can express genes, for example to synthesize ROS-reducing enzymes and replacing all damaged components with a freshly synthesized component. Thus, we assume that Δ*t* (*R*, *T*) *∝* [ROS]*/r_g_*(*T*) (a cell synthesizes proteins at a rate *r_g_*(*T*) (a.u. / time) and an amount of [ROS] (a.u.) tasks (repair/protein synthesis) must be completed). Moreover, we have experimentally measured the intracellular ROS concentrations in cells, which is approximately normally distributed on a log-scale (see Supplementary Fig. S9). Finally, our measurements suggest that there is some threshold concentration of ROS above which duplications become very unlikely (Fig. 3b). We therefore assume that we can describe the distribution of intracellular ROS concentrations according to ln[ROS] *~ N*(*R*, 1), where *R* is some parameter describing environmental factors (see Fig. 2a,c). Thus, Δ*t* (*R*, *T*) is stochastic and described by,

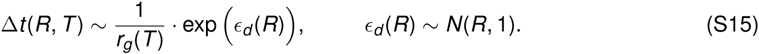

The time duration that increases the doubling time above its minimum thus varies between cells because of some ROS-dependent parameter that is log-normally distributed.

### Stochastic model for the doubling time dictated by temperature and ROS

Using the ROS-independent minimum doubling time (equation S14) and the ROS-dependent duration of the doubling time (equation S15), we obtain te doubling time as function of temperature *T* and environmental parameter *R*,

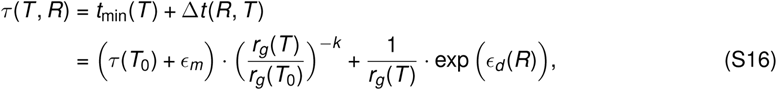

where *τ* (*T*_0_) and *r_g_*(*T*_0_) are the experimentally observed minimum doubling time and protein synthess rate at a temperature *T*_0_ respectively, *k* is the exponent that describes the scaling between the minimum doubling time and the protein synthesis rate, and (from equation S10),

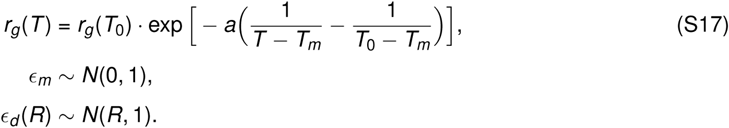

with *a* describing the protein synthesis rate as function of temperature, and the temperature *T_m_* below which all protein synthesis stops. With normally distributed *ɛ_d_* (for fixed *R*), we find that cells have longer doubling times but occur less frequently when *ɛ_d_* increases above *R*. Thus, the doubling time is a tradeoff between the speed of duplications and the likelihood of such a cell occurring. In practice, the values of *ɛ_d_* are bounded because there exists a longest possible doubling time *t*_max_ due to an experimentally observed threshold value of *ɛ_d_* (threshold of ROS concentrations) beyond which cells cannot duplicate and will die. The parameter *R* is the only free variable in the model, and all other parameters will be constrained next.

### Model parameters

We choose the known temperature *T*_0_ to be 5.0 °C, where we have experimentally measured *r_g_*(*T*_0_) *≈* 11.23 a.u. / day (Supplementary Fig. S30) and the experimentally measured minimum doubling time *τ* (*T*_0_) *≈* 2.5 days (with added GSH: *~*25 hours DNA duplication, *~*22 hours DNA segregation, *~*4 hours in *G*2, and a median G1 duration of *~*9 hours (see Supplementary Fig. S22). We used the median G1 duration due to the shape of the distribution to more accurately estimate the minimum time spent in G1 with added GSH). We fitted the power-law type scaling to the minimum doubling time as function of the protein synthesis rate to obtain the exponent *k ≈* 0.77 (Fig. 6b, Pearson correlation-coefficient *β* = 0.9979). Only single-cell data was used to determine the actual minimum doubling times (assumed to be equal to the average single-cell doubling time at 30 °C, the average single-cell doubling time with added GSH at 5.0 °C, and the minimum doubling times at 1.0 °C that exclude the duration of G1). As a lower temperature limit *T_m_* for protein synthesis we took *T_m_* = *−*10 °C, motivated by the fact that the yeast growth medium freezes at *−*2 °C (and thus limits protein synthesis by for example restricting access to nutrients, leading to deprivation of oxygen and limiting movement of (say) ribosomes) and because important enzymes such as Catalase stop functioning at *−*6 °C [18] and other essential enzymes inactivate at 0 °C [19,20]. It is therefore safe to assume that the temperature below which yeast’s protein synthesis stops lies above *−*10 °C. Having fixed *T_m_*, we fitted the temperature dependence of the protein synthesis rate using equation S11 from which we find that *a ≈* 120.72 (Fig. S30, Pearson correlation-coefficient *β* = 0.9954). Substituting the values of all known parameters yields the model that was used for all simulations,

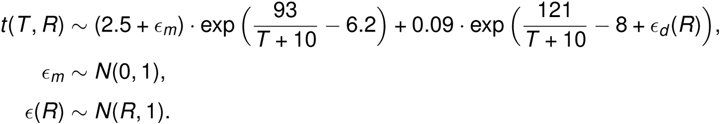

This model describes the single-cell doubling time as function of temperature *T* and parameter *R* being the only free parameter of the model describing the intracellular ROS concentrations.

In practice, we constrained *R* by dictating the average doubling time of the population (knowing the experimental average value of *τ* (*T*, *R*) at a given *T* constrains *R*). We constrained the parameter *R* as follows. We determined the average single-cell doubling time with and without added GSH at 5.0 °C (Supplementary Fig. S11). We then chose *R* such that the predicted doubling time of each respective population was equal to the measured experimental doubling time. At 5.0 °C, this yielded *R* = 3.2 for populations without added GSH, and *R* = 0.35 for populations with added GSH (yielding Fig. 6c). We then simulated *R* for different temperatures by taking *R* = 3.2 (without GSH) and decreasing *R* by 1.5% for every 0.1 °C increase in temperature (yielding Fig. 6d). Finally, we note that cells are extremely unlikely to duplicate for sufficiently high intracellular ROS concentrations (Fig. 3b). We therefore assume that the cells beyond the 98.5-th percentile of intracellular ROS concentrations in the model cannot duplicate, yielding the slowest possible doubling times of the model.

## Details of Methods

### Yeast strains

The ”wild-type”, haploid yeast strain that we used is from Euroscarf with the official strain name ”20000A”. It is isogenic to another laboratory-standard haploid yeast ”W303a”, and has the following genotype: *MATa; his3-11 15; leu2-3 112; ura3-1; trp1*Δ*2; ade2-1; can1-100*. For engineering the strains that express a fluorescent protein, we first built a strain ”AB01” having a functional *ADE2* gene and thus synthesizes adenine. This strain therefore no longer accumulates red pigments that would accumulate without the functional *ADE2* [21] and would interfere with detecting fluorescent proteins. To build this strain, we used PCR to insert the functional *ADE2* gene into the locus of the defective *ade2* gene in the wild-type strain by homologous recombination. To obtain the strain that we used for translation rate measurements, we started from AB01 and constructed a ”mCherry-inducible” strain. Specifically, the mCherry expression is controlled by the *GAL10* promotor, and was on a yeast-integration plasmid that constitutively expressed *URA3* (from *C. albicans*). This construct was integrated into the non-functional *his3* locus of AB01 by homologous recombination. To express mCherry, the ”mCherry-inducible” strain was grown on the trisaccharide raffinose (2% w/v) and expresses mCherry upon addition of galactose (2% w/v) to the growth medium.

To detect whether cells secrete glutathione to their extracellular environment, we used a mutant (*gsh1*Δ strain) that we had previously constructed [5]. This mutant lacks the *GSH1* gene for glutathione biosynthesis, and therefore must import glutathione from the extracellular environment for survival and growth. In short, we amplified the HygB selection marker (hygromycin B phosphotransferase) by PCR using primers whose ends were homologous to the flanking regions of the *GSH1* gene. We then used this PCR product to knock out *GSH1* via homologous recombination. Mutants were selected on YPD selection plates containing hygromycin B and verified the knockout by PCR.

To follow the cell-cycle progression in live cells, we constructed a strain that had several cell-cycle markers tagged with a fluorescent protein. Specifically, we fused the *GFP* gene to *HTB2* and then fused mCherry to *WHI5*. To do so, we started from AB01 and replaced the stop codons of the genes of interest with a cassette coming from yeast integration plasmids (pkt127, [22]). In short, for *HTB2*, the cassette contained a linker sequence followed by the *GFP* gene without a start codon and the KanMX selection marker. We amplified this construct by PCR using primers whose ends were homologous to the flanking regions of the stop codon of the *HTB2* gene. We then used this PCR product to replace the stop codon of *HTB2* with the linker-*GFP* and KanMX construct by homologous recombination. This created a *HTB2-linker-GFP* fusion gene. Mutants were selected on YPD plates containing Geneticin (G418) and verified by PCR and microscope. Next, we used the *HTB2-GFP* mutant to analogously construct the *WHI5-linker-mCherry* fusion gene, except that the integration cassette contained the NatMX selection marker and mutants were selected using YPD plates that contained nourseothricin. The resulting ”cell-cycle marker” strain contained two fluorescent fusion proteins, encoded by *HTB2-GFP* and *WHI5-mCherry*.

### Measuring genome-wide transcription rate with 4tU labelled RNA (see 1-4)

#### (1)#Sample preparation

We prepared large cultures of our wild-type yeast similarly to the description in ”growth experiments”. Specifically, we incubated single yeast colonies in Erlenmeyer flasks containing 100 mL of minimal medium at 30 °C for ~24 hours. We then took aliquots (~25 mL) of these cultures containing ~100 million cells, spun them down using a centrifuge and dissolved the yeast pellet with 250 mL fresh SC medium in 500 mL Erlenmeyer flasks. We then transferred the cultures to a shaker set to 150 rpm that we kept at (say) 5 °C. To ensure that the yeasts had sufficient nutrients, we refreshed the growth media several times during the two weeks of incubation at 5 °C (on days 7, 10 and 13). Specifically, we spun down the cultures using a centrifuge (Eppendorf, 5810R) that we pre-cooled to 5 °C for 5 min at 4, 000 rpm, discarded the supernatant, and re-suspended the pellet into fresh, pre-cooled minimal medium. We ensured that the cultures never ran out of nutrients by checking that the discarded supernatant contained sufficient nutrients to allow for growth of yeast – the supernatant contained enough nutrients to always grow the wild-type yeast to a density of *>* 1, 000, 000 cells / mL at 30 °C.

#### (2)#Sample collection

Previous work suggests to prepare a 2 M 4-thiouracil (4tU, Sigma Aldrich, cat. no. 440736-1G) solution in DMSO that is then added to the growth medium at a final 5 mM concentration [15]. We found that the 4tU solution precipitates and does not dissolve when added to the growth media at *≤* 5 °C. To circumvent this problem we added the 4tU solution to minimal medium at 37 ^°^C to a final 7.5 mM concentration (1.5x). We then cooled this 4tU medium overnight to 5 °C without the 4tU precipitating. After two weeks, we first took aliquots without 4tU as a control. We then added the pre-cooled, fresh medium containing 7.5 mM 4tU to the cultures resulting in the cultures having a final concentration of 5 mM 4tU. Samples were subsequently collected after 1.5, 3, 6, 12, 24, 48 and 96 hours of incubation with 4tU (or 0, 6, 12, 24, 48 and 96 hours at 1 °C or 0, 5, 10 and 20 min at 30 °C). During sample collection at 1 °C or 5 °C, we transferred aliquots of the cultures into 50 mL falcon tubes and spun them down with a pre-cooled centrifuge for 5 min at 4, 000 rpm. We discarded the supernatant and re-suspended the pellet in 1 mL RNAlater (Cat. No. AM7021, Thermo Fischer Scientific). To wash the samples, we again spun down the samples for 2 min at 4, 000 rpm, removed the supernatant and re-suspended the pellet in 1 mL RNAlater. During sample collection at 30 °C, we collected aliquots of the cultures into 50 mL falcon tubes and directly flowed the aliquots through 0.2 μm bottle-top filters with a vacuum. The cells are too small to pass through the filter and remain on the membrane. We then washed the cells with 50 mL ice-cold PBS and then removed the vacuum. The pellet was re-suspended with ~5 mL RNAlater, and cells were collected in 50 mL falcon tubes. Finally, right before storage, we took a 20 μL aliquot of the collected cells in RNAlater to determine the cell density with a flow cytometer (to know the total number of collected cells). All samples were stored at 5 °C for a couple of hours to ensure that RNAlater would penetrate the cells before storing the samples at *−*20 °C for further processing. As a spike-in of 4tU labelled RNA we used cells from *Schizosaccharomyces pombe* (YFS110) analogously to previous work [15]. Specifically, we grew *S. pombe* cells in 300 mL YPD on a shaker set to 150 rpm at 30 °C to a density of ~5, 000, 000 cells / mL. We then incubated the cells with 5 mM 4tU for 15 min to label the RNA with 4tU, collected the cells as described and stored the cells in 5 mL RNA later at *−*20 °C.

#### (3)#Sample processing

Before continuing with the RNA extraction we added a fixed amount of *S. pombe* cells to each sample (in a *~*1 : 6 ratio). We then spun down our samples in a pre-cooled centrifuge, removed the RNAlater, and proceeded with RNA extraction as described in the paragraph ”preparing cells and RNA extraction”. The RNA was extracted within three days of the last samples were collected. We simultaneously extracted the RNA of all time-points of a single sample to minimize the effect of processing on the observed transcription rate. After RNA extraction, we proceeded with biotinylation and purification of the 4tU labelled RNA. Here we followed the protocol described in previous work with minor modifications [15]. Briefly, the RNA was biotinylated using a biotinylation buffer (a solution containing 100 mM Tris, 10 mM EDTA, pH 7.6 (Cat. No. BP2475, Thermo Fischer Scientific)) and biotin-HPDP (dissolved in DMSO) at a final volume of 1 mL. After incubation, the excess biotin-HPDP was removed by mixing in and washing with 1 mL of chloroform. The RNA was then precipitated with 100 μL of 5 M NaCl and 1 mL isopropanol and spun down using a pre-cooled centrifuge for 45 min at 18, 000 x g (used throughout). The RNA was washed with 1 mL of 75% ethanol and re-dissolved in 100 μL DEPC treated water (Invitrogen, 750023). After biotinylation, we directly continued with purification of the 4tU labelled RNA. To do so, we incubated the biotinylated RNA with 100 μL streptavidin-coated magnetic beads (cat. No. 120-001-017, Miltenyi Biotec) for at least 90 min at room temperature. During that time, we placed micro-columns to a magnetic stand (microMACS separator (cat. No. 130-042-602, Miltenyi Biotec)) and pre-washed the columns with 1 mL washing buffer (a solution containing 100 mM Tris, 10 mM EDTA and 1 M NaCl, pH 8.0 (Cat. No. BP2479, Thermo Fischer Scientific), and 0.1% Tween-20 (cat. No. BP337-100, Thermo Fischer Scientific)). After incubation we applied the RNA to the columns. We re-applied the flow-through to the columns as the columns do not capture all magnetic beads during the first flow-through. After washing the columns 5 times with 1 mL washing buffer, we eluted the 4tU labelled RNA twice using 200 μL of 0.1 M DTT (Cat. No. 43816, Sigma Aldrich). To precipitate the RNA, we then added 40 μL of 3 M NaOAc (Cat. No. S7899, Sigma Aldrich) and 1.3 mL ice-cold 100% ethanol (Cat. No. 1.00983.1011, Merck KGaA) to the eluted RNA. Finally, we added 2 μL of 20 mg/mL RNA-grade glycogen (Cat. No. R0551, Thermo Fischer Scientific) to aid precipitation, and left the RNA to precipitate overnight (at least 15 hours) at *−*20 °C. Finally, the 4tU labelled RNA was resuspended in 15 μL of DEPC-treated, RNAse-free water. We found that the RNA extracted from *~*25, 000, 000 cells of *S. pombe* that were incubated with 5 mM 4tU for 15 min (our spike-in) was generally enough to yield *~*15 μL of *>* 40 ng/μL 4tU labelled RNA. This 4tU labelled RNA has poor RIN numbers and rRNA ratio’s (due to incomplete synthesis of the purified ribosomal RNA) but is enough and of sufficient quality for RNA sequencing.

#### (4)#Data normalization

We performed a quality check of each sample with FastQC and processed all sequencing data with the Salmon tool to quantify gene expression levels [23]. Specifically, we obtained reference transcripts for *S. cerevisiae* from Ensembl (assembly R64-1-1) and built a Salmon index containing the transcripts (cDNA and ncRNA) from *S. cerevisiae* with the whole genome as a decoy. We then quantified the transcripts in each sample using this index. Finally, we converted the transcript levels for *S. cerevisiae* to gene expression levels (Transcripts Per Million, TPM) and merged all samples using the package tximport from Bioconductor [24]. We then performed between-sample normalization by rescaling the TPM values for each gene with the sum of the TPM values of all *S. pombe* genes. The same amount of cells from the same culture *S. pombe* were added to each sample as a spike-in, such that the total transcript levels of *S. pombe* should be identical across samples (the spike-in thereby eliminates experimental variations between samples and ensures that the time-lapse represents the relative number of 4tU-labelled transcripts). We also rescaled the gene expression levels in the time-lapses with the number of estimated alive cells in the population for each time point (to ensure that measuring higher transcript levels is not due to having more cells in the population as a result of population growth). These cell numbers were corrected for the percentage of dead cells in the population as measured by PI staining (typically between ~2% and ~20%). These rescaling’s gave us the relative number of 4tU transcripts per cell over time. Finally, we converted these TPM values to ”# of RNA / cell” with a calibration curve that we constructed using the stead-state transcript levels that we had also sequenced. Specifically, during the 4tU time-lapses, we took aliquots of the populations of cells that were at log-phase growth (steady state gene expression) right before the 4tU labelling. We then performed single-molecule RNA FISH on these aliquots for several endogenous yeast genes (*RPS3, RPL3, RPB1, RPB3*). We then used the measured (average) number of mRNA per cell and the measured TPM value of the steady-state RNA-seq samples to create a calibration curve that converts steady-state transcript levels (from total RNA-seq) to integer number of RNA per cell (from RNA FISH, see Supplementary Fig. S29). We then used this calibration curve to convert our 4tU time-lapses to ”# of RNA / cell”. These transcript levels were used for further processing and analysis. (Also see Supplementary figs. S27–S29 for further details).

