## Supplementary Figs and Model for "Fundamental limits to progression of cellular life in frigid environments"

---

**This document contains:**

**-Supplementary Figures 1 - 33**

**-Supplementary Text (starts on Pg. 55)**

#### Supplementary Figures

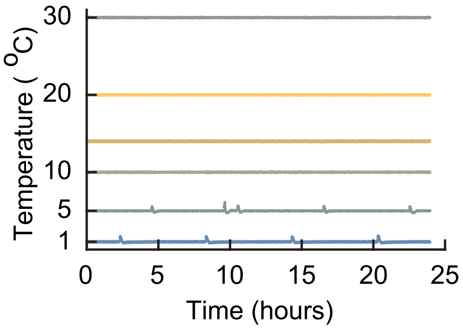


**Figure S1: Temperature remains stable during all our growth experiments (Related to Figure 1a).** All growth experiments were performed with liquid cultures of cells incubated in compressor-cooled, high-precision thermostatic incubators (Memmert ICPs). The ICPs had a precise temperature-recording device whose temperature readings we additionally verified using a separate thermocouple device and aligned with the temperatures of several other incubators. The temperature was monitored over the entire course of our experiments. As examples, shown here are the temperatures as recorded by the incubator's temperature sensor for 24 hours during six different growth experiments. Starting from the top, the curves show the temperature of the incubator set at 30.0 °C, 20.0 °C, 14.0 °C, 10.0 °C, 5.0 °C and 1.0 °C. Throughout our experiments, the incubators had a typical standard deviation of 0.033 °C when the incubation temperature was above 10.0 °C. The standard deviation was 0.09 °C when the incubation temperature was below 10.0 °C. This slightly larger standard deviation was due the incubator undergoing short thaw-cycles when the incubation temperature was 5.0 °C to prevent freezing of its components (deviation measured over several days).

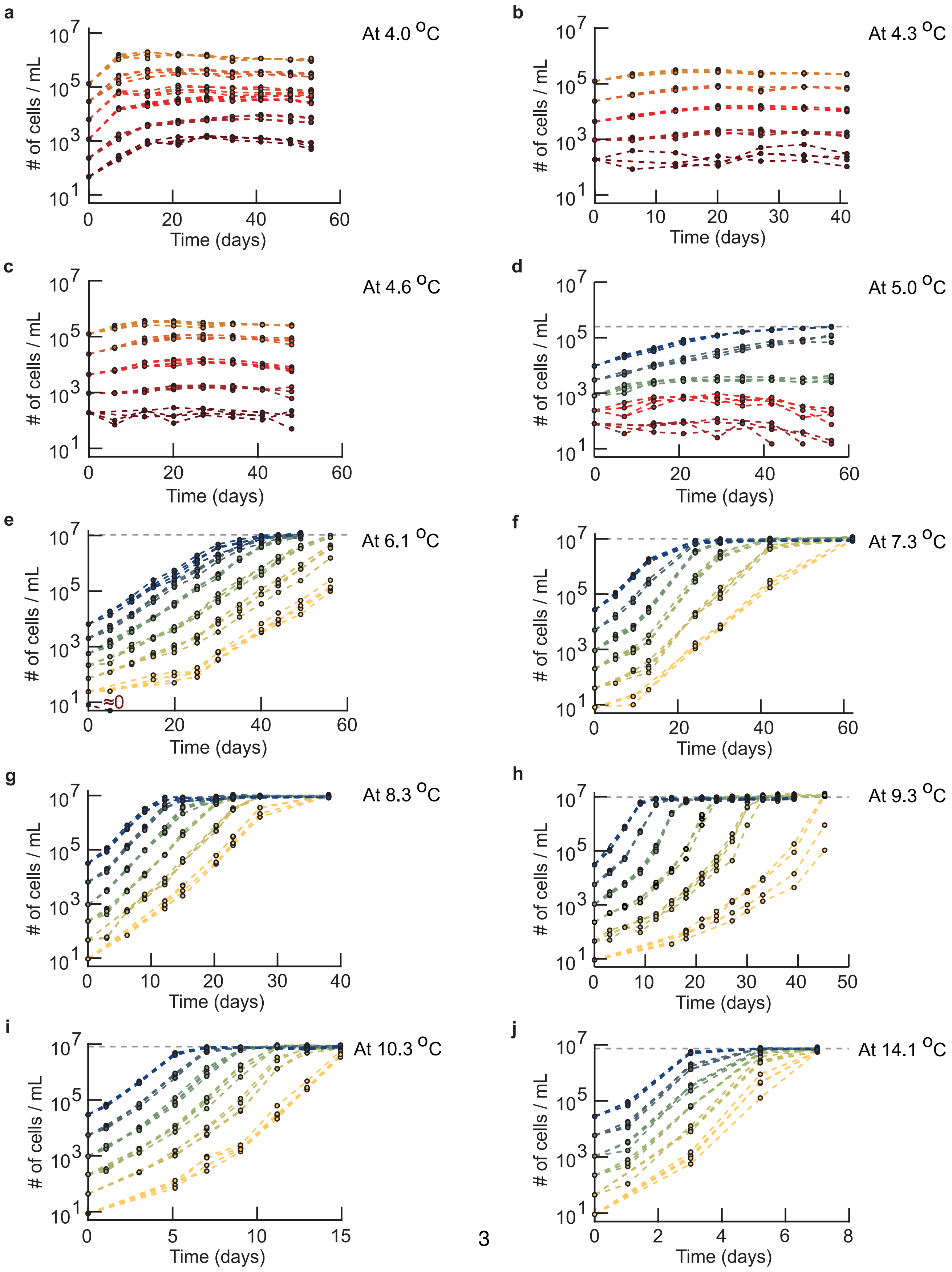
a
At 4.0 °C
b
At 4.3 °C
c
At 4.6 °C
d
At 5.0 °C
e
At 6.1 °C
≈0
f
At 7.3 °C
g
At 8.3 °C
h
At 9.3 °C
i
At 10.3 °C
j
At 14.1 °C
### of cells / mL
Time (days)

**Figure S2: Growth curves for widely varying initial population densities and temperatures (Related to Figure 1a-b).** Population density (number of cells / mL) measured over time with a flow cytometer. Shown are populations of wild-type yeast with differing initial densities incubated at 4.0 °C **(a)**, 4.3 °C **(b)**, 4.7 °C **(c)**, 5.0 °C **(d)**, 6.1 °C **(e)**, 7.3 °C **(f)**, 8.3 °C **(g)**, 9.3 °C **(h)**, 10.3 °C **(i)** and 14.1 °C **(j)**. Different colors represent different initial population-densities. The grey line shows the carrying capacity that we estimated from the final densities of the populations. To construct the phase diagram (Figure 1b), we used the growth-kinetics data as shown here to determine whether a population with a given initial population-density should be characterized as growing ("growth" phase) or non-growing ("no growth" phase). An initial population-density was characterized as being in the growth phase (Fig. 1b – blue region) if all replicate populations that started with that density exponentially grow over time and reach the carrying capacity for that temperature. An initial population-density was characterized as being in the no-growth phase (Fig. 1b – red region) if all replicate populations that started with that density did not grow during several weeks of incubation, except for some initial transient growth that results from the cells having been transferred from 30.0 °C. This transient growth typically lasted a few days. For example, every population in (c) belongs to the no-growth phase because no population grew during the ~6 weeks of incubation at 4.6 °C as can be seen by every population density barely increasing over time. As another example, after some lag-time, every population in (g) grew exponentially and identically over time to reach the carrying capacity at 8.0 °C. To draw the phase boundary that separates the growth phase from the no-growth phase in the phase diagram (Fig. 1b), we connected the maximum initial density for which a population does not grow for each temperature. This curve almost overlaps with the minimum initial density that leads to population growth for each temperature because we sampled the initial densities close to each other for each temperature. Similarly, we drew the boundary curve that separates the growth phase from the region of the phase diagram where nutrients are lacking by connecting the data points that represent, for each temperature, the measured carrying capacity (the grey lines in d-j). Finally, we determined the temperature below which no population growth is possible — and thus only the no-growth phase exists below this temperature -— by identifying the highest temperature (i.e., 4.6 °C (c)) where populations with different starting densities always reach differing final densities (as opposed to the common, carrying capacity). In fact, these populations do not grow at all beyond the initial, transient growths (a-c). All panels show $n$ = 4 replicate populations for each initial population-density (color).

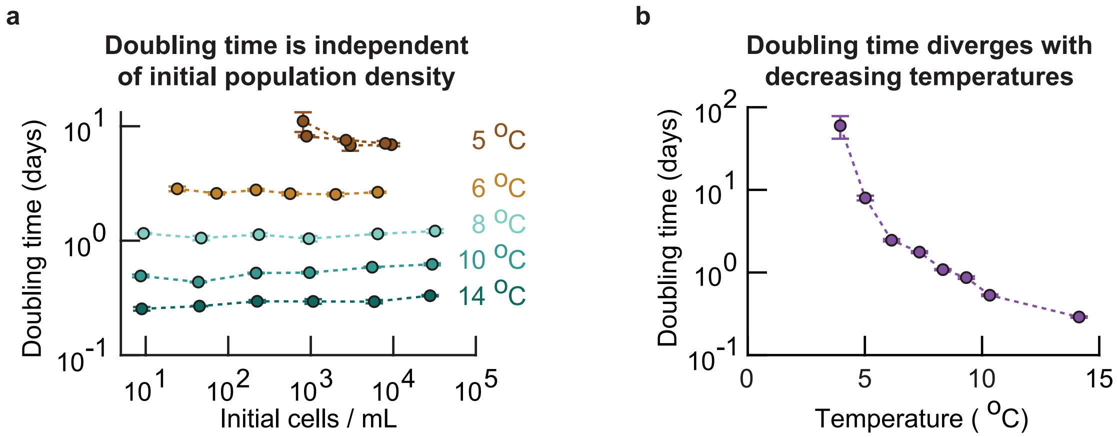


**Figure S3: Population's doubling time is independent of its initial density at near-freezing temperatures (Related to Figure 1b).** Summarizing Supplementary Fig. S2. From the growth curves of populations at various temperatures, we determined each population's doubling time. To obtain the doubling time of the populations that grew (i.e., populations in the "growth" phase of the phase diagram, Fig. 1b), we excluded the initial transient growth that typically lasted from a few days to one week. For non-growing populations (e.g., populations in the "no-growth" phase at 4.0 °C) we only took the growth rates that were positive and excluded the initial transient growth (i.e., transient growth in the first week of incubation). **(a)** Population doubling time as a function of initial density and temperature for growing populations. The doubling time does not depend on initial density. Error bars show the mean with s.e.m., and each data point represents at least $n = 4$ replicate populations. **(b)** Since the doubling time does not depend on initial density, we pooled all observed doubling times for each temperature. The population doubling time diverges as the temperature decreases. Error bars represent the mean with s.e.m., having at least $n = 16$ biological replicates per data point.

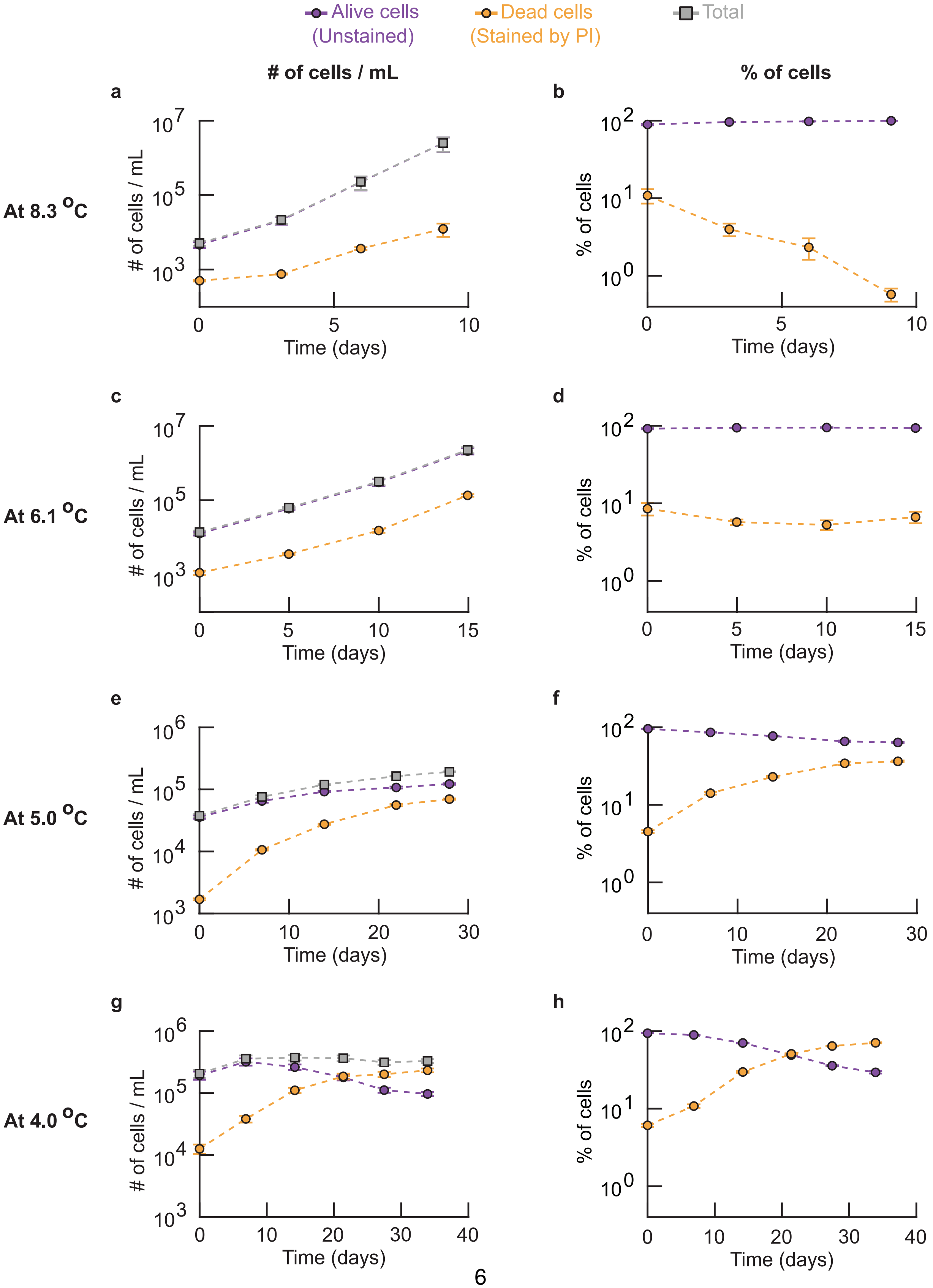
Alive cells
(Unstained)
Dead cells
(Stained by PI)
Total
### of cells / mL
% of cells
a
b
c
d
e
f
g
h
At 8.3 °C
At 6.1 °C
At 5.0 °C
At 4.0 °C
### of cells / mL
% of cells
Time (days)

**Figure S4: Population extinction occurs below 5 °C because as the temperature decreases, the average doubling time of a cell increases and eventually, just below ∼5 °C, becomes larger than the average cell's lifespan (Related to Fig. 1b).** Left column **(a, c, e, g)**: number of dead cells (yellow), number of alive cells (purple) and total number of cells (grey) in a population. Right column **(b, d, f, h)**: percentages of dead (yellow) and alive (purple) cells in a population. For each population, we took an aliquot of the liquid culture at various times and incubated it with 1 μg / mL of propidium iodide (PI) for 20 minutes at room temperature. We then flowed this aliquot through a flow cytometer to measure the total number of cells and the number of cells that were stained (red) and unstained (yellow) by PI. PI does not stain cells with an intact membrane [1]. In contrast, PI enters cells with a damaged membrane and stains their DNA. Thus, propidium iodide stains a cell if and only if it's membrane is permeable. Hence, alive cells are commonly assumed to be impermeable to PI. Shown here are populations of wild-type yeast incubated at 8.3 °C **(a-b)**, 6.1 °C **(c-d)**, 5.0 °C **(e-f)** and 4.0 °C **(g-h)**. The curves show the total population-density and the density of stained and unstained cells over time **(a, c, e** and **f)**, together with the percentage of cells that were stained and unstained in the population **(b, d, f** and **h)**. All populations were already in a steady-state at the respective temperature before the PI staining on day 0 in the graphs (i.e., the populations had already spent ∼6 days at 8.3 °C (a-b), ∼10 days at 6.1 °C (c-d), and ∼14 days at 5.0 °C (e-f) or 4.0 °C (g-h)). Error bars show the mean with s.e.m., with $n$ = 4 replicate populations per data point. **(a-b)** Alive and dead cells over time at 8.3 °C. All populations grow exponentially over time. Both the density of alive and dead cells increase exponentially over time, with the density of alive cells increasing faster than the density of dead cells. Specifically, the percentage of dead cells in the population decreases exponentially over time. Thus, at 8.3 °C, alive cells overtake the population as cell duplications occur more frequently than cell deaths (purple curve in (b) stays at ∼100%). **(c-d)** Alive and dead cells over time at 6.1 °C. Similar to the results at 8.3 °C. The main difference here is that the density of alive cells and the density of dead cells increases exponentially with the same rate (the two curves in (c) are nearly parallel), whereas the density of alive cells increased faster than the density of dead cells at 8.3 °C (see (a)). Indeed, the percentage of dead cells remains constant over time at 6.1 °C (see d). Like in 8.3 °C, nearly 100% of the population consists of alive cells at 6.1 °C. **(e-f)** Alive and dead cells over time at 5.0 °C. The density of alive cells increases over time. However, the percentage of dead cells in the population increases while the percentage of alive cells decreases over time. Still, the populations consists of more alive cells than dead cells after ∼6 weeks of incubation at 5.0 °C (see (f)). **(g-h)** Alive and dead cells over time at 4.0 °C. The population does not grow and the density of alive cells decreases over time (g) (i.e., less than one cells divides per cell that dies). The percentage of dead cells increases exponentially over time while the percentage of alive cells decreases, leading to the population eventually consisting mostly of dead cells (the population becomes extinct as this trend continues over time) (see (h)). ***(caption continues on the next page)***

**Figure S4** ***(caption continued from the previous page)***: Together, (a-h) show that as the temperature decreases, the average doubling time of a cell increases and eventually becomes larger than the average lifespan of a cell. Having a doubling time that is larger than the average lifespan means that the average cell does not have time to duplicate before it dies. This leads to a population extinction. In other words, populations grow more slowly as temperature decreases and eventually, at around 5.0 °C, populations cannot exponentially grow because the average doubling time nearly matches the average lifespan of cells. Finally, at 4.0 °C, the doubling time exceeds the lifespan of cells and thus populations cannot grow.

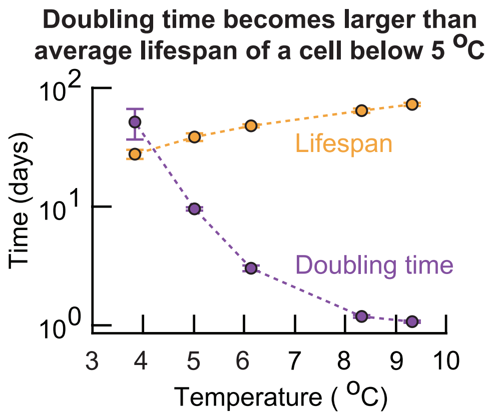


**Figure S5: Doubling time of a cell increases and eventually becomes larger than the averace cell's lifespan as the temperature decreases below 5 °C (Related to Figure 1b).** Summarizing Supplementary Fig. S4. We used our measurements of the number of alive and dead cells in populations at each temperature to extract the average doubling time and the average lifespan of cells. Specifically, we fitted a simple growth model to the data (also see Supplementary Theory). Shown is the average cell's doubling time (purple) and lifespan (orange) at various temperatures. The average doubling time exceeds the average lifespan when the temperature is below 5.0 °C. Consequently, below 5.0 °C, a cell is expected to die before it has a chance to duplicate and hence the population approaches extinction. Error bars represent the mean with s.e.m., having at least $n$ = 16 biological replicates per data point.

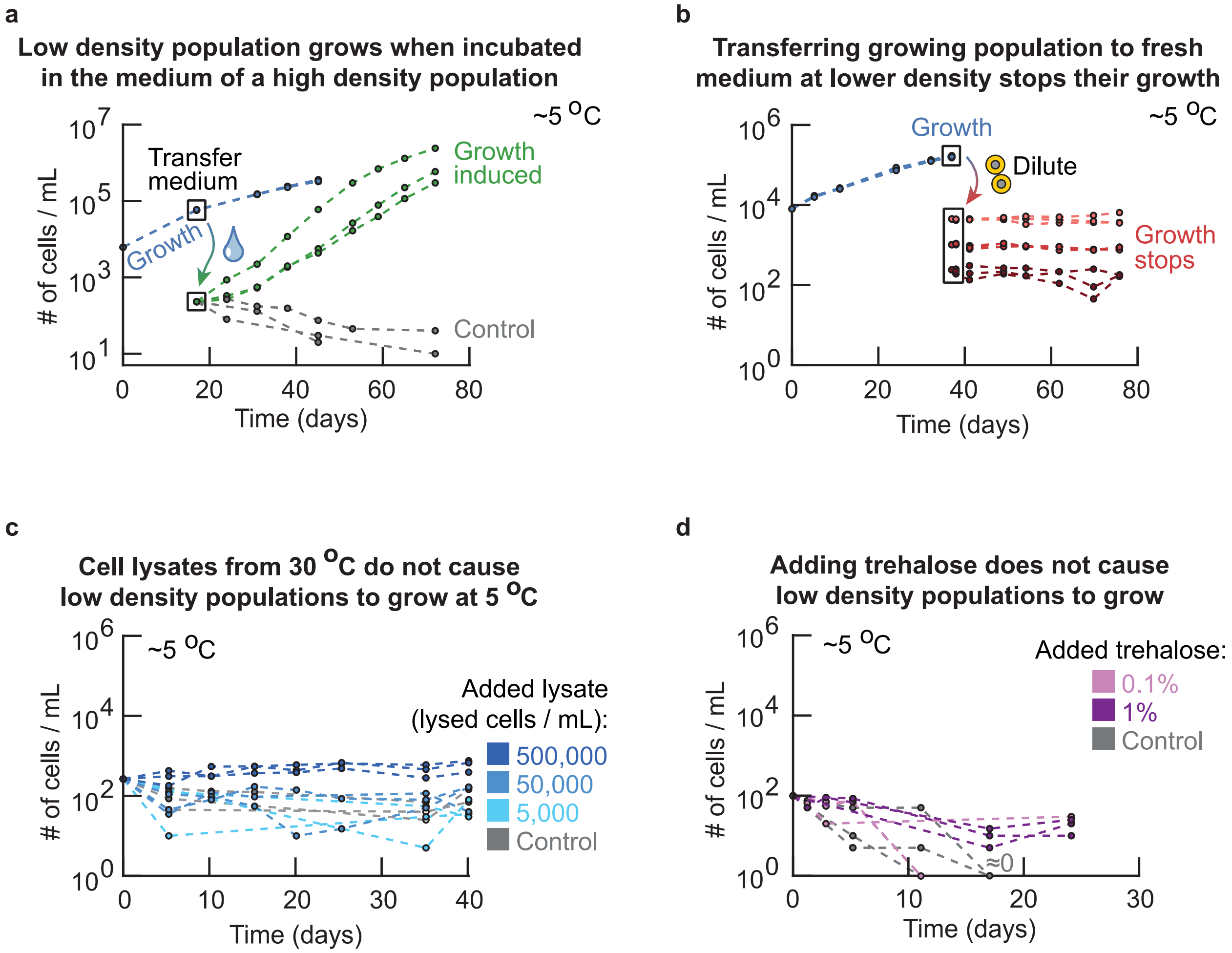


**Figure S6: Secreted factor in extracellular medium determines whether a population grows or not at freezing temperatures (Related to Figure 2a).** Testing why high-density populations grow whereas low-density populations do not grow at the same near-freezing temperature (5.0 °C). **(a)** To test whether the cell's ability to duplicate at 5.0 °Cis dictated by secreted factor(s) in the extracellular medium, we took the growth medium of a high-density (growing) population (blue curves: initially $\sim$6, 250 cells / mL) after several weeks of growth at 5.0 °C. Boxed data points on the blue curves show when the growth medium was taken. We flowed the high-density culture through a membrane filter with 0.2 μm pores so that growth medium taken from this culture was free of cells, which we confirmed with a flow cytometer (i.e., no cells were detected). Next, into this filtered medium, we incubated a fresh, low-density population of cells that were growing in 30 °C (green curves: initially $\sim$250 cells / mL). As a control, we also incubated the same, low-density of cells in a fresh growth medium (grey curves: initially $\sim$250 cells / mL). We incubated the two, low-density cultures at 5.0 °C and measured their cell numbers over time. The populations of fresh cells grew in the filtered growth medium of the high-density population at 5.0 °C (green curves). But the low-density (control) populations in the fresh medium did not grow (grey curves). ***(caption continues on the next page)***

**Figure S6** ***(caption continued from the previous page)***: This experiment shows that cells change their growth medium (e.g., secreted factor(s)) such that they can grow at 5.0 °C and that a sufficiently high density of cells is required for the change to be sufficient for a population growth. **(b)** To test whether the ability to duplicate at 5.0 °C is determined solely by intracellular factor(s) (e.g., heritable trait), we took an aliquot of cells from a high-density (growing) population at 5.0 °C and diluted it by various amounts into fresh, pre-cooled media at 5.0 °C to test whether the resulting low-density population could grow. The high-density population of wild-type cells (blue curves, initially ∼10,000 cells / mL) grew at 5.0 °C. After ∼35 days of growth, we took an aliquot of cells from this culture and diluted it by ∼50x (light red curves, initially ∼10,000 cells / mL), ∼250x (red curves) or ∼1250x (dark red curves). Ordinarily, populations that start at these "low densities" do not grow at 5.0 °C (Fig. 1b). We incubated the diluted, low-density populations in fresh medium at 5.0 °C and measured their density over time (red curves). None of these populations grew at all during more than one month of incubation at 5.0 °C. This experiment shows that the ability to duplicate at 5 °C is not determined by any intracellular factor(s). **(c)** To test whether the extracellular factors that enable population growth come from dying cells that may be lysing, we incubated populations in media with cell lysates. We obtained the cell lysates from a population of wild-type cells growing in log-phase at 30.0 °C (∼5,000,000 cells / mL). We added glass beads to this liquid culture and placed it on a vortex for 20 min to mechanically lyse the cells. After this, we flowed the liquid media containing the lysed cells with the beads through a 0.45 μm pore filter to remove the glass beads and any intact cells that remained. We then added the filtered cell lysate to a fresh medium in amounts dictated by the cell densities before lysis (e.g., fresh medium having 500,000 lysed cells / mL contains 90% fresh medium and ∼10% cell lysate that had ∼5,000,000 cells / mL before the lysis). Finally, we incubated fresh cell populations in these growth media at ∼250 cells / mL and at 5.0 °C. The curves here show populations with ∼500,000 lysed cells / mL (dark blue curves), ∼50,000 lysed cells / mL (blue curves), ∼5,000 cells / mL (light blue curves) and, as control, fresh cells in a fresh medium without any cell lysate (grey curves). None of these populations grew at 5.0 °C. This experiment shows dying cells that may be lysing are not responsible for inducing population growths at 5.0 °C. **(d)** Trehalose is a common cryoprotectant that is abundant in yeasts at low temperatures and is thought to protect the cells against freezing [2]. Its precise mechanism is incompletely understood. We reasoned that yeasts may be secreting trahalose to help them grow at 5.0 °C. To test this idea, we incubated a low-density population at 5.0 °C (initially ∼100 cells / mL) with 1% (dark purple curves, percentage in weight per volume), 0.1% (light purple curves) or without trehalose (grey curves). None of the populations grew, indicating that trehalose is not one of the extracellular factor(s) that induce growth at 5.0 °C. All panels show at least $n = 3$ biological replicates per condition (color).

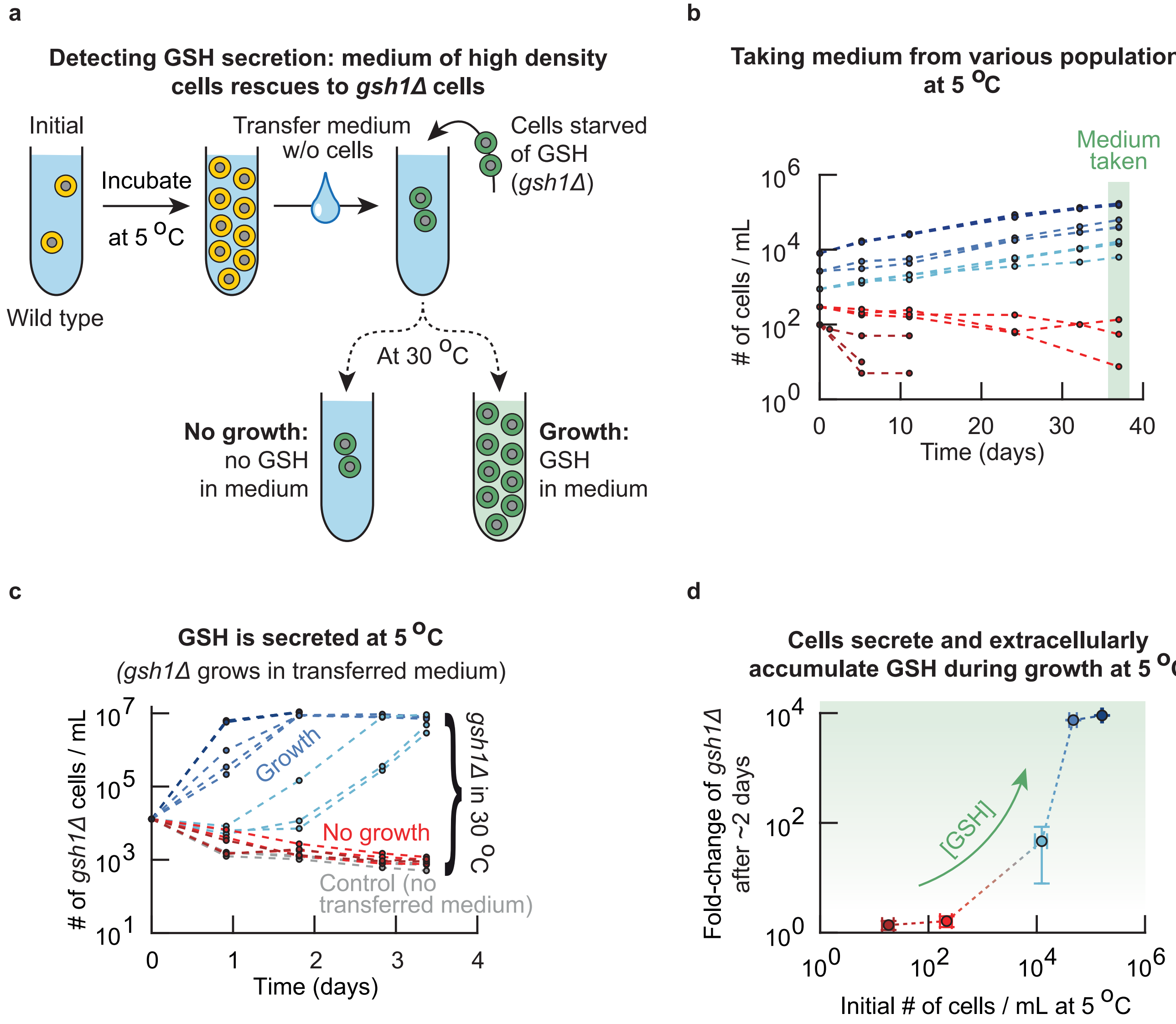


**Figure S7: A non-viable mutant that cannot synthesize glutathione (*gsh1*Δ) is rescued by medium from high-density populations, which confirms that cells in high-density populations secrete and extracellularly accumulate glutathione at 5 °C (Related to Figure 2a). (a)** Schematics of experiments in (b-d). A mutant that cannot synthesize glutathione (*gsh1*Δ-strain) cannot live without supplemented GSH even at 30 °C because GSH is essential for life (e.g., GSH is involved in iron metabolism [3]). The only way for the mutant to survive and grow at any temperature is by supplementing GSH into its growth medium [4, 5]. To test whether cells at 5.0 °C secrete glutathione during log-phase growth and stationary phases, we incubated wild-type yeast populations at 5.0 °C at different initial population-densities. After several weeks of growth, we isolated the growth medium from the wild-type populations. ***(caption continues on the next page)***

**Figure S7** ***(caption continued from the previous page)***: Specifically, we took aliquots of the cultures at 5.0 °C and removed the cells from the growth media by spinning down the aliquot and passing the supernatant through a 0.2 μm pore filter to remove all yeasts. We checked that no cells remained in the filtered media by flowing them through a flow cytometer (no events detected). In the filtered media, we incubated a population of *gsh1*Δ cells that we had starved of glutathione by incubating them overnight in fresh medium without GSH at 30.0 °C (These cells therefore did not grow overnight). If the filtered media did not contain any glutathione, then the starved *gsh1*Δ cells would not grow. If the filtered media did contain glutathione secreted by the wild-type cells, the *gsh1*Δ cells would be able to import the GSH and grow. **(b)** As a first part of the experiment described in (a), we incubated wild-type yeast populations at 5.0 °C and determined whether their growth media contained glutathione after ∼5 weeks of growth. Shown is the wild-type's population density over time for different starting densities (∼300 cells / mL (red curves), ∼900 cells / mL (light blue curves), ∼2,700 cells / mL (blue curves) and ∼8,000 cells / mL (dark blue curves)). Each color shows $n$ = 3 replicate populations. **(c)** To detect extracellular glutathione, we isolated and filtered the growth media from the wild-type populations in (b). Shown is the population density over time at 30.0 °C of glutathione-starved *gsh1*Δ cells that received the filtered media (initially ∼13,000 cells / mL). The colors of the curves here match the colors used for the wild-type populations in (b) (e.g., the red populations received the filtered media from the wild-type populations whose growth curves are shown in red in (b)). As a control, we also incubated *gsh1*Δ cells in fresh medium without any glutathione (grey curves). The *gsh1*Δ cells did not grow in the medium transferred from low-density (non-growing) populations at 5.0 °C. The *gsh1*Δ cells always grew to the carrying capacity in the medium from the high-density (growing) populations at 5.0 °C. Each color shows $n$ = 3 replicate populations. **(d)** Summary of the data from (b) and (c). The x-axis shows the population density of wild-type cells after ∼5 weeks of incubation at 5.0 °C. The y-axis shows, after two days of incubation at 30 °C, the number of *gsh1*Δ cells in transferred media relative to the number of *gsh1*Δ cells in a fresh medium without glutathione. Populations of *gsh1*Δ cells that grew more had more glutathione in their medium (glutathione is the growth limiting factor in the transferred media, see control in (c)). Error bars represent the mean with s.e.m., with $n$ = 3 biological replicates per data point. Together, (a-d) show that the amount of extracellular glutathione increases with population density for growing populations, and that barely any glutathione is secreted for non-growing populations. In summary, cells secrete and extracellularly accumulate glutathione during growth at near-freezing temperatures.

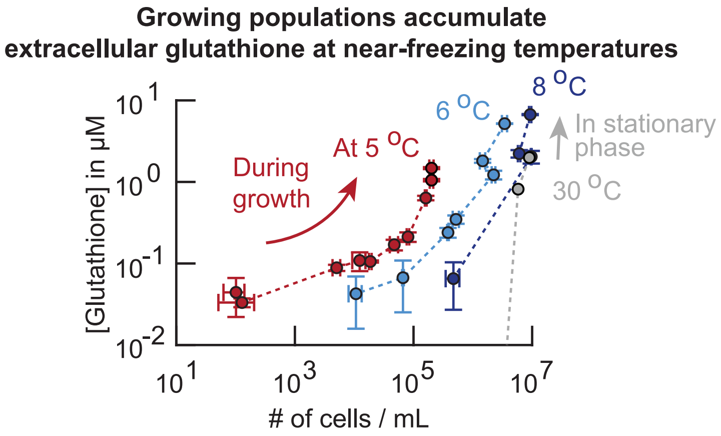


**Figure S8: Cells secrete and accumulate glutathione during growth at near-freezing temperatures (Related to Figure 2a).** Secretion of glutathione into the extracellular medium at various temperatures. We measured the total glutathione concentration in the growth media of wild-type yeast populations at two different times after ∼2 weeks of incubation at various temperatures (see Methods). Shown here is the total extracellular glutathione concentration as function of population density at ∼5.0 °C (red curve, one week between sampling), ∼6.1 °C (light blue curve), ∼8.3 °C (dark blue curve) and ∼30.0 °C (grey curve). The extracellular glutathione concentration increases during population growth at near-freezing temperatures (i.e., 5.0 °C - 8.3 °C), whereas glutathione only accumulates in stationary phase at 30.0 °C as we previously found [5] (data reused). Error bars show the mean with s.e.m., having $n$ = 3 biological replicates per data point.

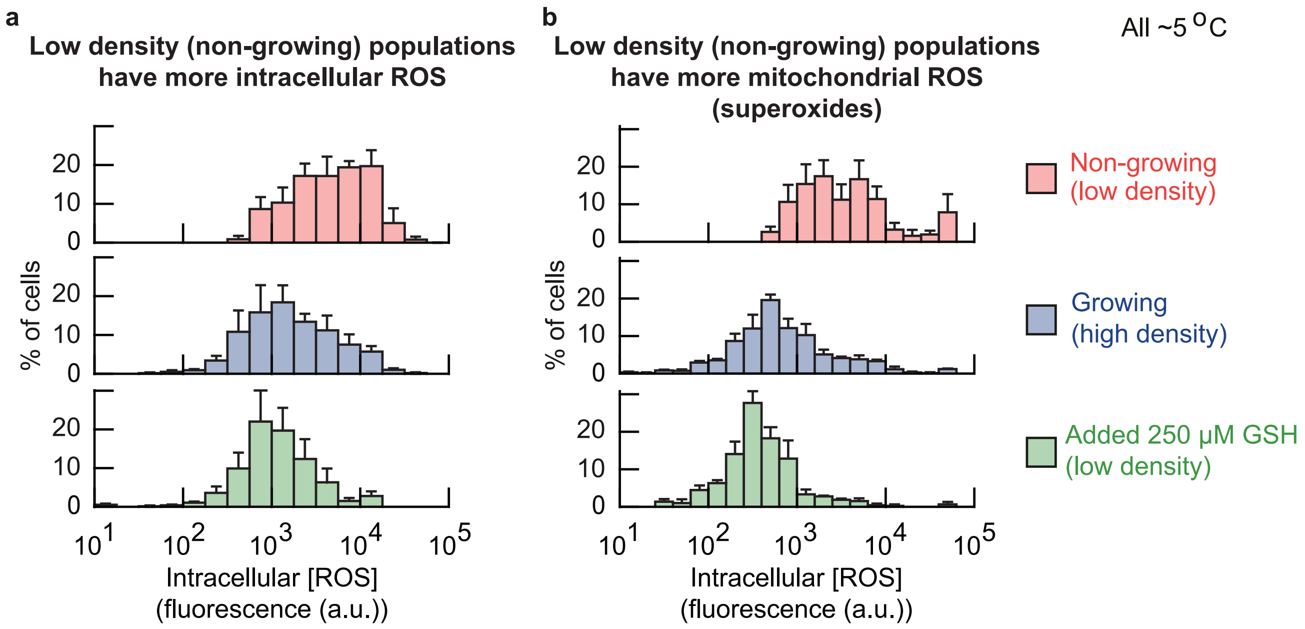


**Figure S9: Cells of low-density (non-growing) populations at 5 °C have more intracellular Reactive Oxygen Species (ROS), including superoxides in their mitochondria, which can be removed by supplementing extracellular reduced glutathione (GSH) (Related to Figure 2d). (a-b)** Intracellular ROS concentrations. We stained intracellular ROS with two different dyes. One dye (cellROX) measures the amount of general, cytoplasmic ROS while another dye (mitoSOX) measures the amount of superoxides in mitochondria. We grew populations of cells at 5.0 °C for two weeks and then stained the cells with the ROS dyes (see Methods). We then used fluorescence microscopy to determine the amounts of intracellular ROS in single cells, as represented by the average fluorescence of each cell (after subtracting background fluorescence). **(a)** Intracellular ROS concentration for general ROS (with cellROX). **(b)** Superoxide concentrations in mitochondria (with mitoSOX). Histograms in (a-b) show cells of low-density (non-growing) populations (red bars, initially ∼250 cells / mL), growing populations (blue bars, initially ∼6,250 cells / mL) and populations that had 250 μM added GSH (green bars, initially ∼250 cells / mL). These histograms show that the low-density (non-growing) populations have higher intracellular ROS concentrations than high-density (growing) populations. Adding extracellular GSH decreases the intracellular ROS concentrations. Error bars show the mean with s.e.m., having $n$ = 3 biological replicates per histogram.

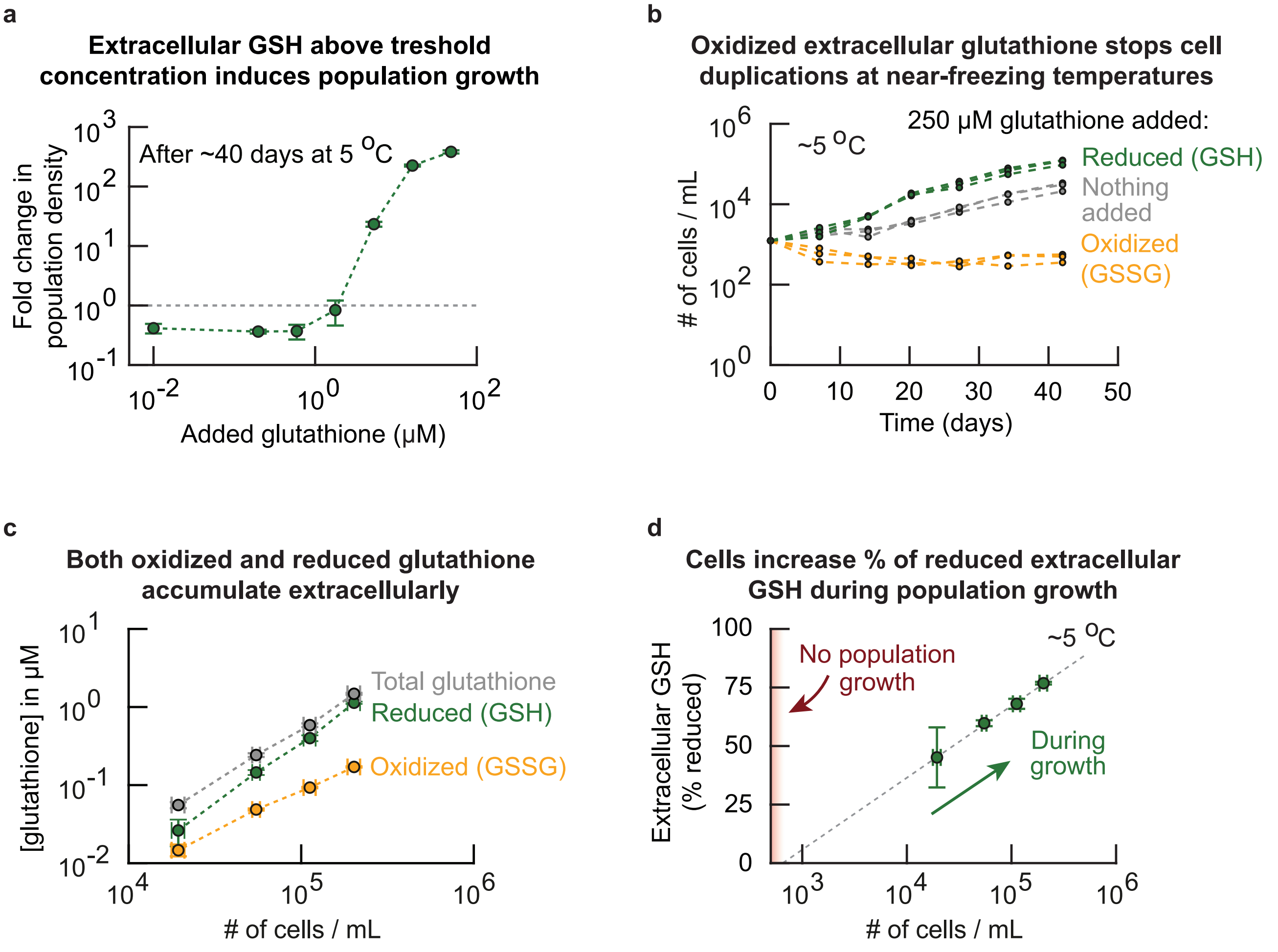


**Figure S10: Reduced form of glutathione, not the oxidized form of glutathione, enables and accelerates duplications at 5 °C above threshold concentration of ∼1 μM (Related to Figure 2d).** Unless stated otherwise, our experiments measured and added the reduced form of glutathione (GSH). But for the experiments in this figure, we used both the reduced glutathione (GSH) and oxidized form of glutathione (GSSG) to determine whether GSSG induces growth at 5.0 °C (answer: it does not). **(a)** We incubated a low-density of wild-type cell population (initially ∼260 cells / mL) in various amounts of added (reduced) glutathione at 5.0 °C. The GSH concentration spans almost four orders of magnitude. Shown here is the fold-change of the population density after ∼40 days of incubation. Populations expand when the extracellular GSH concentration exceeds ∼1 μM, while populations shrink if the extracellular GSH concentration is less than ∼1 μM. Thus, low-density populations require at least 1 μM of GSH to grow at 5.0 °C. This threshold concentration is high compared to the ∼0.1 μM of GSH that the high-density populations accumulate themselves (Fig. 2a). This result suggests that there are other extracellular factors that induce population growths at 5.0 °C, because the concentration of secreted GSH is an order of magnitude below the concentration of the GSH that must be added to cause a low-density population to grow at 5.0 °C. ***(caption continues on the next page)***

Glutathione increases the % of cells that duplicates and decreases their doubling time at 5 °C

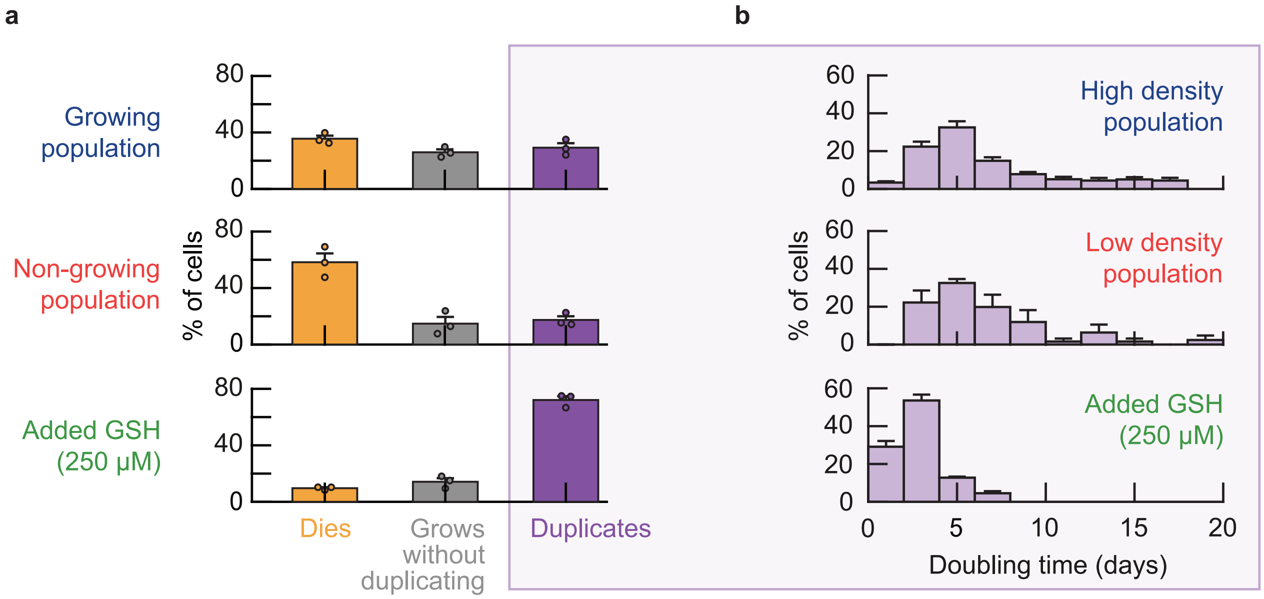


**Figure S11: Glutathione increases the percentage of duplicating cells in a population and decreases their doubling time (Related to Figure 2e).** Comparing the behavior of single cells at near-freezing temperatures with versus without added glutathione ("GSH"). **(a)** We incubated wild-type cell populations at 5.0 °C with or without adding GSH to their growth medium. After two weeks of incubation, we observed the behavior of individual cells with a microscopy time-lapse (see Methods). Shown here are the percentages of cells that duplicate (purple bar), die (orange bar) and grow without duplicating (grey bar) during the ∼20 days of microscope imaging. Three populations were imaged: high-density (growing) population (initially ∼6, 250 cells / mL), low-density (non-growing) population (initially ∼250 cells / mL), and low-density population with 250 μM of added GSH (initially ∼250 cells / mL). Note that 250 μM is higher than the ∼1 μM of glutathione that the high-density population accumulates by itself (Fig. 2a). The added GSH decreases the percentage of cells that die in the low-density population to 9.7 ± 0.7% (compared to ∼35% in high-density populations). Similarly, the added GSH increases the percentage of cells that duplicates in the low-density population to 72.1 ± 2.7% (compared to ∼17% in the low-density populations without added GSH). Thus, extracellular GSH decreases cell deaths and increases cell duplications. Bars show the mean with s.e.m., having $n$ = 3 biological replicates per condition. Dots show raw data. **(b)** Doubling time for cells that duplicate in each population. Histogram shows the percentage of cells with a given doubling time. The average doubling time is 6.5 days for the high-density populations ($n$ = 451 cells) and 7.1 days for the low-density populations ($n$ = 42 cells). Thus, the doubling time of duplicating cells is similar regardless of population density. In contrast, the average doubling time is 2.9 ± 0.1 days for the low-density populations with added GSH ($n$ = 444 cells). ***(caption continues on the next page)***

**Figure S11** ***(caption continued from the previous page)*****:** GSH therefore reduces the time it takes to duplicate by more than half. Data shows the mean with s.e.m. for each bin, having $n$ = 3 biological replicates per condition. Together (a-b) show that both high-density (growing) and low-density (non-growing) populations contain duplicating cells, with nearly identical doubling time distributions. The main differences between the two populations are the percentage of cells in each population that duplicates and the percentage of cells that die. Adding extracellular GSH increases the percentage of duplicating cells (a), and decreases the doubling time of these cells (b). Data for growing and non-growing populations in (a-b) is reproduced here from Fig. 1c for comparison.

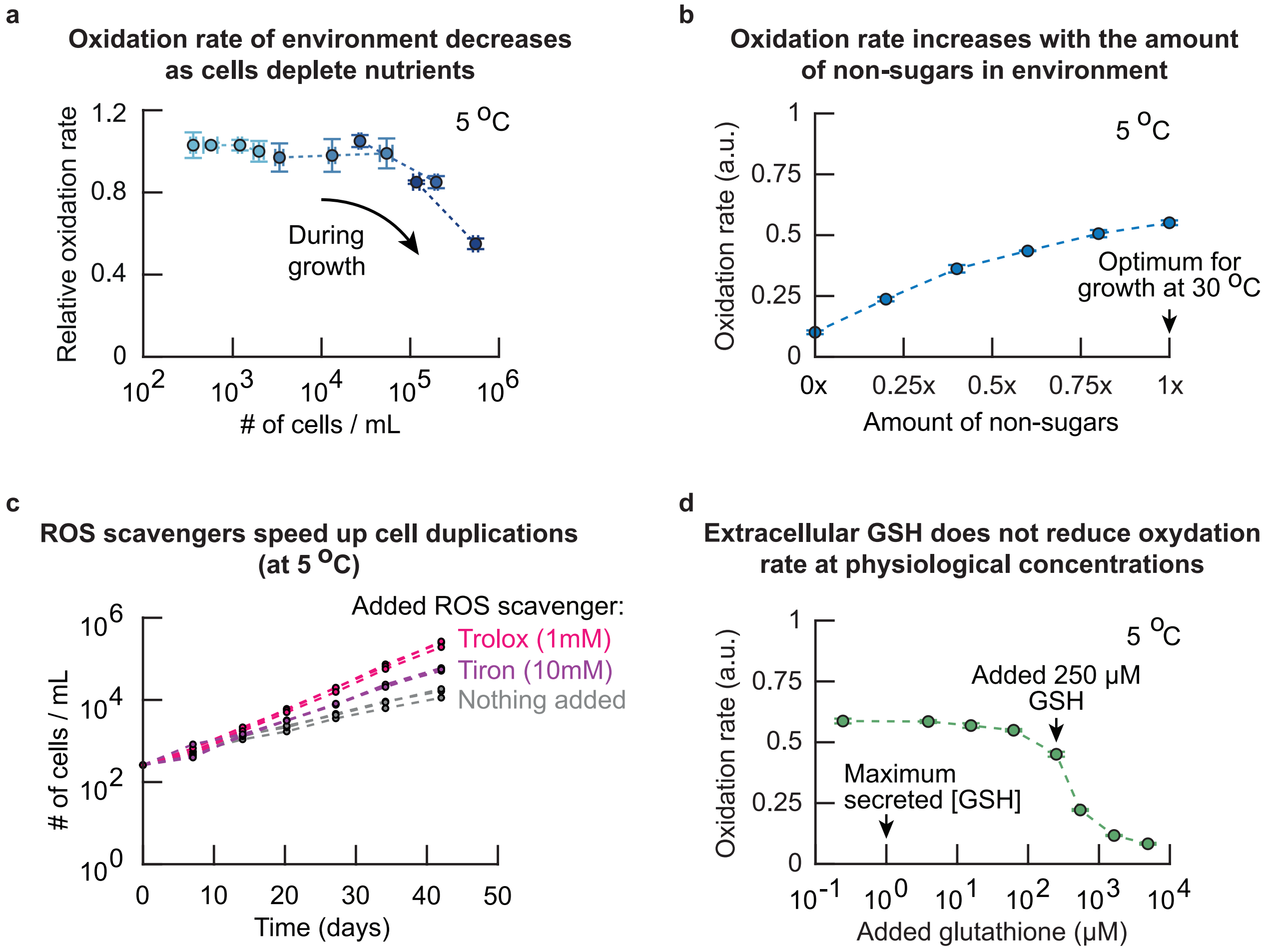


**Figure S12: Non-sugar components of growth medium (e.g., vitamins, amino acids) and oxygen create Reactive Oxygen Species without any cells (Related to Figure 2f).** Together, our results suggest that more extracellular factor(s) – besides reduced glutathione – dictate population growth at near-freezing temperatures (Supplementary Figs. S6 and S10). Knowing that extracellular factor(s) dictate growth at near-freezing temperatures (Fig. S6), cells are stressed by ROS (Fig. 2a, Supplementary Fig. S9) and that it is the antioxidant action of GSH that induces population growth (Supplementary Fig. S10), we hypothesized that the extracellular environment might also be a source of ROS. **(a)** To test whether ROS are present in the extracellular environment of our yeast populations, we used a dye called dihydroethidium (DHE) that becomes fluorescent upon oxidation by ROS (mainly superoxides). We grew wild-type yeast populations with various starting densities at 5.0 °C, and measured the oxidation rate of the growth media relative to the oxidation rate of fresh media. To do so, at every timepoint, we took aliquots of our liquid cultures and flowed the aliquot through a 0.2 μm pore filter. We then measured the oxidation rate in the flow-through – the growth medium without any cells – and compared with the oxidation rate in fresh medium. Shown is the relative oxidation rate as function of population density. ***(caption continues on the next page)***

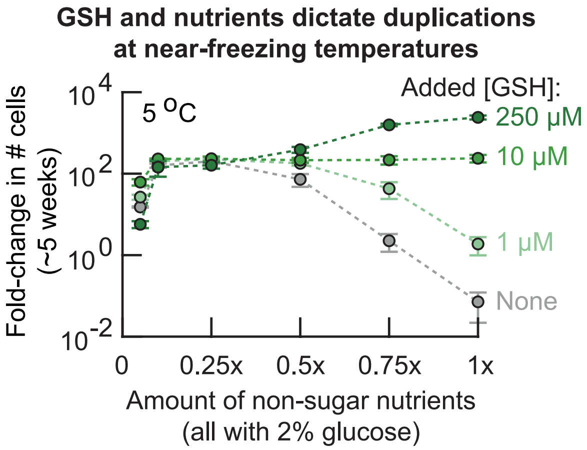


**Figure S13: Ability of cells to duplicate is a balance between extracellular GSH concentration and availability of non-sugar nutrients (Related to Figure 2i).** Studying the combined effect of non-sugar nutrients and extracellular GSH on population growth. We incubated cells in media that had precise amounts of non-sugar nutrients and added GSH. Specifically, for varying the amount of non-sugar nutrients, we formulated the media to contain 0.05x, 0.1x, 0.25x, 0.5x, 0.75x or 1*x* non-sugar nutrients. Here, 1x equals the amount of non-sugar nutrients that is in the medium for yeast grown under standard conditions (see Methods). Each of these media had 2% glucose. Then, for each of these medium compositions, we added either 0 μM, 1 μM, 10 μM or 250 μM of GSH. This procedure created 24 different media (6 different amounts of non-sugar nutrients each with 4 different added GSH concentrations). In these media we incubated low-density populations of cells (initially ~210 cells / mL) at 5.0 °C. These low-density populations should not grow according to our phase diagram (Fig. 1b). Shown is the fold-change in population density as function of the amount of non-sugar nutrients after ~5 weeks of incubation at 5.0 °C. The curves represent 1 μM (light green curve), 10 μM (green curve), 250 μM (dark green curve) or without (grey curve) added GSH. The population in 1x non-sugar nutrients and without added GSH did not grow at all during the 5 weeks of incubation at 5.0 °C, as expected from our phase diagram (Fig. 1b). Further corroborating our other data, we find that at 1x non-sugar nutrients the fold-change in population density increases with the concentration of added GSH (also see Fig. 2e) and that without added GSH, the fold-change in population density increases when the amount of non-sugar nutrients decreases (grey curve, also see Fig. 2g). Then, starting at 1x non-sugar nutrients, all curves converge to a ~100-fold change in population density as the amount of non-sugar nutrients decreases from 1x to 0.25x. Thus, adding more than 10 μM GSH is only beneficial for population growth when sufficient non-sugar nutrients are available (i.e., more than 0.5x non-sugar nutrients). Finally, populations stop growing when the amount of non-sugar nutrients decreases below 0.1x. ***(caption continues on the next page)***

**Figure S13** ***(caption continued from the previous page)***: In summary, this growth experiment shows that either sufficient extracellular GSH ($>$ 1 μM, Supplementary Fig. S10) or low enough amounts of non-sugar nutrients are required for population growth at 5.0 °C. Populations having less non-sugar nutrients require less extracellular GSH for growth, and populations having more extracellular GSH can grow with more non-sugar nutrients. The media with $>$ 0.75x non-sugar nutrients and 250 μM added GSH are special because they permit growths that exceed the ($\sim$100-fold) growths that a population can achieve by secreting $<$ 1 μM glutathione (Fig. 2a) by itself (light green curve). Error bars show the mean with s.e.m., having $n$ = 3 biological replicates per condition.

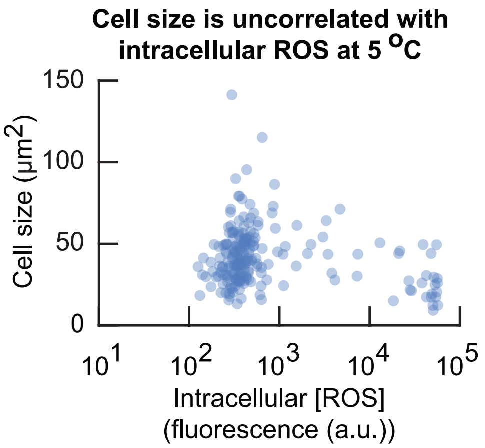


**Figure S14: Cell size does not correlate with the intracellular [ROS] at 5 °C (Related to Figure 3c).** Cell size as a function of intracellular ROS concentration. We incubated populations of cells for two weeks at 5.0 °C (initially ∼6,250 cells / mL). We then measured the size and intracellular ROS concentration in single cells at 5.0 °C (see Methods). This result shows that the cell size is uncorrelated with intracellular ROS concentration. Dots show single-cell data aggregated from $n$ = 3 biological replicates.

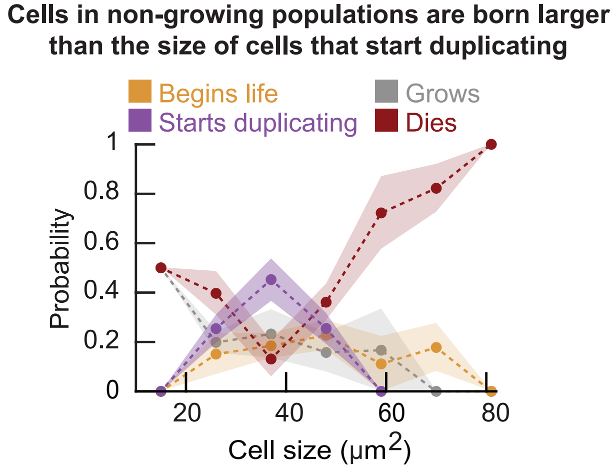


**Figure S15: Newborn cells in low-density (non-growing) populations are larger than those in high-density (growing) populations (Related to Figure 3c).** We incubated populations of wild-type cells at a low density (initially ∼500 cells / mL) for two weeks at 5.0 °C and then imaged these cells in our wide-field microscope for the next several weeks (see Methods). We classified individual cells in the resulting time-lapse movies as being in one of the following classes (see Supplementary Fig. S16): ”begins life” (just-born daughter cell), ”starts duplicating”, ”grows” (without duplicating), or ”dies” (cell just before death). For every cell in each class, we measured its cell size. Specifically, for cells the cells in the ”Begins life” class, we measured the daughter cell's size just after cytokinesis (when the mother's bud neck breaks). For the cells in the ”Starts duplicating” class, we measured the mother cell's size at the time the bud appears. For the cells in the ”Grows” class, we measured their size in the first time point when the movie begins (i.e., while they are in growing). For the cells in the ”Dies” class, we measured their size just before they die (in the last time point before death). We binned the cells according to their sizes. For each bin, we determined the probability that a cell in that size bin would be in each of the four classes. Dots show average probability in each bin for the different events, shaded area represents the s.e.m. for $n$ = 3 biological replicates. This graph shows that newborn cells in low-density (non-growing) populations are on average larger than the cells that duplicate in the same population. Thus, newborn cells in low-density (non-growing) populations are less likely to duplicate than the mother cells in the same population (also see Supplementary Fig. S16).

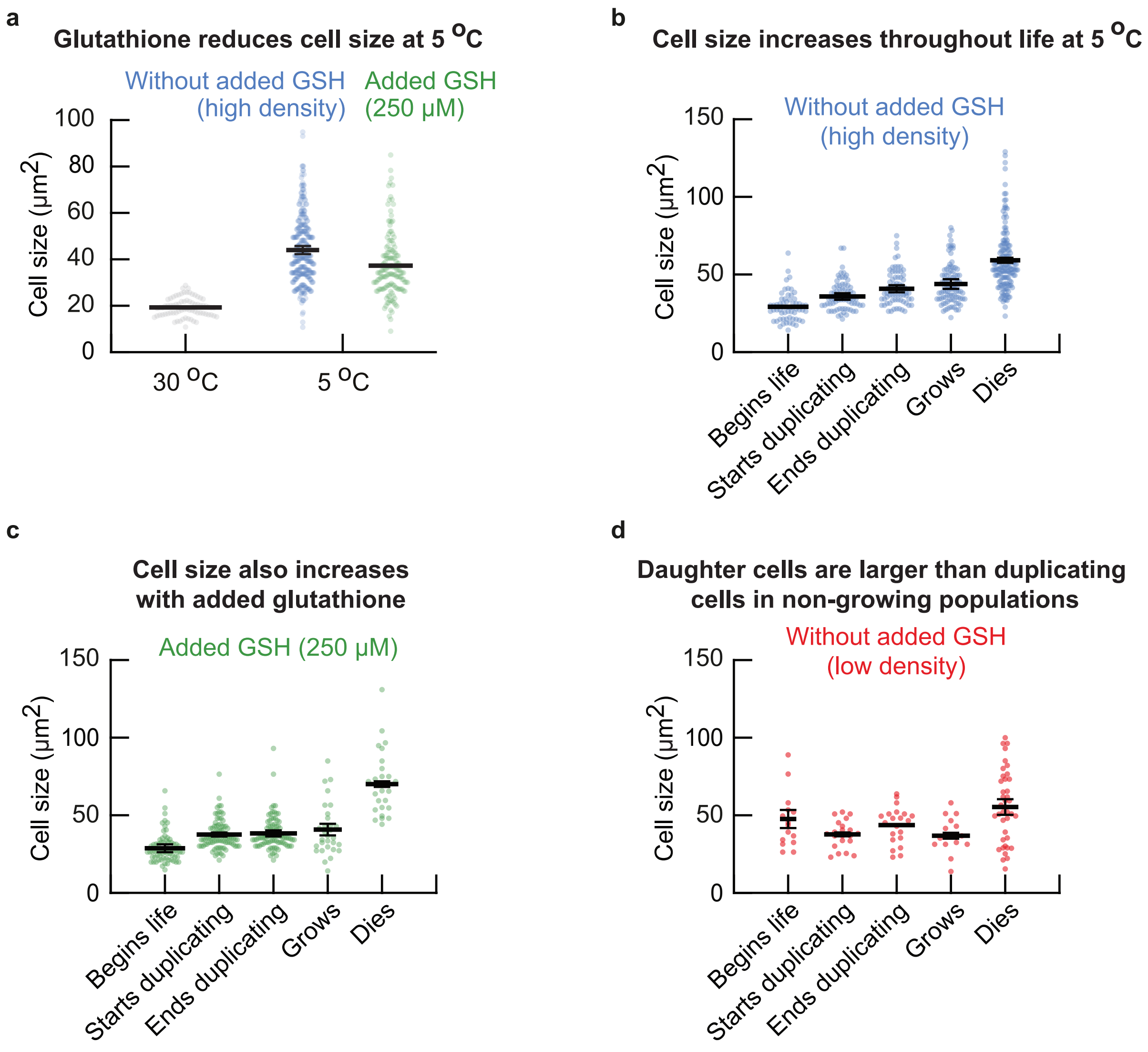


**Figure S16: Cell size monotonically increases over time throughout a cell's life at 5 °C (Related to Figure 3c). (a)** We incubated wild-type cell populations for two weeks at 5.0 °C and then measured the cell sizes with a wide-field microsope. Shown here are the cell sizes of individual cells from high-density populations without any GSH added (blue dots, initially ~8,000 cells / mL) and sizes of individual cells from low-density populations that were incubated with 250 μM GSH (green dots, initially ~420 cells / mL). Grey dots show cells in a log-phase growth at 30.0 °C for comparison. Surprisingly, we find that cells are, on average, more than double the size (cross-sectional area) at 5.0 °C ($44.0 \pm 1.7$ μm$^2$) compared to 30.0 °C($19.3 \pm 0.2$ μm$^2$). Moreover, adding GSH to the growth medium reduces the cell size at 5.0 °C ($37.3 \pm 0.1$ μm$^2$). Error bars represent the mean with s.e.m., having $n = 3$ biological replicates. Dots show data aggregated from the biological replicates. **(b-d)** Next, we incubated populations of wild-type cells for two weeks at 5.0 °C. ***(caption continues on the next page)***

**Figure S16** ***(caption continued from the previous page)***: After two weeks we transferred aliquots of the liquid cultures to microscopy plates that we kept chilled at 5.0 °C before the transfer and took a snapshot with a microscope every day (see Methods). The resulting snapshots form time-lapse movies at 5.0 °C. From these movies, we classified the cells into one of the following events: ”begins life” (size of the daughter cell right after the mother cell finishes cytokinesis), ”start duplicating” (size of the mother cell right after it forms a bud), ”end duplicating” (size of the mother cell right after the mother cell finishes cytokinesis), ”grows” (cells that did not duplicate during the entire time-lapse movie of ~17 days) and ”dies” (cell size just before dying). For all cells in each class, we determined the cell size at the start of each event. Shown here are the cell sizes for populations with high initial density (~8,000 cells / mL) **(b)**, low initial density (~420 cells / mL) with 250 μM added GSH **(c)**, and low initial density (~420 cells / mL) without any GSH added **(d)**. Dots show size of individual cells in each class. For all panels, the error bar shows the mean with s.e.m., having $n$ = 3 biological replicates. Dots are data aggregated from all biological replicates. (b) At high density, newborn cells are the smallest cells in the population (~29.2 ± 0.5 μm$^2$), with cells increasing in size during duplications (from ~35.8 ± 2.1 μm$^2$ at bud formation to ~40.8 ± 2.3 μm$^2$ at cytokinesis). Growing cells are on average 43.8 ± 3.1 μm$^2$. Finally, cells that die are the largest cells in the population (~59.1 ± 1.6 μm$^2$). (c) Similarly, in populations that were incubated in growth medium to which we added 250 μM GSH, we found that newborn cells are the smallest cells (~28.7 ± 2.5 μm$^2$) in the population, with duplicating cells increasing in their size during duplications (starting from ~37.6 ± 1.3 μm$^2$ and ending with ~38.3 ± 2.0 μm$^2$ at the end of division). Growing cells are on average 40.8 ± 3.7 μm$^2$. Finally, with GSH added, cell deaths occur at larger sizes (~70.1 ± 1.8 μm$^2$) compared to cells without added GSH. (d) In non-growing populations (low population density), the newborn cells are on average larger (~47.6 ± 5.8 μm$^2$) than the average size at which the cells start duplicating (~37.9 ± 1.2 μm$^2$), end duplicating (~43.7 ± 0.4 μm$^2$), or are growing (~36.8 ± 1.9 μm$^2$). Finally, cells that die are smaller (~55.3 ± 5.0 μm$^2$) than cells in the growing populations. Together, these results show that newborn cells are the smallest cells in the population. Moreover, the cell size increases monotonically during cell duplications with cells that die being the largest cells in the population. Furthermore, our observations suggest that adding extracellular GSH keeps the cell size approximately constant during cell duplications, but the cell size does increase when cells die. Finally, in low-densty (non-growing) populations without added GSH, the newborn cells are larger than the duplicating cells in the population.

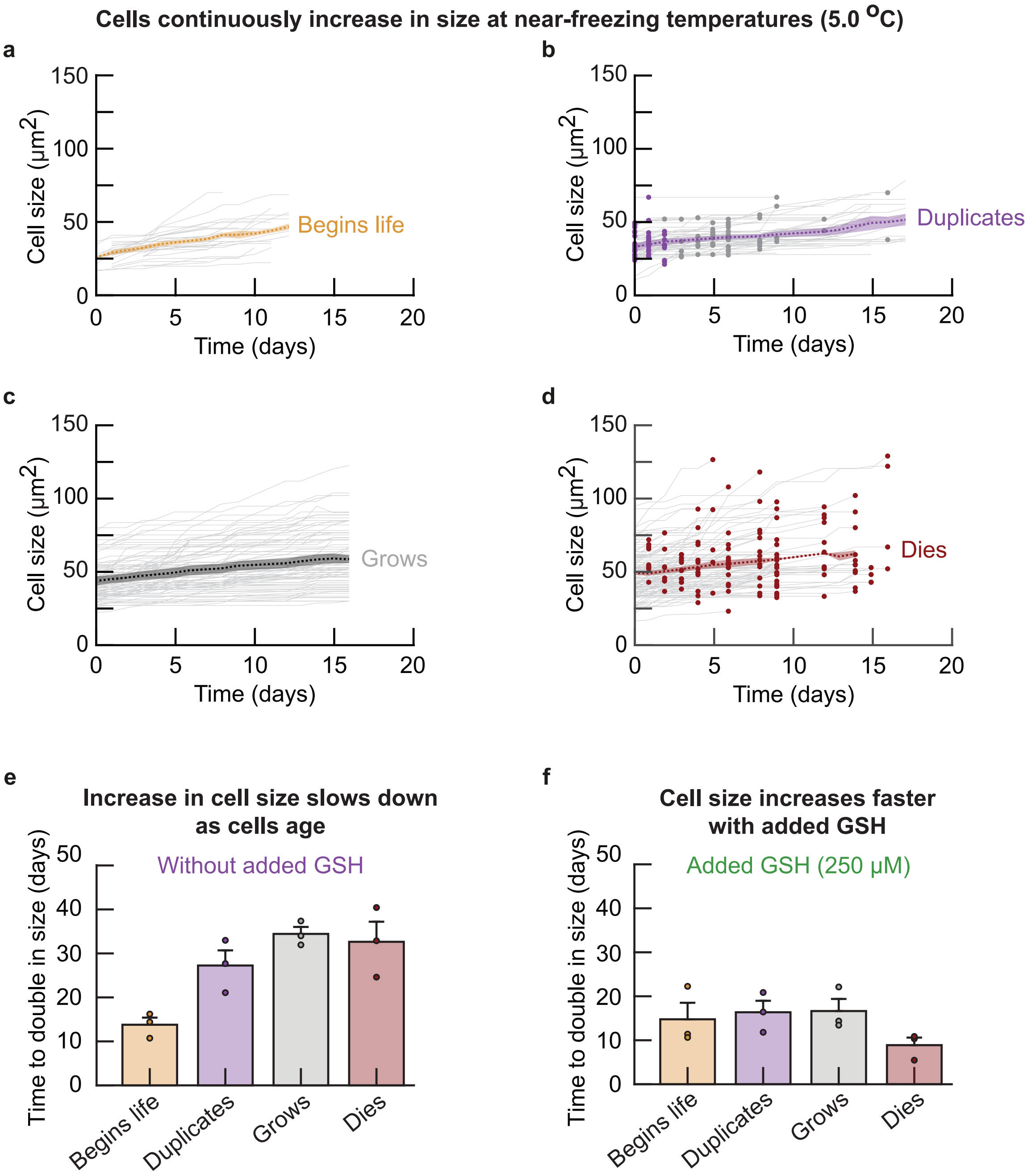


**Figure S17: Cell size monotonically increases over time throughout a cell's life at 5 °C (Related to Figure 3c).** Our data shows that the cell size continuously increases at 5.0 °C starting with the smallest cell size at birth and ending with the largest cell size at death, with a continuous increase in size in between the two events (including during cell duplications, Supplementary Fig. S16). We used time-lapse microscopy to measure how the cell size changes in indiviual cells at 5.0 °C. ***(caption continues on the next page)***

**Figure S17** ***(caption continued from the previous page)*****:** **(a-d)** We used our time-lapse movies of cells in high-density populations (initially $\sim$8,000 cells / mL) to track the cell size over time in single cells and for all cells that were present at the start of the time-lapse movies. To reconstruct every stage of life for cells at near-freezing temperatures (e.g., 5.0 °C here and 1.0 °C shown in a later figure), we categorized all cells in the first time point of our movies into one of four classes (see Supplementary Fig. S16): newborn daughter cells (labelled, ”begins life”, just after cytokinesis), cells that start to duplicate (”duplicates”), cells that did not duplicate for the entire duration of the time-lapse ($\sim$17 days, ”grows”) and cells that died (”dies”). We assumed that the cells in the first time point of the movie are representative samples of populations at 5.0 °C. This is because the cell populations that we imaged were in a steady-state growth when we started the time-lapse of the cells, due to the fact that the population was already incubated at 5.0 °C for two weeks before the first frame of the movie. As an exception, since we did not know which cells were newborn cells when the time-lapse began (”day 0”), we used daughter cells that were newly born within a three-day period during our time-lapse movie. Shown here is the cell size over time for each class: size of a newborn cell **(a)**, size of a cell that begins to duplicate (i.e., when its bud begins to grow) **(b)**, size of a growing cell that does not duplicate **(c)**, and size of a cell just before it dies (almost always by bursting open **(d)**. Grey curves show the cell size over time for individual cells. Blue and grey dots in (b) indicate the time points at which bud formation and cytokinesis occurred respectively. Red dots in (d) indicate the cell size just before death. The dotted line shows the average size of cells each respective class for $n$ = 3 biological duplicates, with the shaded area representing the s.e.m. of the average cell size from each biological replicate. Together, (a-d) show that the cells continuously increase their size at 5.0 °C. Almost all cells continuously increase in size during the entire duration of the time-lapse movies. Given that a newborn cell will either eventually duplicate or will grow without duplicating and that every cell will eventually die, these data together show that cells continuously increase in their size throughout their life at 5.0 °C. **(e-f)** From the measurements in (a-d), we determined the time that cells take to double in size without **(e)** and with 250 μM added GSH **(f)**. Shown here are the time taken to double in size for cells right after cytokinesis (yellow bars), cells that begin to duplicate (purple bars), cells that grow without duplicating (grey bars), and cells just before they die (red bars). Dots show the average time for each biological replicate. Error bars show the mean with s.e.m. of $n$ = 3 biological replicates. These measurements show that, without added GSH, cells take increasingly more time to double in size. This time is the shortest for a newborn cells (13.8 $\pm$ 1.6 days) and increases in the following order: cells that start to duplicate (27.3 $\pm$ 3.4 days), cells that are growing without duplicating (34.4 $\pm$ 1.6 days), and cells that are about to die (32.6 $\pm$ 4.6 days). In contrast, with added GSH, cells take approximately constant time to double in size for newborn cells (14.8 $\pm$ 3.8 days), cells that begin to duplicate (16.4 $\pm$ 2.6 days), and cells that are growing without duplicating (16.6 $\pm$ 2.8 days). Lastly, cells that die take the shortest time to double in size when there is 250 μM of added GSH (8.9 $\pm$ 1.7 days). Together, (e-f) show that cells double in size more slowly as they (chronologically) age, and that adding GSH accelerates cell growth.

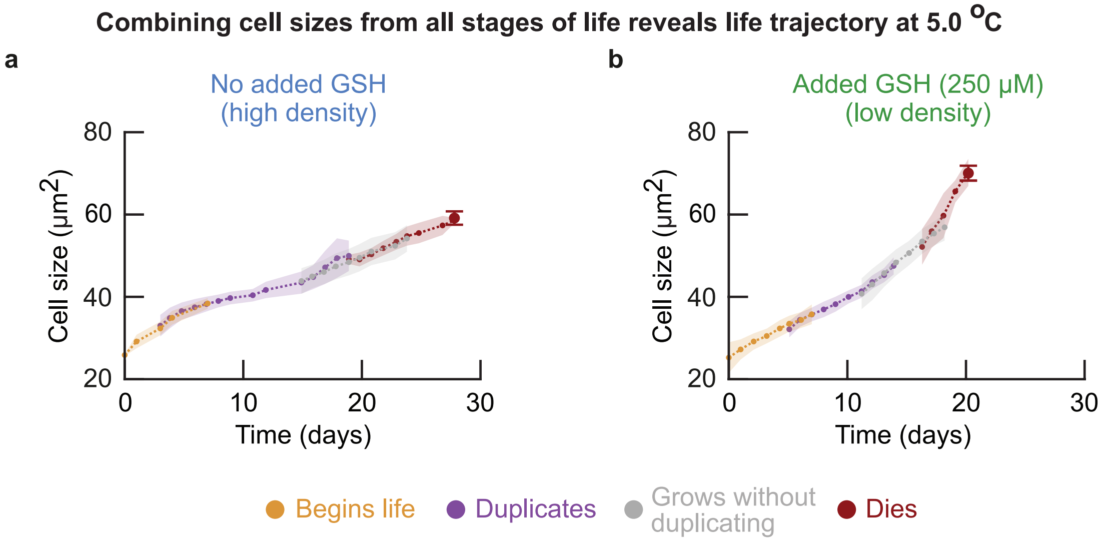


**Figure S18: Continuous cell size growth from each life event reveals "life trajectory" of cells at 5 °C (Related to Figure 3d).** We have now established that cells continuously increase in size throughout their life at 5.0 °C (Supplementary Fig. S17). We also found that the cell size increases with consecutive life events (cells in "begins life" are smaller than cells in "duplicates", followed by "grows" and "dies", Supplementary Fig. S16). Moreover, since a newborn cell will either eventually duplicate or will grow without duplicating, and that every cell will eventually die, there exists a temporal order to these life events. We therefore used the cell size over time for each class to reconstruct the life of cells at 5.0 °C as function of cell size. Specifically, we stitched together the cell size curves of the average cell for each of the consecutive events (based on the order of average cell size: "begins life", "duplicates", "grows" and "dies") by connecting each of the curves with its preceeding curve at the time that the cell sizes at the respective endpoints overlapped most. Shown here are the stitched curves for populations of cells that were incubated without **(a)** and with 250 µM added GSH **(b)**. Each curve overlaps with at least three data points with neighboring curves. The colors of each section indicates the separate life events, showing a cell that begins life (yellow bars), duplicates (purple bars), grows without duplicating (grey bars), and dies (red bars). Large red dot shows average size at just before cell death. Dots show the mean, with shaded areas representing the s.e.m. of $n$ = 3 biological replicates.

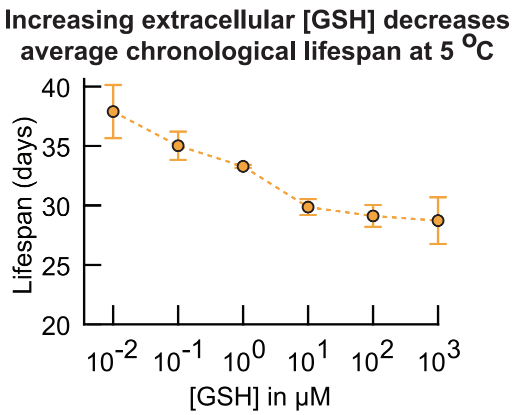


**Figure S19: Average chronological lifespan decreases as the added GSH concentration increases at 5 °C (Related to Figure 3d).** Cell's average chronological lifespan (average chronological age at death) as a function of added extracellular GSH concentration. We incubated wild-type cell populations at 5.0 °C with various amounts of GSH added to the growth medium (all initially at ∼5,680 cells / mL). We took an aliquot of each liquid culture every week and measured the number of alive and dead cells using propidium iodide (see Methods). We then fitted these measurements to the same growth model as before to determine a cell's average chronological lifespan which is plotted here (Supplementary Fig. S5, also see Supplementary Theory). The average chronological lifespan of cells at 5.0 °C decreases from ∼37 days without added GSH to ∼30 days with 100 μM added extracellular GSH. Extracellular GSH therefore decreases the chronological lifespan of cells at near-freezing temperatures. Error bars represent the mean with s.e.m., having $n$ = 4 biological replicates per data point.

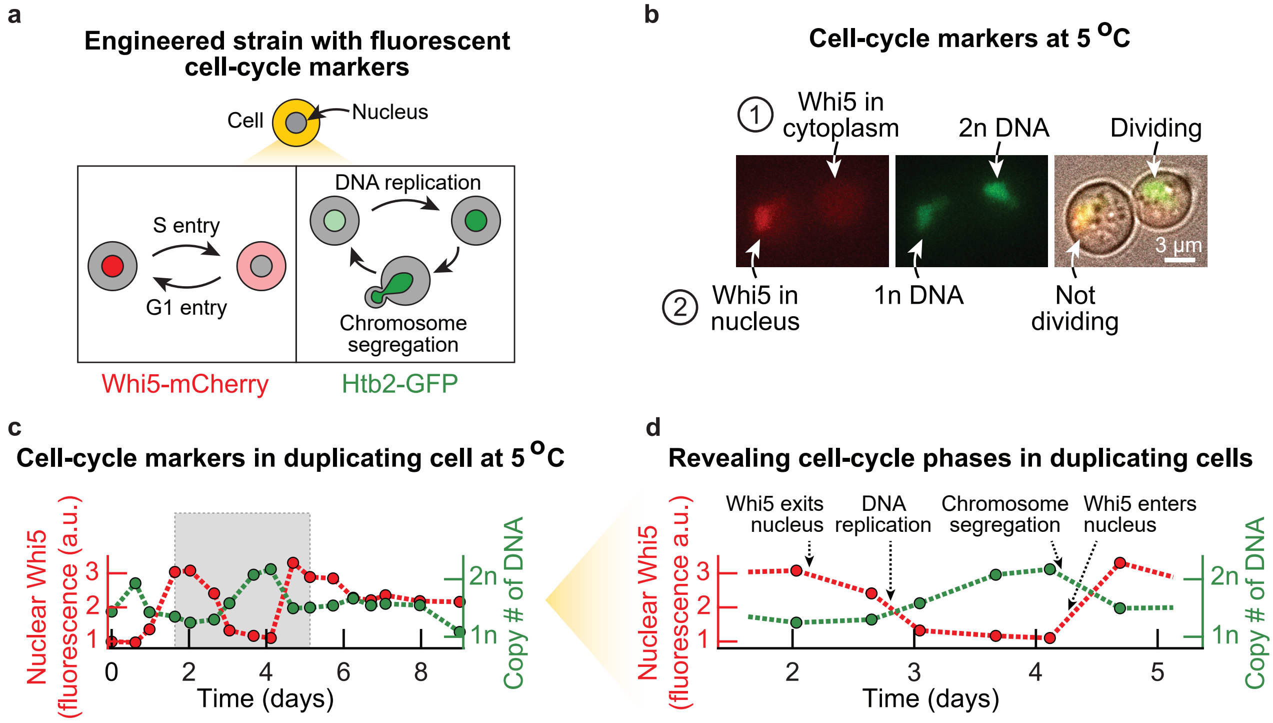


**Figure S20: Time-lapse microscopy on engineered cells with fluorescent reporters for DNA replication (H2B histone fused to GFP) and G1-to-S transition (Whi5 protein fused to mCherry) reveals cell-cycle phases at 5 °C (Related to Figure 4). (a)** Schematic of the engineered strain used for identifying when key cell-cycle phases occur in single cells at 5.0 °C. We engineered yeast to have the G1 transcriptional repressor Whi5 protein fused to mCherry protein (Whi5-mCherry), and the histone protein, H2B (Htb2), fused to GFP (H2B-GFP), also see Methods). Both Whi5 and H2B are well-known cell-cycle markers that indicate G1 phase (Whi5 translocates to the nucleus) and the S-G2-M phases (the amount of H2B is proportional to the amount of DNA) respectively [6, 7, 8]. We used this engineered strain to track the cell cycle at 5.0 °C. **(b)** Microscopy snapshots of the engineered strain. Nuclear fluorescence from Whi5-mCherry (red, left image) and H2B-GFP (green, middle) are shown for two representative cells at 5.0 °C. Also shown is the composite image that includes brightfield (right). Here, one cell is dividing which we can tell, aside from the brightfield image, by finding almost all of the Whi5-mCherry in the cytoplasm and a bright nucleus with H2B-GFP (contains 2n DNA, labelled ”1” in the top row). In this picture, we see that the other cell is not dividing, which we can also tell by finding almost all the Whi5-mCherry to be localized in the nucleus and a dim nucleus (contains 1n DNA, labelled cell ”2” in the bottom row). The dividing cell has a bud. Scale bar is 3 μm. **(c)** Nuclear Whi5-mCherry and H2B-GFP over time shown for a duplicating cell at 5.0 °C. This cell is from a population that was pre-incubated for two weeks at 5.0 °C before starting the timelapse, after which we tracked cells in these populations with a microscope (initially at ∼6,250 cells / mL). We took aliquots of our liquid cultures that we transferred to an imaging plate. *(caption continues on the next page)*

**Figure S20** ***(caption continued from the previous page)*****:** We then kept this plate at 5.0 °C and imaged the cells every ∼12 hours. Finally, we used this microscopy time-lapse to quantify the amount of nuclear Whi5-mCherry (left y-axis, red) and H2B-GFP (righ y-axis, green) over time. The grey area indicates a full cell-cycle, starting from G1 (all of Whi5-mCherry in the nucleus). Specifically, we used the fluorescence of H2B-GFP to locate the nucleus and quantify the amount of DNA in the cells. We first located the nucleus by segmenting the GFP fluorescence of each cell using a threshold GFP fluorescence that we kept fixed for all cells and time points. The nucleus was then the group of pixels whose fluorescences exceeded this threshold. To determine the copy number of DNA in a cell, we determined the total GFP fluorescence within the cell's nucleus. We subsequently rescaled the nuclear GFP between the average minimum and maximum GFP fluorescence that we observed for duplicating cells. Thus, the amount of nuclear H2B was rescaled to a scale with "1n DNA" (average GFP fluorescence in the nucleus of duplicating cells in G1) and "2n DNA" (average GFP fluorescence in the nucleus of duplicating cells in G2). Finally, we used the fluorescence of Whi5-mCherry to quantify the relative amount of nuclear Whi5. To do so, we determined the average mCherry fluorescence in the nucleus and cytoplasm of each cell. The amount of nuclear Whi5 was then given by the ratio of measured nuclear and cytoplasmic mCherry fluorescence. **(d)** Nuclear Whi5 and H2B mark the cell-cycle phases of duplicating cells. Shown is a cell during one cell division (from the grey box of the duplicating cell in (c)). Dotted arrows indicate the end of G1 (Whi5-mCherry exits the nucleus), the S phase (the amount of H2B-GFP increases during replication of DNA), the M phase (H2B-GFP decreases during chromosome segregation) and the start of G1 (Whi5-mCherry begins to enter the nucleus).

**Growing cells that never duplicate and dying cells have Whi5 localized in their nuclei or have 2n DNA at 5 °C**

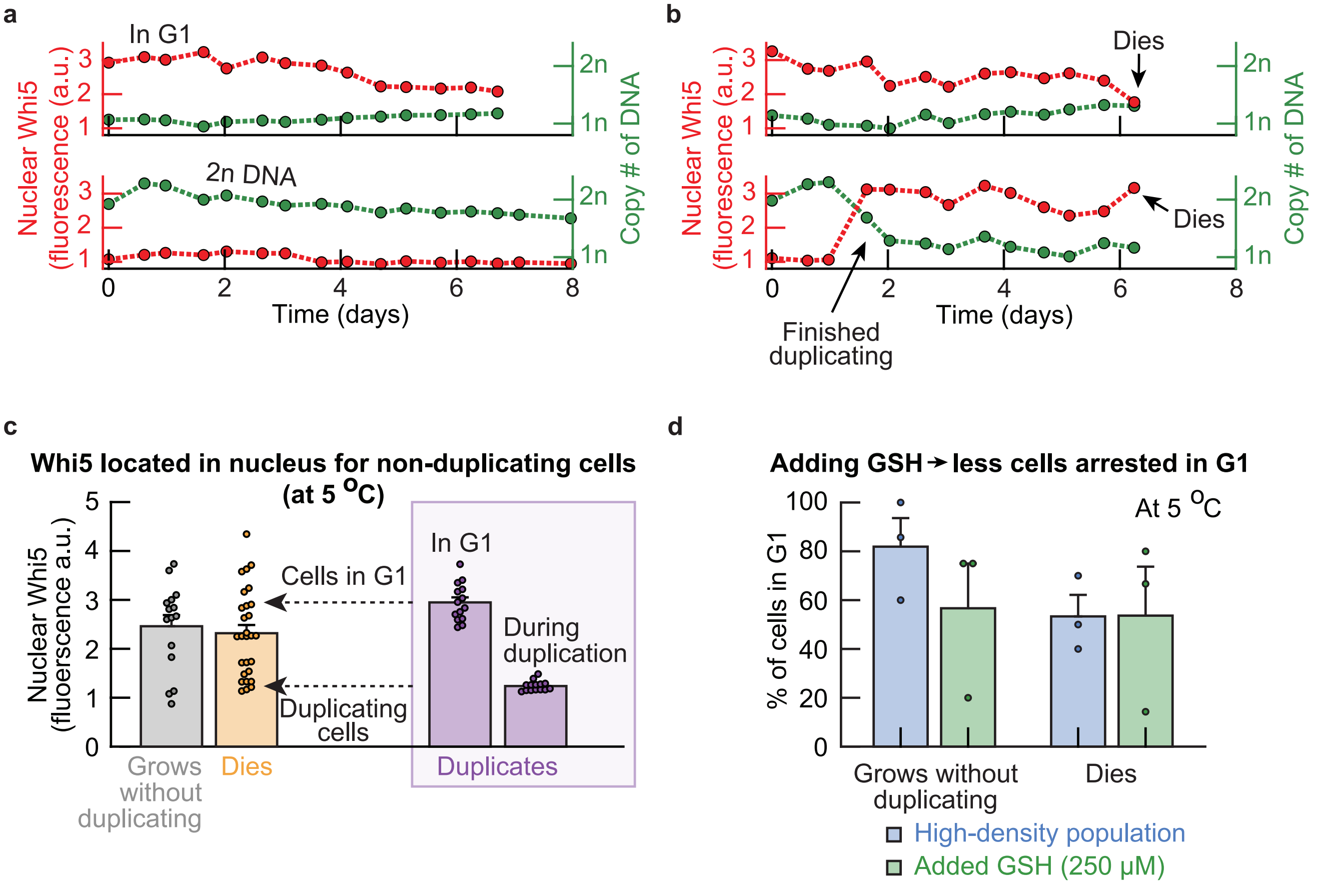


**Figure S21: Growing cells that never duplicate until dying are arrested in G1 phase at 5 °C (Related to Figure 4c). (a-b)** Nuclear Whi5-mCherry and H2B-GFP over time in growing cells that never duplicate until dying and dying cells at 5.0 °C (also see Supplementary Fig. S20). Shown are cells that will never duplicate in G1 or G2 phase **(a)**, and cells just before they die in G1 **(b)**. In (a-b), the amount of nuclear Whi5-mCherry and the amount of DNA remain constant in a growing cell that never duplicates during many days of incubation. **(c)** Cells that will never duplicate and dying cells often have Whi5 located in the nucleus. Shown is the average amount of nuclear Whi5-mCherry in non-duplicating cells (grey bar), in dying cells (orange bar) and for duplicating cells (purple box) when in G1 (left purple bar) or during cell division (S-G2-M phases, right purple bar). Data suggests that most cells that never duplicate stay in G1 (their Whi5-mCherry always stays in the nucleus, compared to the cells in G1 that will duplicate). The average amount of nuclear Whi5-mCherry in cells that will never duplicate was determined by averaging the amount of nuclear Whi5-mCherry for the entire duration of the time-lapse. The average amount of nuclear Whi5-mCherry in dying cells represents the amount of nuclear Whi5-mCherry averaged over the last three frames (~36 hours) before the cell dies. *(caption continues on the next page)*

**Figure S21** ***(caption continued from the previous page)***: The average amounts of nuclear Whi5-mCherry for cells that can duplicate and are in G1 or are dividing were determined as follows. We determined when Whi5-mCherry was located in the cytoplasm (division) or the nucleus (G1). We then averaged the amount of nuclear Whi5-mCherry for each cell while the cell is either in G1 or duplicating (S-G2-M). Dots show raw data ($n$ = 15 non-duplicating cells, $n$ = 28 dying cells, $n$ = 14 duplicating cells). Bars show the mean with s.e.m. of all cells. **(d)** Cells that never duplicate and dying cells mostly remain in G1. Shown here are the percentages of cells in G1 for high-density (growing) populations (blue bars) and low-density populations supplemented with 250 μM GSH (green bars). This plot shows that 80% of the cells that never duplicate remain in G1 until they die. Extracellular GSH decreases the percentage of these cells in G1. Whether a cell is in G1 or not was determined with the average amount of nuclear Whi5-mCherry (in (c)) and the amount of DNA in the cell (a cell was in G1 when the amount of nuclear Whi5-mCherry was above 1.66 and when the amount of DNA was below 2n). Bars show the mean with s.e.m., having $n$ = 3 biological replicates per condition. Dots show raw data.

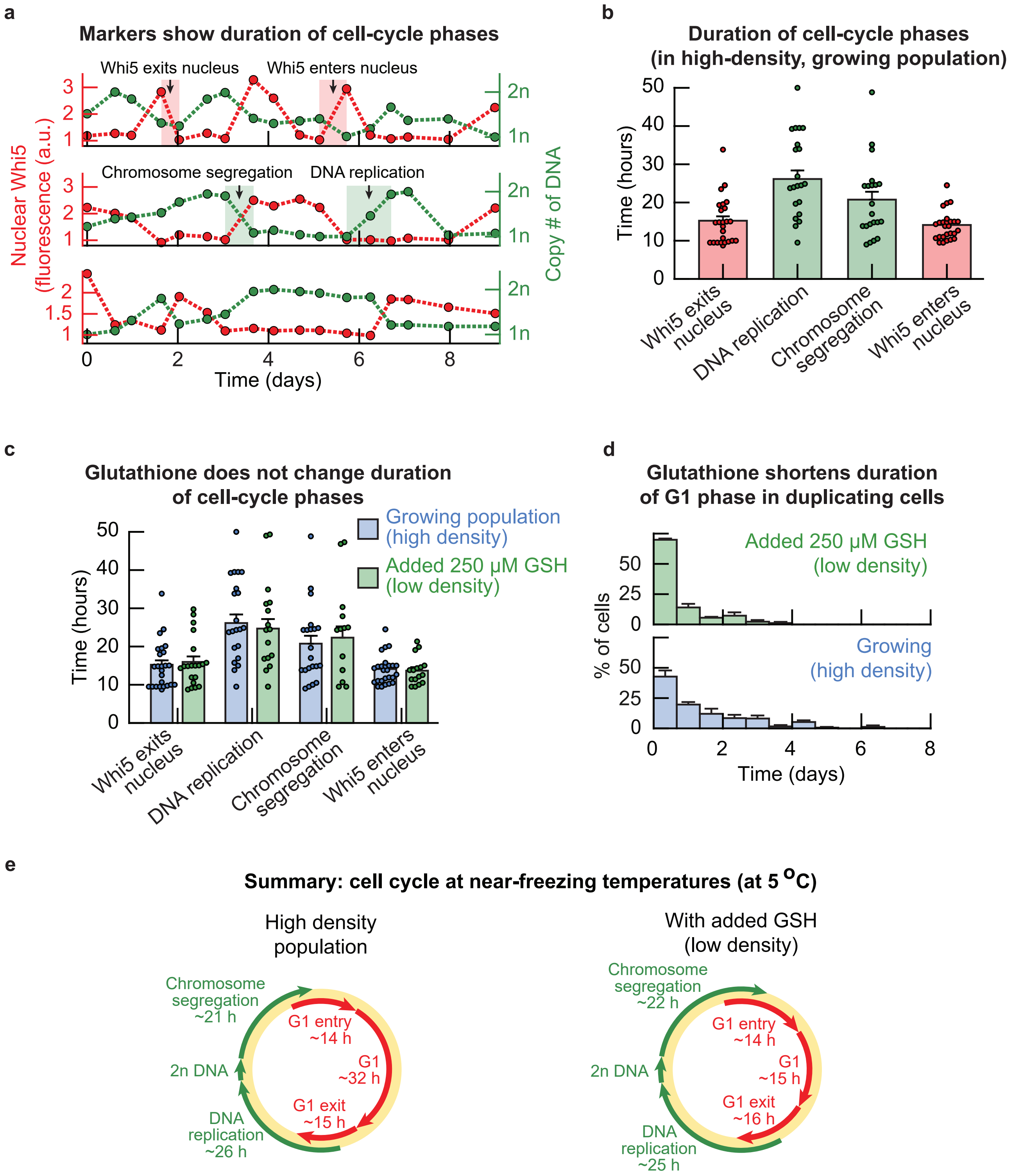

a
Markers show duration of cell-cycle phases
Whi5 exits nucleus
Whi5 enters nucleus
Chromosome segregation
DNA replication
Nuclear Whi5 (fluorescence (a.u.))
Copy # of DNA
1n
2n
Time (days)
b
Duration of cell-cycle phases (in high-density, growing population)
Time (hours)
Whi5 exits nucleus
DNA replication
Chromosome segregation
Whi5 enters nucleus
c
Glutathione does not change duration of cell-cycle phases
Growing population (high density)
Added 250 μM GSH (low density)
Time (hours)
Whi5 exits nucleus
DNA replication
Chromosome segregation
Whi5 enters nucleus
d
Glutathione shortens duration of G1 phase in duplicating cells
Added 250 μM GSH (low density)
Growing (high density)
% of cells
Time (days)
e
Summary: cell cycle at near-freezing temperatures (at 5 °C)
High density population
Chromosome segregation ~21 h
G1 entry ~14 h
G1 ~32 h
2n DNA
G1 exit ~15 h
DNA replication ~26 h
With added GSH (low density)
Chromosome segregation ~22 h
G1 entry ~14 h
G1 ~15 h
2n DNA
G1 exit ~16 h
DNA replication ~25 h

**Figure S22: Glutathione shortens duration of G1 phase, but not S-G2-M phases, at 5 °C (Related to Figure 4c-d). (a)** Nuclear Whi5-mCherry and H2B-GFP over time in duplicating cells that were incubated at 5.0 °C. Red shaded areas indicate examples of the duration of Whi5-mCherry exiting or entering the nucleus. Green shaded areas indicate examples of the duration of DNA replication or chromosome segregation. **(b)** Using time-lapse movies that examined the amounts of Whi5-mCherry and H2B-GFP in single cells, we determined the duration of cell-cycle events in duplicating cells (following the methodology in (a)). Shown here are the durations of Whi5-mCherry exiting the nucleus (15.2 $\pm$ 1.1 hours, red bar), DNA replication (26.1 $\pm$ 2.2 hours, green bar), chromosome segregation (20.8 $\pm$ 2.1 hours, green bar), and Whi5-mCherry entering the nucleus (14.1 $\pm$ 0.8 hours, red bar). Since the time-lapse movies consists of snapshots taken every $\sim$12 hours, this $\sim$12 hours is a lower bound on the true duration of each event. Dots show raw data, having $n$ = 28 cells for the location of Whi5-mCherry, $n$ = 23 cells with DNA replication and $n$ = 24 cells with DNA segregation. Bars show the mean with s.e.m. of all cells. **(c)** Duration of cell-cycle events in cells from low-density populations that were incubated with 250 μM of GSH (green bars). Blue bars show the same data as in (b) for comparison. Shown here are, all with the added GSH, Whi5-mCherry exiting the nucleus (16.0 $\pm$ 1.4 hours), DNA replication (24.7 $\pm$ 2.5 hours), chromosome segregation (22.3 $\pm$ 2.9 hours), and Whi5-mCherry entering the nucleus (13.7 $\pm$ 0.9 hours). There is no reason to assume that the duration of any cell-cycle event changes upon addition of extracellular GSH (p-values are 0.69 for Whi5-mCherry exiting the nucleus, 0.69 for DNA replication, 0.66 for chromosome segregation and 0.72 for Whi5-mCherry entering the nucleus). Blue dots are as in (b). Green dots show raw data, having $n$ = 16 cells for Whi5-mCherry entry and $n$ = 20 cells for Whi5-mCherry exit, $n$ = 17 cells with DNA replication and $n$ = 16 cells with DNA segregation. Bars show the mean with s.e.m. of all cells. **(d)** Duration of G1 in duplicating cells from populations without (blue) and with 250 μM added GSH (green). The average duration of G1 was 32 $\pm$ 3.0 hours without added GSH, and 15.0 $\pm$ 1.5 hours with added GSH (median $\sim$24 hours without and $\sim$9 hours with added GSH). Thus, GSH shortens the duration of G1 in duplicating cells at near-freezing temperatures. With added GSH, the majority of G1 phases in duplicating cells take 12 hours or less. In contrast, the G1 phase in duplicating cells takes up to 6 days without added GSH. Error bar shows mean with s.e.m. of $n$ = 3 biological replicates. **(e)** Summarizing (b-d). Shown are the durations of the cell cycle phases in duplicating cells in populations without (left) and with (right) added GSH. Length of arrows are proportional to the duration of each phase. Red arrows represent Whi5 (indicating events associated to G1), green arrows represent H2B (indicating events associated to S-G2-M). Together, (b-d) show that adding GSH to the growth medium shortens the duration of G1 in duplicating cells, while the duration of cell-cycle transitions remain unchanged upon incubation with GSH.

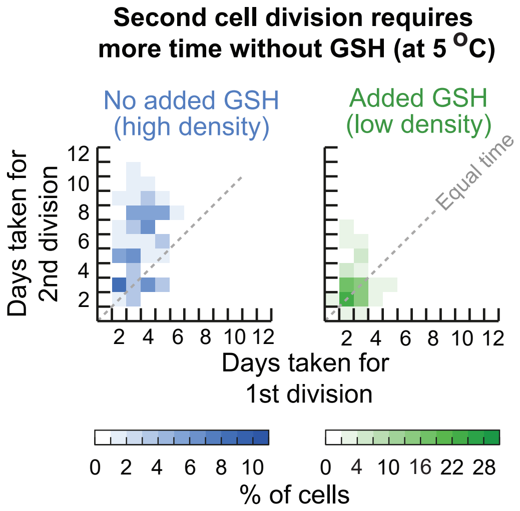


**Figure S23: Consecutive divisions in individual cells slow down at 5 °C (Related to Figure 4d).** We used the single-cell time-lapse movies at 5.0 °C to measure the doubling times of cells that divided multiple times. Heatmaps showing the relation of consecutive cell divisions of single cells at 5.0 °C without added GSH (blue heatmap, left) and with 250 μM added GSH (green heatmap, right). Shown is the number of days taken for the second cell division (y-axis) as function of the number of days taken for the first division (x-axis). Grey dotted diagonal indicates location where consecutive divisions in the same cell have equal doubling times. Data is aggregated from $n$ = 3 biological replicates. These measurements show that the doubling time increases over time in populations without added GSH: the time of a next division is on average larger (above the diagonal) than the time of a preceeding division. With added GSH, the doubling time remains approximately constant for consecutive divisions.

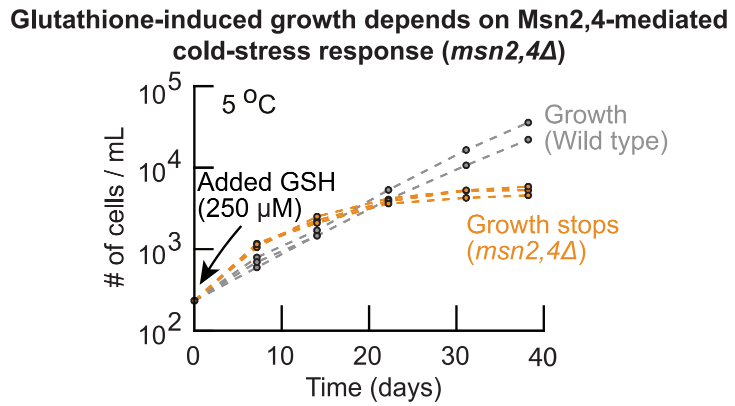


**Figure S24: Glutathione-induced growth at near-freezing temperatures depends on Msn2,4-mediate cold-stress response (Related to Figure 4e).** We constructed a mutant strain that lacks the major stress-response transcriptional activators, Msn2,4 (*msn2,4*Δ strain) [9, 10, 11]. Shown here is the population density over time for the *msn2,4*Δ strain at 5.0 °C with 250 μM added GSH (orange curves, initially ∼250 cells / mL). Grey curves show the wild-type strain for comparison (initially ∼250 cells / mL with 250 μM added GSH). The *msn2,4*Δ strain stops growing after two weeks of incubation with added GSH at 5.0 °C whereas the wild-type strain grows exponentially over time. We already established that GSH removes intracellular ROS and thereby induces the wild-type population to grow (Fig. 2c). However, the *msn2,4*Δ population stops growing at 5.0 °C even with the added GSH. This means that yeasts incubated with extracellular GSH must still experience a low-temperature (cold) stress, despite the added GSH greatly reducing the amount of intracellular ROS that inhibits cell duplications. Together, our results suggest that glutathione-induced population growths at 5.0 °C depend on the Msn2,4 cold-stress response. All colors show $n$ = 3 biological replicates.

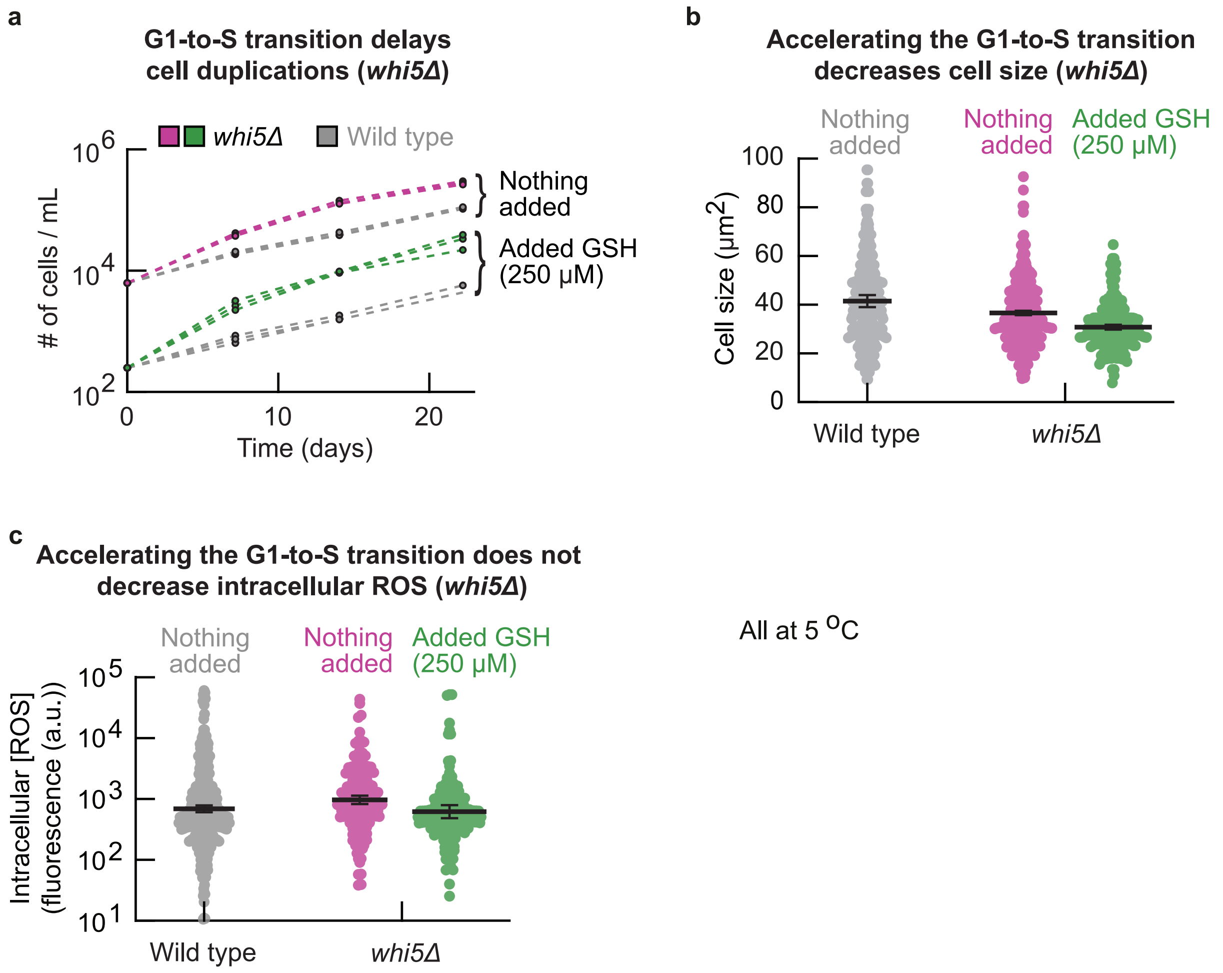


**Figure S25: G1-to-S transition delays cell duplications at 5 °C (Related to Figure 4e). (a)** We constructed a mutant strain (*whi5*Δ strain) whose *WHI5* was knocked out and therefore has an accelerated G1-to-S transition [12]. Shown here are the population densities over time for the *whi5*Δ strain that was incubated at 5.0 °C either without (pink curves, initially ∼6,250 cells / mL) or with 250 μM added GSH (green curves, initially ∼250 cells / mL). Both conditions are compared with the wild-type strain (grey curves). The *whi5*Δ strain grows faster at 5.0 °C (3.4 ± 0.1-fold difference in density after two weeks without added GSH, 5.7 ± 0.3-fold difference in density with added GSH). Thus, knocking out *WHI5* accelerates population growth, suggesting that the G1-to-S transition delays cell duplications. Each color and initial density shows $n$ = 3 replicate populations. **(b)** Cell size of the *whi5*Δ strain. Shown here are the cell sizes of the *whi5*Δ strain in populations without (pink dots) or with 250 added μM GSH (green dots). Wild-type strain without added GSH is shown for comparison (grey). Cell sizes were measured after two weeks of incubation at 5.0 °C. The *whi5*Δ strain has smaller cell size compared to the wild type, with adding extracellular GSH reducing the cell size further. This suggests that accelerating the G1-to-S transition decreases the cell size at 5.0 °C(cells spend less time in the G1 (growth) phase). ***(caption continues on the next page)***

**Figure S25** ***(caption continued from the previous page)***: Error bars show mean with s.e.m., having $n$ = 3 biological replicates. **(c)** Intracellular ROS concentration in the *whi5*Δ strain after two weeks of incubation at 5.0 °C (see Methods). Shown here are populations that were incubated either without (pink dots) or with 250 μM added GSH (green dots). Grey dots show wild-type without added GSH for comparison. The *whi5*Δ strain does not have less intracellular ROS compared to the wild-type strain. Error bars show mean with s.e.m., having $n$ = 3 biological replicates. Together, (a-c) show that accelerating the G1-to-S transition at 5.0 °C accelerates cell duplications and decreases cell size. Researchers found similar effects at 30.0 °C [12, 13]. Simultaneously, accelerating the G1-to-S transition does not decrease intracellular ROS abundance. This suggests that cells do not accumulate intracellular ROS during G1, since shortening G1 does not decrease the intracellular ROS concentration.

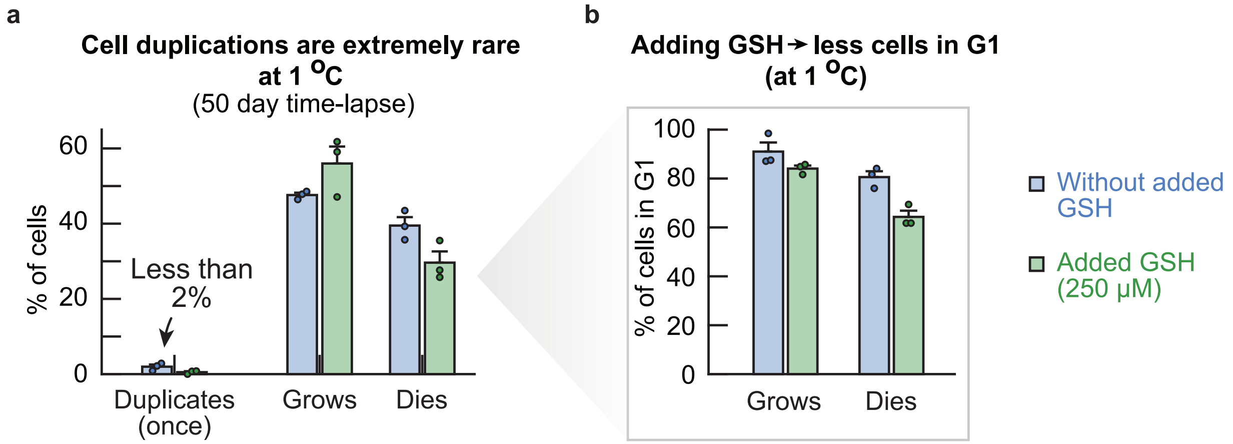


**Figure S26: Cells at 1 °C are arrested in G1 and less than 2% of cells proceeds to duplicate during a 51-day time-lapse (Related to Figure 5a).** **(a)** Percentage of cells that duplicates, grows and dies during a 51 day microscope time-lapse at 1.0 °C. Bars show populations with (blue bars) and without (green bars) 250 μM added GSH. Less than 2% of cells duplicated once (no cell duplicated more than once). **(b)** Percentage of cells that grows without duplicating or dies and that are in G1 for the duration of the time-lapse. Almost all cells that grow without duplicating or die are in G1. Similar to our measurements at 5.0 °C, we found that adding GSH decreases the percentage of cells that are in G1 (Supplementary Fig. S21).

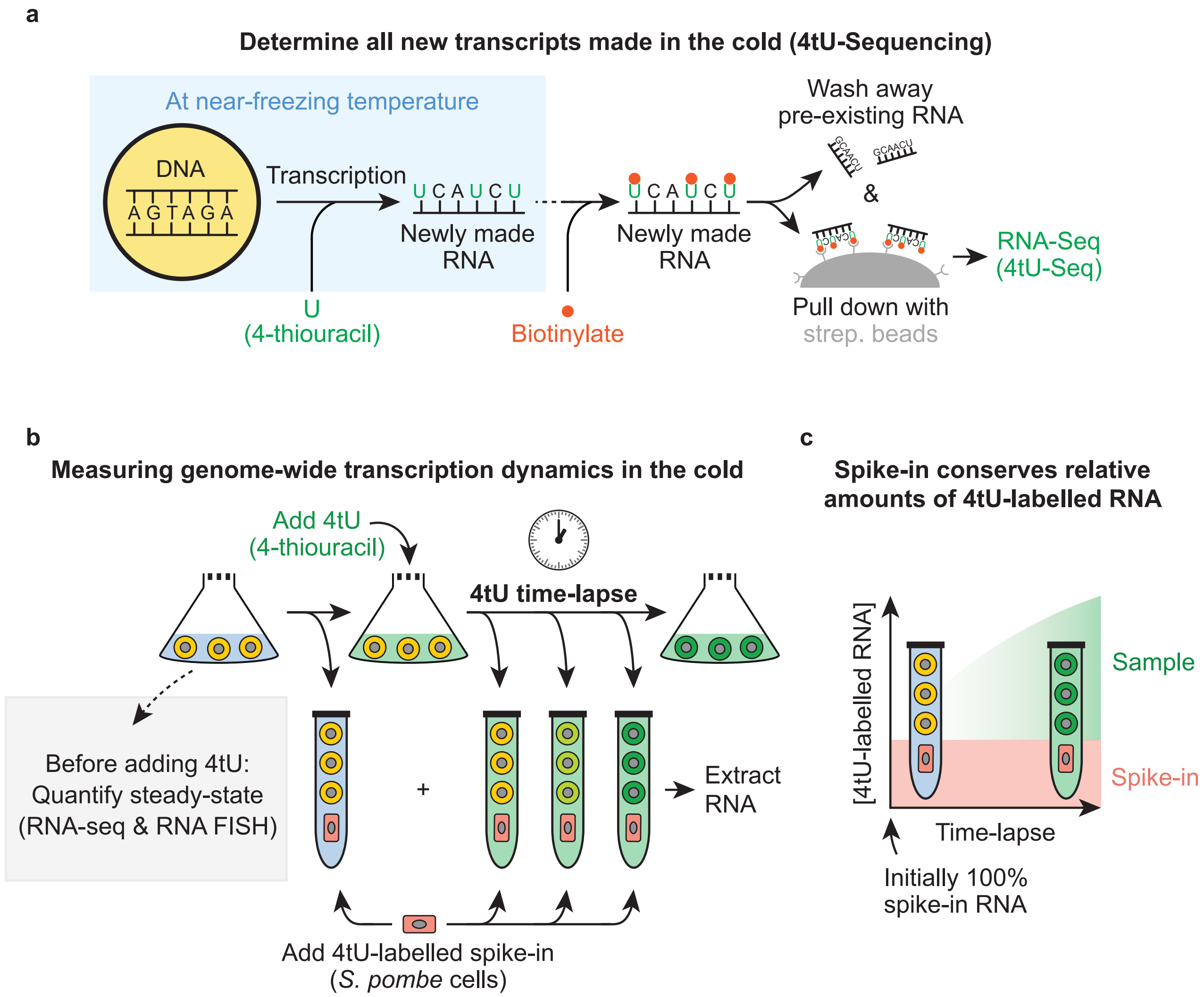


**Figure S27: Measuring the genome-wide transcription at near-freezing temperatures (Related to Figure 5b). (a)** Schematic demonstrating the experimental procedure for performing 4tU-sequencing on newly-made transcripts in populations of *S. cerevisiae* [14, 15, 16]. A synthetic uracil analog (the nucleotide 4-thiouracil, 4tU) is added to the growth medium of cells. Cells incorporate the 4tU into their newly synthesized RNA. We collect cells after the desired time of incubation with 4tU and extract the total RNA. The newly synthesized RNA is linked to biotin by specifically biotinylating the 4tU ("4tU-labelled" RNA). Simultaneously, the pre-existing RNA that does not contain 4tU is not biotinylated. Finally, the 4tU-labelled RNA is separated from the pre-existing RNA without 4tU through a pull-down with magnetic beads containing streptavidin, after which the purified 4tU-labelled RNA is sequenced (see Methods for experimental details). **(b)** Schematic illustrating the experimental procedure for measuring the genome-wide transcription dynamics at near-freezing temperatures. We incubated liquid cultures with populations of wild-type yeasts at the desired temperature (e.g., 5.0 °C). After two weeks of incubation, we first collected two aliquots of our cultures (as "time 0" hours of the time-lapse). ***(caption continues on the next page)***

**Figure S27** ***(caption continued from the previous page)*****:** Directly after, we supplemented the growth medum of the cultures with 4tU at a final 5 mM concentration, and collected aliquots from our cultures at the desired time-points (see Methods for experimental details). After collecting the time-lapse samples, we added a fixed amount of 4*tU*-labelled *S. pombe* (*Schizosaccharomyces pombe*) cells to all 4tU time-lapse samples as a spike-in and proceeded with RNA-extraction. Finally, we used one of the ”time 0” hours aliquots to quantify the steady-state transcript levels (grey box, RNA-seq and RNA FISH). **(c)** The *S. pombe* spike-in that was added to all 4tU time-lapse samples enabled us to normalize the amount of 4tU-labelled transcripts from different time-points. Initially, all 4tU-labelled RNA in the sample is from the spike-in since the cold-incubated cells have not initially synthesized 4tU-labelled RNA. Over time, the cold-incubated cells synthesize new 4tU-labelled RNA and degrade old (unlabelled) RNA, increasing the percentage of 4tU-labelled RNA that is from the cold-incubated cells. Normalizing with the amount of spike-in transcripts therefore ensures that the relative differences between samples in a time-lapse are conserved (since the amount of 4tU-labelled RNA from *S. pombe* cell was the same in every sample). Moreover, this procedure eliminates experimental differences between samples due to sequencing and processing of RNA.

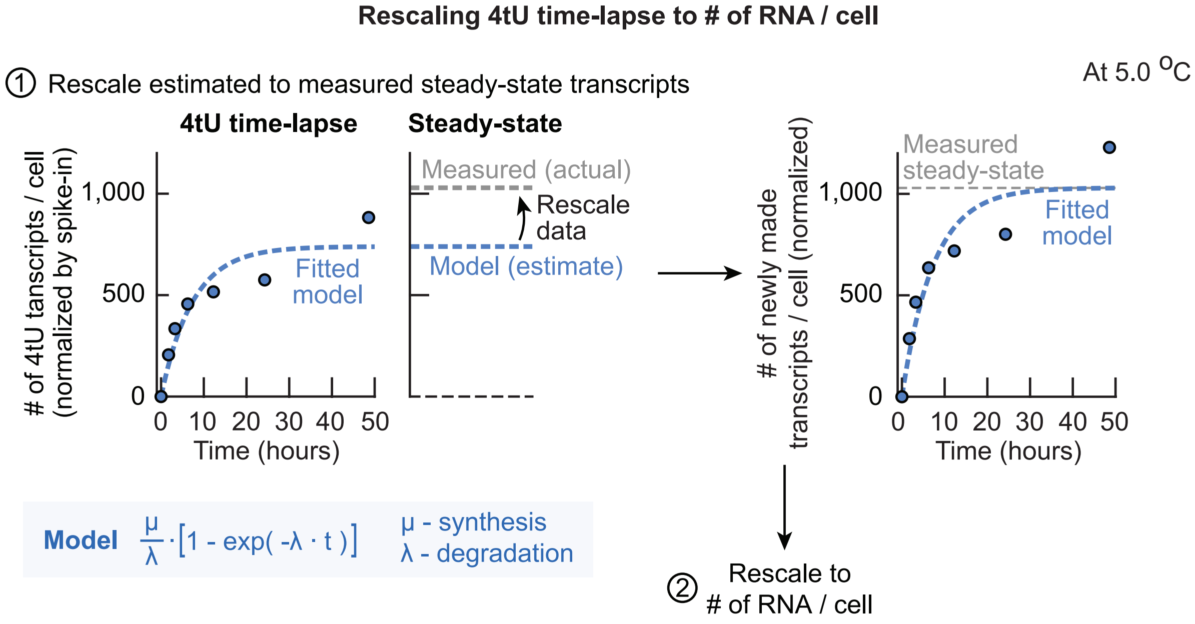


**Figure S28: Procedure for rescaling the 4tU time-lapses to # of RNA / cell using steady-state transcript levels (Related to Figure 5b).** First, the 4tU time-lapse samples were normalized by the number of spike-in transcripts and the number of alive cells in the population, giving the relative amount of 4tU transcripts per cell over time (amount of 4tU-labelled RNA normalized by 4tU spike-in). Next, these relative values are rescaled to obtain the "# of RNA / cell". To do so, the samples are rescaled such that steady-state amount of RNA / cell that is predicted by the 4tU time-lapse is equal to the actual steady-state amount of RNA / cell that we measured through RNA-seq. First, we fitted a mathematical model to the amount of 4tU transcripts / cell. The conventional model to describe RNA synthesis assumes that RNA is synthesized at a constant rate and degraded at a rate that is dependent on the concentration. Thus, the kinetics of 4tU-labelled RNA are given by $\frac{dN}{dt} = \frac{\mu}{\lambda}\left(1 - \exp(-\lambda t)\right)$ with a RNA synthesis rate $\mu$ (in "# of RNA / cell / hour"), a RNA degradation rate $\lambda$ (in "per hour") and time $t$. Over time the relative amount of 4tU transcripts per cell converges to a steady-state $\mu/\lambda$ (in "cell / hour") as $t \to \infty$ by degradation of old RNA and synthesis of new (4tU-labelled) RNA [17, 16]. To rescale the steady-state amount of RNA / cell that is predicted by model to the actual steady-state amount of RNA / cell, we had also sequenced the steady-state transcript levels (total RNA, not 4tU purified RNA) of each time-lapse together with a *S. pombe* spike-in (total RNA) to normalize our time-lapses across temperatures (the steady-state transcript levels can differ per temperature, and the same spike-in RNA was used for all steady-state samples). Thus, we rescaled all 4tU-samples such that the predicted steady-state of the model matched the measured steady-state transcript levels. Shown is a representative example of the rescaling from a 4tU time-lapse without added extracellular GSH at 5.0 °C. Note that this rescaling conserves the relative differences between time points within each time-lapse. ***(caption continues on the next page)***

**Figure S28** ***(caption continued from the previous page)*****:** Together, the previous normalizations resulted in the relative number of newly-made transcripts (4tU labelled RNA) over time between replicates and across temperatures, and that the predicted steady-state of these time-lapses is equal to the measured steady-state of each time-lapse. As a last step, the relative number of newly-made transcripts / cell are rescaled to the actual integer number of RNA per cell. Dots show raw (normalized) data for one replicate. Blue dotted lines shows model fit for that replicate. Grey dotted lines shows measured steady state values.

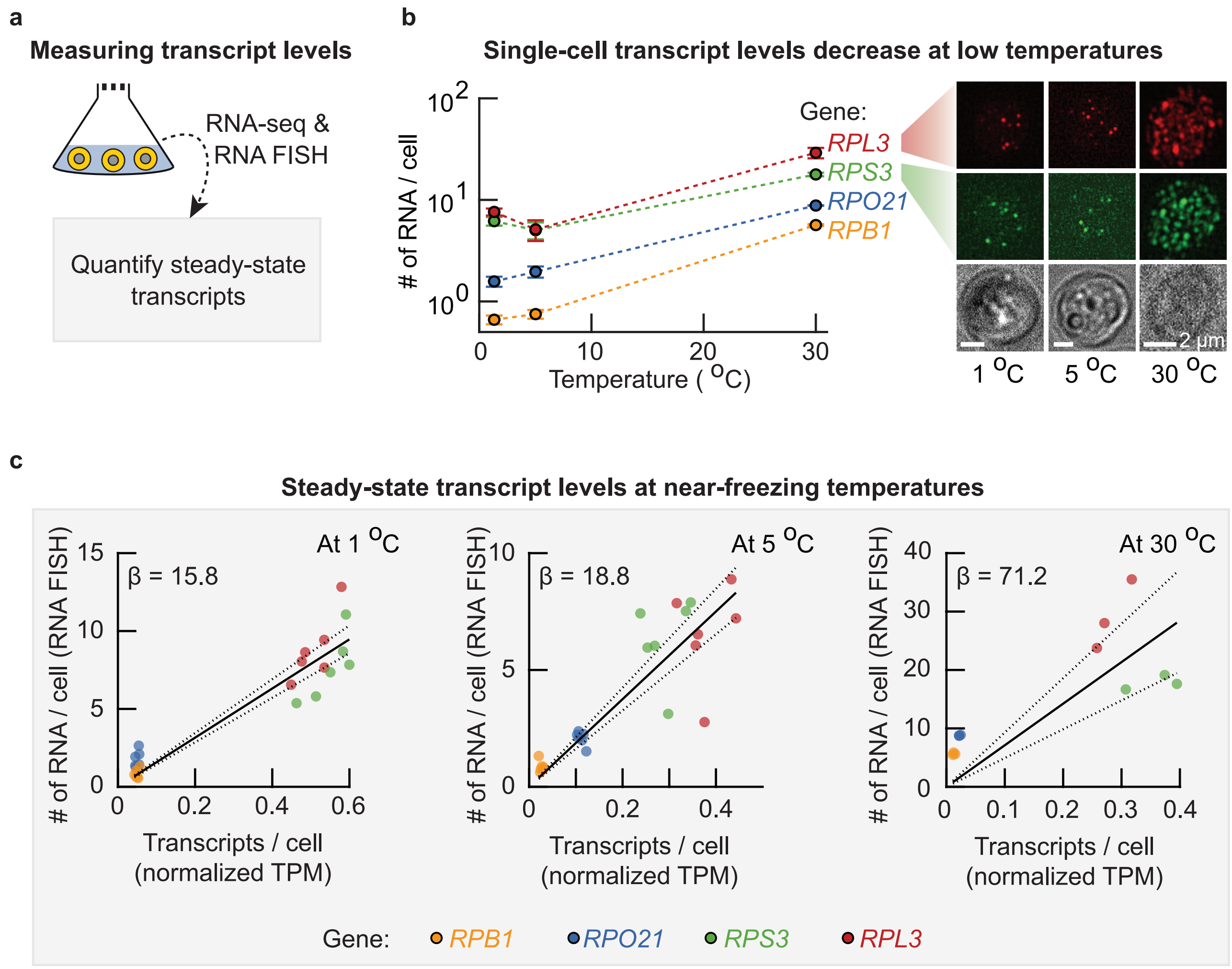


**Figure S29: Steady-state transcript levels decrease with temperature (Related to Figure 5b). (a)** Steady-state transcript levels were quantified in two ways. In one way, we performed regular RNA-seq on the steady-state RNA (normalized across temperatures by a spike-in of total RNA from *S. pombe*). In the another way, we quantified the number of mRNA per cell via single-molecule RNA FISH on cells from the same aliquots on which we performed RNA-seq. These measurements reveal the steady-state transcript levels across temperatures, and can be combined to rescale the number of transcripts / cell (from RNA-seq) to the "# of RNA / cell" as measured by RNA FISH. **(b)** The single-cell transcript levels of several endogenous yeast genes were quantified at 30 °C, 5.0 °Cand 1.0 °C. Shown is the average number of mRNA per cell as function of temperature for *RPL3* (red curve), *RPS3* (green curve), *RPO21* (blue curve) and *RPB1* (yellow curve curve). The average number of mRNA per cell decreases with temperature. Each data point shows the average of $n = 3$ biological replicates, in total having counted at least $n = 616$ cells (for *RPB1* and *RPO21*) or at least $n = 421$ cells (for *RPL3* and *RPS3*). Images show representative examples of labelled RNA in single-cells at 1.0 °C (left), 5.0 °C (middle) and 30 °C (right). ***(caption continues on the next page)***

**Figure S29** ***(caption continued from the previous page)***: Dots are *RPL3* RNA (red fluorescence, top) and *RPS3* RNA (green fluorescence, bottom). Brightfield images are shown as comparison. Scale bar is 2 $\mu$m. **(c)** The average number of RNA per cell (from RNA FISH) as function of the relative steady-state amount of transcripts per cell (from RNA-seq, normalized with the *S. pombe* RNA spike-in) at 1.0 °C (left panel), 5.0 °C (middle panel) and 30 °C (right panel). Shown are the steady-state levels for *RPB1* (yellow dots), *RPO21* (blue dots), *RPS3* (green dots) and *RPL3* (red dots). Black solid line shows linear fit without intercept, dotted lines show 95% confidence interval of fit. Since both RNA FISH and RNA-seq measurements come from the same sample, these linear fits yield a scaling factor to convert the relative amount of transcripts per cell to "# of RNA / cell" ($\beta$ = 15.8 at 1.0 °C (pearson-correlation coefficient $\rho$ = 0.94), $\beta$ = 18.8 at 5.0 °C ($\rho$ = 0.87) and $\beta$ = 71.2 at 30 °C ($\rho$ = 0.79)). All colors show 6 dots, having $n$ = 3 biological replicates for with and without added GSH added.

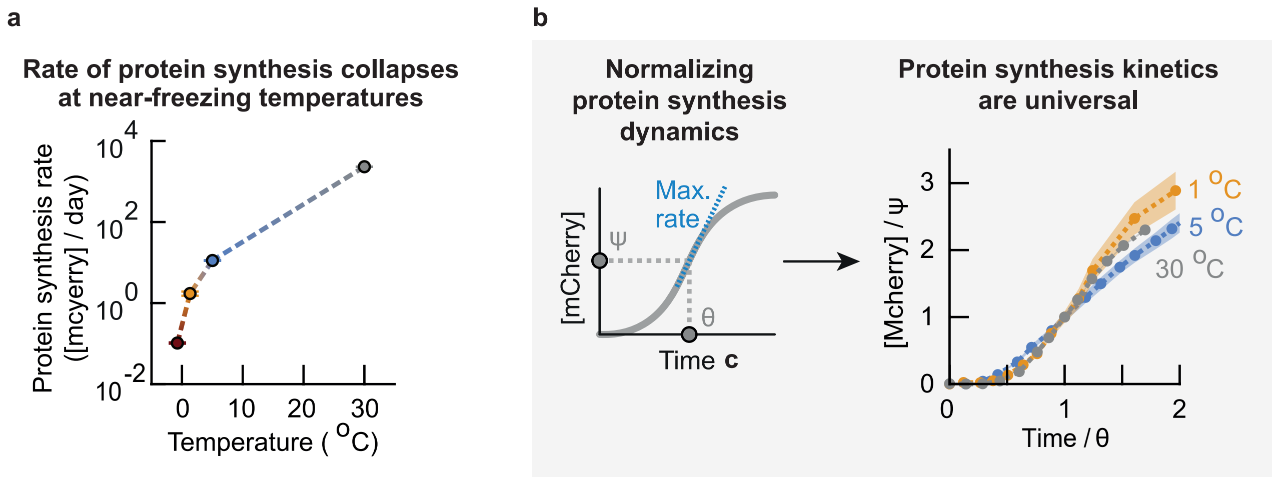


**Figure S30: Protein synthesis dynamics are universal with the protein synthesis rate collapsing around 0 °C (Related to Figure 5c).** We build a synthetic gene-circuit in our wild-type yeast that makes the cells produce a red fluorescent protein (mCherry) upon induction by galactose. Expression of mCherry is controlled by an inducible promoter, *pGAL10*, which is activated by galactose. We first grow the yeast in raffinose and then add galactose to the raffinose-medium to activate the expression of mCherry. **(a)** For each curve in Fig. 5c we determined the maximum rate of mCherry expression (steepest slope in each curve, also see Supplementary Theory). Shown are the protein synthesis rates as function of temperature (the synthesis rate of general proteins can be at least as fast as the synthesis rate of mCherry measured here). **(b)** Protein synthesis dynamics across temperatures. *Left*: Schematics of relevant parameters. At each temperature, we used the characteristic time $\Theta$ and fluorescence $\Psi$ at which the population reaches its maximum rate of protein synthesis to rescale the fluorescence curves. *Right*: Renormalized protein synthesis curves from Fig. 5c. The fluorescence was normalized with the characteristic fluorescence $\Psi$ and time was rescaled with the characteristic time $\Theta$ for each temperature. The renormalized curves collapses onto a single master curve. These renormalized curves suggest that one only needs to know the characteristic fluorescence-scale $\Psi$ and time-scale $\Theta$ for each temperature to fully describe expression of mCherry at that temperature. In both panels the error bars and shaded areas show the mean with s.e.m., having $n = 3$ replicate populations.

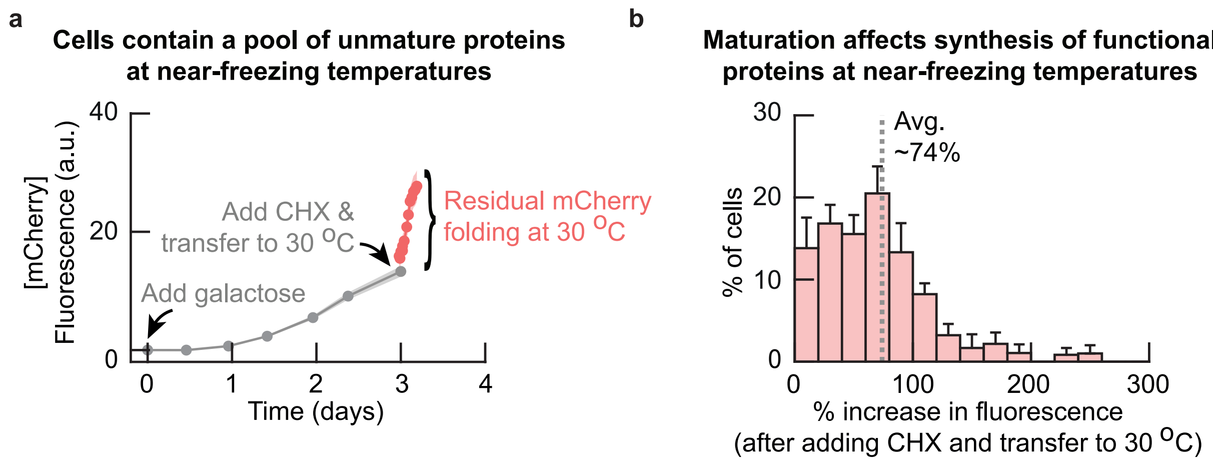


**Figure S31: Time to synthesize functional protein affected by maturation (e.g., folding) time at 5 °C (Related to Figure 5c).** Using the mCherry-inducible strain to quantify processes affecting time to synthesize functinoal proteins at near-freezing temperatures (also see Supplementary Fig. S30). **(a)** We incubated cells of the inducible mCherry strain for two weeks at 5.0 °C. We then added galactose to the growth media and followed the expression of mcherry at 5 °C in single cells over time with a microscope (see Methods). After three days of incubation with galactose, we added cycloheximide (CHX) to the growth medium at a final 100 μg/mL concentration to instantly stop translation in the cells. We then transferred the cells to 30 °C and continued to measure the mCherry fluorescence of the cells to quantify the maturation (e.g., folding) of residual mCherry. Shown is the fluorescence of mCherry over time at 5 °C (grey curve), and the increase in fluorescence at 30 °C after addition of CHX (red curve). The average fluorescence considerably increases after addition of CHX. Cells therefore contain a pool of unmatured mCherry at 5 °C. Dots show average fluorescence, and shaded area shows s.e.m. of fluorescence from the averages in $n = 5$ populations. **(b)** Histogram shows the increase in fluorescence of single cells from (a) after addition of CHX and transfer to 30 °C. The average fluorescence increases by 74% after addition of CHX, showing that roughly half of mCherry remains unmatured during gene expression. Synthesis of functional proteins is therefore affected by the time required for maturation (e.g., folding) at near-freezing temperatures (also see Supplementary Theory). Histogram contains $n = 108$ cells, error bar shows s.e.m. of $n = 5$ populations.

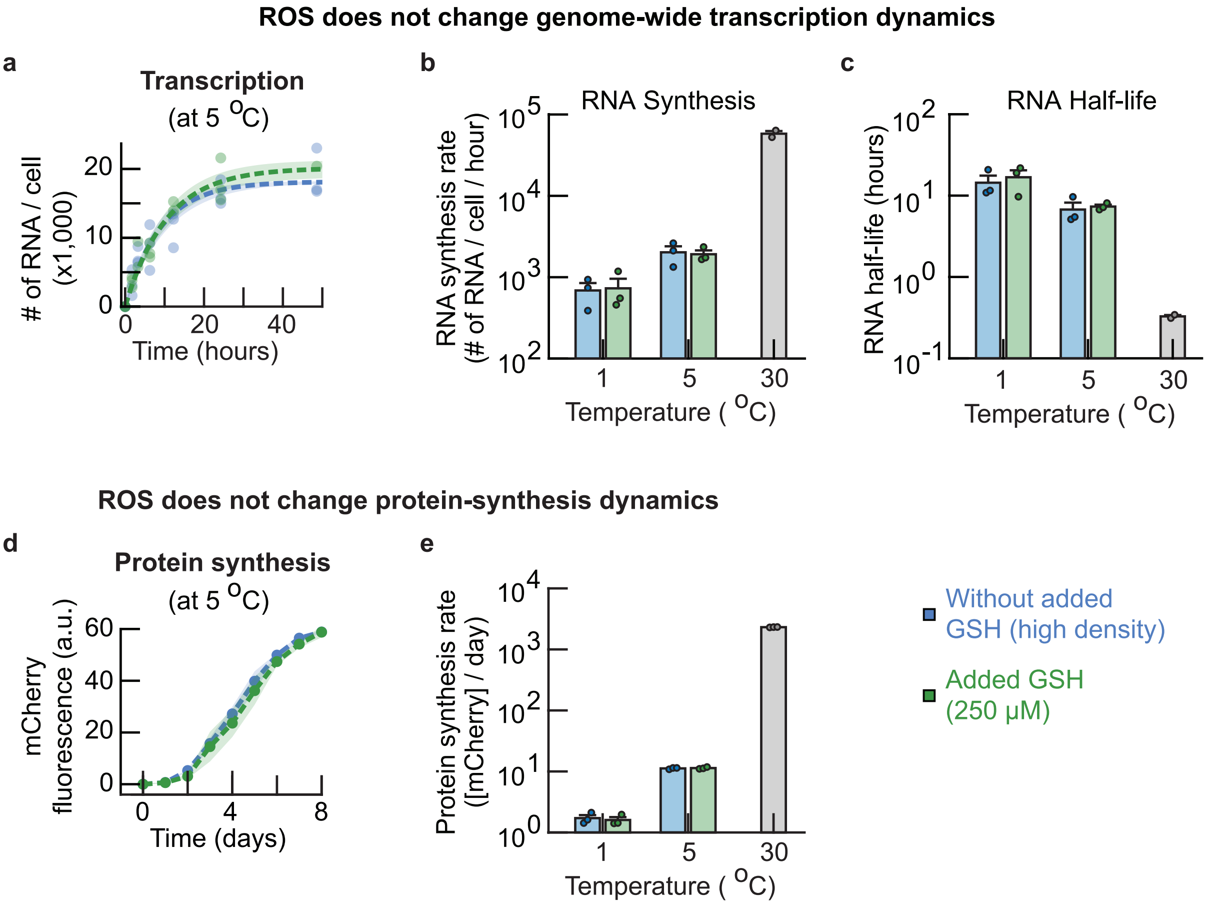


**Figure S32: ROS does not change genome-wide transcription or protein synthesis dynamics at near-freezing temperatures (Related to Figure 5d). (a-c)** ROS does not change rates of genome-wide transcription. **(a)** We used the number of synthesized RNA over time to fit a model for the kinetics of RNA synthesis (also see Supplementary Fig. S28). Shown are the number of newly synthesized RNA per cell over time at 5.0 °C for populations with (blue curve) or without (green curve) 250 μM added GSH. Dots show raw measurements. Dotted line represents kinetics model using average parameter estimates, with n = 3 biological replicates per condition. Shaded area represents the s.e.m. for fitted model parameters. **(b-c)** RNA synthesis rates **(b)** and RNA half-lives **(c)** at 1.0 °C, 5.0 °C as quantified from the fitted model. Included are the rates for populations without (blue bars) and with 250 μM added GSH (green bars). Rates at 30 °C are shown as a comparison (grey bars). The RNA half-lives and RNA synthesis rates decrease with temperature. Without added GSH, the steady-state transcript levels (in # of RNA / cell) are $12,900 \pm 900$ (1.0 °C), $18,300 \pm 800$ (5.0 °C) and $27,300 \pm 1,300$ (30 °C), the synthesis-rates (in # of RNA / cell / hour) are $690 \pm 160$ (1.0 °C), $2020 \pm 370$ (5.0 °C) and $58,100 \pm 5,500$ (30 °C), and the half-lives are (in hours) $14.4 \pm 3.9$ (1.0 °C), $6.7 \pm 1.8$ (5.0 °C) and $20 \pm 1$ (30 °C, in minutes). There is no reason to assume that the RNA synthesis rates are different upon addition of GSH (p-value is 0.82 at 5.0 °C and 0.89 at 1.0 °C). ***(caption continues on the next page)***

**Figure S32** ***(caption continued from the previous page)***: Error bars show mean with s.e.m., having $n$ = 3 biological replicates per condition at 1.0 °C and 5.0 °C, and $n$ = 2 biological replicates at 30 °C. Dots show raw data. **(d-e)** ROS does not change rate of protein synthesis. **(d)** The synthesis of mCherry over time at 5.0 °C compared for populations without (blue curve) and with (green curve) added 250 μM GSH (also see Supplementary Fig. S30). The dots show average fluorescence, with shaded representing the s.e.m. of n = 3 biological replicates. **(e)** Protein synthesis rates at 5.0 °C and 1.0 °C for populations that were incubated without (blue bars) or with (green bars) 250 μM added GSH. Grey bar shows synthesis rate of mCherry at 30 °C as comparison. The protein synthesis rate decreases with temperature. Without added GSH, the protein synthesis rates (in mCherry fluorescence / day (a.u.)) are 1.7 $\pm$ 0.2 (1.0 °C), 11.2 $\pm$ 0.2 (5.0 °C) and 2314 $\pm$ 10 (30 °C). There is no reason to assume that the protein synthesis rates are different upon addition of GSH (p-value is 0.66 at 5.0 °C and 0.69 at 1.0 °C).

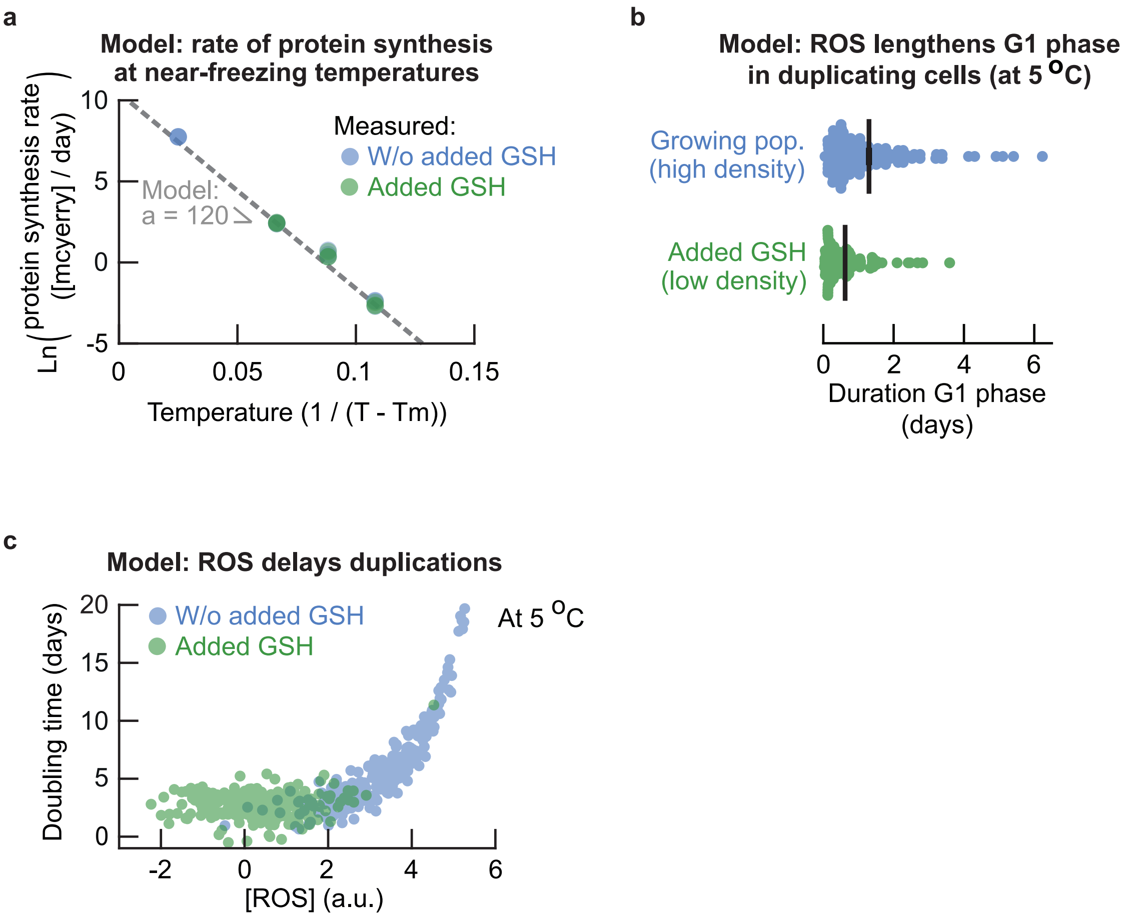


**Figure S33: Stochastic model for single-cell duplications at near-freezing temperatures (Related to Figure 6). (a)** Fitting the protein synthesis rate $r_g(T)$ as function of temperature $T$ using an Arrhenius-type equation ($\ln r_g(T) \propto -a/(T - T_m)$, see equation S11 and Supplementary Theory for derivation). This model assumes a minimum temperature $T_m = -10$ °C below which protein synthesis does not occur. Fitting a linear model yields the parameter $a \approx 120.72$ that fully describes the protein synthesis rate as function of temperature. **(b)** Model simulation of ROS-dependent G1 duration in single cells at 5.0 °C (equal to the delay in the model, compare with Fig. 4c). The only free model parameter (intracellular ROS concentration) was chosen such that the average delay equals the average measured G1 duration for each condition respectively (Fig. 4c, see Supplementary Theory, parameter $R$ in equation S16). This yields $R = 2.4$ for high-density (growing) populations and $R = 1.55$ for low-density populations with added GSH. Error bar shows mean with s.e.m. of $n = 3$ replicate simulations that each have $n = 50$ cells. Dots show raw single-cell delays. **(c)** Model simulation of ROS-dependent doubling times in single cells at 5.0 °C without (blue dots) and with added GSH (green dots). Initially, for relatively low ROS concentrations, the ROS-dependent duration is negligible compared to the minimum doubling time that is set by the protein synthesis rate. ***(caption continues on the next page)***

$$\frac{\partial A}{\partial t} = A(t) \cdot (\mu - \lambda). \tag{S1}$$

Similarly, the total number of cells changes according to,

$$\frac{\partial N}{\partial t} = \mu A(t), \tag{S2}$$

since we add freshly born cells to the total number of cells and dead cells are not removed from the total number of cells in the population. Solving the differential equation S1 yields,

$$A(t+dt) = A(t) \cdot \exp\Big((\mu - \lambda) \cdot dt\Big). \tag{S3}$$

Using equation S3 to solve equation S2 we obtain,

$$\begin{aligned} N(t+dt) &= N(t) + \int_0^{dt} \mu A(\tau) d\tau \\ &= N(t) + \mu A(t) \int_0^{dt} \exp\Big((\mu - \lambda)\tau\Big) \\ &= N(t) + \frac{\mu}{\mu - \lambda} A(t) \Big[\exp\Big((\mu - \lambda)\tau\Big)\Big]_0^{dt} \\ &= N(t) + \frac{\mu}{\mu - \lambda}\Big(A(t+dt) - A(t)\Big) \end{aligned} \tag{S4}$$

In our experiments we can measure the number of alive cells and the number of dead cells and the total number of cells over time. We therefore have $A(t)$ and $N(t)$ available to fit the growth rate $\mu$ and death rate $\lambda$ with our model. Taking the logarithm of equation S3 describing the number of alive cells in the population and rewriting yields,

$$r_1 := \frac{1}{dt}\Big(\ln A(t+dt) - \ln A(t)\Big) = \mu - \lambda \tag{S5}$$

Similarly, we can rewrite equation S4 describing the total number of cells in the population as,

$$r_2 := \frac{N(t+dt) - N(t)}{A(t+dt) - A(t)} = \frac{\mu}{\mu - \lambda}. \tag{S6}$$

Finally, we use the numbers $r_1$ (from equation S5) and $_r2$ (from equation S6) to find the rates $\mu$ and $\lambda$,

$$\begin{aligned} \mu &= r_1 \cdot r_2, \\ \lambda &= r_1 \cdot (r_2 - 1). \end{aligned}$$

We used the above equations together with equations S5 and S6 to estimate the growth rate $\mu$ and death rate $\lambda$ from the total number of cells and the number of alive cells in a population.

**Model for protein synthesis rate (Related to Fig. 5).** Here we describe a simple kinetics model for the synthesis of mCherry that we measured experimentally by following the mCherry fluorescence in cells over time. Assume that mCherry is synthesized by first being translated into an unfolded polypeptide chain, followed by folding into a fluorescent protein (Supplementary Fig. S31). We let $u$ denote the amount of unfolded mCherry and $f$ denote the amount of folded mCherry in a cell. We further assume that unfolded mCherry is synthesized through translation at a constant rate $m$ (a.u. / day), and that the unfolded mCherry is folded at a rate $\gamma$ (day$^{-1}$). Then the amounts of unfolded and folded mCherry over time $t$ are described by,

$$\frac{du}{dt} = m - \gamma \cdot u, \tag{S7}$$

$$\frac{df}{dt} = \gamma \cdot u. \tag{S8}$$

Solving equation S7 yields $u(t) = \frac{m}{\gamma} \cdot \Big(1 - \exp(-\gamma \cdot t)\Big)$, and substitution of $u(t)$ into equation S8 yields,

$$f(t) = \frac{m}{\gamma} \cdot \Big(\exp(-\gamma \cdot t) - 1\Big) + m \cdot t. \tag{S9}$$

Equation S9 describes the amount of folded mCherry over time and provides a model for the experimentally measured mCherry fluorescence in cells. Initially translation of mCherry has not occured, such that a cell does not have unfolded mCherry and no folded mCherry is being produced. At time $t = 0$ the cell starts synthesizing unfolded mCherry, whose concentration is described by $u(t) = \frac{m}{\gamma} \cdot (1 - \exp(-\gamma \cdot t))$. Thus, the amount of unfolded mCherry initially increases by translation and eventually reaches a steady-state that is determined by an inflow from translation and an outflow to folded mCherry ($u(t) \rightarrow \frac{m}{\gamma}$ as $t \rightarrow \infty$). At this steady-state, the equation S7 for unfolded mCherry satisfies $\frac{du}{dt} = 0$, such that $\gamma \cdot u = m$ and $\frac{df}{dt} = m$ (from equation S8). Thus, this derivation shows that the slope of mCherry fluorescence over time converges to the translation rate $m$ ($\frac{df}{dt} \approx m$ for sufficiently large $t$). In practice, we measured the synthesis rate $m(T)$ for each temperature $T$ by taking the maximum slope of the increase of mCherry fluorescence over time, and then used these $m(T)$ to further fit the model in equations S7 to the data.

$$\tau(T,R) = t(T) + \Delta t(R,T).$$

where $R$ is some environmental parameter that depends on, for example, ROS and extracellular GSH. Here $\Delta t > 0$ such that the doubling time is at least the minimum time for a given temperature ($tau(T,R) > t(T)$).

**Modeling temperature dependence of the protein synthesis rate** Recall that the protein synthesis rate decreases as function of temperature (Supplementary Fig. S30). To describe the protein synthesis as function of temperature with a model we assume a lower temperature limit $T_m$ below which no protein synthesis is possible. Then we can describe a rate as function of temperature with a common Arrhenius-type model $r_g(T) = c \cdot \exp(-a/(T - T_m))$ for some parameter $a$ and minimum temperature $T_m$, such that $r_g(T) \to 0$ as the temperature decreases to $T_m$ (the Arrhenius equation would have $T_m = -273$ °C). We can rewrite this model using the experimentally measured protein synthesis rate at a known temperature $T_0$,

$$r_g(T) = r_g(T_0) \cdot \exp\Big[ -a\Big(\frac{1}{T - T_m} - \frac{1}{T_0 - T_m}\Big)\Big], \tag{S10}$$

providing a simple equation that describes the protein synthesis rate as function of temperature. Taking the logarithm of equation S10 we find that,

$$\ln r_g(T) \propto -a \cdot \frac{1}{T - T_m}, \tag{S11}$$

which is a model that is linear in $1/(T - T_m)$ and that we fit to the measured protein synthesis rates at various temperatures to find the slope $a$.

**Model of minimum time limited by the protein synthesis rate** We have experimentally measured both the minimum time $t(T)$ by taking away ROS with GSH (Supplementary Fig. S11). We have also measured the ROS-independent protein synthesis rates (Fig. 5d, Supplementary Fig. S32). Then both the minimum time $t(T)$ and protein synthesis rate are independent of ROS, and we assume without loss of generality that the minimum time $t(T)$ is limited by the protein synthesis rate $r_g(T)$ at each temperature $T$ (i.e., cell division requires protein synthesis, so that the timescale of cell division must be longer than the timescale of protein synthesis). Here, we seek to describe the relation between the minimum doubling time and the protein synthesis rate.

Experimentally we have found that the observed doubling time slows down with the protein synthesis rate via a power-law type scaling (Fig. 6b). One reason why the relation may be of power-law type is the following. We assume that the rate of each process depends on the temperature according to an Arrhenius-type equation $r_p(T) = A_p \exp\left(-\frac{E_p}{RT}\right)$, where $r_p(T)$ is the rate of the process $p$ (protein synthesis or cell division) dependent on and constants $E_p$, $R$ and $A_p$. Isolating the reciprocal temperature gives $\frac{1}{T} = -\frac{R}{E_p}\Big(\ln(r_p) - \ln(A_p)\Big)$. Denoting the rate of cell division as $r_d(T)$ and the rate of protein synthesis with $r_g(T)$, we can express the rate of cell division in terms of the rate of protein synthesis (temperature dependence of the rates omitted for readability),

$$\begin{aligned}\ln(r_d) &= -\frac{E_d}{R}\Big(\frac{1}{T}\Big) + \ln(A_d) \\ &= -\frac{E_d}{R}\Big( - \frac{R}{E_g}(\ln(r_g) - \ln(A_g))\Big) + \ln(A_d) \\ &= \frac{E_d}{E_g}\ln(r_g) + c_1, \end{aligned} \tag{S12}$$

with $c_1$ a constant of aggregated rest terms. Let $k = E_d/E_g$ be some factor such that by taking the exponent of equation S12 we obtain, $r_d = c_2 \cdot r_g^k$. Then, by substituting the minimum time $t(T)$ for the growth rate via $t(T) = \ln(2)/r_d(T)$ we obtain,

$$t(T) = c \cdot r_g(T)^{-k}, \tag{S13}$$

for some constant $c$ and exponent $k$. Note that the above derivation is not restricted to the choice for a temperature scale. Taking the logarithm of equation S13 we find that,

$$\ln t(T) \propto -k \ln r_g(T),$$

which is a simple linear model that we fit to the measured minimum times and protein synthesis rates at various temperatures to find the value of the exponent $k$. The power law with exponent $k$ in equation S13 also illustrates how the doubling time scales with protein synthesis. For example, if the protein synthesis rate changes by two-fold, then the doubling time changes by $2^{-k}$-fold, such that fold-change in the doubling time is smaller than the fold-change in protein synthesis for exponents $k < 1$. Of note, it is only the fold-change of the protein synthesis rate that sets how the doubling time changes, irrespective of their actual values (the doubling time and protein synthesis rate change proportionally).

In summary, with an Arrhenius-type equation describing rates as function of temperature we can derive that cell division and protein synthesis may follow a power-law relation. We have therefore now experimentally measured and theoretically motivated a power-law type scaling of the minimum time $t(T)$ and the protein synthesis rate. Finally, using equation S13, we rewrite the minimum time $t(T)$ using a minimum observed time $t(T_0)$ at a known temperature $T_0$,

$$t(T) = t(T_0) \cdot \Big(\frac{r_g(T)}{r_g(T_0)}\Big)^{-k}.$$

We know from our single-cell data that the experimentally measured minimum doubling time is stochastic and approximately normally distributed (e.g., the single-cell doubling times of the wild type at 5.0 °C with added GSH, see Supplementary Fig. S11). We therefore assume that $t(T_0)$ is stochastic and equal to the experimentally measured minimum doubling time $\tau(T_0)$ plus some normally distributed noise

$\epsilon_m \sim N(0, 1)$, such that,

$$t(T) \sim \Big(\tau(T_0) + \epsilon_m\Big) \cdot \Big(\frac{r_g(T)}{r_g(T_0)}\Big)^{-k}, \qquad \epsilon_m \sim N(0, 1). \tag{S14}$$

We then use the $t(T)$ for $\epsilon_m = 0$ as the deterministic minimum doubling time $t_{\min}(T)$ from the model. The minimum doubling time is thereby the same for all cells in a population at a given temeperature, and only changes proportionally to the protein synthesis rate as temperature changes. Cell-to-cell variability due to other factors is simulated with the normally distributed noise.

**Stochastic model of the ROS-dependent duration of the doubling time** Given the ROS-independent description of the minimum doubling time (equation S14), we here seek to describe the ROS-dependent additional time $\Delta t(R, T)$ that increases the doubling time above this minimum. To describe this ROS-dependent additional time, we assume that $\Delta t(R, T)$ is described by the time it takes a cell to remove sufficient intracellular ROS and repair enough damages to divide. For example, we have experimentally measured that cells with low intracellular ROS concentrations are likely to duplicate whereas cells with high intracellular ROS concentrations are unlikely to duplicate (Fig. 3b). We can therefore assume that a cell has some amount of doubling time delaying tasks to perform before it can duplicate and that this amount of time is proportional to the abundance of ROS. We further assume that these tasks can be performed in a time that is proportional to how fast a cell can express genes, for example to synthesize ROS-reducing enzymes and replacing all damaged components with a freshly synthesized component. Thus, we assume that $\Delta t(R, T) \propto [\text{ROS}]/r_g(T)$ (a cell synthesizes proteins at a rate $r_g(T)$ (a.u. / time) and an amount of [ROS] (a.u.) tasks (repair/protein synthesis) must be completed). Moreover, we have experimentally measured the intracellular ROS concentrations in cells, which is approximately normally distributed on a log-scale (see Supplementary Fig. S9). Finally, our measurements suggest that there is some threshold concentration of ROS above which duplications become very unlikely (Fig. 3b). We therefore assume that we can describe the distribution of intracellular ROS concentrations according to $\ln[\text{ROS}] \sim N(R, 1)$, where $R$ is some parameter describing environmental factors (see Fig. 2a,c). Thus, $\Delta t(R, T)$ is stochastic and described by,

$$\Delta t(R, T) \sim \frac{1}{r_g(T)} \cdot \exp\Big(\epsilon_d(R)\Big), \qquad \epsilon_d(R) \sim N(R, 1). \tag{S15}$$

The time duration that increases the doubling time above its minimum thus varies between cells because of some ROS-dependent parameter that is log-normally distributed.

**Stochastic model for the doubling time dictated by temperature and ROS** Using the ROS-independent minimum doubling time (equation S14) and the ROS-dependent duration of the doubling time (equation S15), we obtain te doubling time as function of temperature $T$ and environmental parameter $R$,

$$\begin{aligned}\tau(T,R) &= t_{\min}(T) + \Delta t(R,T)\\ &= \Big(\tau(T_0) + \epsilon_m\Big)\cdot\Big(\frac{r_g(T)}{r_g(T_0)}\Big)^{-k} + \frac{1}{r_g(T)}\cdot\exp\Big(\epsilon_d(R)\Big),\end{aligned} \tag{S16}$$

where $\tau(T_0)$ and $r_g(T_0)$ are the experimentally observed minimum doubling time and protein synthess rate at a temperature $T_0$ respectively, $k$ is the exponent that describes the scaling between the minimum doubling time and the protein synthesis rate, and (from equation S10),

$$\begin{aligned} r_g(T) &= r_g(T_0)\cdot\exp\Big[-a\Big(\frac{1}{T-T_m} - \frac{1}{T_0-T_m}\Big)\Big],\\ \epsilon_m &\sim N(0,1),\\ \epsilon_d(R) &\sim N(R,1).\end{aligned} \tag{S17}$$

with $a$ describing the protein synthesis rate as function of temperature, and the temperature $T_m$ below which all protein synthesis stops. With normally distributed $\epsilon_d$ (for fixed $R$), we find that cells have longer doubling times but occur less frequently when $\epsilon_d$ increases above $R$. Thus, the doubling time is a tradeoff between the speed of duplications and the likelihood of such a cell occurring. In practice, the values of $\epsilon_d$ are bounded because there exists a longest possible doubling time $t_{\max}$ due to an experimentally observed threshold value of $\epsilon_d$ (threshold of ROS concentrations) beyond which cells cannot duplicate and will die. The parameter $R$ is the only free variable in the model, and all other parameters will be constrained next.

**Model parameters** We choose the known temperature $T_0$ to be 5.0 °C, where we have experimentally measured $r_g(T_0) \approx 11.23$ a.u. / day (Supplementary Fig. S30) and the experimentally measured minimum doubling time $\tau(T_0) \approx 2.5$ days (with added GSH: ~25 hours DNA duplication, ~22 hours DNA segregation, ~4 hours in $G2$, and a median G1 duration of ~9 hours (see Supplementary Fig. S22). We used the median G1 duration due to the shape of the distribution to more accurately estimate the minimum time spent in G1 with added GSH). We fitted the power-law type scaling to the minimum doubling time as function of the protein synthesis rate to obtain the exponent $k \approx 0.77$ (Fig. 6b, Pearson correlation-coefficient $\beta = 0.9979$). Only single-cell data was used to determine the actual minimum doubling times (assumed to be equal to the average single-cell doubling time at 30 °C, the average single-cell doubling time with added GSH at 5.0 °C, and the minimum doubling times at 1.0 °C that exclude the duration of G1). As a lower temperature limit $T_m$ for protein synthesis we took $T_m = -10$ °C, motivated by the fact that the yeast growth medium freezes at −2 °C (and thus limits protein synthesis by for example restricting access to nutrients, leading to deprivation of oxygen and limiting movement of (say) ribosomes) and because important enzymes such as Catalase stop functioning at $-6$ °C [18] and other essential enzymes inactivate at 0 °C [19, 20]. It is therefore safe to assume that the temperature below which yeast's protein synthesis stops lies above $-10$ °C. Having fixed $T_m$, we fitted the temperature dependence of the protein synthesis rate using equation S11 from which we find that $a \approx 120.72$ (Fig. S30, Pearson correlation-coefficient $\beta = 0.9954$). Substituting the values of all known parameters yields the model that was used for all simulations,

$$
\begin{aligned}
t(T, R) &\sim (2.5 + \epsilon_m) \cdot \exp\left(\frac{93}{T + 10} - 6.2\right) + 0.09 \cdot \exp\left(\frac{121}{T + 10} - 8 + \epsilon_d(R)\right), \\
\epsilon_m &\sim N(0, 1), \\
\epsilon(R) &\sim N(R, 1).
\end{aligned}
$$

This model describes the single-cell doubling time as function of temperature $T$ and parameter $R$ being the only free parameter of the model describing the intracellular ROS concentrations.
